# ProA and ProB repeat sequences shape 3D genome organization in eukaryotes

**DOI:** 10.1101/2023.10.27.564043

**Authors:** Konstantinn Acen Bonnet, Nicolas Hulo, Raphaël Mourad, Adam Ewing, Olivier Croce, Magali Naville, Nikita Vassetzky, Eric Gilson, Didier Picard, Geneviève Fourel

## Abstract

Genome organization is partially conserved across cell types, yet its DNA-encoded determinants remain incompletely understood. Here we define ProA and ProB repeat sequences (RepSeqs) as two classes of cis-elements that promote A/euchromatin or B/heterochromatin compartment identity. We show that relative ProA/ProB density predicts Hi-C compartment profiles, indicating that compartmental propensity is largely encoded in sequence composition, and point to specific chromatin-based mechanisms underlying these effects. ProA RepSeqs are predominantly Alu elements, whereas ProB RepSeqs comprise young LINE-1s, selected ERVs, AT-rich microsatellites, and satellite repeats. RepSeqs of more indefinite character, including transcriptional enhancers, can switch between ProA and ProB functions to open or close chromatin domains in a context-dependent manner. In cancer, CpG methylation loss disproportionately impacts ProB RepSeqs, weakening the B compartment and thereby contributing to genome unfolding and cancer cell plasticity.

## INTRODUCTION

In differentiated eukaryotic cells, the genome is partitioned into two major nuclear compartments, euchromatin and heterochromatin. These are thought to correspond approximately to A and B compartments, respectively, as defined experimentally by Hi-C, and within which interactions between distant loci are preferentially detected ^1,2^. B is significantly more compact and is unfavorable to gene expression such that the vast majority of expressed genes in differentiated cells are in A. In a browser view, this translates into a succession of A and B domains as indicated by the Hi-C eigenvector profile (Hi-C EV) ^1^, in a characteristic pattern for a given cell type (Fig. 1). The A and B regions may be further divided into subdomains that seemingly constitute the elementary unit of unfolding/refolding of the genome ^3^. In vertebrates, these domains are called topologically associated domains (TADs) and are often delimited by boundary elements associated with DNA-binding factors, and in a high proportion of cases the transcription factor (TF) CTCF ^4,5^. In a living cell, the chromatin fiber is pervasively set in motion due to the action of “active effectors” ^6,7^, in particular cohesin rings that relentlessly extrude DNA, forming chromatin loops that enlarge until the cohesin ring detaches or arrests. This occurs preferentially when a cohesin ring encounters a bound CTCF molecule and is responsible for TADs appearing as triangles with a marked apex in contact maps ^8-10^. Furthermore, pervasive chromatin loop extrusion by cohesins behaves as a force which uncouples contiguous chromatin segments, and globally opposes the formation of both A and B compartment, therefore breathing dynamics into A/ B partitioning ^11-15^.

**Fig. 1:**
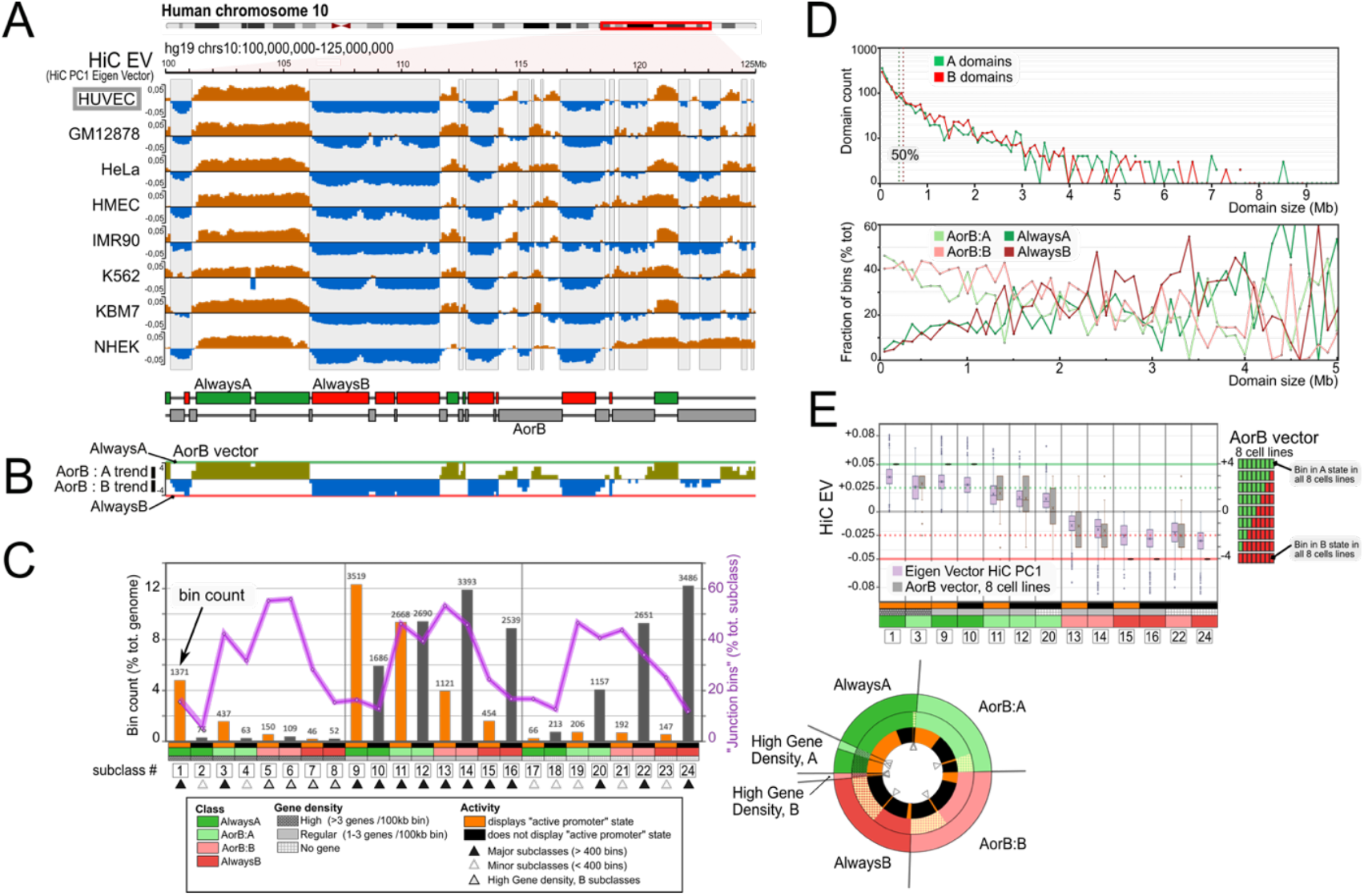
High conservation of the A/B compartment profile points to the existence of pivotal DNA determinants. (A) The eigenvector of the Hi-C PC1 is aligned for eight ENCODE human reference cell lines, over a 25 Mb region of human chromosome HUVEC and NHEK are primary, expanded cells; IMR90 is derived from fetal lung; HMEC and GM12878 are immortalized by SV40 T antigen and EBV, respectively; the others are cancer cell lines. Bottom: Regions that are A (respectively B) in all eight cell lines are indicated as “AlwaysA” green boxes (“AlwaysB” red boxes, respectively). (B) The “AorB vector” function sums the compartment state of the eight cell lines as shown in A. Its value is +4 for AlwaysA regions and -4 for AlwaysB regions, and has intermediate values otherwise. (C) Partitioning of the human genome into 24 subclasses of 100 kb bins according to compartment, gene density, and “activity”. The latter corresponds to the presence of at least one segment of “active promoter state” (ChromHMM state 1 or 2 ^28^). For each subclass, the absolute value of bin counts is shown at the top of histogram bars and as % of the total bin count on the left y-axis; the pink line indicates the fraction of bins in a subclass lying within 200 kb of an A/B transition (“junction bins”; right y-axis). Scheme on the right: Pie representation of the 24 subclasses. (D) Domain size as a function of number of domains with a certain size, shown independently for A domains and B domains (top); domain size as a function of fraction of total bins (bottom). The total population of bins belonging to domains of a certain size is considered. Shown is the fraction of such bins belonging to each of the four bin classes (AorB:A, AorB:B, AlwaysA, AlwaysB). A domain is defined as a group of contiguous bins between two A/B transitions. Dotted line, median values. (E) Boxplot representation of the AorB vector distribution shown for the 13 major bin subclasses alongside with HUVEC Hi-C EV. In order to better take into account genome heterogeneity, we further partitioned the hg19 version of the human genome into 24 subclasses of 100 kb bins according to gene density and whether the H3K4me3 histone mark typically found at active promoters is detected in the bin (“active” or “inactive” bins) (Fig. 1C; Extended Data Fig. 1B; see Resources & Methods section). As anticipated, a majority of active bins are found in A subclasses (Fig. 1C; orange bar), whereas a majority of gene-empty bins are found in B subclasses. The counts for the latter would be even higher if one used more recent versions of the human genome than hg19, which contains limited portions of the (peri)centromeric sequences. That some gene-empty, mostly inactive bins harbor active promoter marks is due to the fact that such marks may be found outside promoters, at potent enhancers in particular. In the following, we will mainly focus on 13 major subclasses, each represented by more than 400 bins (Fig. 1C). The case of AlwaysB bins featuring active promoter marks and/or a high gene density will be treated separately (Extended Data Figs. 2 and 3). These largely overlap with so-called “gene complexes”, which emerged by repeated segmental duplications.

While the genome itself is generally thought to be essentially identical between all or almost all cells of an organism and throughout development, A/B compartmentalization is largely undetectable in the early embryo and in the early G1 phase just after mitosis. Compartments arise gradually in both cases alongside with the emergence and reconstitution, respectively, of the H3K9me3/HP1a -based heterochromatin system ^16-19^, pointing to a key role of heterochromatin and associated coalescence forces in B compartment formation, and more generally in genome compartmentalization ^20,21^, as supported by modeling ^22,23^. The mechanisms responsible for the chromatin states underlying A and B compartments are now well understood. For each of A and B, they consist of a system of histone marks, chromatin modifiers and chromatin-binding factors, with the A and B systems being symmetrical but incompatible (“mutual opposition”) ^24^. Moreover, interwoven feed-forward loops animate each of the A and B systems, enabling a memory of chromatin states at distinct time and physical scales, which is responsible for the apparent stability of A and B states ^24-27^. A/B partitioning is thus governed by a toggle-switch scheme as described for many metastable two-state systems, in which mutual opposition dictates the existence of two alternative states, whereas the relative strength of the underlying interactions dictates the dynamics of alternation between them (Fig. 2). However, what causes A- or B-type chromatin states to be found at a given location in the genome largely remains a conundrum.

**Fig. 2:**
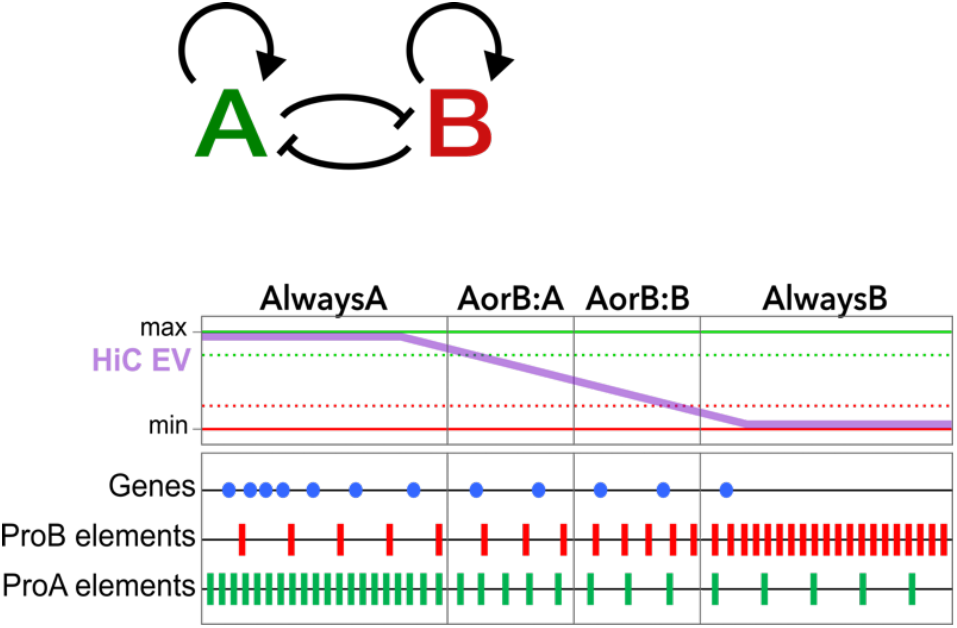
A/B compartment toggle-switch points to ProA and ProB determinants as being enriched in the cognate compartment and having densities correlating with Hi-C EV values. A/B compartmentalization is driven by a toggle-switch principle (upper part) which allows prediction of the distribution of ProA and ProB elements, defined as cis-determinants of A/B partitioning promoting A and B compartments, respectively.

Epigenomics has produced a wealth of data over the last 20 years depicting chromatin fiber composition and long-range interactions throughout the human genome. Epigenomics is unbiased as to the nature of the underlying sequences, however interpretation often turns out to be gene-centered, with more attention being paid to active chromatin states, associated with gene activity, rather than to inactive chromatin states that actually occupy most of the genome ^28,29^. In addition, repeat sequences (RepSeqs) have largely been left out due to technical hurdles. In this article, we revisit with fresh eyes a set of epigenomic data in an attempt to identify general rules and key players responsible for human genome organization. We first show that genome organization is largely inscribed in the DNA sequence, and then set out to identify the DNA elements involved. We incriminate RepSeqs, providing strong evidence that they are responsible for the formation of a default chromatin landscape largely invariant across cell types, which predetermines some domains for compartments A or B, and dictates the propensity of other regions to shift between the A and B states. In these latter regions, DNase I hypersensitive sites (DHS), which are DNA elements that bind TFs and, in most cases, correspond to the so-called transcriptional ‘enhancers’, hitherto thought to activate gene transcription primarily by directly targeting promoters, also appear to drive chromatin domain opening.

## RESULTS

### Genome organization is largely inscribed in the DNA sequence

Let’s observe the Hi-C EV profile obtained with a 100 kb resolution along a representative 25 Mb region of chromosome 10 (Fig. 1A and Extended Data Fig. 1A). In Fig. 1A, the Hi-C EV is aligned for 8 ENCODE reference cell lines. HUVEC shown at the top is a primary cell “line” (cord blood endothelial cells, expanded in culture), and it will serve as our reference cell in this study. Most striking is the general conservation of the profile. In particular, one can distinguish regions that belong to the same compartment in all eight cell lines and cover a majority of the 25 Mb region. These regions will be referred to as “AlwaysA/AlwaysB” (Fig. 1A, bottom part). Conversely, “AorB” are regions that are either A or B depending on the cell line (Fig. 1A, bottom part), and of those, regions that are A (respectively B) in our reference line HUVEC will be referred to as AorB:A (respectively AorB:B). The left half of the chromosome segment shown in Fig. 1A is enriched in AlwaysA/AlwaysB regions whereas the right half is enriched in AorB regions (Extended Data Fig. 1B). In HUVEC, AlwaysA, AlwaysB, AorB:A, AorB:B regions cover each about 25% of the genome (Fig. 1C). Noteworthy, the absolute values of the Hi-C EV appear lower in the AorB regions when compared to the AlwaysA/AlwaysB regions where it is fairly uniformly high (Fig. 1A and E).

Approximately half of the genes in the human genome are contained within AorB regions (Extended Data Fig. 1C). These regions belong to either the A or B compartment depending on tissue type, developmental stage, or alterations associated with carcinogenesis. Accordingly, AorB regions are enriched in genes involved in evolutionarily specialized yet diverse functions, in contrast to housekeeping genes, which are preferentially enriched in AlwaysA regions, as revealed by Gene Ontology (GO) analysis (Extended Data Figs. 1C and 4). AorB bins are found in approximately equal numbers in two situations: (i) at junctions between A and B domains, where the Hi-C EV curve crosses the value 0 (Fig. 1A and C, pink curve; see Extended Data Figs. 3 and 5 for illustration) and (ii) in relatively small domains that contain mostly AorB bins (Fig. 1A, compare right and left parts of the 25 Mb region; Fig. 1C, pink curve; Fig. 1D; see Extended Data Figs. 6 and 7 for illustration). Thus, whereas AorB regions found at A/B transitions may rightfully appear as the place where A and B influences from adjacent regions confront each other, these observations suggest that AorB regions also represent a specific habitat, favorable for genes whose expression must be strongly regulated, as independently supported by a recent report ^30^.

To be more nuanced than just mentioning AorB, we created an indicator referred to as “AorB vector” (AorBvec) scoring the A or B status for each 100 kb bin in the eight cell lines (Fig. 1B and E). The value of the AorB vector is +4 when the considered bin is A in all eight lines, i.e. AlwaysA, +3 when the considered bin is A in 7 lines and B in 1 line, and so forth down to -4 for AlwaysB. The AorB vector is therefore a sort of probability function that a bin is part of either the A or B compartment in a given cell line, which could be rendered more accurate by including a larger number of cell lines. AorB regions therefore display AorBvec values between -3 and +3. The AorB vector profile is overall strikingly similar to the Hi-C EV profiles of the cell lines and in particular to our reference HUVEC Hi-C EV (Fig. 1A, B, and E), which has several implications. First, AorB regions, which alternate between A and B compartments between cell lines, largely coincide with regions displaying low absolute values of the Hi-C EV, i.e. low apparent contact preference between A and B compartments <u>in a given cell line</u> (“low A character” or “low B character”). AorB regions therefore display high chromatin “plasticity” both within an individual cell and between tissues. AlwaysA and AlwaysB regions instead contribute to the similarity of the Hi-C EV and AorBvec profiles by featuring high absolute values for both parameters, indicative of low plasticity. Second, an AorB:A domain in HUVEC is more likely to be AorB:A than AorB:B in another cell line, especially when it contains genes: there is an “A trend” - and conversely, an AorB:B domain in HUVEC will most often be B in another lineage. This translates into identical signs for Hi-C EV and AorBvec (Fig. 1E) (“directionality”). In conclusion, the striking conservation of the Hi-C EV profiles between cell lines indicates that major determinants of Hi-C EV, and altogether of A/B partitioning, are to be found in the DNA sequence as suggested earlier ^31,32^, determining both compartment directionality and the degree of plasticity.

### The identification of ProA and ProB elements in the human genome points to a key role of repeat sequences in genome organization

#### Identifying cis-determinants of A/B partitioning: ProA and ProB elements

Since genome organization appears largely encoded in the DNA sequence, we decided to identify sequence determinants of A/B compartmentalization. One may think essentially of two types of DNA features: GC content (%GC), and the density of different types of RepSeqs, and genes - which can be viewed as a particular type of repeat sequence. It has long been known that the genome partitions into domains of varying GC content ^33^, which closely coincide with the A and B domains (see Extended Data Figs. 2, 3, 5-7 for illustration), with a higher %GC consistently found in the A compartment. Therefore, GC content is a good predictor of A/B partitioning ^33^. Strikingly, high AT content appears to promote the spreading of distinct, recently described heterochromatin types in prokaryotes ^34,35^, and there is evidence suggesting that this may also be true in eukaryotes ^36-38^. Thus, one reason for the high %AT and, consequently, low %GC in the B compartment might be that it promotes the binding of structural heterochromatin proteins to the chromatin fiber through mechanisms which await characterization. AT-binding factors, including both structural proteins and RNA, as well as remodeler machineries, may be involved in this process ^39,40^. Alternatively, the formation of nucleosome clutches, thought to be stabilized by structural heterochromatin factors, or even the binding of DNA by these proteins, may also be directly or indirectly promoted by sequences with high AT content ^41-43^.

Considering this background knowledge about %GC, we will focus on interspersed repeats to identify *cis-* determinants of A/B partitioning. In other words, we aim to identify what will hereafter be called in the following ProA and ProB DNA elements, defined as elements that locally promote the formation of A and B compartments, respectively. We reasoned that the density of such individual determinants of Hi-C EV should correlate (or anti-correlate) with the Hi-C EV value (Fig. 2). Conversely, knowing such densities should enable one to predict the Hi-C EV profile. We therefore first assessed the copy count in 25 kb bins for each of the 1,395 RepSeq subfamilies of the RepBase library, and classified these subfamilies according to their degree of Spearman correlation with the Hi-C EV in HUVEC (Fig. 3A; Table S1; see Resources & Methods section). We arbitrarily retained families scoring above 0.01 or below - 0.01, for which the calculated correlation value is highly significant (P < 0.01), and called these two sets CorrA RepSeqs and CorrB RepSeqs, containing 250 and 367 subfamilies, respectively (Fig. 3A; Table S1). As expected, considering the procedure by which CorrA and CorrB RepSeq sets were defined, their density profiles approximately follow the Hi-C EV profile (see below and Extended Data Figs. 2, 3, 5-7 for illustration). For each RepBase subfamily, we further assessed the correlation with the HUVEC ChIP signals for H3K9me3 and H3K27me3, which are typifying histone marks for the HP1a -based and Polycomb-based heterochromatin systems, respectively (Table S1). We also assessed the enrichment in A or B compartments, and in AlwaysA or AlwaysB regions (Table S1).

**Fig. 3:**
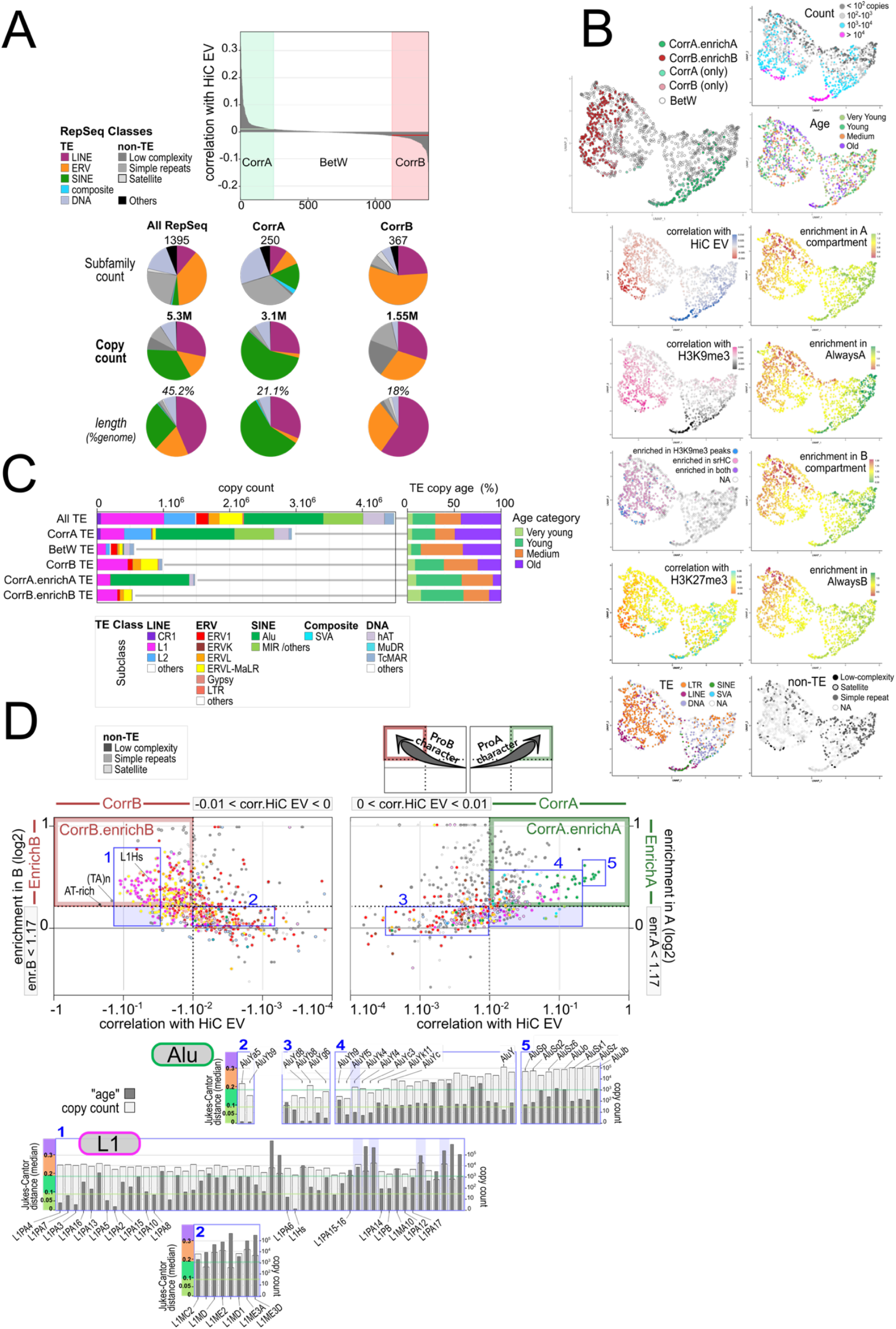
Identifiying ProA and ProB RepSeq in the human genome. (A) Top: All RepSeq subfamilies from RepBase ordered by Spearman correlation with the value of the Hi-C EV obtained with the HUVEC cell line, with bins of 25 kb. Graph showing distribution of CorrA and CorrB RepSeq subsets, with correlation values above 0.01 and below 0.01, respectively. Pie charts at the bottom: Composition of the full complement of RepSeq in the human genome (“All RepSeq”, hg19 version), and of CorrA and CorrB sets, according to RepSeq classes. The category “Others” includes snRNA, srpRNA, tRNA, rRNA, scRNA. Copy count in millions (M), length covered in % of total genome length (hg19). Note that “copy” refers to a segment of continuous homology with a subfamily consensus sequence. TE inserts frequently appear as a suite of more than one such segment (see Resources and Methods section). See Extended Data Fig. 9 for a version of the figure including insert count. (B) UMAP (Uniform Manifold Approximation and Projection) analysis of all RepSeq subfamilies from RepBase, based on the following parameters: age and count, enrichment (in A, in B, in AlwaysA, in AlwaysB); correlation (with Hi-C EV; with H3K9me3 ChIP-seq signal; with H3K27me3 ChIP-seq signal); 100 kb binning; data from HUVEC. Other panels highlight RepSeq classes, and subsets of subfamilies as follows: subsets defined below for panel D using thresholds for correlation with Hi-C EV and/or enrichment in A or B (larger panel at upper left); subsets enriched in biochemically defined fractions of the human genome ^44^, sonication-resistant heterochromatin (srHC), or H3K9me3 ChIP peaks in srHC-free regions. (C,D) Composition and characteristics of RepSeq sets. (D, upper part) Each RepSeq subfamily from RepBase is displayed as a point, with its color indicating TE or non-TE class, on a Euclidean plane according to correlation with Hi-C EV and enrichment in compartments A (right) or B (left). Delimitations of CorrA and CorrB spaces, and EnrichA and EnrichB spaces are indicated on the respective axes. CorrA.enrichA and CorrB.enrichB spaces are highlighted with green and red boxes, respectively. Scheme upper right, ProA and ProB trends (see text). There is a strong overlap between CorrB.enrichB and the L1 family, and between CorrA.enrichA and the Alu family. See Extended Data Fig. 10 for a per class breakdown. (C) TE copy count in RepSeq sets, displayed by subfamily (on the left in absolute counts) or age category (on the right as % of total count in the set considered). All TE, full complement of TE in the human genome. BetW RepSeq, correlation with Hi-C EV between -0.01 and 0.01. The median Jukes-Cantor distance (JCD) calculated as in Ref. ^29^ is used as a proxy for the age of a subfamily. Very young, JCD < 0.09; Young, 0.09 <JCD < 0.18; Medium 0.19 <JCD < 0.3; Old, JCD > 0.3. 0.09 was chosen to demarcate Young and Very Young TEs, because it is an intermediate value between the 0.08 median JCD of the L1PA7 subfamily, the “oldest” among the young L1 to display significant KAP1 binding at a subfamily level, as defined in ES cells ^443^, and that of the older L1PA8A (median JCD 0.10) which does not. (D, lower part) Alu and L1 RepSeq subfamilies falling in the blue boxes noted 1 to 5 in the upper part are displayed as for their copy count and age (JCD median), ordered by increasing correlation with Hi-C EV, from left to right. Purple shading of the background denotes enrichment below 1.17 (in B for box 1, in A for box 4). The youngest L1 subfamilies are among the ones scoring best as CorrB.enrichB, whereas the youngest Alu subfamilies are not (Blue box 2) or little (Blue box 3) enriched in A, and the ones scoring best as CorrA.enrichA (Blue box 5) are among the “less young” Alu (Alu elements and AluJ clades). Notably however, all Alu subfamilies are Young or Very Young. See Extended Data Fig. 10 for a full display of subfamily age by TE class. It has long been recognized that Alu repeats are enriched in euchromatin and L1 repeats in heterochromatin, and that, therefore, the distribution of TEs is heterogeneous throughout the human genome ^20,61-63^. Here, we show more precisely that the vast majority of RepSeqs in the human genome, including both TEs and non-TEs, fall into RepSeq subfamilies whose density significantly correlates either with the A or B character. Thus, the composition of RepSeqs is remarkably distinct between the A and B compartments.

A UMAP analysis of the RepSeq subfamilies using these parameters, along with copy count and estimated evolutionary age (see Resources & Methods section), shows a butterfly-shaped pattern. The subfamilies in the left wing exhibit trends expected for the B compartment (such as high correlation with H3K9me3 and low correlation with Hi-C EV), while those in the right wing show trends typical of the A compartment (Fig. 3B). A significant portion of the RepSeq subfamilies forming the left wing have been independently shown to be enriched in genome fractions biochemically characterized as embedded in heterochromatin structures ^44^ (Fig. 3B). Thus, RepSeq subfamilies can be neatly partitioned into those associated with A character and those associated with B character. The fact that the CorrA and CorrB RepSeq sets each form a distinct cluster on the outer edges of the corresponding wings confirms a strong association (Fig. 3B). Correlation with Hi-C EV is largely conserved between cell lines for RepSeq subfamilies with values below -0.01 or above 0.01 such that the CorrA and CorrB sets would only slightly differ if a different cell line had been selected as a reference (Extended Data Fig. 8; Table S2).

CorrA RepSeq subfamilies primarily consist of transposable elements (TEs) covering more than two-thirds of the TE inserts in the genome, and contain the vast majority of short interspersed elements (SINE) subfamilies (Fig. 3C). CorrB TE RepSeq subfamilies mainly consist of endogenous retroviruses (ERVs) and long interspersed elements (LINE) L1 (Fig. 3C). Together, the CorrA RepSeq and CorrB RepSeq sets account for 90% of the 3.4 million TE inserts in the human genome, covering 37.6% of the hg19 reference genome length, whereas the TE subfamilies absent from both lists (and therefore more evenly distributed throughout the genome) largely consist of relatively young ERV subfamilies with low insert counts (‘BetW’, Fig. 3C-D; Extended Data Figs. 9 and 10; Table S1). The CorrB RepSeq set further includes simple repeats (also called ‘microsatellites’ or ‘STRs’ for short tandem repeats), low complexity sequences, and pericentromeric satellites which together represent about one-third of the CorrB RepSeq copies (Fig. 3A; Extended Data Fig. 10). A salient feature of the CorrB non-TE RepSeqs is their predicted propensity to adopt non-B DNA structures: (i) CorrB simple repeat sequences consist of alternating purine and pyrimidine, with only C (or G) on one strand when not composed solely of A and T, which predicts their ability to adopt a Z-DNA conformation ^45,46^ (Table S1); (ii) such A-rich and AT-rich low complexity sequences can form triplexes with RNAs via Hoogsteen pairing, in particular with the lncRNA KCNQ1OT1, which mediates heterochromatin-dependent repression of its targets ^47,48^. In addition, simple repeats are commonly transcribed, and the high AT skewing of CorrB non-TE RepSeq suggests that they are prone to forming R-loops upon transcription ^49-51^. Simple repeats and satellites are further known to specifically bind a host of TFs ^52-54^. Notably, all types of TEs may also display Z-DNA and R-loop formation potential ^51,55-57^, and L1/Alu elements harbor A-rich tracts capable of forming triplexes, similar to simple repeats ^48^. Satellite sequences building up constitutive heterochromatin at pericentromeres and telomeres (which fall into AlwaysB) are poorly represented in hg19. R-loop formation is a conserved feature of pericentromeric satellites, and is thought to serve a physiological, regulatory role in promoting specific DNA transactions ^57-60^.

#### Knowing RepSeq composition and density suffices to accurately predict Hi-C EV

As anticipated based on our definition of CorrA and CorrB elements, using CorrA and CorrB RepSeq sets together with gene densities in a linear model gives a very good score for predicting the Hi-C EV, and the resulting profile most closely resembles that of AorBvec (Fig. 4B, line 12). This score is better than that obtained using %GC as the only parameter (Fig. 4B, line 15), suggesting that RepSeqs have a contribution to the prediction beyond their GC content. Furthermore, this score is in the same range as those based on chromatin features known to be associated with gene activity and highly enriched in the A compartment (Fig. 4A; Extended Data Fig. 1A). This is the case in particular for the count of DNase I hypersensitive sites (DHS), which has a very strong predictive power as already reported ^64,65^ (Fig. 4B, line 11), or the GROseq signal, which reflects gene transcription activity itself (Fig. 4B, line 5). As expected, in the linear models, the multiplicative coefficients associated with the CorrA RepSeq density, gene density and chromatin features associated with gene activity are positive, whereas the coefficients associated with CorrB RepSeq density are negative, which can be interpreted as the former seemingly promoting A character whereas the latter would seemingly promote B character. Furthermore, combining chromatin features with DNA features improves the prediction score. Intriguingly, in this case the features associated with gene promoter activity, in particular H3K4me3 ChIP peak density, contribute to the models with *negative* coefficients (Fig. 4B, line 1-2), which suggests that while DHS and enhancers are clearly ProA features, gene promoters in an active state behave to some extent as ProB features. In conclusion, the distribution of most RepSeq subfamilies in the human genome is sufficiently heterogeneous with respect to compartmentalization such that one can define two sets of RepSeqs on the basis of correlation with the Hi-C EV, together covering roughly 90% of RepSeq inserts in the genome, and use that information to predict the organization of the genome into A and B compartments.

**Fig. 4:**
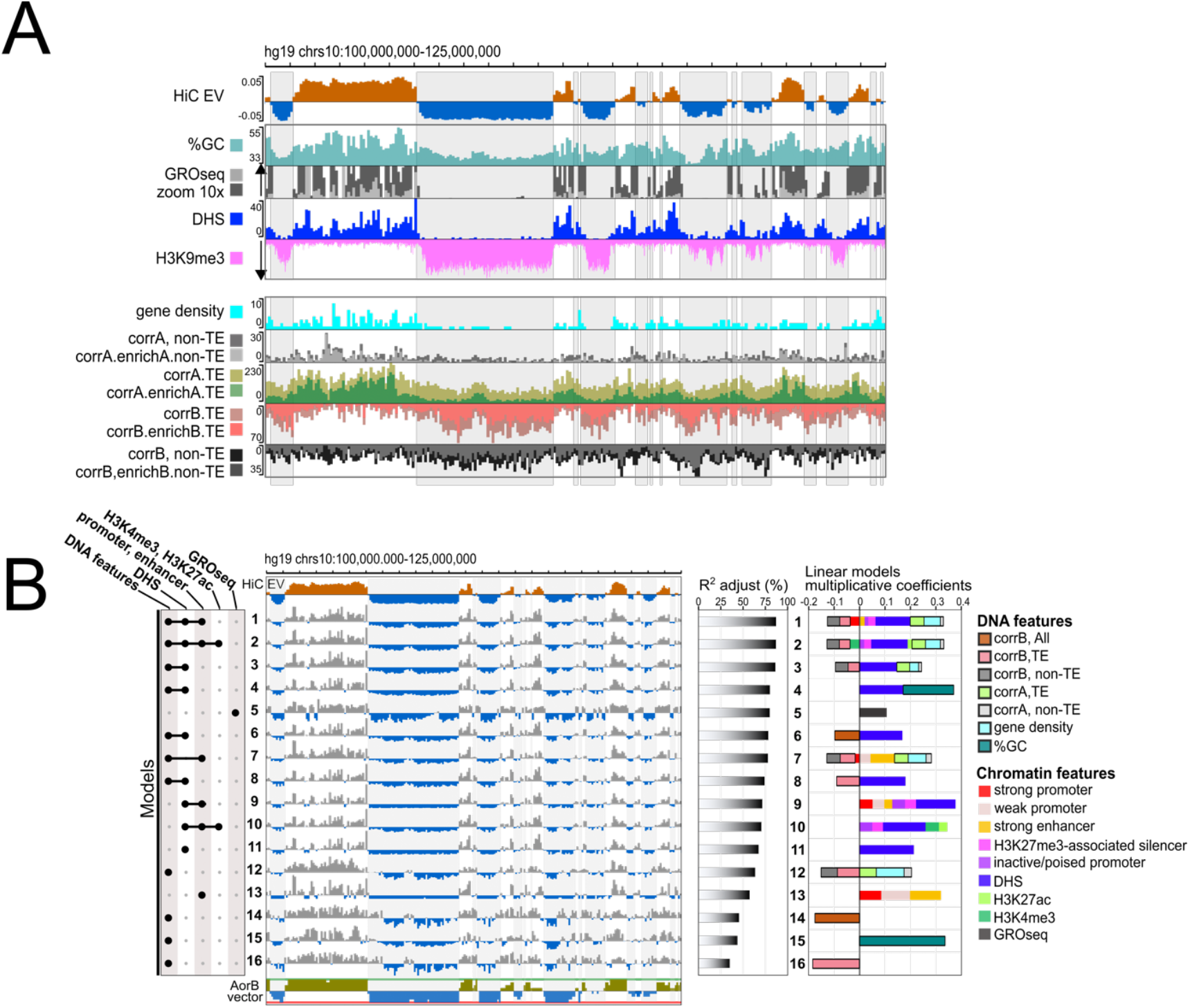
CorrA and CorrB RepSeq local densities predict Hi-C EV. (A) Browser view along the same 25 Mb region of chromosome 10 as in Fig. 1, for features as indicated. GRO-seq, signal summed over 100 kb; H3K9me3, ChIP-seq signal; %GC, averaged over 100 kb; Gene and RepSeq sets, DHS, density over 100 kb. (B) Linear models developed using combinations of DNA features and chromatin features (left) recapitulate to some extent HUVEC Hi-C EV (middle) with a performance indicated by adjusted R^2^ (middle right), expressed as % of the performance obtained by combining all features. Multiplicative coefficients in the linear equation (right) indicate the relative importance of the features used in the prediction (indicated by colors as in the legend, far right), as well as the directionality, positive (resp. negative) coefficients being suggestive of a ProA (resp. ProB) role. Chromatin features (calculated for the HUVEC cell line): ChromHMM segment count, for chromatin states associated with active promoters or enhancers, as well as for inactive/poised, H3K27me3-associated promoters; silencer count, for H3K27me3-associated DHS correlating with gene downregulation ^439^; H3K27ac and H3K4me3, peak density over 100 kb (see Resources and Methods section). Other details are as in panel A.

#### Identifying core sets of ProA and ProB elements: CorrA.enrichA and CorrB.enrichB RepSeqs

Can it, however, be claimed that CorrA and CorrB RepSeqs have a role in establishing the A and B compartments, respectively? ProA and ProB elements are expected not only to locally promote the cognate character within their corresponding compartment, but also to antagonize the opposite character according to the toggle-switch scheme (Fig. 2). Consequently, they should be depleted from the opposite compartment. In other words, ProA and ProB elements should be significantly enriched in the A and B compartments, respectively, which is clearly not the case for all CorrA and CorrB RepSeq subfamilies (Fig. 3D). We therefore applied an additional filter, retaining only those CorrA (or CorrB) RepSeq subfamilies showing enrichment above the empirically selected threshold value of 1.17 in the A compartment (or B compartment) (Fig. 3D; Extended Data Fig. 10). The density profiles of the resulting CorrA.enrichA and CorrB.enrichB subsets follow the Hi-C EV curve much more closely than the complete CorrA and CorrB sets (Fig. 4A). These two “Corr.enrich” RepSeq sets thus clearly meet the expectations for sequences that promote A and B character, respectively.

This does not preclude the possibility that additional RepSeq subfamilies may also play a ProA or ProB role in genome organization. Two types of putative RepSeq profiles would clearly escape our correlation-/enrichment-based selection: (i) “compartment mitigators”, i.e. RepSeqs that promote B character specifically within an A context and vice versa; and (ii) “chameleons”, which would behave as ProA in an A context and as ProB in a B context. Moreover, some RepSeq subfamilies may exert a role through only a subset of their members, without reaching our threshold values when analyzed as a group. The CorrA.enrichA and CorrB.enrichB RepSeqs should therefore be viewed as archetypal ProA and ProB sequences, displaying the most pronounced and context-consistent ProA and ProB characters, respectively. These are the profiles that should be prioritized in future efforts to elucidate the molecular mechanisms by which RepSeqs may contribute to genome compartmentalization.

#### CorrA.enrichA and CorrB.enrichB TEs largely correspond to Alu and SVA subfamilies, and to young L1 and ERV subfamilies, respectively

CorrA.enrichA and CorrB.enrichB TEs represent 1.3 million and 0.3 million inserts, respectively, each covering approximately 12% of the hg19 genome, and are essentially composed of Alu and SVA sequences for ProA, and of ERV and L1 elements for ProB (Fig. 3C; Extended Data Figs. 9 and 10). By comparison with CorrA and CorrB sets, both CorrA.enrichA and CorrB.enrichB RepSeq sets are depleted of relatively old subfamilies, and are therefore correspondingly enriched in relatively young subfamilies (Fig. 3C; Extended Data Fig. 10A). Without being a demonstration, this is consistent with the idea that ProA and ProB TEs might promote the A and B compartments, respectively, via their *cis-*regulatory elements and associated TF binding sites, which degenerate and lose functionality with age (Box 1). Thus, similarly to the *cis*-regulatory elements of genes known as promoters and enhancers, RepSeq harbor *cis*-regulatory elements, which bind both activators and repressors, locally anchoring transcriptional activating and repressing systems, respectively ^66-74^. Repression of young, potentially active TEs appears especially crucial to maintain genome stability by inhibiting both transcription and recombination, chiefly involving HP1a /H3K9me3-based heterochromatin in somatic cells ^4,66,75,76^ (Box 2). Remarkably, the most prominent chromatin marker in the B compartment is precisely the H3K9me3 mark (Fig. 4A). This suggests, without presuming the underlying mechanisms, that somehow H3K9me3 and the associated chromatin state spread from ProB TEs and affect the entire B compartment. Notably, although TE *cis*-regulatory sequences tend to decay and lose functionality over evolutionary time as the TE sequence drifts, heterochromatin marks may still be observed long after a TE has seemingly lost any transcription potential ^66,77^. This suggests ongoing positive selection (Box 1), which we propose reflects selection for ProB function. L1 inserts perfectly illustrate these findings as the youngest L1 subfamilies are enriched among the top RepSeq scores as ProB, with a trend by which scores tend to diminish with age (Fig. 3D, Extended Data Fig. 10A).

Interestingly, the youngest Alu elements score on the ProB half of Fig. 3D without showing marked polarity, whereas only young Alu subfamilies that have significantly drifted score as CorrA.enrichA (Fig. 3D, Extended Data Fig. 10A). This is consistent with a scenario in which new L1 inserts are initially born as strong ProB elements, whereas new Alu inserts may instead initially feature both ProA- and ProB-type potential and gradually lose the latter through sequence drift, eventually being selected against in the B compartment and selected for in the A compartment. This scenario, which we will substantiate with further molecular details in the following sections and Box 2, is reminiscent of early hypotheses by others regarding the positive and/or negative selection of Alu inserts ^61,78^. In conclusion, there is clear evidence that CorrA.enrichA and CorrB.enrichB RepSeqs have been selected for over evolutionary time within their respective compartments.

#### Molecular mechanisms responsible for active and repressed chromatin states can mediate ProA and ProB functions

While the majority of ERV subfamilies score on the B side of Fig. 3D, no clear trends are observed for the ERV1 and HERVK subclasses. Young ERV1 and HERVK subfamilies predominantly exhibit a mixed character, with intermediate values of correlation and compartment enrichment, similar to relatively young Alu elements (Extended Data Fig. 10A). The LTRs of young ERVs are GC- and CpG-rich sequences (Extended Data Fig. 11), and can behave as *bona fide*, potent enhancers of gene expression, as demonstrated both in classical reverse genetics assays and by *in situ* neutralization of individual inserts, with subfamilies displaying distinctive tissue-specificity associated with the binding of tissue-specific TFs ^69,70,79-83^. Young ERV inserts are therefore markedly bipotential, possessing strong enhancer capacity while also being prominent targets for heterochromatin-mediated repression, similar to other young TE inserts, particularly L1s ^70,75,84^. Correspondingly, a large number of ERV LTRs inserts are found associated with either an active or a repressed chromatin state in various tissues ^29^. Notably, while the reported gene *cis*-regulatory activating potential of L1 and Alu elements is less spectacular, the idiosyncratic evolution of any individual TE insert, through acquisition and loss of TF binding sites (Box1), may allow the emergence of elements with high enhancer activity and therefore similar bipotentialities as the LTR of ERV inserts, as indeed observed ^72,73,83,85-87^.

The observation that DNA segments harboring active chromatin states, typically associated with enhancer elements, behave as ProA elements in linear models (Fig. 4B), while strong ProB TEs are characteristically repressed by HP1α/H3K9me3-based heterochromatin, further suggests that the molecular mechanisms responsible for active and repressed chromatin states can mediate ProA and ProB functions, respectively, as proposed earlier ^20,24^. Conversely, the indefinite ProA/ProB character of many ERV subfamilies likely reflects the fact that their members are “convertible RepSeqs,” capable of switching between active and repressed chromatin states, and therefore between ProA and ProB functions, depending on cellular or chromosomal context, potentially acting as “mitigators” or “chameleons”.

#### TEs as composite elements harboring both of ProA and ProB capacities

A dynamic view of molecular events taking place along the chromatin fiber suggests that active and repressed chromatin states constantly confront one another at every TE insert, a phenomenon captured in specific settings ^24,84^. We therefore propose that the majority, if not all, TEs contribute to genome organization through a combination of both ProA and ProB capacities, with the “slider” locked toward ProA or ProB only for the core sets of RepSeqs that predominantly adopt ProA or ProB functions, respectively, across most contexts. These latter RepSeqs will hereafter be referred to as “Constitutive ProA” and “Constitutive ProB” elements, respectively, a category that notably includes a large fraction of CorrA.enrichA and CorrB.enrichB RepSeq subfamilies.

### Compositional genomics reveals general trends for the local densities of ProA and ProB elements, and a marked heterogeneity between functionally similar territories

#### Local densities suggest that relative proportions of ProA and ProB RepSeqs determine proneness toward A or B compartments

By way of extension, the CorrA.enrichA and CorrB.enrichB sets will hereafter be referred to simply as ProA and ProB RepSeq sets, although they in fact represent a subset of archetypal ProA and ProB elements, which have the capacity to perform these functions in a largely constitutive manner. A compositional analysis of the human genome reveals a trend in which the density of both TE and non-TE ProB RepSeqs gradually increases from high-A-character subclasses to high-B-character subclasses, while the reverse is observed for ProA RepSeqs and for chromatin features associated with gene activity, particularly DHSs. AorB subclasses display intermediate profiles (Fig. 5A; see Extended Data Fig. 13A for all 24 subclasses). This global pattern aligns remarkably well with the expectations outlined in Fig. 2.

**Fig. 5:**
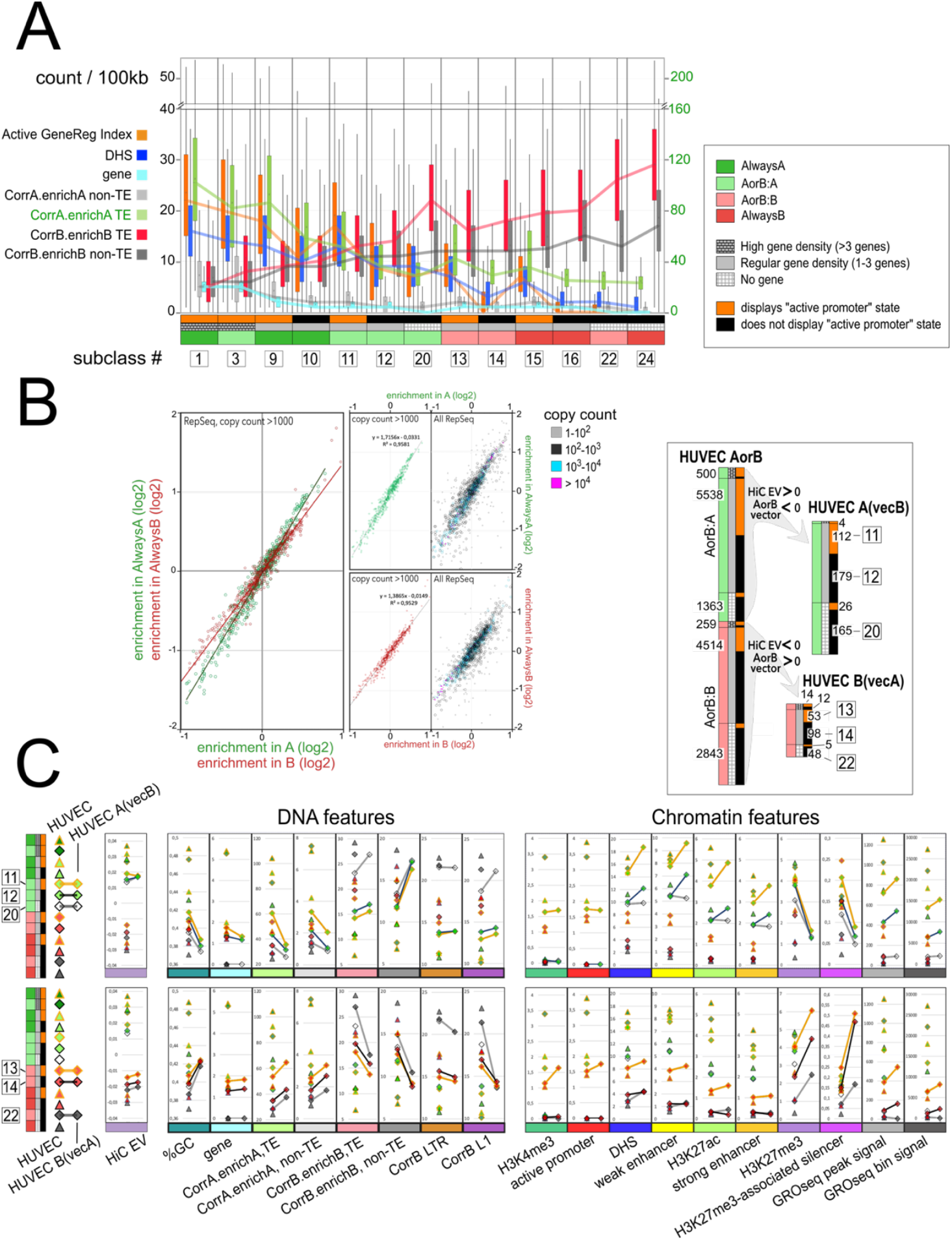
Compositional genomics reveals general trends for the local densities of ProA and ProB elements, and an outlier composition for outlier bins. (A) Boxplot representation of the density of DNA and chromatin features in the 13 major bin subclasses. Active GeneReg Index is a sum calculated from ChromHMM segment counts as follows: 0.5*WeakPromoter + 1.5*StrongPromoter + 0.5*WeakEnhancer + 1.5*StrongEnhancer. Lines connect successive median values. Note the scale is different for CorrA.enrichA TE density, as indicated in green on the right of the graph. For other details, see also legend to Fig. 4. (B) Direct mathematical relationship between enrichment in AlwaysA and A, and AlwaysB and B, for each RepSeq subfamily. Enrichment calculation is not reliable for subfamily counts below 1000 (right panels), which are discarded for the linear regression shown on the left with an overlay of the data points for copy counts >1000 from the central panels. (C) Outlier AorB domains in HUVEC display a chimeric RepSeq composition. Outlier domains in HUVEC are AorB domains for which Hi-C EV has a sign opposite to AorBvec, meaning that the compartment “color” is different in HUVEC from the majority of the other seven reference cell lines shown in Fig. 1. Only “outstanding outlier” bins with the following characteristics were retained: (i) a marked difference between Hi-C EV and AorBvec (see Resources and Methods section); (ii) not found at an A/B junction (within 100 kb); (iii) part of domains with more than two successive bins showing such characteristics. Upper part, partitioning of AorB domains, left, and of HUVEC A(vecB) and HUVEC B(vecA) outliers, right. Only subclasses containing more than 40 bins are considered in the analysis. Lower part, mean values of Hi-C EV, DNA features, and chromatin features as indicated, calculated for individual bin subclasses and represented with the same symbols shown on the very left. Values are shown for all 13 major bin subclasses in HUVEC, and for the selected outlier subclasses, three HUVEC A(vecB) subclasses and three HUVEC B(vecA) subclasses. The same symbols are used in both cases, connected with a line. For other details, see also legend to Fig. 4B.

In addition, there is a direct relationship between the enrichment value of a ProA RepSeq subfamily in AlwaysA regions and in A regions more generally, and likewise for ProB subfamilies in AlwaysB and in B regions (Fig. 5B). Thus, the marked functional character of Always regions with respect to compartmentalization coincides with a marked character in their ProA and ProB RepSeq composition.

Because the absolute counts of ProA TEs along these trend lines are much higher than those of ProB RepSeqs, we divide ProA TE counts by 3 (referred to as ProA* counts) in subsequent analyses and representations to facilitate comparison. Using this arbitrary adjustment to compute a ProA/ProB RepSeq ratio, the ProA “weight” has a median value of 3.6 (5.2 when only TEs are considered) in the AlwaysA subclasses with the strongest A character – representing an approximately 20-fold increase relative to its value in the AlwaysB subclasses with the strongest B character. The mirror image holds for the ProB weight, which reaches a median value of 8.4 (5.4 when only TEs are considered) in the AlwaysB subclass composed of inactive, gene-devoid bins (Extended Data Fig. 13A). Together, these reciprocal trends illustrate a robust polarity in the distribution of ProA and ProB elements across compartments. Interestingly, when using ProA* and ProB counts, the total RepSeq density remains approximately constant across subclasses (Extended Data Fig. 13A, middle panel).

These observations are consistent with a model in which the relative proportions of ProA and ProB RepSeqs play a crucial role in determining the propensity of a chromosomal segment to adopt either an A or a B configuration. However, it is important to note that these patterns represent general trends: each subclass displays a broad range of values for each feature, indicating substantial heterogeneity between bins within a subclass and notable overlap between subclasses (Fig. 5; Extended Data Fig. 13C). A similar pattern emerges when %GC classes are examined (Extended Data Fig. 13B).

The fact that both absolute and relative ProA and ProB densities follow these trends supports a model in which ProA and ProB elements operate as two systems. In this model, functionally interchangeable elements cooperate over distance within each system, with greater efficiency when in closer proximity, and collectively follow the toggle-switch logic shown in Fig. 2. Cooperation among ProB elements may occur through previously described principles ^20^, forming a network that collectively promotes HP1α-based heterochromatin establishment and cohesion within the nucleosomal fiber ^21,88-90^. ProA elements, in turn, are envisioned to cooperate in antagonizing the ProB system.

#### Outlier regulatory patterns correlate with outlier RepSeq compositions

Is the heterogeneity observed between bins within a chromatin subclass compatible with the notion of RepSeqs as key players in genome organization, or might it even have a specific role? To address this question, we analyzed AorB bins that behave as outliers in HUVEC relative to seven additional reference cell lines (i.e., HUVEC.A(vecB) bins that score as A in HUVEC but display a B trend in most other cell lines, and conversely HUVEC.B(vecA) bins). Only “outstanding outliers” in HUVEC - those displaying the largest discrepancies between Hi-C EV and AorBvec values - were considered (see Resources and Methods).

HUVEC.A(vecB) and HUVEC.B(vecA) regions show strikingly low and high levels of Polycomb-associated facultative chromatin markers, respectively, while otherwise displaying the expected average levels of chromatin features associated with gene activity for A and B bins (Fig. 5C; Extended Data Figs. 5, 6). However, they also reveal a remarkable chimeric RepSeq composition. For HUVEC.A(vecB), the %GC and overall RepSeq composition generally follow the trends of the corresponding HUVEC B subclasses, consistent with these regions belonging to the B compartment in a majority of cell lines. The exception lies in ProB RepSeqs: ProB TE density approaches the lower levels seen in A subclasses, whereas ProB non-TE density tends to be particularly high, matching the levels observed in AlwaysB regions (Fig. 5C; Extended Data Fig. 7).

The mirror image is observed for HUVEC.B(vecA) regions, which largely display a typical A-like composition with elevated ProA RepSeq densities, while simultaneously exhibiting ProB RepSeq densities only slightly lower, on average, than those of the corresponding HUVEC B subclasses (Fig. 5C; Extended Data Fig. 5). Notably, many HUVEC.A(vecB) and HUVEC.B(vecA) regions exhibit intermediate values for the various DNA features, a pattern consistent with the observation that these regions do not show extended stretches with a clear A or B trend, as reflected by the low absolute values of the Hi-C EV and AorB vector (see Extended Data Fig. 6 for illustration).

Altogether, bins that are outliers in HUVEC with respect to compartmentalization, i.e. scoring B in HUVEC but A in most other cell lines or vice versa, also emerge as outliers in terms of RepSeq composition. The broad dispersion of RepSeq composition observed within each subclass may therefore play a role in enabling the local emergence of diverse regulatory subfunctionalities.

#### Gene complexes as extreme cases of outlier RepSeq composition

A distinctive type of outlier region consists of gene complexes, arising from segmental duplications. These complexes represent an impressive 13.4% of the human genome ^91^ and harbor families of genes, often at high densities, that may have diverged functionally and in their regulation. Gene complexes predominantly fall within B subclasses 5-8 and 13-16, and accordingly display a high density of ProB elements (Extended Data Figs. 2, 3, 13A).

Among these, KZFP complexes stand out with an exceptionally chimeric composition, displaying very high densities of both ProA and ProB RepSeqs - a feature strikingly associated with strong, constitutive transcriptional activity (Extended Data Figs. 3,13C). In this context, HP1α-based heterochromatin likely exhibits dual functionality, acting both to establish a robust regulatory environment and to prevent recombination between closely related sequences, as previously described for Drosophila chromosome 4 ^92,93^. HOX complexes (particularly HOXA and HOXB), which score as AlwaysA in our analysis, represent a notable exception: they exhibit an almost complete absence of TEs, a feature noted early ^61^, together with a high GC content (illustrated for HOXA in Extended Data Fig. 14). This composition correlates with prominent, evolutionarily conserved control by Polycomb heterochromatin. Notably, their immediate genomic neighborhoods display the typical combinatorial composition of other gene complexes, with relatively high densities of both ProA and ProB RepSeqs.

Gene complexes thus provide extreme examples of outlier regulatory behavior, in both transcription and recombination, that correlate with atypical RepSeq compositions, ranging from unusually high densities of repeats to an almost complete absence of them.

#### Heterogeneity in RepSeq composition within chromatin subclasses as a reflection of the fractal organization of the genome

Alternatively, the heterogeneity in RepSeq composition may also reflect fundamental physical principles that govern genome organization. Although RepSeq- and gene-density-based linear models capture the overall shape of the Hi-C EV curve, the resulting curves are noticeably more jagged than the remarkably smooth Hi-C EV profile (Fig. 4B). This observation is consistent with the idea of long-distance cooperation among elements within the ProA and ProB subsystems, extending far beyond the 100-kb binning length used here. For instance, the smoothing of the Hi-C EV curve across a small A domain that is highly enriched in DHSs can be understood as the result of cooperation among surrounding B domains, which collectively tend to impose a B-like chromatin configuration (Extended Data Fig. 6). Conversely, neighboring bins that share similar RepSeq compositions may exhibit very different Hi-C EV values if one lies near the center of a small B domain while the other is positioned at its border, as illustrated in Extended Data Fig. 5. More broadly, genome organization is known to obey fractal principles, with self-similar domain patterns recurring across scales, as inferred in particular from Hi-C analysis ^1,5,94,95^. A large dispersion in the elementary parameters governing the system measured at a 100-kb resolution is therefore not unexpected, and may reflect features associated with multiscale, fractal-like organization.

In conclusion, we provide strong evidence that the local dosage of ProA and ProB DNA features coarsely dictates the propensity of a chromatin region to adopt an A- or B-type configuration, or to instead display plasticity. This principle should not be viewed as strictly deterministic.

Rather, the considerable heterogeneity observed in ProA and ProB RepSeq densities can, in part, be interpreted as a reflection of the fractal-like, multiscale organization of the genome, in which ProA and ProB elements are envisioned to functionally interact across multiple spatial dimensions in nuclear space, and in a highly dynamic manner.

### DHSs open domains by opposing B compartment

#### DHSs act as an opposing force to B-compartment formation

DHS density emerges as the single ProA chromatin feature with the strongest predictive value in our linear model approach (Fig. 4B). DHSs correspond to DNA segments actively freed from one or two nucleosomes by the interplay of TFs that bind to them and recruit chromatin remodelers, which in turn promote nucleosome sliding or eviction to both induce and sustain DHS existence ^96-102^. DHSs are largely tissue-specific, and approximately two thirds of DHSs in a differentiated somatic cell coincide with active gene enhancers ^103^. Analyses across multiple tissues and cell lines have established an initial compendium of 3.6 million DNA elements actuated as DHS in the human genome ^103^, and more recent work suggests that more than 550,000 DHSs are active specifically in the human cortex ^87,104^. The number of these elements is strikingly in the same range as the total RepSeq count of chromosome arms, as discussed above.

As expected for a feature associated with gene activity, DHS density is on average systematically lower in a chromatin subclass where bins display no active promoter marks, compared with the subclass with similar gene density and chromatin color but displaying activity (Fig. 6A). Moreover, DHS density is generally at a background level in the B compartment except for a minority of regions displaying active marks (Fig. 6A), which are mostly found at A/B junctions (Extended Data Figs. 2, 5, 6). In active, high gene density regions, DHS density tends to increase with domain size, albeit only mildly (Fig. 6B). In a seemingly paradoxical twist, smaller domains in certain A subclasses tend to exhibit higher DHS density. This tendency reaches statistical significance for inactive A bins containing few or no genes (Fig. 6B, subclasses 12 and 20; see Extended Data Fig. 6 for illustration). In addition, both DHS and active enhancer densities are especially high in gene-containing bins that score A in HUVEC but B in a majority of the 7 other cell lines analyzed (Fig. 5C, compare HUVEC.A(vecB) bins and HUVEC A bins; Extended Data Figs. 6 and 7). Altogether, these observations suggest that the factors and events associated with a DHS constitute an opposing force, enabling the formation of an A domain within a “hostile,” B-dominated chromatin environment.

**Fig. 6:**
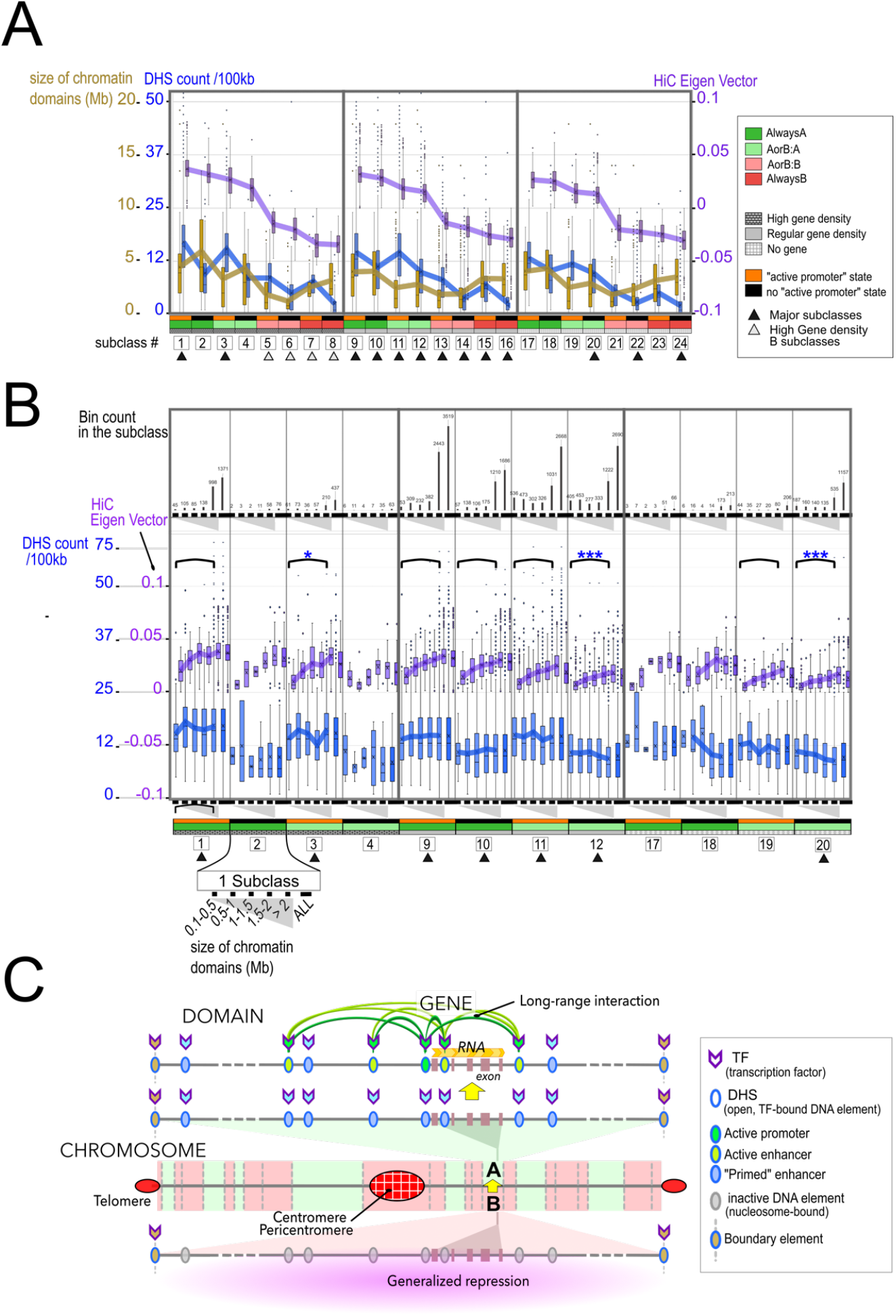
Evidence that DHS open domains. (A) Boxplot showing size of chromatin domains (brown), DHS counts (blue), and Hi-C EV (purple) in the 24 bin subclasses. Note the different scales of the Y axes. Lines connect mean values. (B) Boxplot as in A where each subclass is further subdivided in groups of bins according to the size of the domain in which the bin resides (lower part, zoomed out for subclass 2). Upper part: corresponding bin counts. Lower part: P-value calculated for comparison between domains with sizes comprised between 0.1 and 0.5 Mb with domains above 2Mb; statistical significance for this comparison is reached for subclasses 12 and 20 (p <0.05, three asterisks) with bins that belong to smaller domains containing more DHSs on average than bins in larger domains, and for subclass 3 (p<0.1, one asterisk) in the other direction, i.e., larger domains displaying more DHSs. There are too few bins in subclasses 2,4,17 and 18 to assess the statistical significance (no parenthesis). (C) *DHS occurrence correlates with domain opening*. Schematic representation of a prototype gene, embedded in a prototype domain displaying either A or B state, itself embedded in a prototype chromosome, each displaying general trends. Upper part: In a steady state situation, the gene is transcribed and the domain is part of the A compartment. Promoter activity depends on enhancers which are contained within the chromatin domain (although this is not always the case). The domain may be delimited by clear boundaries, in which case these boundaries also depend on TFs. Although some TFs are specific for a given type of *cis*-element, there is overall a large overlap between TFs, which can perform distinct functions depending on which cofactors they associate with. Lower part: Earlier in development, the domain is part of the B compartment. The DNA displays low accessibility to *trans*-acting factors, including nucleases, and the gene is silent and its regulatory elements are occupied by nucleosomes, with chromatin marks largely indistinguishable from nondescript chromatin in the neighborhood (“generalized repression”, also referred to as “closed configuration”). Middle: When development proceeds and expression of the gene is required for differentiation in a specific cell lineage, for instance, there is a transition stage preceding transcription during which the domain adopts all characteristic features of the A compartment (“open configuration”). In particular, the domain unfolds in nuclear space, and internal contacts within the domain increase while contacts with B domains in the surroundings decrease. There is also a heightened sensitivity to nucleases. This stage strongly correlates with the loss of nucleosomes at enhancers, which is enacted by nucleosome remodeling machineries as recruited by TFs, with a key role for so-called pioneer TFs ^444^. Enhancers then manifest as DHSs, further associated with typical enhancer marks on neighboring nucleosomes, such as H3K4me1, H2A.Z, low levels of H3K27ac, and various additional marks that are thought to arise mostly through the non-targeted activity of enzymes, chromatin modifiers, remodelers, but also Pol II (referred to as “open”or “primed enhancer”) ^119,120,445,446^. These histone modifications are nevertheless believed to redundantly promote DHS formation and prime subsequent conversion to the “active enhancer” state ^119^. Only in a distinct, final stage, coinciding with the onset of gene transcription, chromatin marks typical of active enhancers or active promoters such as high levels of H3K27ac appear at the corresponding DHSs. Enhancer DNA typically shifts from methylated to unmethylated in the DHS/primed enhancer state, as primarily enacted and maintained by TET enzymes, with an intermediate, transient stage where 5hmC is detected. While loss of DNA methylation is generally assumed to aid in the DNA binding of TFs, it strikingly appears not to be an absolute prerequisite for DNA binding of a large number of TFs, even factors known to be methylation-sensitive. Thus, demethylation may occur either before, or concomitantly, or with some delay relative to the appearance of a DHS, the first option being characteristic of sites bound by certain pioneer factors such as Klf4 ^126,214,217,394,447^.

#### DHS emergence accompanies B-to-A domain transitions

Strong support for the notion that DHSs act as an opposing force to B-compartment formation is to be found during development, or by experimental manipulation, in which the shift of domains from a B/closed to an A/open configuration, and relocation away from a heterochromatin-dense compartment, have been reported to coincide with the appearance of DHSs scattered across the domain, in the absence of any gene transcription ^3,105-110^ (illustrated in Fig. 6C). Some of these DHSs will convert to *bona fide* enhancers, associated with typical markers such as high levels of H3K27ac and RNA (eRNA) production, and possibly contacting a target gene promoter ^3,106,108,111-115^. However, this occurs only in a second step, at the same time as the activity of the cognate gene takes off, and generally entails a relay in transcription factors and their cofactors binding the DHSs ^102,116-122^. Notably, the converse is also observed: the targeted closing of enhancers distributed along a domain by a specific transcription factor is associated with concomitant closure of the domain itself ^121,123^. Strikingly similar sequences of events are observed on shorter time scales, for instance for regions that switch between A and B compartments over a circadian cycle ^124-126^, or upon neuronal stimulation and memory encoding ^127^, or during inflammatory priming of domains for rapid reactivation ^128^.

#### Enhancers primarily act through domain opening

Some active enhancers lack an obvious target gene, as they do not exhibit preferential contact with any gene in a given cell type despite strong enrichment for canonical active enhancer features ^83^. Even for well-established enhancer–gene pairs, the determinants of interaction frequency appear elusive ^129^, casting doubt on the view that the primary function of enhancers lies in direct physical contact with promoters.

Consistent with this idea, enhancers are frequently broadly distributed around target genes, a configuration more compatible with a role in chromatin domain opening than with promoter activation through discrete looping interactions ^130^. Thus, accumulating evidence supports DHS/enhancers as primarily mediating domain opening, enabling transcription but also recombination and initiation of DNA replication ^131,132^, and conceivably any process that is inhibited by HP1a-based heterochromatin in the B compartment. This function of active enhancers may be described as “generalized activation,” in reference to the “generalized repression” observed in the B compartment, also referred to as “silencing,” which is a hallmark of heterochromatin action. According to this perspective, the ability of enhancers to directly regulate gene transcription comes second.

#### Long-range enhancer cooperation drives anti-silencing

Intriguingly, it was reported that perturbing a single enhancer can substantially reduce target gene expression during B-to-A transitions, whereas active enhancers generally display higher redundancy in steady-state conditions ^130^. Clustered DHSs have been observed co-actuating at a single-molecule level ^133,134^, indicating potential cooperation among distant DHS/enhancers. The widespread distribution of enhancers across domains and their essential role in chromatin state transitions further support the idea that DHS/enhancers cooperate over long distances to counteract the formation of B chromatin, promoting chromatin opening and A domain establishment. This remarkably parallels the cooperation among ProA RepSeqs, and also among ProB RepSeqs, as discussed above.

DHS/enhancers thus appear to behave as inducible ProA elements, acting to open domains by cooperatively opposing the forces responsible for B compartment establishment and likely requiring continuous action, though not constant binding by TFs. This anti-silencing role is likely amplified when a basic DHS evolves into a classical active enhancer, as supported by extensive evidence ^135^. As a consequence, the well-documented cooperative interactions among enhancers within a domain, especially the synergy between distant enhancers ^136-138^, may partly result from their cooperation in opening domains via anti-silencing, rather than by direct activation of gene promoters. In the Discussion, we will propose molecular mechanisms by which ProA elements counteract the establishment and maintenance of the B compartment, integrating our findings with existing literature.

### DHS/enhancers are of RepSeq origin, switch from a ProB to a ProA configuration upon induction by TFs, and can globally unfold the genome

#### DHS/enhancers are of RepSeq origin

A growing body of literature shows that, in all tissues where this has been investigated, a significant proportion of DHS/enhancers active in a given cell type are in fact RepSeqs ^29,139-142^. Mounting evidence further suggests that all DHS/enhancers originate from RepSeq, deriving either from TE or non-TE sequences. Thus, recently evolved enhancers, which are specific to an organism or a particular cell type, mostly coincide with young TEs, which may become essentially undetectable across evolutionary time, due to sequence drift and possibly accelerated evolution associated with selection (see Box 1) ^52,85,87,143-148^. Furthermore, strong evidence suggests that gene promoters also primarily originate from RepSeqs, as promoters are thought to derive predominantly from enhancers or TEs ^69,149-151^.

#### DHS/enhancers are dual-function elements that switch from a ProB to a ProA state upon induction

In cases where the DHS/enhancer considered is a very young TE, typically a human- or primate-specific TE, actuation of the DHS/enhancer in the course of development or differentiation actually consists in a clear switch from an initial, H3K9me3-associated ProB chromatin state, to an active DHS/enhancer chromatin state ^69^. Interestingly, integrating switches from H3K9me3 to active DHS/enhancer state on a genome-wide scale allows a compelling prediction of cell fate trajectories ^109^.

However, most DHS/enhancer elements exhibit a nondescript, seemingly neutral chromatin state prior to induction, blending into the landscape ^66^. We surmise that a large fraction of DHS/enhancers nonetheless perform a typical ProB function prior to induction. There are at least three possible explanations for why this is not generally apparent in the form of bound repressor TFs and repressive marks:

(i) A conspicuous background of H3K9me3 signal is observed across the entire genome, visible in both bulk ChIP and single-cell analyses, which may obscure smaller peaks. Conversely, it is often challenging to distinguish a clear H3K9me3 ChIP peak over a ProB RepSeq, even in regions exhibiting marked B character (Extended Data Figs. 2, 3, 7);

(ii) Transient binding of TFs, such as Krüppel-associated box (KRAB) zinc-finger proteins (KZFPs), may evade detection by ChIP while being sufficient to sustain heterochromatin assembly over a region via discrete events of nucleation, due to memory effects within the heterochromatin system of a differentiated cell ^152^ (Box 2). Notably, even prototypical ProB elements, which typically exhibit both H3K9me3 and KZFP ChIP signals, do not appear as DHSs (as illustrated with ZNF274 in Extended Data Fig. 3). This phenomenon can be explained by the repressive chromatin configuration, or alternatively by solitary TF binding at an inter-nucleosomal linker ^153,154^;

(iii) All types of heterochromatin require histone deacetylation for spreading. Targeted histone deacetylation promotes local heterochromatin binding to the chromatin fiber without requiring a dedicated recruitment apparatus, as initially observed in SIR complex-mediated heterochromatin formation in the budding yeast *Saccharomyces cerevisiae* (*S. cerevisiae*) ^155^. It may likewise explain why gene promoters also paradoxically promote the B compartment, as inferred above (Fig. 4B), insofar as active promoters toggle between high and low acetylation states that accompany Pol II transcriptional bursts and intervening inactive periods ^156-158^.

Recent studies have characterized cis-regulatory elements with gene repressive function, in steady-state conditions and specific cell lines or lineages, also exhibiting an absence of notable histone marks ^123,154,159,160^. These silencers bind repressor transcription factors and cofactors, particularly HDACs, display decreased DNA accessibility, and are predominantly located in the B compartment ^123,154,160-162^. Notably, these silencers can convert to an enhancer state in other cell lines or upon induction. These findings provide independent evidence that some DHS/enhancers, in their inactive state, actually perform a ProB function. Altogether, ProA and ProB RepSeqs, along with DHS/enhancers, must therefore be considered part of a single system of elements. In this system, two core sets of RepSeqs, characterized by marked ProA and ProB character and primarily composed of Alu/SVA and L1/ERV elements, function constitutively as ProA and ProB elements, respectively, while DHS/enhancers represent RepSeqs or their derivatives that can exist in either a ProA or a ProB state depending on the cellular context, and will thus be referred to as “convertible RepSeqs”.

#### The shift of DHS/enhancers between ProB and ProA states reflects transcription factor/cofactor switching

Independent support for the ProA/ProB bipotentiality of DHS/enhancers, and for their RepSeq origin, comes from the finding that KZFPs are major contributors to interindividual epigenetic variability in mice and human ^84,163-165^. Thus, KZFPs - which constitute, for the most part, pivotal transcription-factor effectors of heterochromatin-mediated RepSeq repression, and therefore of ProB function in general (Box 2) - were also found to be prime trans-determinants of DHS/enhancer activity ^163,164^. This may be interpreted as revealing that, even when a DHS/enhancer is active, it nonetheless carries a concurrent ProB functionality that negatively modulates its ProA function, thereby tuning its ProA output, as supported experimentally (Fig. 7A) ^163,166^. However, there is also evidence that KAP1, the main corepressor of KZFP can switch to a coactivator state upon phosphorylation or loss of sumoylation ^167-171^, potentially promoting a shift from ProB to ProA function.

**Fig. 7:**
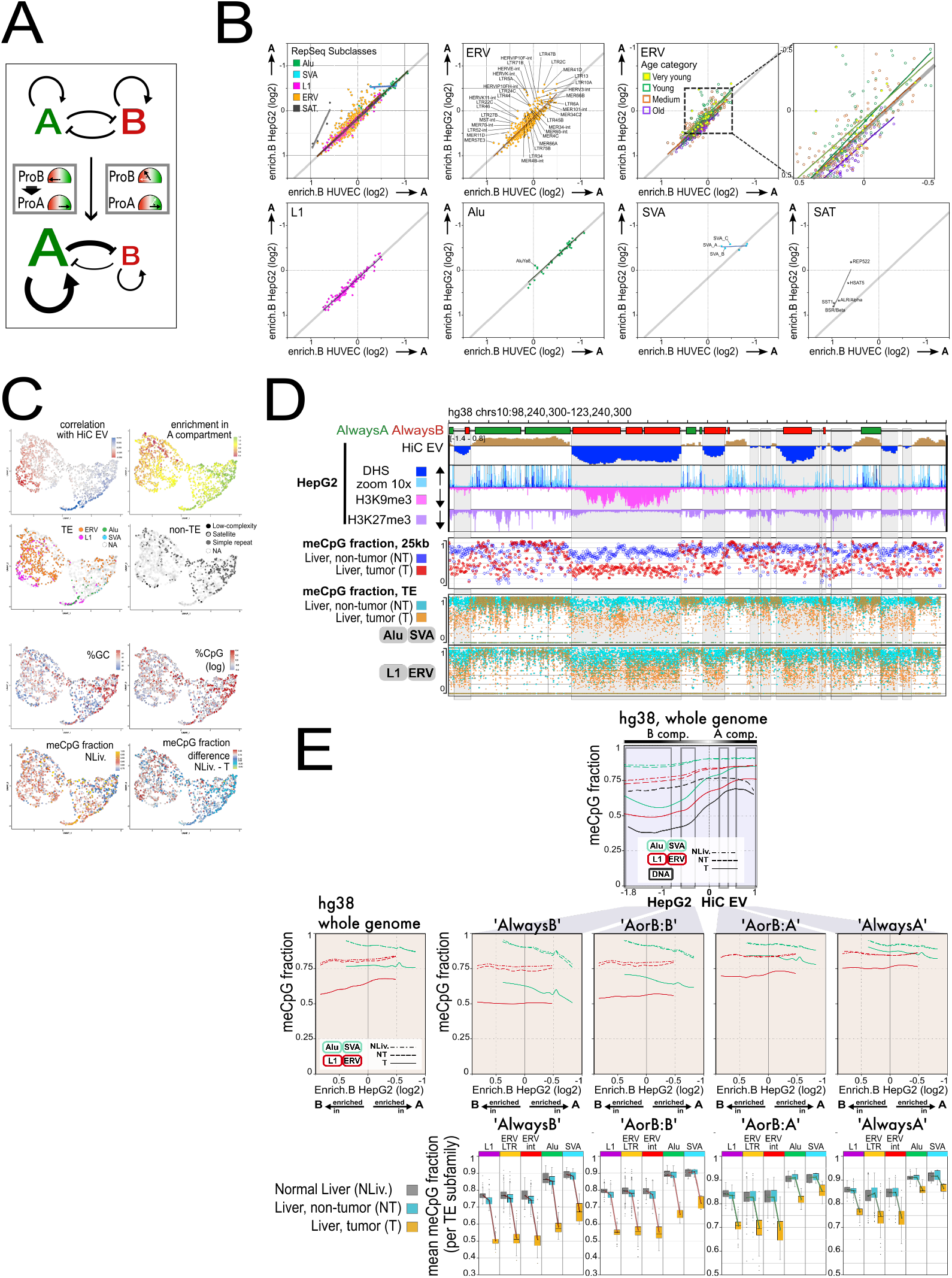
The function of ProB RepSeqs is impaired in cancer, driving genome opening. (A) Modulating or switching RepSeqs from a ProB state to a ProA state, of the type found at active DHS/enhancers, tips the genome-partitioning toggle-switch toward A, opening the genome at all scales. RepSeq inserts should be viewed as composite elements endowed with both ProA and ProB potentials, schematized as a dial with green (ProA) and red (ProB) sectors. Switching multiple RepSeqs located along a chromatin segment from a ProB state, typically marked by H3K9me3, to a ProA state characteristic of active DHS/enhancers, or even simply increasing the ProA output of a set of RepSeqs and/or reducing the ProB capacity of others, tips the toggle-switch toward A, unfolding the segment. This principle applies at every scale of genome organization, ultimately yielding global genome opening. (B) A subset of young ERVs is more enriched in the A compartment in liver cancer, consistent with a shift in their ProB/ProA balance toward ProA. Each RepSeq subfamily (subclasses as indicated) is plotted according to the mean B-compartment enrichment in HUVEC and HepG2 cells. A thick grey line (slope = 1) denotes the approximate trend for Alu, SVA and L1 combined, slightly offset from the diagonal. Only subfamilies with >200 copies are shown. All five subclasses are displayed together in the top-left panel, and separately in the remaining panels. Two upper-right panels: ERV subfamilies are colored by median age (as in Fig. 3), with a magnified central field; age-dependent trend lines are shown with matching colors. Labels are provided only for subfamilies deviating by >0.2 (absolute log2) from the grey trend line toward A - i.e., losing ProB and gaining ProA character in HepG2 relative to HUVEC. (C) Altered DNA methylation of RepSeqs in cancer. UMAP analysis of all RepSeq subfamilies (RepBase), as in Fig. 3B, computed using enrichment (in A, B, AlwaysA, AlwaysB); correlations with Hi-C EV and H3K9me3/H3K27me3 ChIP-seq signal, and RepSeq subfamilies age and copy-number categories. HUVEC data, 100-kb binning. Upper panels: RepSeq families highlighted. Coloring corresponds to compartment enrichment or correlation with Hi-C EV (as in Fig. 3B). Lower panels: Coloring corresponds to mean %GC or %CpG (log10), or mean meCpG fraction in normal liver, or meCpG loss in tumor, calculated as (NLiv − T). Negative values indicate increased methylation in tumor. Most RepSeq subfamilies gaining methylation in tumor are GC-rich simple repeats with low physiological methylation, typically located at CpG islands (see also Ext. Fig. 10B). (D) CpG methylation loss in cancer predominantly occurs in the B compartment, affecting both total DNA and RepSeqs. Browser view of a 25-Mb region of chromosome 10 (as in Fig. 1). The “AlwaysA/AlwaysB” track (same as Fig. 1A) highlights segments consistently embedded in the A or B compartment across eight cell lines. Epigenomic tracks: chromatin features (as indicated) in HepG2. meCpG fraction, 25 kb: nanopore-derived ratio of methylated to total CpGs per 25-kb bin in tumor (T) and adjacent non-tumor (NT) tissue. meCpG fraction, TE: nanopore-derived meCpG fraction for individual TE copies. Points denote Alu/SVA (upper track) and L1/ERV (lower track) copies. (E) TE methylation and its loss in cancer are largely determined by chromatin context. Upper panel: Regression curves generated by plotting each TE copy by its HepG2 Hi-C EV value and its meCpG fraction in T, NT and NLiv. Curve for total DNA (black), derived as in Extended Fig. 15A, is shown for reference. Middle row: Same analysis, except the x-axis is mean B-compartment enrichment of the TE subfamily in HepG2. Curves are shown for the whole genome (left) or for four chromatin classes: ‘AlwaysB’ (EV < −0.8), ‘AorB:B’ (−0.6 < EV < −0.3), ‘AorB:A’ (0.2 < EV < 0.4), ‘AlwaysA’ (EV > 0.55). B-enrichment increases from right to left, placing A-enriched subfamilies to the right of 0. Points indicating the mean meCpG fraction for each subfamily appear in Extended Data Fig. 16A. Bottom row: Boxplots of mean meCpG fraction per subfamily in T, NT and NLiv for L1, ERV, Alu and SVA, in the same four chromatin classes. ERV-LTR and ERV-internal (int) sequences analyzed separately. Only subfamilies with >30 copies in the region considered are included. Extended Fig. 16C presents the same analysis stratified by age.

This notion is further supported by the observation that a minor subset of KZFPs are primarily recognized as transcriptional activators ^77^.

More broadly, most TFs can act in either activating or repressing modes through distinct cofactor partnerships ^123,153,161,172,173^. The effect of TF binding at a given site is therefore the net outcome of the balance between activating and repressing cofactors recruited locally ^172,174^. A classical example is provided by CREB1, SRF, and ELK1, which bind immediate early genes (IEG) enhancers even prior to stimulation ^175,176^. IEG loci, located in AlwaysA regions, are highly accessible (Extended Data Fig. 5) and their promoters carry both active and repressive marks, in a ready-to-go configuration. Upon MAPK- and PKA-dependent phosphorylation, CREB1/SRF/ELK1 switch from recruiting predominantly HDACs/LSD1-associated corepressors to recruiting CBP/p300 HATs, thereby converting key enhancers from repressive to activating and enabling rapid IEG induction ^175,177,178^.

In conclusion, TFs can mediate activating or repressive outputs depending on their cofactor partnerships. TF combinations modulate the behavior of convertible RepSeqs, collectively imposing predominantly either a ProA/enhancer or a ProB configuration on a given element. Transitions between these states may arise from shifts in the predominant cofactors, but more often also involve changes in TF composition itself, as supported by extensive evidence ^173,179^.

#### Viruses drive global genome opening by switching or tuning selected sets of DHS/enhancers towards a ProA state

Viruses massively reprogram cells, a phenomenon intensely studied for decades. We revisited this process for selected viruses to gain additional insights into general principles of genome organization. Cell infection by viruses has been reported to drive subsets of TE inserts - most commonly ERV LTRs - into a transition from a repressed chromatin state to an active enhancer or promoter state, in which they stably persist, accompanied by loss of DNA methylation and, for a subset, active transcription ^180,181^. This is particularly evident for Epstein–Barr Virus (EBV) infection of resting B cells or epithelial cells ^181-184^, where numerous ERV subfamilies of all ages contain members that are induced or over-activated, even though only a minority of inserts within each subfamily are affected (Extended Data Fig. 12B). These induced LTRs are indistinguishable from conventional host cell enhancers except that they rely on the collaboration between host tissue-specific TFs, and virus-encoded TFs or cofactors (“viral transactivators”) ^185,186^. Induced LTRs may also interact in trans with non-integrated EBV episomes ^183,185^. Such neo-enhancers appear to switch entire domains from B to A ^183,184^, mirroring the behavior of endogenous enhancers as described above (Fig. 6C; Box 3).

EBV infection of B cells triggers their transformation from quiescence to a state resembling activated, self-renewing B cells, characterized by a globally decompacted genome and enlarged nuclei ^184,187-189^. The TFs NF-κB and MYC, together with MAPK signaling, have been implicated in both situations, with MYC being directly upregulated by viral transactivators, whereas NF-κB is activated by an EBV-encoded membrane protein ^190^. A plausible explanation for such widespread genome unfolding is that converting large numbers of RepSeqs genome-wide into DHS/enhancer states shifts the ProA/ProB equilibrium, and thus the A/B equilibrium, toward A, thereby flipping the genome-partitioning toggle switch (Fig. 7A). Thus, genome-wide chromatin opening would arise from a cascading collapse of cooperative interactions within the ProB system, across all scales. This interpretation is strongly supported by experimental evidence in the mouse early embryo: switching only a limited number of RepSeqs from a ProB to a ProA state, whether or not accompanied by transcriptional activation, was sufficient to globally decompact the genome ^191^, whereas strengthening the ProB state of these same elements had the opposite effect ^191^. This provides compelling evidence that tuning the ProA/ProB state of RepSeqs constitutes a general mechanism by which cells adjust genome folding at all scales. A comparable global opening of the host genome has been observed upon infection by many unrelated viruses, and also upon viral reactivation from latency. This opening is associated with exit from cell quiescence, which appears necessary for productive viral replication ^184^. Viruses deploy numerous strategies, often simultaneously, to tilt the chromatin toggle-switch toward A. Small DNA tumor viruses - including simian virus 40 (SV40), polyomaviruses, human papillomaviruses (HPV), and the hepatitis B and C viruses (HBV, HCV) - rapidly drive p53 degradation upon infection ^192^, and p53 is a pivotal TF for the repression of RepSeqs ^193-197^. Its inactivation also prevents the p53-driven stress response typically triggered by viral infection ^192^. Some viruses (e.g., HTLV-I, HBV) express “promiscuous transactivators” that potentiate the activity of specific host TFs and chromatin modifiers acting at active enhancers or promoters ^198-201^. EBV EBNA3C similarly converts CtBP from a corepressor into a coactivator ^186^. Still other viruses interfere with transcription, exemplified by influenza A virus (IAV), which inhibits transcription termination ^202^. The resulting readthrough transcription across extended regions exerts intrinsic ProA/anti-silencing effects, switching entire domains from B to A ^202^.

Notably, heterochromatin functions primarily as a virus-restriction system, silencing nucleic acids recognized as invasive. Accordingly, viruses have evolved antagonistic strategies that promote the degradation of heterochromatin components, thereby globally weakening ProB function as a collateral effect. Targets include Smc5/6 (degraded by HBV), the HUSH complex (targeted by HIV-2 and HSV-1), and components of PML bodies, that foster heterochromatinization ^203-206^, notably SP100 and PML itself (targeted by HSV-1 ^207^), as well as ATRX/DAXX, targeted by adenovirus ^203^. While some viral genome-opening mechanisms persist throughout infection, others are transient. For example, both herpesviruses and papillomaviruses transiently induce the cellular TF DUX4, which shifts hundreds of ERVs and Alu repeats, toward a ProA state ^208^. Remarkably, this mirrors the global genome decompaction driven by DUX4 during early embryogenesis ^191,209,210^.

### CpG methylation modulates both ProA and ProB function

#### CpG methylation opposes the ProA capacity of DHS/enhancers and DMRs, possibly through nucleosome stabilization

The human genome of a healthy, somatic, differentiated cell typically exhibits a relatively homogeneous level of cytosine methylation at CpG dinucleotides (meCpG) (∼70% in liver) (Extended Data Figs. 2, 3, 5-7 and 15A (upper panel)) ^211^. The B compartment, however, exhibits a subtle yet clear hypomethylation compared to the A compartment (Extended Data Figs. 2, 3, 5-7; 15A), a phenomenon that emerges at the earliest stages of development ^211^.

A highly localized loss of meCpG accompanies the emergence of a DHS/enhancer, which manifests as a cluster of differentially methylated CpG sites (Differentially Methylated Region, DMR) (Box 3; Fig. 6C). This loss of methylation arises primarily from the action of ten-eleven translocation (TET) methylcytosine dioxygenases, which are recruited by DNA-bound TFs ^212,213^. While the loss of meCpG may precede the onset of a DHS, especially when a TET-recruiting pioneer TF such as Klf4 is involved, it may also occur after DHS formation ^213,214^, such that the two processes are not strictly coupled.

Strikingly, within a given tissue such as the human motor cortex, approximately tenfold more DMRs are detected than DHS/enhancers ^87^, suggesting functions beyond serving as intermediates in DHS actuation. DMRs lacking DHS overlap may function as ProA units, contributing to domain opening in a manner similar to DHS/enhancers, albeit more weakly (Box 3). Such DMRs frequently overlap RepSeqs from all TE classes, even more so than DHS/enhancers, and tend to be less cell-type-specific ^87^.

While it has long been established that methylated CpG dinucleotides can be recognized as an epigenetic tag by a variety of chromatin factors and can also impede the DNA binding of certain transcription factors, it is increasingly clear that CpG methylation exerts additional effects through other mechanisms ^215-217^. High nucleosome occupancy is frequently observed at enhancers prior to actuation and appears to be largely determined by their sequence, particularly their GC content ^121,218,219^. All evidence points to CpG methylation of Alu sequences stabilizing embedded nucleosomes, with Alu repeats standing out as the genomic sequences with the highest nucleosome occupancy (Box 2). Therefore, it is reasonable to propose that CpG methylation may also contribute to stabilizing nucleosomes over enhancer sequences in their closed configuration, as well as over DMRs, although this remains to be investigated. In this model, loss of methylation would facilitate, though not be strictly required for, the nucleosome mobilization observed upon DHS/enhancer actuation ^99^, thereby promoting their ProA capacity.

Conversely, the nucleosome stabilization putatively associated with CpG methylation may also support the ProB capacity of both DHS/enhancers and DMRs when they are in a closed state, reminiscent of the mode of action of SCAN-domain–containing zinc-finger proteins (SZFPs). SZFPs are nucleosome binders that prevent their targets, in particular TEs and TE-derived elements, from gaining chromatin accessibility, independently of canonical heterochromatin pathways ^220^.

#### CpG methylation promotes the ProB capacity of young TEs, presumably by promoting HP1-based heterochromatin

CpG methylation has long been recognized for its role in TE repression, notably serving as a stable memory mark within the HP1α/H3K9me3 heterochromatin system ^221,222^ (Box 2). Thus, targeted CpG methylation plays a pivotal role in repressing L1s and ERVs, particularly those belonging to younger subfamilies, whose regulatory regions are characterized by high GC and CpG densities and elevated methylation levels in normal tissues (Extended Data Figs. 11 and 16) ^223^. Partial or complete loss of meCpG at these TEs is associated with disruption of their heterochromatin-associated repressive state and with RNA production from a subset of them (Extended Data Fig. 12) ^75,222,224,225^.

It is notable that the TE subfamilies with the strongest ProB character in our analysis are precisely found among these young, GC-rich L1s and ERVs (Fig. 3D; Extended Data Fig. 10), suggesting that CpG methylation contributes to their ProB capacity, most likely through its established role in promoting HP1-based heterochromatin. Importantly, a less frequently emphasized aspect is that the high GC content of these TEs also makes them prone to assembling Polycomb-type heterochromatin, which manifests specifically when CpG methylation is lost ^226^.

Although age is a major determinant of CpG methylation levels in L1s and ERVs, with younger inserts generally being more methylated, the chromatin context of the insert also appears to have a strong impact (Extended Data Fig. 15B). Specifically, L1s/ERVs tend to exhibit higher methylation levels in regions with higher Hi-C EV values, from AlwaysB toward AlwaysA contexts (Extended Data Fig. 15B-C). While this pattern may appear counterintuitive, given the view that meCpG promotes the ProB function of L1s/ERVs, it is nonetheless consistent with the general hypomethylation of the B compartment relative to the A compartment (Extended Data Fig. 15B-C).

Additionally, the lines appear approximately flat in Extended Data Fig. 15C (L1/ERV panel), indicating that, to a first approximation, CpG methylation is relatively independent of the ProA or ProB character of the subfamily to which the insert belongs, with the exception of a slight increase at the highest B-enrichment values, corresponding to the higher methylation of younger, CorrB.enrichB L1s and ERVs (Extended Data Fig. 16B).

#### CpG methylation promotes the ProA capacity of Alu repeats, presumably by stabilizing nucleosome

Alu/SVA elements display very high levels of CpG methylation (Extended Data Fig. 11C), and appear, on average, only slightly less methylated in the B compartment (Extended Data Fig. 15B), consistent also with the relatively modest tissue-to-tissue variation of Alu methylation ^227^.

As with L1/ERVs, the youngest Alu subfamilies are the most GC- and CpG-rich and the most methylated, but with exceptionally high methylation levels, frequently exceeding 90% in normal liver tissue (Extended Data Fig. 11B-C; Extended Data Fig. 16). This also applies to SVAs, except that SVAs are human-specific composite elements and are therefore all very young (Extended Data Fig. 16). These levels are actively maintained, as Alu repeats essentially lose methylation within days upon dual depletion of DNMT1 and DNMT3B ^228^, as also observed for other repeats.

One difference that is visually apparent in Extended Data Fig. 15C concerns the influence of chromatin context. Whereas the L1/ERV curves are all approximately flat, the Alu/SVA curves are systematically tilted, becoming increasingly so from the AlwaysA curve toward the AlwaysB curve. This indicates that younger Alu repeats, which are progressively more enriched in B (Fig. 3D), escape the general hypomethylation of the B compartment, whereas Alu subfamilies with the highest ProA character display the lowest methylation levels when located in AlwaysB regions, where their meCpG levels approach that of L1/ERVs. Notably, SVA subfamilies, which all show a similar enrichment in the A compartment in HepG2 context (B-enrichment ∼-0.6), also show a decrease in methylation in B, though less pronounced than for Alu repeats, as evidenced by the red lines, especially the AlwaysB line, forming a clear bump at this level (Extended Data Figs. 15C, upper panel; 16A, B).

A plausible explanation for these observations is that Alu methylation is actively opposed in the B compartment, in a manner that scales with their ProA character. Considering that approximately 40% of the roughly one million Alu sequences in the human genome lie within the B compartment, the most straightforward interpretation is that partial loss of Alu meCpG is required to preserve B-compartment integrity. In this view, CpG methylation directly contributes to the intrinsic ProA capacity of Alu repeats. Nucleosomes are markedly stabilized over Alu sequences, and this positioning is highly dependent on their methylation, with even a mild loss of methylation being sufficient to disrupt this property (Box 2). Accordingly, partial demethylation of Alu repeats in the B compartment is expected to reduce their ability to position nucleosomes. This suggests that nucleosome positioning may be the key property through which highly methylated Alu sequences exert a constitutive ProA effect. An independent line of evidence supporting this interpretation relates to the potential consequences of such nucleosome positioning on heterochromatin spreading, as discussed later.

Conversely, the youngest Alu subfamilies are, like young L1s and ERVs, targets of CpG-methylation-dependent silencing. Their extremely high methylation levels, together with the presence of strongly positioned nucleosomes, contribute to making them archetypal “chameleon” RepSeqs. Indeed, CpG methylation is predicted to enhance their ProA capacity when the local environment disfavors heterochromatin formation (i.e., in A regions, similar to CorrA.enrichA Alu repeats), and to enhance their ProB capacity in B regions by facilitating heterochromatin assembly. Notably, the very high GC and CpG content of the youngest Alu subfamilies (Extended Data Fig. 11C; Extended Data Fig. 16C) promotes DNMT enzymatic activity ^229^ and thereby likely supports the maintenance of their extreme CpG methylation. These properties are partially lost as sequence drift accumulates over evolution, although CorrA.enrichA Alu subfamilies remain among the most GC- and CpG-rich elements (Extended Data Fig. 11C).

Altogether, these findings support the view that shifts in meCpG levels, whether or not accompanied by a DHS, may encode adjustments in the ProA or ProB output of a DNA element, regardless of whether the element is clearly RepSeq-related or displays a distinctive histone-mark profile.

### RepSeq ProB function is impaired in cancer by both targeted and global mechanisms, resulting in genome opening and plasticity

#### Targeted ProB-to-ProA switching drives local genome unfolding in cancer

Cancer cells exhibit a less compact and more homogeneous chromatin organization than healthy differentiated cells, with increased intermingling of A and B compartments, reminiscent of progenitor or embryonic stem cells ^16,230-232^. These features are indicative of global genome decompaction and suggest that subsets of ProB RepSeqs switch toward a ProA configuration, thereby locally shifting the ProA/ProB balance and unfolding chromatin domains (Fig. 7A), in a manner analogous to genome reorganization observed upon viral infection. Comparative analysis of RepSeq subfamily enrichment in the A and B compartments of HepG2 and HUVEC cells supports this model. While L1s, Alus, and old ERVs display similar B-compartment enrichment in both cell types, a large number of young and middle-aged ERVs are preferentially enriched in the A compartment in the liver cancer cell line HepG2 (Fig. 7B), consistent with a role as drivers of compartment switching and genome unfolding.

Independent studies have shown that numerous ERV LTRs, together with some L1s, are transcriptionally induced in liver cancer, in both human and mouse, through transactivation by MYC in cooperation with liver-specific transcription factors, notably the pioneer factor FOXA3 ^67,83,233^. MYC, a central oncogenic driver of hepatocarcinogenesis, is frequently upregulated in liver cancer from early stages and potently induces tumors upon overexpression in mouse liver ^234-239^. ERV upregulation is paralleled by induction of specific KZFPs, establishing regulatory feedback loops in which ERV inserts act both as repression targets and regulatory elements, and the corresponding KZFPs exert tumor suppressor functions ^240^ (Extended Data Fig. 3). These observations support a model in which MYC, in cooperation with frequent p53 loss, drives widespread ProB-to-ProA switching of RepSeqs during liver tumorigenesis, thereby promoting global genome unfolding.

This framework provides a mechanistic basis for MYC oncogenic activity “without target genes”, a long-standing but unresolved concept ^241^, and explains, at least in part, the well-documented genome-unfolding activity of MYC during lymphocyte activation. In this context, MYC-driven metabolic rewiring additionally contributes to nucleus-wide nucleosomal fiber unpacking through widespread histone hyperacetylation ^188,242-244^. Similar RepSeq-driven mechanisms are likely engaged during iPS reprogramming, which is promoted by MYC overexpression, p53 inactivation, and weakening of HP1α/H3K9me3 heterochromatin, and involves RepSeq subfamilies responsive to Yamanaka factors ^245-247^.

Notably, TEs derepression emerges as a common feature of cancer, involving distinct TE subsets depending on tumor type and driven by tumorigenesis-dependent transcription factor activity (Box 4).

#### Loss of CpG methylation at ProB RepSeqs reflects compartment-specific erosion in cancer

ProB RepSeqs that switch toward a ProA configuration generally undergo CpG demethylation (Box 3). This is recapitulated in an in vitro model of multistage carcinogenesis, in which CpG demethylation affects subsets of RepSeqs early, progressively amplifies, and at later stages may be accompanied by transcriptional activation ^248^. To quantify this effect genome-wide in liver cancer, we measured the fraction of methylated CpG dinucleotides (meCpG fraction) for each individual RepSeq copy in normal liver and in a liver tumor sample using Nanopore sequencing data (Fig. 7C-E) ^223^. Strikingly, not only ERVs but the majority of RepSeq subfamilies displaying a ProB-biased profile, clustered in the left wing of the UMAP, exhibit substantial loss of DNA methylation in the tumor, whereas ProA-biased subfamilies in the right wing are largely spared or even gain methylation.

Analysis of RepSeq DNA methylation across the same 25-Mb region as above reveals that this apparent ProB-selective demethylation is primarily dictated by compartmental context (Fig. 7D). Accordingly, restricting the analysis to Alu/SVA elements (constitutive ProA subclasses) and L1/ERV elements (constitutive ProB subclasses) shows that both categories undergo marked CpG demethylation within the B compartment, but not in the A compartment, in the tumor. Visual inspection of meCpG levels in 25-kb DNA windows confirms this compartment-specific pattern. By contrast, A-compartment regions, including the 5-Mb AlwaysA domain on the left of the 25-Mb region, frequently display equal or higher CpG methylation in the tumor than in adjacent non-tumoral tissue (Fig. 7D; Extended Data Fig. 15A), consistent with previous observations in cancer genomes ^249,250^.

Plotting RepSeq meCpG fraction as a function of local Hi-C EV values further clarifies these trends (Fig. 7E). Despite large differences in absolute methylation levels, Alu/SVA and L1/ERV elements lose CpG methylation to a comparable extent within the B compartment, while exhibiting milder changes in the A compartment. Moreover, CpG loss in cancer correlates more strongly with the local chromatin environment of RepSeq copies than with subfamily age or intrinsic ProB character (Fig. 7E; Extended Data Fig. 16). Thus, ProB RepSeqs appear preferentially demethylated in cancer largely because they are enriched within the B compartment, which undergoes global CpG erosion during tumorigenesis. Centromeric and pericentromeric satellites follow a similar trend (Extended Data Fig. 17), consistent with their increased accessibility and frequent transcriptional activation in tumors ^251-256^.

RepSeq age nonetheless influences CpG methylation. Younger TEs, which are more GC- and CpG-rich, are more heavily methylated in normal tissue, as discussed in a previous section. In particular, very young (VY) L1s and ERVs exhibit higher methylation levels than their older counterparts (Extended Data Fig. 16). Both categories undergo CpG methylation loss in the tumor B compartment; however, this loss is more pronounced for VY L1s, likely reflecting their lower GC content compared to VY ERVs (Extended Data Fig. 16).

#### Sustained signaling reshapes CpG methylation landscapes and redirects Polycomb heterochromatin toward the B compartment

Unexpectedly, analysis of adjacent non-tumoral tissue revealed that most RepSeq subfamilies, across age categories, display a rise in mean CpG methylation within the A compartment, with the notable exception of most L1s, which are markedly AT-rich (Fig. 7E; Extended Data Fig. 16C). This gain in A-compartment methylation is accompanied by a reciprocal, mild decrease within the B compartment. Notably, the cancer meCpG profile strikingly resembles that of the non-tumoral tissue, but with a markedly increased differential between the A and B compartments, a contrast known to progressively amplify during tumorigenesis ^222,230,232,257^. In parallel, in cancer cells, Polycomb-associated chromatin becomes detectable within B-compartment regions, preferentially affecting domains with intermediate GC content and mixed RepSeq composition, which are overrepresented at A/B boundaries (Extended Data Figs. 2, 6) ^230,257,258^. In these regions, Polycomb invasion correlates with partial loss of CpG methylation and repression of a subset of genes (e.g. IFI16), whereas others display both elevated meCpG levels and expression (e.g. WDR11). By contrast, Polycomb heterochromatin is partially depleted from canonical targets such as HOX clusters and the SOX2 locus, coincident with a significant gain in CpG methylation (Extended Data Figs. 7, 14). In normal tissue, these loci are characterized by exceptionally low CpG methylation levels, permissive for efficient Polycomb heterochromatin encroachment ^226^.

The similarity between CpG methylation profiles in non-tumoral and tumor samples likely reflects the fact that in both contexts, cells are exposed to sustained signaling activity, albeit of distinct origins. In peritumoral tissue, this state is plausibly driven by chronic paracrine stimulation and enhanced cell–cell communication within a heterogeneous peritumoral environment ^230-232^, whereas in cancer cells it most often results from constitutive activation of one or more signaling- or hormone-responsive pathways due to oncogenic mutations ^259-262^. Downstream of these pathways, gene expression programs are rewired through two complementary mechanistic strategies. First, sequence-specific TFs induced by signaling cascades directly modulate the activity of cis-regulatory elements, essentially within the A compartment ^260^. Second, signaling also engages locus-independent, genome-wide processes that transiently alleviate heterochromatin-mediated repression within the A compartment, most notably via phosphorylation and acetylation of both histone and non-histone chromatin components, thereby facilitating broad TF access to open regulatory elements ^188,263-265^. These global processes manifest as transient bursts of chromatin accessibility and genome-wide increases in histone acetylation at open regulatory elements, accompanied by a background of rapid, stochastic gene expression upon stimulation ^176,266-268^.

Consistent with this framework, increased methylation of Alu and SVA repeats is observed in AorB:A regions in non-tumoral tissue, bringing their methylation levels close to those of AlwaysA regions (Fig. 7E, lower panel; Extended Data Fig. 16C). This shift is predicted to enhance their ProA capacity (Box 2) and likely to contribute to heterochromatin displacement from the A compartment.

Under physiological conditions, acute stimulation-associated epigenome remodeling is followed by restoration of A-compartment organization over hours to days ^176,266^, including redeployment of Polycomb heterochromatin to re-secure repression of inactive genes. However, when Polycomb eviction from A chromatin is prolonged, antagonistic mechanisms are expected to progressively take over and become durably established. Chief among these are NSD1/2-dependent deposition of H3K36me2 and recruitment of DNMT3A, which together promote CpG methylation and antagonize Polycomb spreading over extended genomic regions ^226,264,269-271^. Sustained activity of this axis provides a parsimonious framework to account for the progressive gain of CpG methylation within the A compartment under chronic signaling conditions.

#### Compensatory repression of ProB RepSeqs by Polycomb heterochromatin in the B compartment likely aggravates meCpG erosion in cancer

Why Polycomb heterochromatin displaced from A-compartment regions subsequently invades the B compartment remains incompletely understood. Importantly, however, contrary to prevailing views, HP1α/H3K9me3- and Polycomb/H3K27me3-based heterochromatin are not mutually exclusive systems but can functionally cooperate ^152^. Although often detected separately by ChIP, co-occupancy of H3K27me3 and H3K9me3 is observed at specific loci, including olfactory receptor (OR) and KZFP gene clusters (Extended Data Figs. 2–3) ^272^. In addition, H3K27me3 “shoulders” are systematically detected at A/B transition zones (see Extended Data Fig. 5 for illustration), indicating functional coupling between Polycomb- and HP1α/H3K9me3-based heterochromatin at compartment boundaries, reminiscent of the mosaic organization observed across the inactive X chromosome in female vertebrates ^273^.

GC-rich ProB RepSeqs that lose CpG methylation in cancer, primarily as a consequence of their location within the B compartment, emerge as particularly favorable substrates for Polycomb encroachment. Such elements are therefore envisioned to act as anchoring sites for Polycomb heterochromatin assemblies within B-compartment regions, thereby providing compensatory repression of ProB RepSeqs ^75^ (Box 2), while at the same time reinforcing a feed-forward process in which Polycomb heterochromatin progressively establishes and comes to dominate the B compartment. The accompanying erosion of CpG methylation in the B compartment may reflect functional interplay between Polycomb chromatin and TET enzymes operating within this newly permissive environment. However, that such CpG methylation loss can occur even in the absence of TET enzymes ^225^ indicates that, while this mechanism may contribute, it is unlikely to represent the primary driver of meCpG erosion.

#### DNMT3B redistribution as a driver of meCpG loss in B compartment during tumorigenesis

As noted above, L1 elements, which are poorer Polycomb substrates than ERVs due to their much lower GC content, exhibit a markedly more pronounced loss of CpG methylation than ERVs within the B compartment in liver cancer (Extended Data Fig. 16C). This differential sensitivity strongly argues against a Polycomb-driven mechanism and instead implicates a primary defect in DNA methylation maintenance as the dominant driver of meCpG erosion in the B compartment during tumorigenesis. Remarkably, the epigenomic alterations universally observed in cancer - including reduced compartmentalization strength without major changes in overall compartment identity, shifts in A/B boundaries accompanied by Polycomb encroachment into the B compartment, increased accessibility of the A compartment, and loss of B compartment and ProB RepSeqs CpG methylation - closely mirror those observed across multiple experimental settings in which both DNMT1 and DNMT3B activity is altered ^228,272,274,275^. DNMT1 functions during S phase as part of a maintenance mechanism preserving methylation memory, and additionally contributes to *de novo* methylation of TEs ^228,276^. DNMT3B is a multi-isoform DNA methyltransferase essential for maintaining global CpG methylation within the B compartment, particularly at L1, ERV, and satellite repeats, acting in cooperation with HP1α, SUV39H1, DNMT3L, and G9a ^152,228,274^. Notably, DNMT1 and DNMT3B functionally cooperate in many contexts, and both enzymes participate in CpG methylation across transcribed gene bodies and at Alu elements in the A compartment ^228,269,277^.

We propose that increased transcriptional activity in the A compartment, together with sustained near-saturation methylation of the extremely abundant Alu repeats, creates a dominant sink for DNMT3B, and possibly also for DNMT1. This redistribution progressively depletes DNMT3B from the B compartment, resulting in passive erosion of CpG methylation that phenocopies experimental DNMT3B depletion. Within this framework, Polycomb invasion of the B compartment is best interpreted as an opportunistic, compensatory response to DNMT3B-driven loss of DNA methylation as recapitulated experimentally ^278 75,228^, although it is likely that both processes operate concomitantly during tumorigenesis and reinforce one another. Notably, in cancer, Polycomb heterochromatin appears to crucially compensate for both the loss of RepSeq repression and the weakening of B-compartment cohesion resulting from meCpG erosion, thereby limiting global genome unfolding ^75,279^.

#### Altered meCpG at ProB RepSeqs as a key driver of cancer cell plasticity

In differentiated cells, chromatin is largely accessible within the A compartment, where the primary role of Polycomb heterochromatin is to maintain non-expressed genes at extremely low levels of transcriptional noise while preserving open promoters poised for acute induction ^280^. This mode of repression is highly dynamic, enabling rapid gene activation, such as that of immediate early genes, within minutes of cellular stimulation (Extended Data Fig. 5). Redistribution of Polycomb heterochromatin toward the B compartment therefore appears sufficient to account for the emergence of elevated transcriptional background noise and entropy in cancer cells, comparable to that observed during early phases of cellular stimulation and in stem cell contexts ^256,266,281-283^.

Beyond Polycomb-dependent deregulation within the A compartment, HP1α/H3K9me3-based heterochromatin, which dominates the B compartment but also pervades the genome, is itself profoundly affected by multiple convergent mechanisms in cancer. First, cell stimulation, whether driven by the tumor microenvironment or by tumor-intrinsic constitutive activation of signaling pathways, is accompanied by global post-translational modifications of H1.0 and HP1α, as well as of nucleosomes and other associated partners, which collectively weaken the HP1α/H3K9me3-based heterochromatin system ^244,263,284-287^. Not only chromosome arms, but also pericentromeres and telomeres can be affected ^256,285,288,289^. Second, tumors commonly harbor mutations or functional alterations in key components of this system, directly compromising both the establishment and maintenance of the ProB state and impairing heterochromatin spreading. Canonical examples include mutation or loss of p53 ^195,259^; H1.0 ^290^; core histones ^291^; the ATRX/DAXX remodeler-chaperone pair required for deposition and maintenance of H3.3K9me3 at RepSeqs ^292-294^; DNMTs; and the HELLS/LSH remodeler specifically required for CpG methylation maintenance at repressed RepSeqs ^295-298^. Finally, as shown above, the global loss of CpG methylation within the B compartment disproportionately affects ProB-type RepSeqs.

While transcription of ProB RepSeqs remains constrained by compensatory repression mechanisms, notably Polycomb heterochromatin (Box 2), the resulting ProB repression is neither as efficient nor as qualitatively robust as in healthy cells. Crucially, CpG demethylation of ProB RepSeqs abolishes the stable memory of the repressed state ^152,221,299^ (Box 2), that is, of the ProB state itself. Beyond enforcing efficient transcriptional inhibition within the B compartment, the ProB system is thought to organize a functional heterochromatin network that makes an essential contribution to three-dimensional genome folding and, more broadly, to genome organization at the scale of the entire nucleus. Loss of CpG methylation at ProB RepSeqs is therefore predicted to weaken the robustness of this global network, emerging as a central determinant of genome-wide organizational plasticity in cancer and facilitating the transcriptional and structural state transitions that characterize tumor cells ^123,300,301^, although this remains to be formally demonstrated.

All of the above conclusions are broadly consistent with observations reported across multiple forms of cancer, but also in ageing. However, while substantial evidence indicates that, in ageing, nuclear three-dimensional reorganization primarily results from a global collapse of the ProB/heterochromatin system ^302-305^, in cancer it most often relies primarily on mechanisms that globally amplify chromatin accessibility and transcriptional activity within the A compartment. In advanced metastatic stages, transcriptional programs normally active during early development and associated with undifferentiated cellular states and specific behaviors, such as increased motility, are frequently reactivated, including epithelial– mesenchymal transition (EMT) ^143,256,306,307^. EMT is driven by pioneer transcription factors such as ZEB1 and TWIST1; remarkably, however, EMT in this context also appears to be accompanied by profound remodeling of the B subcompartment corresponding to constitutive heterochromatin, inducing marked structural genomic instability and likely also directly affecting higher-order genome organization through the network of ProB elements ^256,294,308-310^. In aggressive cancers and metastatic cells, the cancer genome appears globally unfolded within an enlarged nucleus, strikingly reminiscent of pluripotent embryonic stem cells, tightly coupling the loss of chromatin compartmentalization with the emergence of an ultimately lethal cell-fate plasticity ^16,230,259,262,311^.

## DISCUSSION

### A repeat-sequence scaffold underlies euchromatin/ heterochromatin compartmentalization

Eukaryotic genomes are universally partitioned into a euchromatin/A compartment and a heterochromatin/B compartment in the nucleus of differentiated cells, with euchromatin/A being conducive to gene transcription and heterochromatin/B largely non-permissive to transcription. The observations reported in this article, based on simple premises, suggest a model in which the core ProA and ProB repeat sequences (RepSeqs) of a genome, defined as specifically promoting the A and B compartments, respectively, shape a basic A/B landscape that, as a first approximation, can be considered invariant between different cells of an organism, and onto which genes settle. “Always” regions exhibit a strong propensity to be either A or B, whereas “AorB” regions are more plastic, coinciding with a more balanced RepSeq composition (Fig. 8A). While subsets of RepSeq subfamilies as defined herein exhibit pronounced ProA or ProB functions, essentially every RepSeq insert in the genome must be regarded as a composite element with both ProA and ProB capacities, the relative influence of which may vary depending on cellular and genomic contexts and thus on the activities of TFs binding that particular copy.

**Fig. 8:**
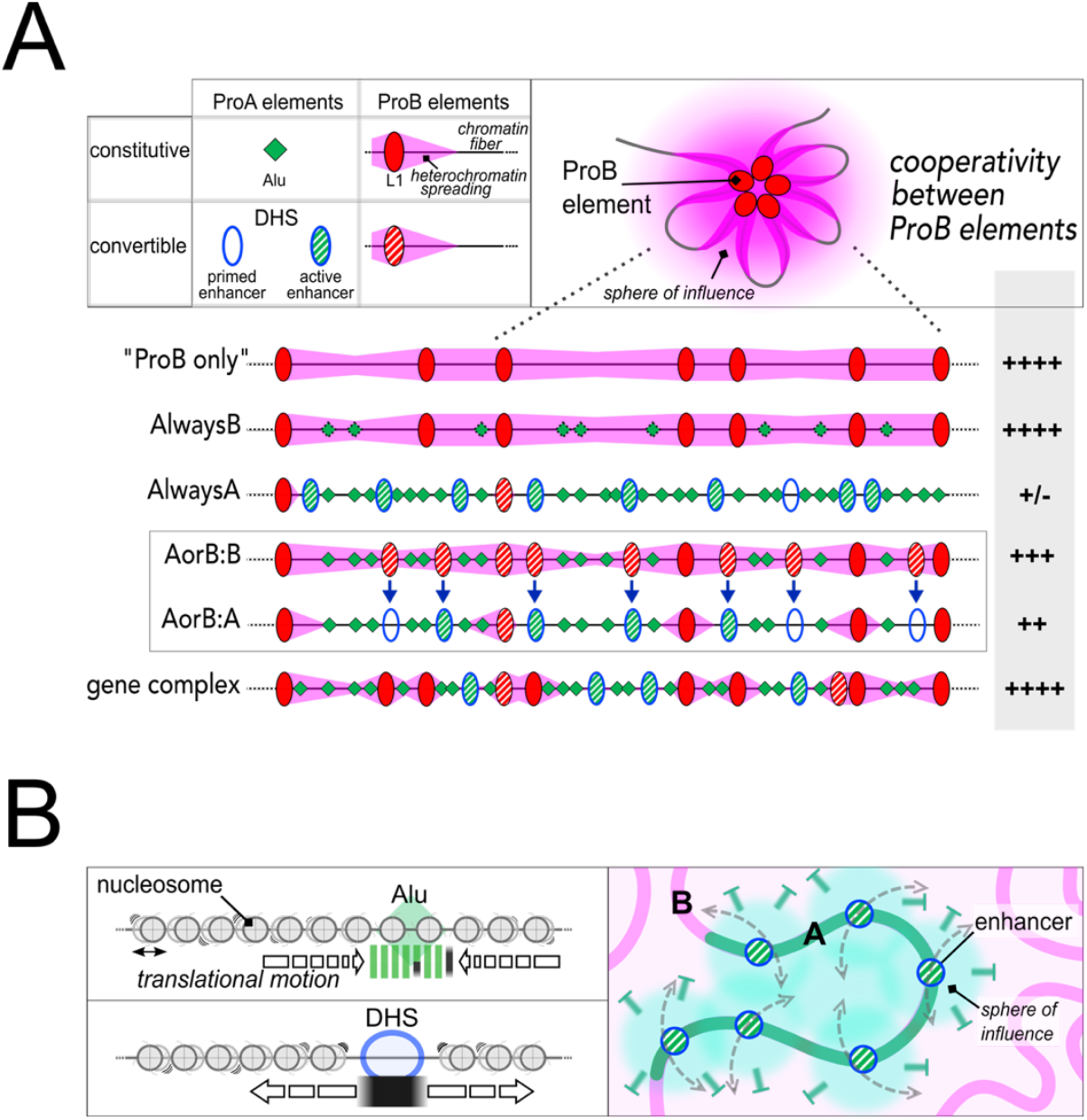
General principles by which ProA and ProB element densities determine A/B partitioning. **(A) Conceptual illustration of two core principles of genome organization.** ProB elements (e.g., young L1s) cooperate to impose a B-type, heterochromatin-based chromatin system over a domain. ProB elements can be constitutive, or convertible into ProA elements through TF action. ProA elements, both constitutive ProA (such as Alu) and convertible ProA (such as DHSs and enhancers), cumulatively disrupt this cooperativity and thereby promote an A-type system over a domain. Active genes (not shown) exert similar ProA effects. In the absence of significant ProA influence, the B state emerges as a default owing to the pervasive presence of ProB elements in the genome. This limited toolkit of ProA and ProB elements underlies the A/B toggle-switch behaviour shown in Fig. 2, as supported by modelling ^20,24,371^. Molecular mechanisms are detailed in Extended Data Fig. 18. **Top, left**. ProB elements act as nucleation points from which heterochromatin appears to “spread” along the chromatin fiber. Pink horizontal triangles indicate the operational extent of spreading, defined as chromatin segments characterized by high levels of heterochromatin marks (e.g., H3K9me3) and strong transcriptional inhibition - two hallmarks of the B compartment. **Top, right**. Conceptual representation of cooperativity among ProB elements (coalescence model, adapted from: ^20^). ProB elements distributed across a domain collectively behave as a higher-order silencing structure, generating a local “sphere of influence” enriched in heterochromatin factors. **Bottom**. When spreading is unimpeded - e.g., in a hypothetical “ProB-only’’ locus - heterochromatin extends across the entire stretch of chromatin embedded in the sphere of high influence. *AlwaysB regions* contain high ProB and low ProA densities. Alu elements mildly attenuate spreading, but ProB cooperativity remains strong. Alu ProA capacity is partially reduced due to lower CpG methylation (green diamonds with dotted contour). *AlwaysA regions* display the opposite pattern, with additional contribution from active genes (not shown). ProB cooperativity is weak, and heterochromatin spreading is inefficient. *AorB regions* contain intermediate levels of both classes of elements, permitting strong ProB cooperativity while spreading is partially mitigated by ProA elements. Convertible elements (e.g., TF-induced DHSs/enhancers) can undergo ProB-to-ProA transitions, collectively shifting a region from B to A (boxed). *Gene complexes* embedded within the B compartment typically display high densities of both ProA and ProB RepSeqs, enabling strong ProB cooperativity while permitting the emergence of localized, transcription-permissive subdomains. **(B) Hypothetical molecular mechanisms by which three types of ProA elements alter the chromatin fiber, thereby interfering with heterochromatin spreading**. Alu repeats and DHSs impede nucleosome translational motion and thereby inhibit heterochromatin spreading - strongly in the case of DHSs, owing to nucleosome eviction, and more moderately for Alu elements, which typically carry one or two well-positioned nucleosomes. CpG methylation supports the ProA function of Alu elements, whereas active DNA demethylation at DHSs may facilitate nucleosome displacement. Active enhancers generate a local “sphere of influence” enriched in transcriptional cofactors (green cloud) that counteract heterochromatin installation (⊣). While primed enhancers (DHSs) distributed along a domain are sufficient to switch a B domain into an A domain, factors associated with active enhancers more effectively secure the A state by cooperatively repelling B influence across the domain (see Extended Data Fig. 18).

pronounced ProA or ProB functions, essentially every RepSeq insert in the genome must be regarded as a composite element with both ProA and ProB capacities, the relative influence of which may vary depending on cellular and genomic contexts and thus on the activities of TFs binding that particular copy.

### DHS/enhancers are inducible ProA elements that open domains

Genes and their regulatory elements located in AorB regions behave as inducible ProA elements, switching these regions to A when they become active and thereby forming new A domains (Fig. 8A). Notably, “activity” here refers not primarily to transcription, but to an active chromatin state (A-type chromatin state), associated with TF binding and nucleosome eviction, producing DHS sites, which appear evenly scattered along the domain. A major fraction of DHSs corresponds to enhancers, known to recruit TFs and cofactors to activate target gene promoters that can be identified as looping partners ^312-314^. In combination, these lines of evidence suggest that another mechanism by which enhancers and bound TFs stimulate gene transcription is separate from *bona fide* transcriptional activation and consists in opposing the formation of B-type chromatin and generalized repression. Such anti-silencing activity of TFs and enhancers was actually postulated and experimentally demonstrated almost three decades ago ^24,105,315-317^. What has become appreciated more recently is that when a gene embedded in a B domain is silent yet should be expressed, there exists an intermediate stage in which the gene is not yet expressed but the domain has already switched to an A configuration (Fig. 6C). This intermediate stage coincides with the appearance of enhancers in an immature form (primed enhancers), which manifest as DHSs marked by H3K4me1 and low levels of H3K27ac on adjacent nucleosomes. Here, we provide additional evidence that a high density of DHSs of all types repels the B compartment, suggesting that an A domain will emerge on a B background precisely over a segment where DHS density is sufficient to oppose the forces driving B-compartment formation (Fig. 8A).

There is indeed evidence for cooperation between DHSs in this process ^130,134^, many of which correspond to enhancers in steady-state conditions. This framework is suggestive of a feed-forward loop whereby DHSs cooperatively enhance accessibility over a region, which in turn promotes TF binding at DHSs, further consolidating their presence. Physically, this mechanism appears to reflect the ability of enhancers to extract the chromatin fiber from the repressive environment formed by large nucleosome clutches, whose cores are enriched in B compartment-typical markers, also conferring a degree of fluidity to the A-type chromatin fiber, whereas B-type chromatin appears more gel-like ^318,319^. Notably, this phenomenon likely underlies the generalized accessibility of open gene domains that was already described more than four decades ago ^320,321^.

Consistent with these notions, enhanced opening of specific domains in two common chronic diseases - schizophrenia and bipolar disorder - correlates with coordinated, tissue-specific upregulation of enhancers scattered along these domains ^322^. Nucleotide polymorphisms predisposing to these diseases are enriched in a number of these enhancers ^322^. Similar findings have been reported in other disorders ^138,323,324^. In particular in cancer, mutations appear to commonly increase the binding affinity of specific TFs at driver gene loci ^325^. These lines of evidence suggest more broadly that a variety of disease-associated enhancers - particularly those for which the search for a single cognate target gene has remained unsuccessful ^314,326,327^ - may act primarily through domain-level anti-silencing activity.

### DHS/enhancers are RepSeqs derivatives that switch between ProA and ProB states

There is a growing awareness that enhancers originate from RepSeqs, and that a significant proportion of enhancers actually are *bona fide* RepSeqs ^69,83,85,87,140,142,328,329^. Furthermore, the literature contains countless examples of situations in which RepSeqs, particularly ERVs, switch from a conspicuously repressed or otherwise nondescript chromatin state to a DHS or “enhancer” state, for instance during viral infection or in cancer, as reported herein, blurring the line between ProA and ProB identity. This is in fact true for every tissue- or developmental stage-specific enhancer, whether or not it is of recognized RepSeq origin, suggesting that DHS/enhancer induction may more generally consist in a switch from a ProB to a ProA state, even in the absence of detectable TF binding and repressive marks prior to induction ^330^.

### A paradigm shift: from gene-centric models to RepSeq-driven genome organization

The genome must therefore be envisioned as a supersystem, with RepSeqs as the main constituents responsible for essentially all levels of organization. Schematically, three major subsystems can be distinguished, each composed of functionally redundant, cooperating elements: RepSeqs with marked ProA or ProB character, which broadly delineate A/B partitioning, and RepSeqs with more indefinite characteristics. In human, a large proportion of the latter are ERV LTRs. These can shift between ProA and ProB functions (“convertible RepSeqs”), depending on the cellular context and the TFs expressed, thereby cooperatively opening or closing AorB domains. Such ERVs and their derivatives, together with derivatives of other RepSeqs that have emerged through idiosyncratic evolution and selection, are better known as transcriptional enhancers. Within this framework, genes and their promoters appear as a special class of RepSeqs - “special guests at the genome party” - which, in their active, transcribed state, reinforce the openness of their surroundings. “Constitutive” ProA and ProB RepSeqs instead form a scaffold of genome organization shared across all cell types, with their relative proportions dictating the propensity of a region to adopt an A- or B-type chromatin state. This perspective essentially confirms the visionary prediction of Emile Zuckerkandl, who wrote almost these exact words in 1974, before any DNA sequence was known ^331^.

In human, constitutive ProA RepSeqs consist primarily of Alu elements, whereas constitutive ProB RepSeqs comprise young L1 elements, ERVs, as well as a panel of AT-rich microsatellites, and telomeric and pericentromeric satellites. The latter form the main components of constitutive heterochromatin, forming megabase-scale blocks of DNA that likely play a specific role in the ProB subsystem by massively assembling heterochromatin material (see below). This remains to be explored using the now available fully sequenced human genomes ^54,91^.

This perspective, which assigns primacy in genome organization to RepSeqs, is consistent with conclusions drawn from more than a century of research on nuclear architecture, as well as with the bioinformatic analyses presented here ^20,24,331-333^. Oddly, however, it represents a major paradigm shift compared to mainstream gene-centered models. Along these lines, another fundamentally new insight concerns the function of enhancers. While enhancers are widely believed to act primarily by directly stimulating cognate promoters, the primary role that emerges from our model is that enhancers act to open domains ^68,69,334^, with their ability to regulate transcription directly coming second. The well-known cooperativity between distant enhancers in gene activation may therefore arise, at least in part, from cooperative effects in domain opening at multiple scales of genome organization ^136,138^.

### What are the molecular mechanisms for ProA and ProB functions?

We have identified subsets of RepSeq subfamilies in the human genome that exhibit the expected molecular features of ProA and ProB elements. These are primarily the Alu and SVA subfamilies for ProA and the young L1 and ERV subfamilies for ProB. However, many additional RepSeqs may contribute in less apparent ways, either because they display a more mixed ProA/ProB character, or because heterogeneity among individual inserts obscures their identification at the subfamily level.

The ProB subfamily archetypes are young L1 and ERVs, with inserts known to be predominantly associated with high levels of H3K9me3/HP1α heterochromatin-related markers; this points to a scenario in which repression of TEs is not only a necessity for the cell to maintain genome stability, but also a prime opportunity for genome organization. Pericentromeric satellites, which are the main constituents of constitutive heterochromatin sequences, and which notoriously nucleate HP1α-based heterochromatin assembly across species in spite of highly divergent sequences ^20,59,335-337^, score among the strongest ProB RepSeqs in our analysis. There is evidence that simple repeats and low-complexity sequences scoring as ProB can also trigger local heterochromatin assembly, in particular via DNA:RNA hybrids or mechanisms fostered by non-B DNA ^48,57,59^. Finally, artificially targeting the KRAB domain of KAP1, a heterochromatin nucleator (see Box 2), to sites along a domain can switch it from A to B ^338^, altogether making a strong case for an equivalence between ProB function and a capacity to nucleate HP1α-based heterochromatin assembly (Fig. 8A, top; Extended data Fig. 18).

The ProA function of Alu may similarly relate to the binding of TFs and associated cofactors (Box 2). Alu subfamilies originate from the 7SL RNA gene that is transcribed by RNA polymerase III (Pol III) from an internal promoter that they inherit, and therefore Alu elements can bind the Pol III-specific initiation factor TFIIIC (Box 2). A ProA-type, antisilencing function has been demonstrated for Pol III-independent TFIIIC-binding elements, and is conserved throughout evolution ^339-342^. All five TFIIIC subunits have histone acetyltransferase activity (HAT) ^343^, and each of these as well as ADNP, known to bind Alu elements as part of the ChAHP complex ^74,344-346^, can interact with the TF MYC ^347,348^. Both TFIIIC and MYC can recruit HATs, suggesting the assembly of a branching, dynamic structure that enables very high local concentrations of HATs at Alu inserts, especially in cells with high MYC expression ^73,188,241^, or upon cell stimulation that activates MYC by phosphorylation ^349^. Pol III is likewise activated upon cell stimulation, and subsets of both tRNA and Alu repeats are rapidly induced ^50,350,351^. Histone acetylation prevents heterochromatin spreading, presumably by inhibiting the formation of nucleosome clutches ^244,352^. This provides a plausible mechanism by which Alu elements can counteract B-type chromatin installation, a mechanism that also applies to active enhancers, which typically recruit HATs, particularly CBP/p300 ^73,312^ (Fig. 8B; Extended Data Fig. 18). As MYC is detected at essentially all active enhancers and promoters in non-quiescent cells, cooperation between these ProA elements in destabilizing nucleosome clutches likely contributes to the massive genome opening observed upon MYC induction ^188,241,353^. Conversely, “low-MYC” conditions may promote genome compaction through the opposite mechanism, as seen in quiescent cells, whose chromatin displays globally low acetylation ^188^.

However, no clear histone acetylation signal is detected at Alu elements in steady-state conditions, nor at extra-TFIIIC sites in the genomes of either *Schizosaccharomyces pombe (S. pombe*) or *Caenorhabditis elegans* ^339,342^. Evidence for the conservation of ProA activity for SINE elements in other species ^62,354^, which also derive from Pol III transcripts, suggests an additional, mechanistically distinct possibility - one that does not rely on TFs or their cofactors. A body of evidence points to a nucleosome-positioning capacity of Alu elements, similar to that of many Pol III-transcribed genes and dependent, at least in the case of Alu, on CpG methylation (Box 2; Fig. 8B; Extended Data Fig. 18). By contrast, heterochromatin assembly is thought to require ATP-dependent translational nucleosome motion, mediated by chromatin remodeling machineries acting together with HMG B-box proteins as cofactors, presumably to optimize HP1α binding and nucleosome bridging ^355-358^. Supporting this view, the FACT complex, best known for for its role in transcription elongation, was recently shown to be essential for heterochromatin maintenance and spreading in *S. pombe* ^359,360^, and to be strongly enriched, together with remodelers such as RSC, ISWI1/2, INO80 and CHD1, in a reconstituted SIRc-heterochromatin assembly in *S. cerevisiae* ^361^. The HMGB factor Nhp6A is similarly enriched in this system ^361^, and HMGB1/2 were found in natural pericentromeric heterochromatin from mouse ES cells ^278^. In the same vein, the variant histone H2A.Z, primarily associated with the priming of transcription elongation at +1 nucleosomes ^362^, appears, paradoxically, to promote heterochromatin spreading and, more generally, the establishment of a B-type chromatin over a domain ^363-366^. H2A.Z-containing nucleosomes display redistributed histone-DNA contacts ^367^, facilitating Pol II passage and nucleosome mobilization, consistent with the notion that nucleosomes must be mobile during heterochromatin installation, as in transcription.

A likely scenario therefore is that Alu element, and presumably also SVA elements, impede the translational remodeling of nucleosomes, thereby interfering with heterochromatin spreading (Fig. 8B; Extended Data Fig. 18). Even minor CpG hypomethylation, as observed in AlwaysB domains in healthy tissues, is expected to reduce the nucleosome-positioning capacity of Alu repeats ^368^ and thus their ProA potential, which may explain the surprising presence of Alu elements in many AlwaysB regions (Extended Data Fig. 7). Non-linear effects are also conceivable: negligible when Alu density is low, as in the AlwaysB subcompartment, but becoming prominent at loci that must remain open or responsive, such as immediate early genes or SOX2 super-enhancers, which exhibit particularly high Alu densities (Extended Data Figs. 5 and 7). The intriguing coexistence of high ProA and high ProB RepSeq densities in gene complexes, as over KZFP gene clusters, embedded within the B compartment, may reflect a genomic environment compatible both with transcription, through local insulation, and with strong repressive control of gene expression and recombination ^20^ (Fig. 8A; Extended Data Fig. 3).

Altogether, the main molecular mechanism by which Alu inserts exert their ProA function likely stems from their occupancy by one or two nucleosomes that are difficult to mobilize. By contrast, DHSs represent interruptions in the “beads-on-a-string” organization of the nucleosomal fiber, and are associated with rapid nucleosome turnover to maintain a nucleosome-free DNA segment. Thus, the ProA capacity of DHSs may similarly derive from the fact that nucleosome translocation through a DHS is virtually impossible - albeit through a mirror-image mechanism to that operating at Alu inserts (Fig. 8B; Extended Data Fig. 18). Such a mechanism was proposed early on for TFs that potently recruit remodelers, to account for their capacity to uncouple adjacent chromatin domains and prevent heterochromatin spreading, as well as for tRNA genes in *S. cerevisiae* ^24,369,370^. The ability of multiple DHSs scattered along a domain to switch it from B to A was further demonstrated by *in silico* modeling ^371^.

Additional details on how these mechanisms translate into distinct chromatin landscapes shaped by varying proportions of ProA and ProB RepSeqs are provided in Fig. 8 and Extended Data Fig. 18. In short, the fundamental principle underlying non-linear behaviors, whereby modest increases in DHS/enhancer density can tip a B domain into an A domain, is the strong cooperativity of ProB elements within the domain, that ProA elements counteract. This cooperativity is likely underpinned, first, by the collective recruitment of condensate-forming heterochromatin components, including HP1α, by ProB elements (Extended Data Fig. 18), and, critically, by the ability of HP1α to stabilize nucleosome clutches. In one-dimensional representations, this effect appears as simple heterochromatin “spreading” along the chromatin fiber; in reality, however, it endows the system with strong synergistic cooperativity that scales up to the chromatin domain level (Fig. 8A; Extended Data Fig. 18). ProA elements interfere with this process through an activity that can be broadly described as anti-silencing, as previously proposed ^20,24^, thereby breaking ProB cooperativity and disengaging chromatin loops from nucleosome clutches ^318^.

### Evolutionary dynamics: how order emerges from repeat-driven genome flux

Genomes are in flux, as recognized by Barbara McClintock decades ago ^372^, with TEs inserting more or less randomly, being possibly fixed in a population, and eventually drifting along evolutionary times. Enhancers emerge essentially from TE inserts, which is how she discovered TEs, but turns out to be a generality, not an anecdote. TEs harbor binding sites for tissue-specific TFs and may acquire others over time by genetic drift. This is consistent with the fact that enhancer sequences are largely species-specific, being conserved only between close phyla, and are fast-evolving sequences in sharp contrast to genes ^85,87,328,373^. Simple repeats may also be involved ^52,143,144^. Any young TE is subjected to repression upon insertion and can seemingly behave as a ProB element. Our observations are consistent with a model in which ProB RepSeqs operate as a system of redundant elements that cooperate by little, dynamic touches across scales and in particular at a domain level ^20,374,375^. The same can be said for ProA elements, except that ProAs seemingly cooperate to oppose the genome driving force that emerges from the ensemble function of ProBs. Sequence drift upon TE aging weakens ProB function, and a TE will be counter-selected in the B compartment if a ProA capacity emerges and takes over, which would no longer be in accordance with the genomic environment. Such a scenario appears to account for the observed distribution of Alu repeats, with the youngest inserts being progressively lost from B, in particular on the Y chromosome, as recognized early on ^78,376^. It should therefore be appreciated that the selection pressure that acts to maintain an adequate RepSeq composition in any DNA segment over the genome also proceeds by small touches. These operate on populations of interchangeable elements, which can be replaced when one or several gradually disappear in the course of evolution, to generate a habitat, which is diffusely more conducive to the dynamic installation of a chromatin system of a particular flavor. This is how order emerges out of chaos within eukaryotic genomes, populated for the most part by repeated sequences, which were until recently denied the right to a functional existence to the point of being described as junk DNA. Repeated sequences operate in a collective manner and thus are subject to selection as an ensemble according to a logic that is totally different from the individualistic logic that operates at the level of genes. Using this conceptual framework and unprecedented technical possibilities, we can now read and contemplate genomes as never before.

#### Box 1: Life history of retrotransposable elements

The life history of retrotransposable elements includes the following steps:

1. **An RNA is produced from a source** full-length TE insert.
2. **Reverse copying by a reverse transcriptase (RT)**, encoded in cis or in trans, **and insertion into the genome**, producing a new TE insert. The new insert may include a promoter, especially when the original promoter is internal to the source TE (e.g. full-length L1, Alu), as well as an enhancer (e.g. the LTR of full-length ERVs, or solo LTRs derived from ERVs by recombination between LTRs). These regulatory sequences are hereafter collectively referred to as RepSeq Regs. TEs display varied levels of GC content, with L1s being overall AT-rich, whereas ERVs are GC-rich (Extended Data Fig. 11). However, RepSeq Regs are usually GC-rich and therefore appear in L1 elements as GC-rich windows within an otherwise AT-rich sequence ^223^.
3. **Regulated expression**. RepSeq Regs are controlled by both transcriptional activators and repressors, as are cis-regulatory gene elements, hereafter referred to as Gene Regs. In particular, ERV subfamilies are expressed in specific cell lineages due to the binding of tissue-specific transcription factors to the LTR enhancer. However, RepSeq Regs are, more than Gene Regs, subject to specific, targeted heterochromatin-mediated repression, which stably suppresses potentially harmful transcription but also recombination between repeats, thereby preserving genome stability ^377^. Heterochromatin targeting at TEs involves dedicated repressive systems and relies on both H3K9me3 deposition and CpG methylation, together with their cognate binding partners (see Box 2).
4. **Sequence drift**, including in particular progressive loss of CpGs (converted to TpGs as a consequence of CpG methylation; see Box 2), correlates with both loss of transcriptional capacity and weakening of repressive mechanisms. Remarkably, some heterochromatin marks can still be observed at TE inserts that have lost all transcriptional capacity, and there are instances in which positive selection for such combinations has been demonstrated ^66^.
5. **TEs as a source of innovation for gene regulation**. At all stages of the idiosyncratic evolution of TE inserts, RepSeq Regs can modulate the activity of neighboring gene promoters. RepSeq Regs are therefore subjected to selection pressure as silencers or enhancers, seemingly acquiring binding sites for some transcription factors while losing others. It has recently become appreciated that gene enhancers (i) are fast-evolving sequences that are, or derive from, RepSeq Regs, particularly from ERV LTRs in human ^87^, and (ii) intrinsically display both silencer and enhancer capacity and can switch between these functions depending on cellular context ^378,379^.

#### Box 2: CpG methylation in repression of L1s and ERVs, and in activation of Alu elements: mechanisms and functions.

The cytosine of CpG dinucleotides is subject to methylation by dedicated enzymes known as DNA methyltransferases (DNMTs). Over evolutionary time, CpGs are progressively lost due to pervasive methylation and the increased likelihood of spontaneous deamination of methylated cytosines (mC) to thymine, resulting in a global depletion of CpGs in the genome relative to other dinucleotides such as GpC ^380^. Despite this overall erosion, RepSeqs, which account for ∼56% of the human genome, harbor ∼61% of all CpG dinucleotides, with a large fraction of the remaining CpGs located in gene exons and their regulatory elements ^381^. Pharmacological inhibition of DNMTs using 5-azacytidine (AZA) or related compounds leads to rapid, global loss of DNA methylation in actively dividing cells. This treatment results in transcriptional activation of a minority of ERVs (Extended Data Fig. 12) and L1 inserts ^382^, enriched, but not restricted to, young subfamilies, whereas transcription of the vast majority of Alu inserts appears largely unaffected ^71,75,228,383^. Strikingly, while levels of ongoing transcription correlate negatively with CpG methylation for L1 elements, as expected, they correlate positively for Alu elements, and this positive correlation is also observed for RNA polymerase III occupancy at Alu loci, as assessed by Pol III ChIP-seq ^71,381^. Finally, the GC and CpG composition of Alu and SVA elements clearly distinguish them from other TE classes (Extended Data Fig. 11), collectively pointing to fundamentally distinct roles of CpG methylation in L1/ERV elements versus Alu/SVA elements, as previously suggested ^29^.

**CpG methylation as a memory mark for heterochromatin-mediated repression of L1s, ERVs, and pericentromeric satellites**

The expression of RepSeqs is inhibited by an interplay of interwoven and largely redundant repressive mechanisms ^384^, with a central role played by H3K9me3/HP1α-based heterochromatin in repressing transcription ^76,195,278,346,385,386^. Target specificity for heterochromatin assembly is primarily conferred through sequence-specific TFs, most notably members of the KZFP multigene family, which recruit the heterochromatin scaffold KAP1, as well as through RNA-based pathways such as the PIWI and HUSH systems. These latter mechanisms are particularly prominent in cells with highly plastic chromatin states, such as during early development or gametogenesis, where they silence transcription-competent inserts ^66,387-391^.

CpG methylation by DNMTs represents an additional key layer enforcing RepSeq repression in mammals ^222^. Methylated CpGs stabilize the heterochromatic state of both genes and RepSeqs within a cell cycle and across cell generations by introducing feed-forward and reciprocal regulatory loops into the heterochromatin regulatory network ^24,215,392^. As such, CpG methylation acts as a molecular memory of repression. This principle has been formally demonstrated by experimental systems in which DNMTs, or TET enzymes that reverse CpG methylation, are artificially targeted to reporter genes, revealing that meCpG confers long-term heritable repression ^152,221,299,384,393^. Consistent with this essential role, DNMT loss is embryonically lethal in mice due to the inability of embryonic cells to stably silence prior or alternative cell identities ^394^. Cultured cells treated with AZA not only display altered TE repression but also exhibit loss of CpG methylation at pericentromeric satellites and subtelomeric RepSeqs ^274^, a phenotype that mirrors observations in patients with Immunodeficiency with Centromeric instability and Facial dysmorphia (ICF) syndrome. In this condition, mutations in DNMT3B or in components of the CDCA7–HELLS complex, which enables DNMT access to heterochromatin-embedded repeats, lead to marked decompaction of specific pericentromeric regions ^395^.

CpG methylation is particularly critical for repression of young ERV subfamilies in differentiated cells (Extended Data Figs. 12 and 15). As progressively older TEs are considered, the gradual loss of CpGs due to sequence drift appears to be compensated by an increased contribution of H3K9me3-based repression, until this mechanism is eventually lost as well ^75^. Notably, a subset of ancient TEs nevertheless retains detectable KAP1 occupancy in an idiosyncratic manner ^66,75^. Intriguingly, this sequence of events, which unfolds over evolutionary timescales, is recapitulated in vitro over a period of weeks when cultured cells are treated with AZA, with H3K9me3 initially increasing at young ERVs while CpG methylation decreases ^75^.

A comparable sequence of events is also observed during early mammalian development, when the genome undergoes global, massive DNA demethylation ^396^. TEs also lose CpG methylation, albeit to more limited and variable extents compared to the rest of the genome, and a subset becomes transcriptionally active, a process that appears necessary for totipotency ^191,393,394,396^. Shortly thereafter, RepSeqs are re-silenced through the ensuing upregulation of KZFPs and associated cofactors, notably the histone methyltransferase SETDB1, resulting in widespread deposition of H3K9me3 at TEs ^69,140^. CpG methylation is subsequently re-established as cells exit pluripotency - of which embryonic stem (ES) cells represent an equivalent - and eventually reaches the meCpG patterns and levels characteristic of differentiated tissues during gastrulation ^394,397^.

In cancer, CpG methylation is broadly lost across the B compartment (Fig. 7; Extended Data Fig. 15A), while H3K9me3 levels generally increase and subsets of KZFPs may be upregulated ^398-400^, echoing the sequence of events observed during evolution and early development. This apparent compensatory rise in H3K9me3 may be further promoted by anti-cancer treatments and likely reflects both an intrinsic cellular response to RepSeq transcription, via activation of heterochromatinization pathways, and selective pressure against RepSeq expression, which would otherwise trigger innate immunity surveillance pathways and elimination of the cancer cell ^204,401-408^ (Box 4).

Interestingly, across evolution, early development, and cancer, loss of RepSeq repression upon CpG demethylation is also compensated by HP1α/H3K9me3-independent mechanisms, most notably through the engagement of Polycomb heterochromatin ^75,228,279,409^. Polycomb heterochromatin assemblies involving PRC1 and PRC2, while more dynamic and typically prominent in the A compartment, are adapted to engage unmethylated, GC-rich sequences ^226,279^. Notably, while alternative repression mechanisms may suffice to limit immune activation and allow tumor cell immune evasion ^401,410^, the quality and robustness of repression itself are clearly not equivalent in the absence of CpG methylation–based memory mechanisms ^299^.

**CpG methylation likely enforces nucleosome positioning over Alu**

There is clear evidence that a large fraction of Alu repeats can be transcribed by RNA polymerase II. This reflects two distinct situations. First, Alu elements may reside near gene enhancers or promoters and be present in eRNA or uaRNA, respectively, which has been shown to structurally stabilize enhancer–promoter interactions ^411^. Alternatively, Alu elements can evolve to become *bona fide* enhancers, binding Pol II–related transcription factors ^72-74,85,87^.

However, Alu elements are primarily transcribed by RNA polymerase III, as they originated by reverse transcription from the non-coding 7SL RNA and therefore retained its internal Pol III promoter. Pol III– transcribed genes are typically short, approximately the size of a nucleosome, and often display a positioned nucleosome, as exemplified by the 5S rRNA gene, a classically used nucleosome-positioning sequence ^412^.

Unlike Pol II, Pol III transcription is compatible with, and can even be promoted by, nucleosome occupancy: (i) TFIIIC, which scaffolds Pol III pre-initiation complex assembly, can bind nucleosome-embedded DNA; (ii) Pol III transcription can proceed without nucleosome eviction; (iii) in vitro reconstitution experiments show that TFIIIC-dependent transcription is more efficient in the presence of a nucleosome ^413^. While some highly transcribed Pol III genes, such as tRNA genes in *S. cerevisiae*, exhibit nucleosome eviction ^370^, the flanking nucleosomes remain strongly positioned and H2A.Z-depleted, a configuration required for efficient transcription ^414^.

Alu elements correspond to dimers of 7SL cDNA, with the internal Pol III promoter conserved in the first monomer. They display among the highest nucleosome occupancies in the human genome, with one positioned nucleosome on each monomer ^368^. The two monomers are separated by an A-rich linker and followed by a poly(A) tract, both of which strongly exclude nucleosomes and are therefore predicted to contribute to precise nucleosome positioning over Alu ^219^. Consistent with a key role of nucleosome positioning, positive selection for mutations predicted to enhance this property has been documented in specific Alu inserts ^219,415,416^.

Given that Alu repeats are among the most highly CpG-methylated DNA sequences in the human genome (Fig. 7; Extended Data Fig. 16), we favor the model that CpG methylation alters DNA structure in a manner that promotes nucleosome positioning and occupancy. This is supported by multiple observations: (i) the CpG distribution within Alu displays a remarkable and conserved grammar, shared by both monomers, with a predominant 8-nt spacing and additional 22- and 44-nt periodicities ^71^, reminiscent of the Widom 601 nucleosome-positioning sequence ^355^; (ii) methylcytosine exhibits distinctive structural properties, including increased base stacking, duplex stability, curvature, and torsional rigidity relative to unmethylated CpG ^215^, and is predicted to subtly alter DNA groove geometry ^368^; (iii) CpG methylation of natural CpG-rich sequences promotes nucleosome stability in vitro and correlates with nucleosome occupancy in vivo ^368,417^.

Collectively, these lines of evidence lend strong support to the idea that the *raison d’être* of cytosine methylation at Alu elements is to promote stable nucleosome positioning and occupancy.

#### Box 3: A scenario for the transition of DNA elements from ProB to ProA configuration ultimately leading to domain opening

Transition of a DNA element from a ProB status to a ProA status may be conceived as a continuum, of which the DMR status may be regarded as an intermediate stage, according to the following scenario. The loss of DNA methylation and the eviction of a nucleosome, initially driven by pioneer TFs and associated cofactors binding to the element, are envisioned as highly dynamic processes. In the initial stages, these events are presumably compensated by reverse reactions, with the loss of methylation/DMR status becoming gradually detectable over time. A minor subset of these elements in a B chromatin segment would clearly emerge as DHS, upon the pivotal binding of some additional TFs, only when a threshold density of such elements is reached within the domain. This would tip the domain into an A configuration and lead in turn to the all-or-none co-actuation of these DHSs by cooperative effects, as experimentally supported by a recent publication ^134^. The other DMRs, which are much more abundant than DHSs, would instead remain as DMRs, possibly progressing to a clearer demethylated stage without ever reaching a detectable DHS stage. Loss of methylation at DMR elements lacking overlap with a DHS, many of which overlap with RepSeqs, may thus contribute to domain opening in a more subtle way than DHSs, by virtue of modulating their ProB function without a clear switch to ProA.

In summary, this model provides insight into how continuous phenomena, such as variations in the concentration of specific TFs, may drive domain switching from a B to an A state, which often appears essentially binary. This likely reflects mutual reinforcement between domain opening and the formation of DHS, concurring to the apparent self-sustainment of the A state at the domain level (Fig. 7A). Whether RepSeqs whose ProB function is simply modulated without switching to an overt ProA DHS or active enhancer state, as is believed to be the case for DMR elements lacking overlap with a DHS, play an important role in this picture remains to be investigated.

#### Box 4: RepSeq transcription is the Achilles’ heel of cancer cells

A whole body of literature shows that RepSeq transcription is a hallmark of cancer. Combinatorial signatures based on a limited set of RepSeqs can be defined, much like gene panels used to score mutations, transcriptional deregulation, or altered chromatin states, enabling tumor sample clustering with high predictive value ^254,255,418-421^. RepSeq repression is secured by redundant pathways in healthy cells (Box 2) such that RepSeq transcription can occur only when several of them are debilitated or dampened, in particular CpG methylation due to either targeted or global deregulation, as discussed in the main text. In addition, the constitutive cancer-associated activation of signaling pathways translates into the binding of TFs that activate specific RepSeq subsets ^193^. However, within a given tumor, RepSeqs that exhibit transcriptional activity typically represent only a minor fraction of all RepSeq inserts that switch to a promoter- or enhancer-associated chromatin state ^422,423^, similar to what we describe herein during development or viral infection. RepSeq derepression, whether or not associated with transcription, can now be understood in terms of a switch from a ProB to ProA function, or an increase in ProA character, with transcription enforcing a marked ProA status.

RepSeq transcription at a high level is also detected in healthy stem and progenitor cells, both in early development and in tissues ^424,425^. Interestingly, while genome unfolding is prominent both in aggressive forms of cancer and healthy stem cells, TE RNA levels may be surprisingly low in cancer cells. This appears to be accounted for by post-transcriptional suppression of TEs by a variety of mechanisms ^16,424 386,426,427^. More broadly, the innate immunity pathway, highly conserved in the animal kingdom and best known for its defense role against microbes, also constitutes a first line immune surveillance system against cancer by detecting TE-derived dsRNA and RNA-DNA hybrids, notably in inflammatory contexts ^423,428^. Physiologically, the detection and tagging of TE-derived nucleic acids as abnormal activity is suppressed in healthy stem and progenitor cells, thereby accommodating the prominent levels of RepSeq transcription associated with physiological genome unfolding and rewiring ^429-432^. The mechanisms involved, which normally ensure such tolerance in stem cells may be subverted by cancer cells as an immune evasion strategy ^426,427^. In addition, the innate immunity system itself is commonly crippled in cancer, via both genetic and epigenetic alterations of gene expression ^256,433,434^, although it is rarely completely ablated. Acutely treating cancer cells with 5-azacytidine, a DNA demethylating agent, induces a sudden and massive burst of RepSeq transcription (Extended Data Fig. 12A). It was first reported in 2015 that this arouses what remains functional of the innate immunity system, turning the tumor into an immunogenic body: the cancer cell becomes unmasked, at least transiently, before having a possibility to adapt ^75,193,383,418,423^. This is the rationale underlying anti-cancer therapeutic strategies that combine immunotherapy with a range of epidrugs, in particular DNMT inhibitors and HDAC inhibitors, which undermine RepSeq repression ^434-438^. This combination may seem paradoxical at first, as these latter treatments are often independently associated with oncogenic effects. Notably, normal cells exhibit much lower levels of RepSeq induction when subjected to the same treatments and show little to no induction of genes involved in the innate immunity response, whereas both RepSeq transcription and innate immunity are strongly boosted by epidrugs in cells exposed to inflammatory cues ^423,438^. Weakened repression of ProB repeat sequences thus emerges as an Achilles’ heel of cancer cells, serving both as a necessity for cancer cell plasticity and, at the same time, generating ‘sneaks’ that alert the immune system to abnormal genome unfolding ^406^.

## RESOURCES & METHODS

### Resources

Data pertaining to chromatin-related features (histone marks, chromatin sensitivity to DNase I) were gathered from validated ChIP-seq and DNase-seq experiments in the HUVEC reference epigenome series of the ENCODE project (ENCSR194DQD), namely for H3K4me3 (ENCSR578QSO), H3K27ac (ENCSR000ALB), H3K27me3 (ENCSR000DVO) and DHS (ENCSR000EOQ). GRO-seq cumulated counts over a 100 kb bin were used as a proxy for RNA polymerase activity, and therefore transcription intensity. GRO-seq data and H3K9me3 data for HUVEC were downloaded from the Gene Expression Omnibus (GEO) under the dataset GSE94872. The samples selected for GRO-seq were the 4 normoxia tracks (GSM2486801 to GSM2486804). A list of regulatory elements that operationally behave as Polycomb-responsive silencers in HUVEC was established from the literature (Table S2 in Ref:^439^) and named “Silencer”. Note that most of these elements score as enhancers in other cellular contexts ^439^. RepeatMasker annotations for hg19, hg19 RefSeq genic feature locations, hg19 GC content, hg19 chromosome size, and ChromHMM ^28^ tracks for HUVEC were downloaded from the UCSC Genome Browser.

Hi-C data were retrieved from the Juicebox database via Juicer Tools (https://hicfiles.s3.amazonaws.com/hiseq). Juicer Tools was used to compute Hi-C EV from Hi-C data at 25, 50, and 100 kb resolutions, and EV values were used to define A and B compartments as described ^1^. Hi-C EV was thus computed for eight human cell lines:

- IMR90 are primary, expanded fibroblasts derived from embryo lung; HUVEC are primary, expanded umbilical vein endothelial cells; NHEK are expanded normal human epidermal keratinocytes;
- HMEC were obtained by SV40 T-antigen-mediated immortalization of endothelial cells; GM12878 is a lymphoblastoid cell line obtained by EBV-mediated immortalization of normal B cells;
- KBM7, K562, and HeLa are cancer-derived cell lines established from chronic myeloid leukemia (KBM7, K562), and cervical carcinoma (HeLa) samples, respectively. HeLa cells harbour HPV-18 sequences and in particular express the HPV-E6 protein.

### AorB vector

The Hi-C EV for the 8 human cell lines was used to generate an aggregate parameter, denoted “AorB.vec” (for “AorB vector”), which conveys the inherent propensity of a genomic region to display A (EV>0) or B (EV<0) chromatin identity, as follows. The value 4 was attributed to bins that display (EV>0) in the eight cell lines, 3 if (EV>0) in 7 cell lines, and so on up to -4 if (EV>0) in 0 cell lines which equates (EV<0) in the eight cell lines. In particular, we could thereby delineate 3 main classes of bins: AlwaysA (EV>0 in 8/8 cell lines), AlwaysB (EV<0 in 8/8 cell lines), AorB (both EV>0 and EV<0 values among the eight cell lines for this bin). In addition, AorB bins display either EV>0 (AorB:A) or EV<0 (AorB:B) in a given cell line. In the HUVEC cell line, AlwaysA, AlwaysB, and AorB fraction represent respectively 22%, 23%, and 55% of hg19 genome, which is largely devoid of pericentromeric and subtelomeric regions. Our analysis therefore applies only to chromosome arms.

### RepSeq insert count and copy count in the human genome

RepSeq were identified in the hg19 genome as segments of homology with the consensus sequences of subfamilies (“NAMEs”) found in the RepBase repository. Individual inserts may be counted multiple times if they are split into more than one segment, mostly due to short INDELs or to the insertion of other TEs, in particular Alu elements. We therefore obtained a proxy for the count of inserts by joining any two segments that are assigned to the same NAME and within 350 bp of each other. Note that 350 bp is slightly above the size of Alu sequences. Counting ERV occurrence requires a specific approach in that an ERV insert may consist either of a full-length provirus made of two LTRs flanking an internal region (“INT”), or of a solo LTR derived by recombination between two identical LTRs in the course of evolution. Indeed, LTR and INT sequences appear as distinct NAMEs in RepBase. ERV insert count was obtained by subtracting INT insert counts from LTR insert counts. “copy count”, which is the metric classically used in bioinformatic analyses refers to the number of segments. We refer to the above proxy for insert counts as “insert count”.

### Correlation with Hi-C EV

We computed the Spearman correlation between the Hi-C eigenvector (Hi-C EV) value in the 8 human cell lines and repeat sequence density (copy counts, 25 kb bins) along the hg19 genome (Table S1, Table S2). The 1,395 subfamilies of RepSeqs found in the RepBase repository (NAMEs) were used. Only the correlation with the Hi-C EV in the HUVEC cell line is used in this study. Correlation values and order in the classification were very similar when the eigenvector derived from another cell line was used (Extended Data Fig. 8; Table S2). Of note, Spearman correlation with the Hi-C EV is negatively impacted when a RepSeq subfamily is absent from a large number of bins, which occurs more often for subfamilies having a low number of copies in the genome.

### Assessing feature density or summed signal in 100 kb bins

Apart from RepSeq, features used in multivariate linear models (see below) are peaks (DHS or ChIP peaks, for instance), intervals (ChromHMM states; Genes), or signals (GRO-seq). Assessing features in 100 kb bins was done with the BEDtools software suite.

#### Gene density

was established as follows. Genes of the same symbolic name on the same chromosome and strand were merged together. Overlapping genes on the same chromosome and strand, which share a common prefix of at least one letter were also merged together. Note that in this work, merging is defined as summarizing several overlapping intervals by a single interval with a starting position as the minimum start of its components, and an end position as the maximum end of its components. Assessing gene 5’ end density allowed to distribute bins into “No gene” (0 gene /100 kb), “Regular gene density” (1-3 genes /100 kb), and “High gene density” (>3 genes /100 kb) subclasses.

#### DHS count

We used the broadPeak file associated with GSM736575, in which the threshold is set at a lower value than with narrowPeak. The idea was to be more sensitive and to detect small peaks that would be missed by narrowPeak, and accordingly the former detects twice more peaks than the latter (25.5 × 10^4^ versus 12 × 10^4^). However, such sensitivity is also associated with a higher background of false peaks, estimated at about 2-5 peaks/100 kb, more pronounced in the B compartment than in the A compartment.

**GRO-seq signals**of the four experimental replicates for HUVEC cultivated in normoxic conditions (see above) were concatenated, and a sum was calculated, which conveys global transcription intensity over a bin. Outliers displaying abnormally strong signals were brought down to a threshold defined according to the Tukey’s Fences method with k=5 upper bound.

#### ChromHMM segments

The HUVEC ChromHMM track was split into individual files for each state. ChromHMM state 4 and state 5 intervals, corresponding to strong enhancers found within or outside of a gene, were gathered into a single list (“Strong.Enhancers”). ChromHMM state 6 and state 7 intervals, corresponding to weak and poised enhancers, respectively, were also grouped together in a single list (“Weak.Enhancers”). Most intervals detected as “Strong promoter state” (state 1) are bracketed by adjacent “weak promoter state” intervals (state 2), which in fact relate to the same cis-regulatory element. In order to avoid overestimating the number of active promoters, we decided to merge state 2 intervals within 1000 bp of a state 1 interval with the latter. The name “Strong.Promoter” was used for the resulting list of merged state 1 and state 2 intervals, and “Weak.Promoter” for the list of state 2 intervals depleted of intervals included in the “Strong.Promoter” list. A bin was considered as “Active” when it contains at least one “Strong.Promoter” or one “Weak.Promoter” interval. Note that ChromHMM state 1 and 2 do not always overlap with gene promoters.

### Multivariate linear model of Hi-C EV profile

Multivariate linear models were generated using either density or mean value for a number of features over 100 kb bins in order to predict the HUVEC Hi-C EV. Both DNA sequence features and epigenomic features as detailed above were used in all possible combinations. DNA features comprised: copy count for different RepSeq subsets, and gene density. Epigenomic features comprised DHS peaks, Strong.Promoter, Weak.Promoter, Strong.Enhancers, Weak.Enhancers, silencer intervals, H3K4me3, H3K27ac, and H3K27me3 peaks, and GRO-seq summed signal. Models were established using values of features as assessed using the hg19.noflank version of the genome, with the idea that trends might be clearer using only the core of chromatin domains, and discarding transition bins. Models were then compared using the adjusted square of the sample Pearson correlation coefficient to estimate the fraction of the variance in the Hi-C EV that is explained by a list of selected features as parameters, whilst adjusting for the number of parameters in the model. Those models with the highest adjusted R-squared value were defined as the most performing.

### GO analysis

A GO analysis was performed for gene-containing bin subclasses 1-16 as defined in our study using data obtained with HUVEC cells. The mapping of GO terms on human genes was obtained from the R package “org.Hs.eg.db” version 3.12. Overrepresentation of GO terms in bin subclasses as defined with HUVEC data was assessed using the classical approach as described ^440^. P-values were calculated with the one-sided Fisher’s exact test and adjusted for multiple comparisons with the BH method. GO terms referred to herein as “enriched GO terms” (or eGO) are those for which enrichment reaches statistical significance (P < 0.05) at the level of the subclass considered, amounting to 1938 GO terms in total for the 16 subclasses. In order to determine whether genes embedded in individual bin subclasses display any specific pattern with regard to functional specialization, we created a list of 34 terms more cross-sectional than GO terms, dubbed “GF-GO.slim”, such that it is possible to assign a unique GF-GO.slim to each GO term by following a procedure detailed below. Although there is some overlap, GF-GO.slim are distinct from classical GO.slim in that a GO term is commonly associated with several GO.slim. First, the full list of GO terms identified in our analysis as enriched in at least one bin subclass was searched for a number of strings, and a GF-GO.slim was attributed in case the string was identified, as detailed below. When the string “signal” was identified, the GF-GO.slim “signaling” was attributed. When either of the strings “stress”, “stimul”, “response” were identified, the GF-GO.slim “stress, stimulus” was attributed, except for GO terms, which are immunity-related, in which case the GF-GO.slim “immunity” was attributed. When either of the strings “nerv”, “neuron”, “axon”, “synapse”, or “dendrit” were identified, the GF-GO.slim “nervous system” was attributed, except if the GO.terms included the words development, differentiation, or commitment, in which case the GF-GO.slim “development, differentiation” was attributed. When either of the strings “skin”, “keratin”, or “corn” were identified, the GF-GO.slim “skin” was attributed, except if the GO.terms included the words development, differentiation, or commitment, in which case the GF-GO.slim “ development, differentiation” was attributed. When the string “musc” was identified, the GF-GO.slim “muscle” was attributed, except if the GO.terms included the words development, differentiation, or commitment, in which case the GF-GO.slim “development, differentiation” was attributed. When the string “ plasma membrane” was identified, the GF-GO.slim “ plasma membrane” was attributed, except if the GO.terms included the words transport, regulation, adhesion, in which case the GF-GO.slim “intracellular transport, localization, cytoskeleton”, “regulation, system process”, or “cell motility, adhesion, polarity, localization”, respectively, were attributed. When the string “regulation”, or exactly the wording “system process” were identified, the GF-GO.slim “regulation” or “system process”, respectively, were attributed, irrespective of the process targeted by the regulation in question. When the string “checkpoint”, “homeosta”, “viral”, or “chromosome” were identified, the GF-GO.slim “checkpoint”, “homeostasis”, “virus related”, or “chromatin, transcription, chromosome”, respectively, were attributed. When none of these possibilities were met, GF-GO.slim were manually attributed in a unambiguous manner, i.e. such that priority matters did not have to be considered. Notably, the GF-GO.slim “development, differentiation” includes terms related to morphogenesis. GF-GO.slim were grouped into four clades: I (01-04): Crosscutting principles; II (05-16): Typical housekeeping functions, such as metabolism and intracellular transport, also including cell cycle, apoptosis, autophagy; III (17-21): Housekeeping functions associated with a specific cellular substructure, possibly associated with a cell differentiated phenotype; IV (22-34): Cell communication with its environment (23,24), chromatin and transcription (25), development and differentiation, and differentiated cell functions, in particular related to Immunity (34). Clade IV thus encompasses what may be called “evolutionarily specialized function”, as opposed to housekeeping functions, which are represented most prominently in clade II and to some extent also in clade III.

### Identification of “outlier bins”: HUVEC.A(vecB), HUVEC.B (vecA)

There are 1601 bins, which display a Hi-C EV sign in HUVEC different from the sign of the AorB vector, i.e. which adopt a chromatin configuration in HUVEC, which does not follow the general trend of the majority of tissues (“outlier bins”). In order to study the characteristics of these regions and to observe marked features, we focused only a subset of these bins (“outstanding outliers”) by applying the following selection criteria: (i) We discarded outlier bins that lie within -100/+100 kb of an A/B junction, as well as isolated bin outlier, i.e. we retained only those bins that form an outlier cluster containing at least 2 bins in a row; (ii) we retained only those outlier bins for which not only the Hi-C EV sign and AorBvec sign are different, but for which the difference in value is significant. To do this, we reduced AorBvec values to -0.05/+0.05 (AorB.vec*), and retained only those outlier bins for which the difference between HUVEC Hi-C EV and AorBvec* is above 0.03 (for HUVEC.A(vecB) bins) and below -0.03 (for HUVEC.B(vecA) bins).

### Analysis of CpG methylation

Oxford Nanopore Technologies (ONT) PromethION data for liver samples were sourced from a previous publication ^223^. CpG Methylation and canonical bases were called from fast5 files using ONT guppy version 6.2.1 using the model “dna_r9.4.1_450bps_modbases_5mc_cg_sup_prom.cfg”. Methylation statistics were generated using methylartist “segmeth” ^441^ for RepeatMasker annotations downloaded from the UCSC Genome Browser assembly hg38 ^442^. RepeatMasker classes retained for analysis were: LTR/ERV1, LTR/ERV1?, LTR/ERVK, LTR/ERVL, LTR/ERVL?, LTR/ERVL-MaLR, LINE/L1, SINE/Alu, and Retroposon/SVA. The ratio of CpG dinucleotides with cytosine methylation to the total number of CpG dinucleotides was calculated for total DNA over 25 kb chromosome segments along the genome (meCpG fraction, 25kb binning) and for each individual RepSeq copy. Hi-C EV for the HepG2 cell line, obtained from the 4D Nucleome website (4DNESC2DEQIJ), was used as a reference.

## Supporting information

Supplementary.Text, Figures_Graphical.Abstract

## Code availability

The code for constructing linear models from feature counts, including links to the publicly available data used in the analysis, is published as free software under the MIT licence and available at https://github.com/qwx9/chrab.

## SUPPLEMENTARY INFORMATION

This manuscript includes supplementary information, namely eighteen Extended Data Figures and two Extended Data Tables.

**Supplementary Table S1. Quantitative features of RepSeq subfamilies**

This table reports quantitative features computed for the 1,395 RepSeq subfamilies annotated in Repbase. RepSeq genomic coordinates and chromatin-related metrics were mapped onto the hg19 human genome assembly and analyzed in HUVEC cells at 100-kb resolution. For each RepSeq subfamily (Class/Family/Name), the table provides Spearman correlation values with the Hi-C eigenvector (corr. Hi-C EV) and with heterochromatin-associated marks (H3K9me3 and H3K27me3), enrichment scores across A/B compartment categories (A and B compartment; Always A and Always B subcompartments), and assignment to the CorrA.enrichA or CorrB.enrichB sets. Additional parameters include Jukes–Cantor distance metrics (JCD mean/min/max/median; proxy for RepSeq age), age category, genomic copy number, and sequence composition (GC and CpG densities). CpG methylation (mean meCpG fraction) was computed from hg38-based datasets and is reported for normal liver (NL) and hepatocellular carcinoma (HCC) samples.

The table contains two worksheets: the first reports the full set of 1,395 RepSeq subfamilies, ordered by Class/Family/Subfamily; the second reports non-TE RepSeqs only, ranked by increasing correlation with the Hi-C eigenvector.

This table provides the primary quantitative input for downstream analyses, including the UMAP representations shown in Fig. 3B and Fig. 7C.

**Supplementary Table S2. RepSeq correlation with Hi-C eigenvector across ENCODE reference cell lines**

This table reports Spearman correlation values between the Hi-C eigenvector (EV) and the genomic distribution of the 1,395 RepSeq subfamilies annotated in Repbase, computed at 25-kb resolution across the eight ENCODE reference cell lines analyzed in Fig. 1A. For each RepSeq subfamily, correlation coefficients are provided together with the corresponding p-values. Data are supplied as CSV files and as an Excel file, in which a color code facilitates visualization of conserved patterns across cell lines, highlighting RepSeq subfamilies displaying cell-type-specific correlation profiles. This table was used for Extended Data Fig. 8.

## ACKNOWLEDGEMENTS

G.F. was supported by INSERM. This work was a labor of love and was not externally funded. K.B. was a student in the Lyon 1 Bioinformatics Master’s program. Communication fees were covered by grants ANR-21-CE13-0037-02 and ANR-21-CE45-0011-01. We thank the teams of Daniel Jost and Olivier Gandrillon for hosting G.F. and for fruitful discussions, and especially Cesar Lopez and Hossein Salari for assistance with figures. We are grateful to Geof Faulkner, Bing Ren, Katherine Chiappinelli, and Tomas Kanholm for providing access to data. We also thank Erica Pehrsson, Armelle Corpet, Sarah Elgin, Helmut Schiessel, Claire Vourc’h, Andras Paldi, Léa Prochasson, Benjamin Audit, Cédric Vaillant, Jeremy Just, and Mohammed Bendahmane for their insightful discussions. Finally, we express our gratitude to Jerry Crabtree, Sylvie Mader, Haitham Shaban, and Toufick Kismoune for their support. We feel deeply indebted to predecessors who shared similar insights. This article is dedicated to the memory of Giorgio Bernardi and Emile Zuckerkandl.

## AUTHOR CONTRIBUTIONS

G.F. conceived and led the project. N.V, M.N., and O.C. performed preliminary computational analysis. Most of the computational analysis, particularly linear models, was carried out by K.B. Spearman correlations were established by R.M. N.H. performed the UMAP analysis. A.E. and N.H. conducted the computational analysis of CpG methylation. G.F. and D.P. wrote the manuscript. E.G. and D.J. contributed with valuable comments on the study, manuscript editing, and provided funding to cover communication fees. All authors discussed the results and reviewed the manuscript.

## REFERENCES

1. Lieberman-Aiden, E. et al. Comprehensive mapping of long-range interactions reveals folding principles of the human genome. Science 326, 289–93 (2009).

2. Harris, H.L. et al. Chromatin alternates between A and B compartments at kilobase scale for subgenic organization. Nat Commun 14, 3303 (2023).

3. Brown, J.M. et al. A tissue-specific self-interacting chromatin domain forms independently of enhancer-promoter interactions. Nat Commun 9, 3849 (2018).

4. Haws, S.A., Simandi, Z., Barnett, R.J. & Phillips-Cremins, J.E. 3D genome, on repeat: Higher-order folding principles of the heterochromatinized repetitive genome. Cell 185, 2690–2707 (2022).

5. Ye, Y., Zhang, S., Gao, L., Zhu, Y. & Zhang, J. Deciphering Hierarchical Chromatin Domains and Preference of Genomic Position Forming Boundaries in Single Mouse Embryonic Stem Cells. Adv Sci (Weinh), e2205162 (2023).

6. Barth, R., Fourel, G. & Shaban, H.A. Dynamics as a cause for the nanoscale organization of the genome. Nucleus 11, 83–98 (2020).

7. Keizer, V.I.P. et al. Live-cell micromanipulation of a genomic locus reveals interphase chromatin mechanics. Science 377, 489–495 (2022).

8. Gabriele, M. et al. Dynamics of CTCF- and cohesin-mediated chromatin looping revealed by live-cell imaging. Science 376, 496–501 (2022).

9. Mach, P. et al. Cohesin and CTCF control the dynamics of chromosome folding. Nat Genet 54, 1907–1918 (2022).

10. Davidson, I.F. et al. CTCF is a DNA-tension-dependent barrier to cohesin-mediated loop extrusion. Nature 616, 822–827 (2023).

11. Nuebler, J., Fudenberg, G., Imakaev, M., Abdennur, N. & Mirny, L.A. Chromatin organization by an interplay of loop extrusion and compartmental segregation. Proc Natl Acad Sci U S A 115, E6697–E6706 (2018).

12. Xie, L. et al. BRD2 compartmentalizes the accessible genome. Nat Genet 54, 481–491 (2022).

13. Mirny, L.A., Imakaev, M. & Abdennur, N. Two major mechanisms of chromosome organization. Curr Opin Cell Biol 58, 142–152 (2019).

14. Belaghzal, H. et al. Liquid chromatin Hi-C characterizes compartment-dependent chromatin interaction dynamics. Nat Genet 53, 367–378 (2021).

15. Dong, P. et al. Cohesin prevents cross-domain gene coactivation. Nat Genet 56, 1654–1664 (2024).

16. Ugarte, F. et al. Progressive Chromatin Condensation and H3K9 Methylation Regulate the Differentiation of Embryonic and Hematopoietic Stem Cells. Stem Cell Reports 5, 728–740 (2015).

17. Magnitov, M.D., Garaev, A.K., Tyakht, A.V., Ulianov, S.V. & Razin, S.V. Pentad: a tool for distance-dependent analysis of Hi-C interactions within and between chromatin compartments. BMC Bioinformatics 23, 116 (2022).

18. Rodriguez-Terrones, D. & Torres-Padilla, M.E. Nimble and Ready to Mingle: Transposon Outbursts of Early Development. Trends Genet 34, 806–820 (2018).

19. Guthmann, M. et al. A change in biophysical properties accompanies heterochromatin formation in mouse embryos. Genes Dev (2023).

20. Fourel, G., Lebrun, E. & Gilson, E. Protosilencers as building blocks for heterochromatin. Bioessays 24, 828–35 (2002).

21. Sanulli, S., Gross, J.D. & Narlikar, G.J. Biophysical Properties of HP1-Mediated Heterochromatin. Cold Spring Harb Symp Quant Biol 84, 217–225 (2019).

22. Haddad, N., Jost, D. & Vaillant, C. Perspectives: using polymer modeling to understand the formation and function of nuclear compartments. Chromosome Res 25, 35–50 (2017).

23. Falk, M. et al. Heterochromatin drives compartmentalization of inverted and conventional nuclei. Nature 570, 395–399 (2019).

24. Fourel, G., Magdinier, F. & Gilson, E. Insulator dynamics and the setting of chromatin domains. Bioessays 26, 523–32 (2004).

25. Rada-Iglesias, A., Grosveld, F.G. & Papantonis, A. Forces driving the three-dimensional folding of eukaryotic genomes. Mol Syst Biol 14, e8214 (2018).

26. Bailey, L.T., Northall, S.J. & Schalch, T. Breakers and amplifiers in chromatin circuitry: acetylation and ubiquitination control the heterochromatin machinery. Curr Opin Struct Biol 71, 156–163 (2021).

27. Bell, O., Burton, A., Dean, C., Gasser, S.M. & Torres-Padilla, M.E. Heterochromatin definition and function. Nat Rev Mol Cell Biol (2023).

28. Ernst, J. et al. Mapping and analysis of chromatin state dynamics in nine human cell types. Nature 473, 43–9 (2011).

29. Pehrsson, E.C., Choudhary, M.N.K., Sundaram, V. & Wang, T. The epigenomic landscape of transposable elements across normal human development and anatomy. Nat Commun 10, 5640 (2019).

30. Carron, L. et al. A layer cake model for plant and metazoan chromatin. bioRxiv, 2023.02.24.529851 (2023).

31. Shopland, L.S. et al. Folding and organization of a contiguous chromosome region according to the gene distribution pattern in primary genomic sequence. J Cell Biol 174, 27–38 (2006).

32. van de Werken, H.J.G. et al. Small chromosomal regions position themselves autonomously according to their chromatin class. Genome Res 27, 922–933 (2017).

33. Jabbari, K. & Bernardi, G. An Isochore Framework Underlies Chromatin Architecture. PLoS One 12, e0168023 (2017).

34. Beaufay, F. et al. Polyphosphate drives bacterial heterochromatin formation. Sci Adv 7, eabk0233 (2021).

35. Kortebi, M. et al. Bacterial chromatin remodeling associated with transcription-induced domains at pathogenicity Islands. bioRxiv, 2025.03.11.642640 (2025).

36. Larson, A.G. et al. Liquid droplet formation by HP1alpha suggests a role for phase separation in heterochromatin. Nature 547, 236–240 (2017).

37. Chapard, C. et al. Exogenous chromosomes reveal how sequence composition drives chromatin assembly, activity, folding and compartmentalization. bioRxiv, 2022.12.21.520625 (2023).

38. Lensch, S. et al. Dynamic spreading of chromatin-mediated gene silencing and reactivation between neighboring genes in single cells. Elife 11(2022).

39. Quante, T. & Bird, A. Do short, frequent DNA sequence motifs mould the epigenome? Nat Rev Mol Cell Biol 17, 257–62 (2016).

40. Kuwayama, N. et al. HMGA2 directly mediates chromatin condensation in association with neuronal fate regulation. Nat Commun 14, 6420 (2023).

41. Sanulli, S. et al. HP1 reshapes nucleosome core to promote phase separation of heterochromatin. Nature 575, 390–394 (2019).

42. Bryan, L.C. et al. Single-molecule kinetic analysis of HP1-chromatin binding reveals a dynamic network of histone modification and DNA interactions. Nucleic Acids Res 45, 10504–10517 (2017).

43. Biswas, S. et al. HP1 oligomerization compensates for low-affinity H3K9me recognition and provides a tunable mechanism for heterochromatin-specific localization. Sci Adv 8, eabk0793 (2022).

44. Becker, J.S. et al. Genomic and Proteomic Resolution of Heterochromatin and Its Restriction of Alternate Fate Genes. Mol Cell 68, 1023–1037 e15 (2017).

45. Kouzine, F. et al. Permanganate/S1 Nuclease Footprinting Reveals Non-B DNA Structures with Regulatory Potential across a Mammalian Genome. Cell Syst 4, 344–356 e7 (2017).

46. Herbert, A. Simple Repeats as Building Blocks for Genetic Computers. Trends Genet 36, 739–750 (2020).

47. Senturk Cetin, N. et al. Isolation and genome-wide characterization of cellular DNA:RNA triplex structures. Nucleic Acids Res 47, 2306–2321 (2019).

48. Zhang, X. et al. KCNQ1OT1 promotes genome-wide transposon repression by guiding RNA-DNA triplexes and HP1 binding. Nat Cell Biol (2022).

49. Grapotte, M. et al. Discovery of widespread transcription initiation at microsatellites predictable by sequence-based deep neural network. Nat Commun 12, 3297 (2021).

50. Maag, J.L. et al. Dynamic expression of long noncoding RNAs and repeat elements in synaptic plasticity. Front Neurosci 9, 351 (2015).

51. Crossley, M.P. et al. R-loop-derived cytoplasmic RNA-DNA hybrids activate an immune response. Nature (2022).

52. Horton, C.A. et al. Short tandem repeats bind transcription factors to tune eukaryotic gene expression. Science 381, eadd1250 (2023).

53. Ma, R. et al. Targeting pericentric non-consecutive motifs for heterochromatin initiation. Nature 631, 678–685 (2024).

54. Franklin, J.M. et al. Human Satellite 3 DNA encodes megabase-scale transcription factor binding platforms. BioRxiv (2025).

55. Herbert, A. Z-DNA and Z-RNA in human disease. Commun Biol 2, 7 (2019).

56. Jiao, H. et al. Z-nucleic-acid sensing triggers ZBP1-dependent necroptosis and inflammation. Nature 580, 391–395 (2020).

57. Gambelli, A., Ferrando, A., Boncristiani, C. & Schoeftner, S. Regulation and function of R-loops at repetitive elements. Biochimie (2023).

58. Liu, Y. et al. Genome-wide mapping reveals R-loops associated with centromeric repeats in maize. Genome Res 31, 1409–1418 (2021).

59. Zhou, J. et al. DDM1-mediated R-loop resolution and H2A.Z exclusion facilitates heterochromatin formation in Arabidopsis. Sci Adv 9, eadg2699 (2023).

60. Niehrs, C. & Luke, B. Regulatory R-loops as facilitators of gene expression and genome stability. Nat Rev Mol Cell Biol 21, 167–178 (2020).

61. Lander, E.S. et al. Initial sequencing and analysis of the human genome. Nature 409, 860–921 (2001).

62. Solovei, I., Thanisch, K. & Feodorova, Y. How to rule the nucleus: divide et impera. Curr Opin Cell Biol 40, 47–59 (2016).

63. Lu, J.Y. et al. Homotypic clustering of L1 and B1/Alu repeats compartmentalizes the 3D genome. Cell Res 31, 613–630 (2021).

64. Moore, B.L., Aitken, S. & Semple, C.A. Integrative modeling reveals the principles of multi-scale chromatin boundary formation in human nuclear organization. Genome Biol 16, 110 (2015).

65. Cao, Q. et al. A unified framework for integrative study of heterogeneous gene regulatory mechanisms. Nature Machine Intelligence 2, 447–456 (2020).

66. Imbeault, M., Helleboid, P.Y. & Trono, D. KRAB zinc-finger proteins contribute to the evolution of gene regulatory networks. Nature 543, 550–554 (2017).

67. Sun, X. et al. Transcription factor profiling reveals molecular choreography and key regulators of human retrotransposon expression. Proc Natl Acad Sci U S A 115, E5526–E5535 (2018).

68. Pontis, J. et al. Hominoid-Specific Transposable Elements and KZFPs Facilitate Human Embryonic Genome Activation and Control Transcription in Naive Human ESCs. Cell Stem Cell 24, 724–735 e5 (2019).

69. Pontis, J. et al. Primate-specific transposable elements shape transcriptional networks during human development. Nat Commun 13, 7178 (2022).

70. Ernst, J. et al. Genome-scale high-resolution mapping of activating and repressive nucleotides in regulatory regions. Nat Biotechnol 34, 1180–1190 (2016).

71. Varshney, D. et al. SINE transcription by RNA polymerase III is suppressed by histone methylation but not by DNA methylation. Nat Commun 6, 6569 (2015).

72. Bouttier, M. et al. Alu repeats as transcriptional regulatory platforms in macrophage responses to M. tuberculosis infection. Nucleic Acids Res 44, 10571–10587 (2016).

73. Policarpi, C. et al. Enhancer SINEs Link Pol III to Pol II Transcription in Neurons. Cell Rep 21, 2879–2894 (2017).

74. Ferrari, R. et al. TFIIIC Binding to Alu Elements Controls Gene Expression via Chromatin Looping and Histone Acetylation. Mol Cell 77, 475–487 e11 (2020).

75. Ohtani, H., Liu, M., Zhou, W., Liang, G. & Jones, P.A. Switching roles for DNA and histone methylation depend on evolutionary ages of human endogenous retroviruses. Genome Res 28, 1147–1157 (2018).

76. McCarthy, R.L. et al. Diverse heterochromatin-associated proteins repress distinct classes of genes and repetitive elements. Nat Cell Biol 23, 905–914 (2021).

77. de Tribolet-Hardy, J. et al. Genetic features and genomic targets of human KRAB-zinc finger proteins. Genome Res (2023).

78. Medstrand, P., van de Lagemaat, L.N. & Mager, D.L. Retroelement distributions in the human genome: variations associated with age and proximity to genes. Genome Res 12, 1483–95 (2002).

79. Fuentes, D.R., Swigut, T. & Wysocka, J. Systematic perturbation of retroviral LTRs reveals widespread long-range effects on human gene regulation. Elife 7(2018).

80. Pal, D. et al. H4K16ac activates the transcription of transposable elements and contributes to their cis-regulatory function. bioRxiv, 2022.04.29.488986 (2022).

81. Fueyo, R., Judd, J., Feschotte, C. & Wysocka, J. Roles of transposable elements in the regulation of mammalian transcription. Nat Rev Mol Cell Biol 23, 481–497 (2022).

82. Carter, T.A. et al. Mosaic cis-regulatory evolution drives transcriptional partitioning of HERVH endogenous retrovirus in the human embryo. Elife 11(2022).

83. Karttunen, K. et al. Transposable elements as tissue-specific enhancers in cancers of endodermal lineage. Nat Commun 14, 5313 (2023).

84. Costello, K.R. et al. Sequence features of retrotransposons allow for epigenetic variability. Elife 10(2021).

85. Zemke, N.R. et al. Comparative single cell epigenomic analysis of gene regulatory programs in the rodent and primate neocortex. bioRxiv (2023).

86. Li, X. et al. LINE-1 transcription activates long-range gene expression. Nat Genet (2024).

87. Zemke, N.R. et al. Conserved and divergent gene regulatory programs of the mammalian neocortex. Nature 624, 390–402 (2023).

88. Lebrun, E., Fourel, G., Defossez, P.A. & Gilson, E. A methyltransferase targeting assay reveals silencer-telomere interactions in budding yeast. Mol Cell Biol 23, 1498–508 (2003).

89. Strom, A.R. et al. Phase separation drives heterochromatin domain formation. Nature 547, 241–245 (2017).

90. Ruault, M. et al. Sir3 mediates long-range chromosome interactions in budding yeast. Genome Res 31, 411–425 (2021).

91. Nurk, S. et al. The complete sequence of a human genome. Science 376, 44–53 (2022).

92. Riddle, N.C. et al. Enrichment of HP1a on Drosophila chromosome 4 genes creates an alternate chromatin structure critical for regulation in this heterochromatic domain. PLoS Genet 8, e1002954 (2012).

93. Riddle, N.C. & Elgin, S.C.R. The Drosophila Dot Chromosome: Where Genes Flourish Amidst Repeats. Genetics 210, 757–772 (2018).

94. Bancaud, A. et al. Molecular crowding affects diffusion and binding of nuclear proteins in heterochromatin and reveals the fractal organization of chromatin. EMBO J 28, 3785–98 (2009).

95. Polovnikov, K.E. et al. Crumpled polymer with loops recapitulates key features of chromosome organization. bioRxiv, 2022.02.01.478588 (2023).

96. Mito, Y., Henikoff, J.G. & Henikoff, S. Histone replacement marks the boundaries of cis-regulatory domains. Science 315, 1408–11 (2007).

97. Vierstra, J. et al. Global reference mapping of human transcription factor footprints. Nature 583, 729–736 (2020).

98. Lazar, J.E. et al. Global Regulatory DNA Potentiation by SMARCA4 Propagates to Selective Gene Expression Programs via Domain-Level Remodeling. Cell Rep 31, 107676 (2020).

99. Comoglio, F. et al. Dissection of acute stimulus-inducible nucleosome remodeling in mammalian cells. Genes Dev 33, 1159–1174 (2019).

100. Paun, O. et al. Pioneer factor ASCL1 cooperates with the mSWI/SNF complex at distal regulatory elements to regulate human neural differentiation. Genes Dev 37, 218–242 (2023).

101. Michael, A.K. et al. Cooperation between bHLH transcription factors and histones for DNA access. Nature (2023).

102. Gamble, N. et al. PU.1 and BCL11B sequentially cooperate with RUNX1 to anchor mSWI/SNF to poise the T cell effector landscape. Nat Immunol (2024).

103. Meuleman, W. et al. Index and biological spectrum of human DNase I hypersensitive sites. Nature 584, 244–251 (2020).

104. Emani, P.S. et al. Single-cell genomics and regulatory networks for 388 human brains. Science 384, eadi5199 (2024).

105. Francastel, C., Walters, M.C., Groudine, M. & Martin, D.I. A functional enhancer suppresses silencing of a transgene and prevents its localization close to centrometric heterochromatin. Cell 99, 259–69 (1999).

106. Schubeler, D. et al. Nuclear localization and histone acetylation: a pathway for chromatin opening and transcriptional activation of the human beta-globin locus. Genes Dev 14, 940–50 (2000).

107. Therizols, P. et al. Chromatin decondensation is sufficient to alter nuclear organization in embryonic stem cells. Science 346, 1238–42 (2014).

108. Karagianni, P., Moulos, P., Schmidt, D., Odom, D.T. & Talianidis, I. Bookmarking by Non-pioneer Transcription Factors during Liver Development Establishes Competence for Future Gene Activation. Cell Rep 30, 1319–1328 e6 (2020).

109. Tedesco, M. et al. Chromatin Velocity reveals epigenetic dynamics by single-cell profiling of heterochromatin and euchromatin. Nat Biotechnol 40, 235–244 (2022).

110. Shah, P.P. et al. An atlas of lamina-associated chromatin across twelve human cell types reveals an intermediate chromatin subtype. Genome Biol 24, 16 (2023).

111. Madsen-Osterbye, J., Abdelhalim, M., Baudement, M.O. & Collas, P. Local euchromatin enrichment in lamina-associated domains anticipates their repositioning in the adipogenic lineage. Genome Biol 23, 91 (2022).

112. Madrigal, P. et al. Epigenetic and transcriptional regulations prime cell fate before division during human pluripotent stem cell differentiation. Nat Commun 14, 405 (2023).

113. Schröder, C.M. et al. An EOMES induced epigenetic deflection initiates lineage commitment at mammalian gastrulation. bioRxiv, 2023.03.15.532746 (2023).

114. Hirabayashi, S. et al. NET-CAGE characterizes the dynamics and topology of human transcribed cis-regulatory elements. Nat Genet 51, 1369–1379 (2019).

115. Barshad, G. et al. RNA polymerase II dynamics shape enhancer-promoter interactions. Nat Genet 55, 1370–1380 (2023).

116. Chen, Z. et al. Increased enhancer-promoter interactions during developmental enhancer activation in mammals. Nat Genet 56, 675–685 (2024).

117. Kitagawa, Y., Ikenaka, A., Sugimura, R., Niwa, A. & Saito, M.K. ZEB2 and MEIS1 independently contribute to hematopoiesis via early hematopoietic enhancer activation. iScience 26, 107893 (2023).

118. Mannens, C.C.A. et al. Chromatin accessibility during human first-trimester neurodevelopment. Nature (2024).

119. Wang, Z. & Ren, B. Role of H3K4 monomethylation in gene regulation. Curr Opin Genet Dev 84, 102153 (2024).

120. Boileau, R.M., Chen, K.X. & Blelloch, R. Loss of MLL3/4 decouples enhancer H3K4 monomethylation, H3K27 acetylation, and gene activation during embryonic stem cell differentiation. Genome Biol 24, 41 (2023).

121. Katsuda, T. et al. Cellular reprogramming in vivo initiated by SOX4 pioneer factor activity. Nat Commun 15, 1761 (2024).

122. Zhang, D. et al. Spatial dynamics of brain development and neuroinflammation. Nature 647, 213–227 (2025).

123. Lim, B. et al. Active repression of cell fate plasticity by PROX1 safeguards hepatocyte identity and prevents liver tumorigenesis. Nat Genet (2025).

124. Menet, J.S., Pescatore, S. & Rosbash, M. CLOCK:BMAL1 is a pioneer-like transcription factor. Genes Dev 28, 8–13 (2014).

125. Furlan-Magaril, M. et al. The global and promoter-centric 3D genome organization temporally resolved during a circadian cycle. Genome Biol 22, 162 (2021).

126. Misra, N., Damara, M., Ye, T. & Chambon, P. The circadian demethylation of a unique intronic deoxymethylCpG-rich island boosts the transcription of its cognate circadian clock output gene. Proc Natl Acad Sci U S A 120, e2214062120 (2023).

127. Marco, A. et al. Mapping the epigenomic and transcriptomic interplay during memory formation and recall in the hippocampal engram ensemble. Nat Neurosci (2020).

128. Larsen, S.B. et al. Establishment, maintenance, and recall of inflammatory memory. Cell Stem Cell 28, 1758–1774 e8 (2021).

129. Jusuf, J.M. et al. Genome-wide absolute quantification of chromatin looping. bioRxiv, 2025.01.13.632736 (2025).

130. Luo, R. et al. Dynamic network-guided CRISPRi screen identifies CTCF-loop-constrained nonlinear enhancer gene regulatory activity during cell state transitions. Nat Genet 55, 1336–1346 (2023).

131. Barajas-Mora, E.M. et al. Enhancer-instructed epigenetic landscape and chromatin compartmentalization dictate a primary antibody repertoire protective against specific bacterial pathogens. Nat Immunol (2023).

132. Sima, J. et al. Identifying cis Elements for Spatiotemporal Control of Mammalian DNA Replication. Cell 176, 816–830 e18 (2019).

133. Pliner, H.A. et al. Cicero Predicts cis-Regulatory DNA Interactions from Single-Cell Chromatin Accessibility Data. Mol Cell 71, 858–871 e8 (2018).

134. Stergachis, A.B., Debo, B.M., Haugen, E., Churchman, L.S. & Stamatoyannopoulos, J.A. Single-molecule regulatory architectures captured by chromatin fiber sequencing. Science 368, 1449–1454 (2020).

135. Vermunt, M.W. et al. Gene silencing dynamics are modulated by transiently active regulatory elements. Mol Cell 83, 715–730 e6 (2023).

136. Lin, X. et al. Nested epistasis enhancer networks for robust genome regulation. Science 377, 1077–1085 (2022).

137. Martinez-Ara, M., Comoglio, F. & van Steensel, B. Large-scale analysis of the integration of enhancer-enhancer signals by promoters. Elife 12(2024).

138. Lin, H. et al. Modular organization of enhancer network provides transcriptional robustness in mammalian development. Nucleic Acids Res 53(2025).

139. Chuong, E.B., Rumi, M.A., Soares, M.J. & Baker, J.C. Endogenous retroviruses function as species-specific enhancer elements in the placenta. Nat Genet 45, 325–9 (2013).

140. Xu, R. et al. Stage-specific H3K9me3 occupancy ensures retrotransposon silencing in human pre-implantation embryos. Cell Stem Cell 29, 1051–1066 e8 (2022).

141. Ito, J. et al. A hominoid-specific endogenous retrovirus may have rewired the gene regulatory network shared between primordial germ cells and naive pluripotent cells. PLoS Genet 18, e1009846 (2022).

142. Roller, M. et al. LINE retrotransposons characterize mammalian tissue-specific and evolutionarily dynamic regulatory regions. Genome Biol 22, 62 (2021).

143. Balestrieri, C. et al. Co-optation of Tandem DNA Repeats for the Maintenance of Mesenchymal Identity. Cell 173, 1150–1164 e14 (2018).

144. Mangan, R.J. et al. Adaptive sequence divergence forged new neurodevelopmental enhancers in humans. Cell 185, 4587–4603 e23 (2022).

145. Andrews, G. et al. Mammalian evolution of human cis-regulatory elements and transcription factor binding sites. Science 380, eabn7930 (2023).

146. Zu, S. et al. Single-cell analysis of chromatin accessibility in the adult mouse brain. Nature 624, 378–389 (2023).

147. Li, Y.E. et al. A comparative atlas of single-cell chromatin accessibility in the human brain. Science 382, eadf7044 (2023).

148. Matsushima, W., Planet, E. & Trono, D. Ancestral genome reconstruction enhances transposable element annotation by identifying degenerate integrants. Cell Genom, 100497 (2024).

149. Young, R.S. et al. The frequent evolutionary birth and death of functional promoters in mouse and human. Genome Res 25, 1546–57 (2015).

150. Carelli, F.N., Liechti, A., Halbert, J., Warnefors, M. & Kaessmann, H. Repurposing of promoters and enhancers during mammalian evolution. Nat Commun 9, 4066 (2018).

151. Oomen, M.E. et al. An atlas of transcription initiation reveals regulatory principles of gene and transposable element expression in early mammalian development. Cell 188, 1156–1174 e20 (2025).

152. Tatarakis, A. et al. Requirements for establishment and epigenetic stability of mammalian heterochromatin. Mol Cell 85, 3388–3406 e12 (2025).

153. Berest, I. et al. Quantification of Differential Transcription Factor Activity and Multiomics-Based Classification into Activators and Repressors: diffTF. Cell Rep 29, 3147–3159 e12 (2019).

154. Hofbauer, L. et al. A genome-wide screen identifies silencers with distinct chromatin properties and mechanisms of repression. Mol Cell 84, 4503–4521 e14 (2024).

155. Radman-Livaja, M. et al. Patterns and mechanisms of ancestral histone protein inheritance in budding yeast. PLoS Biol 9, e1001075 (2011).

156. Ma, L. et al. Co-condensation between transcription factor and coactivator p300 modulates transcriptional bursting kinetics. Mol Cell 81, 1682–1697 e7 (2021).

157. Lim, B. & Levine, M.S. Enhancer-promoter communication: hubs or loops? Curr Opin Genet Dev 67, 5–9 (2021).

158. Sood, V., Holewinski, R., Andresson, T., Larson, D.R. & Misteli, T. Identification of molecular determinants of gene-specific bursting patterns by high-throughput imaging screens. Mol Cell (2025).

159. Gisselbrecht, S.S. et al. Transcriptional Silencers in Drosophila Serve a Dual Role as Transcriptional Enhancers in Alternate Cellular Contexts. Mol Cell 77, 324–337 e8 (2020).

160. Zhu, X. et al. Uncovering the whole genome silencers of human cells via Ss-STARR-seq. Nat Commun 16, 723 (2025).

161. Weidemuller, P., Kholmatov, M., Petsalaki, E. & Zaugg, J.B. Transcription factors: Bridge between cell signaling and gene regulation. Proteomics, e2000034 (2021).

162. Shaban, H.A. et al. Individual transcription factors modulate both the micromovement of chromatin and its long-range structure. Proc Natl Acad Sci U S A 121, e2311374121 (2024).

163. Baker, C.L. et al. Tissue-Specific Trans Regulation of the Mouse Epigenome. Genetics 211, 831–845 (2019).

164. Hawe, J.S. et al. Genetic variation influencing DNA methylation provides insights into molecular mechanisms regulating genomic function. Nat Genet 54, 18–29 (2022).

165. Bertozzi, T.M., Elmer, J.L., Macfarlan, T.S. & Ferguson-Smith, A.C. KRAB zinc finger protein diversification drives mammalian interindividual methylation variability. Proc Natl Acad Sci U S A 117, 31290–31300 (2020).

166. Mowery, C.T. et al. Systematic decoding of cis gene regulation defines context-dependent control of the multi-gene costimulatory receptor locus in human T cells. Nat Genet (2024).

167. Yang, F. et al. Shape of promoter antisense RNAs regulates ligand-induced transcription activation. Nature 595, 444–449 (2021).

168. Kuang, M. et al. XAF1 promotes anti-RNA virus immune responses by regulating chromatin accessibility. Sci Adv 9, eadg5211 (2023).

169. Schmidt, N. et al. An influenza virus-triggered SUMO switch orchestrates co-opted endogenous retroviruses to stimulate host antiviral immunity. Proc Natl Acad Sci U S A 116, 17399–17408 (2019).

170. Wei, K. et al. TRIM28 is an essential regulator of three-dimensional chromatin state underpinning CD8(+) T cell activation. Nat Commun 16, 750 (2025).

171. Wang, L. et al. An eRNA transcription checkpoint for diverse signal-dependent enhancer activation programs. Nat Genet 57, 962–972 (2025).

172. Jacobs, J., Pagani, M., Wenzl, C. & Stark, A. Widespread regulatory specificities between transcriptional co-repressors and enhancers in Drosophila. Science 381, 198–204 (2023).

173. Katsuda, T., Sussman, J.H., Zaret, K.S. & Stanger, B.Z. The yin and yang of pioneer transcription factors: Dual roles in repression and activation. Bioessays 46, e2400138 (2024).

174. Waterbury, A.L. et al. An autoinhibitory switch of the LSD1 disordered region controls enhancer silencing. Mol Cell (2024).

175. Bahrami, S. & Drablos, F. Gene regulation in the immediate-early response process. Adv Biol Regul 62, 37–49 (2016).

176. Fernandez-Albert, J. et al. Immediate and deferred epigenomic signatures of in vivo neuronal activation in mouse hippocampus. Nat Neurosci 22, 1718–1730 (2019).

177. Gerosa, L. et al. SRF and SRFDelta5 Splicing Isoform Recruit Corepressor LSD1/KDM1A Modifying Structural Neuroplasticity and Environmental Stress Response. Mol Neurobiol 57, 393–407 (2020).

178. Agarwal, S. et al. KDM1A maintains genome-wide homeostasis of transcriptional enhancers. Genome Res 31, 186–97 (2021).

179. Sigvardsson, M. Transcription factor networks link B-lymphocyte development and malignant transformation in leukemia. Genes Dev 37, 703–723 (2023).

180. Castro, F.L. et al. Modulation of HERV Expression by Four Different Encephalitic Arboviruses during Infection of Human Primary Astrocytes. Viruses 14(2022).

181. Leung, A. et al. LTRs activated by Epstein-Barr virus-induced transformation of B cells alter the transcriptome. Genome Res 28, 1791–1798 (2018).

182. Zhao, B. et al. Epstein-Barr virus exploits intrinsic B-lymphocyte transcription programs to achieve immortal cell growth. Proc Natl Acad Sci U S A 108, 14902–7 (2011).

183. Okabe, A. et al. Cross-species chromatin interactions drive transcriptional rewiring in Epstein-Barr virus-positive gastric adenocarcinoma. Nat Genet 52, 919–930 (2020).

184. Wang, C. et al. A DNA tumor virus globally reprograms host 3D genome architecture to achieve immortal growth. Nat Commun 14, 1598 (2023).

185. Wang, L. et al. Epstein-Barr Virus Episome Physically Interacts with Active Regions of the Host Genome in Lymphoblastoid Cells. J Virol 94(2020).

186. Ohashi, M., Hayes, M., McChesney, K. & Johannsen, E. Epstein-Barr virus nuclear antigen 3C (EBNA3C) interacts with the metabolism sensing C-terminal binding protein (CtBP) repressor to upregulate host genes. PLoS Pathog 17, e1009419 (2021).

187. Stanfield, B.A. & Luftig, M.A. Recent advances in understanding Epstein-Barr virus. F1000Res 6, 386 (2017).

188. Kieffer-Kwon, K.R. et al. Myc Regulates Chromatin Decompaction and Nuclear Architecture during B Cell Activation. Mol Cell 67, 566–578 e10 (2017).

189. Xu, M. et al. Epstein-Barr virus-encoded miR-BART11-3p modulates the DUSP6-MAPK axis to promote gastric cancer cell proliferation and metastasis. J Virol, e0088123 (2023).

190. Wood, C.D. et al. MYC activation and BCL2L11 silencing by a tumour virus through the large-scale reconfiguration of enhancer-promoter hubs. Elife 5(2016).

191. Jachowicz, J.W. et al. LINE-1 activation after fertilization regulates global chromatin accessibility in the early mouse embryo. Nat Genet 49, 1502–1510 (2017).

192. Aloni-Grinstein, R., Charni-Natan, M., Solomon, H. & Rotter, V. p53 and the Viral Connection: Back into the Future (double dagger). Cancers (Basel) 10(2018).

193. Leonova, K.I. et al. p53 cooperates with DNA methylation and a suicidal interferon response to maintain epigenetic silencing of repeats and noncoding RNAs. Proc Natl Acad Sci U S A 110, E89–98 (2013).

194. Wylie, A. et al. p53 genes function to restrain mobile elements. Genes Dev 30, 64–77 (2016).

195. Tiwari, B. et al. p53 directly represses human LINE1 transposons. Genes Dev (2020).

196. Sun, S. et al. Cancer cells co-evolve with retrotransposons to mitigate viral mimicry. bioRxiv (2023).

197. Isbel, L. et al. Readout of histone methylation by Trim24 locally restricts chromatin opening by p53. Nat Struct Mol Biol 30, 948–957 (2023).

198. Giam, C.Z. & Semmes, O.J. HTLV-1 Infection and Adult T-Cell Leukemia/Lymphoma-A Tale of Two Proteins: Tax and HBZ. Viruses 8(2016).

199. Alarcon, V. et al. The enzymes LSD1 and Set1A cooperate with the viral protein HBx to establish an active hepatitis B viral chromatin state. Sci Rep 6, 25901 (2016).

200. Dias, J.D., Sarica, N., Cournac, A., Koszul, R. & Neuveut, C. Crosstalk between Hepatitis B Virus and the 3D Genome Structure. Viruses 14(2022).

201. Liu, W. et al. Molecular insights into Spindlin1-HBx interplay and its impact on HBV transcription from cccDNA minichromosome. Nat Commun 14, 4663 (2023).

202. Heinz, S. et al. Transcription Elongation Can Affect Genome 3D Structure. Cell 174, 1522–1536 e22 (2018).

203. Ryabchenko, B. et al. The interactions between PML nuclear bodies and small and medium size DNA viruses. Virol J 20, 82 (2023).

204. Corpet, A. et al. PML nuclear bodies and chromatin dynamics: catch me if you can! Nucleic Acids Res 48, 11890–11912 (2020).

205. Yurkovetskiy, L. et al. Primate immunodeficiency virus proteins Vpx and Vpr counteract transcriptional repression of proviruses by the HUSH complex. Nat Microbiol 3, 1354–1361 (2018).

206. Roubille, S. et al. The HUSH epigenetic repressor complex silences PML nuclear bodies-associated HSV-1 quiescent genomes. bioRxiv, 2024.06.18.599571 (2024).

207. Jan Fada, B. et al. A Novel Recognition by the E3 Ubiquitin Ligase of HSV-1 ICP0 Enhances the Degradation of PML Isoform I to Prevent ND10 Reformation in Late Infection. Viruses 15(2023).

208. Neugebauer, E. et al. Herpesviruses mimic zygotic genome activation to promote viral replication. Nat Commun 16, 710 (2025).

209. De Iaco, A. et al. DUX-family transcription factors regulate zygotic genome activation in placental mammals. Nat Genet 49, 941–945 (2017).

210. Sakashita, A. et al. Transcription of MERVL retrotransposons is required for preimplantation embryo development. Nat Genet 55, 484–495 (2023).

211. Zhou, W. et al. DNA methylation loss in late-replicating domains is linked to mitotic cell division. Nat Genet 50, 591–602 (2018).

212. Ginno, P.A. et al. A genome-scale map of DNA methylation turnover identifies site-specific dependencies of DNMT and TET activity. Nat Commun 11, 2680 (2020).

213. Sardina, J.L. et al. Transcription Factors Drive Tet2-Mediated Enhancer Demethylation to Reprogram Cell Fate. Cell Stem Cell 23, 727–741 e9 (2018).

214. Stadhouders, R. et al. Transcription factors orchestrate dynamic interplay between genome topology and gene regulation during cell reprogramming. Nat Genet 50, 238–249 (2018).

215. Rausch, C., Hastert, F.D. & Cardoso, M.C. DNA Modification Readers and Writers and Their Interplay. J Mol Biol (2019).

216. Grand, R.S. et al. BANP opens chromatin and activates CpG-island-regulated genes. Nature 596, 133–137 (2021).

217. Kreibich, E., Kleinendorst, R., Barzaghi, G., Kaspar, S. & Krebs, A.R. Single-molecule footprinting identifies context-dependent regulation of enhancers by DNA methylation. Mol Cell 83, 787–802 e9 (2023).

218. Tillo, D. et al. High nucleosome occupancy is encoded at human regulatory sequences. PLoS One 5, e9129 (2010).

219. Barbier, J., Vaillant, C., Volff, J.N., Brunet, F.G. & Audit, B. Coupling between Sequence-Mediated Nucleosome Organization and Genome Evolution. Genes (Basel) 12(2021).

220. Matsushima, W. et al. Zinc-finger proteins with a co-opted capsid domain anchor nucleosomes over transposon sequences. bioRxiv, 2025.03.03.638093 (2025).

221. Hathaway, N.A. et al. Dynamics and memory of heterochromatin in living cells. Cell 149, 1447–60 (2012).

222. Smith, Z.D., Hetzel, S. & Meissner, A. DNA methylation in mammalian development and disease. Nat Rev Genet 26, 7–30 (2025).

223. Ewing, A.D. et al. Nanopore Sequencing Enables Comprehensive Transposable Element Epigenomic Profiling. Mol Cell 80, 915–928 e5 (2020).

224. Colombo, A.R., Elias, H.K. & Ramsingh, G. Senescence induction universally activates transposable element expression. Cell Cycle 17, 1846–1857 (2018).

225. Lopez-Moyado, I.F. et al. Paradoxical association of TET loss of function with genome-wide DNA hypomethylation. Proc Natl Acad Sci U S A 116, 16933–16942 (2019).

226. Owen, B.M. & Davidovich, C. DNA binding by polycomb-group proteins: searching for the link to CpG islands. Nucleic Acids Res 50, 4813–4839 (2022).

227. Zhou, J. et al. Human Body Single-Cell Atlas of 3D Genome Organization and DNA Methylation. bioRxiv, 2025.03.23.644697 (2025).

228. Scelfo, A. et al. Tunable DNMT1 degradation reveals DNMT1/DNMT3B synergy in DNA methylation and genome organization. J Cell Biol 223(2024).

229. Bonsu, K.A., Trinh, A., Downing, T.L. & Read, E.L. A Tunable, Ultrasensitive Threshold in Enzymatic Activity Governs the DNA Methylation Landscape. bioRxiv, 2024.06.25.600710 (2024).

230. Johnstone, S.E. et al. Large-Scale Topological Changes Restrain Malignant Progression in Colorectal Cancer. Cell 182, 1474–1489 e23 (2020).

231. Xu, J. et al. Super-resolution imaging reveals the evolution of higher-order chromatin folding in early carcinogenesis. Nat Commun 11, 1899 (2020).

232. Xue, Y. et al. Unraveling the key role of chromatin structure in cancer development through epigenetic landscape characterization of oral cancer. Mol Cancer 23, 190 (2024).

233. Hashimoto, K. et al. CAGE profiling of ncRNAs in hepatocellular carcinoma reveals widespread activation of retroviral LTR promoters in virus-induced tumors. Genome Res 25, 1812–24 (2015).

234. Fourel, G. et al. Frequent activation of N-myc genes by hepadnavirus insertion in woodchuck liver tumours. Nature 347, 294–8 (1990).

235. Yang, D., Alt, E. & Rogler, C.E. Coordinate expression of N-myc 2 and insulin-like growth factor II in precancerous altered hepatic foci in woodchuck hepatitis virus carriers. Cancer Res 53, 2020–7 (1993).

236. Etiemble, J. et al. Liver-specific expression and high oncogenic efficiency of a c-myc transgene activated by woodchuck hepatitis virus insertion. Oncogene 9, 727–37 (1994).

237. Kaposi-Novak, P. et al. Central role of c-Myc during malignant conversion in human hepatocarcinogenesis. Cancer Res 69, 2775–82 (2009).

238. Buendia, M.A. & Neuveut, C. Hepatocellular carcinoma. Cold Spring Harb Perspect Med 5, a021444 (2015).

239. Senapedis, W. et al. Targeted transcriptional downregulation of MYC using epigenomic controllers demonstrates antitumor activity in hepatocellular carcinoma models. Nat Commun 15, 7875 (2024).

240. Ito, J. et al. Endogenous retroviruses drive KRAB zinc-finger protein family expression for tumor suppression. Sci Adv 6(2020).

241. Baluapuri, A., Wolf, E. & Eilers, M. Target gene-independent functions of MYC oncoproteins. Nat Rev Mol Cell Biol 21, 255–267 (2020).

242. Dejure, F.R. & Eilers, M. MYC and tumor metabolism: chicken and egg. EMBO J 36, 3409–3420 (2017).

243. Tan, H. et al. Integrative Proteomics and Phosphoproteomics Profiling Reveals Dynamic Signaling Networks and Bioenergetics Pathways Underlying T Cell Activation. Immunity 46, 488–503 (2017).

244. Gibson, B.A. et al. Organization of Chromatin by Intrinsic and Regulated Phase Separation. Cell 179, 470–484 e21 (2019).

245. Hong, H. et al. Suppression of induced pluripotent stem cell generation by the p53-p21 pathway. Nature 460, 1132–5 (2009).

246. Nicetto, D. & Zaret, K.S. Role of H3K9me3 heterochromatin in cell identity establishment and maintenance. Curr Opin Genet Dev 55, 1–10 (2019).

247. Ito, J. et al. Systematic identification and characterization of regulatory elements derived from human endogenous retroviruses. PLoS Genet 13, e1006883 (2017).

248. Kanholm, T. et al. Oncogenic Transformation Drives DNA Methylation Loss and Transcriptional Activation of Transposable Element Loci. Cancer Res (2023).

249. Berman, B.P. et al. Regions of focal DNA hypermethylation and long-range hypomethylation in colorectal cancer coincide with nuclear lamina-associated domains. Nat Genet 44, 40–6 (2011).

250. Feinberg, A.P. The Key Role of Epigenetics in Human Disease Prevention and Mitigation. N Engl J Med 378, 1323–1334 (2018).

251. Narayan, A. et al. Hypomethylation of pericentromeric DNA in breast adenocarcinomas. Int J Cancer 77, 833–8 (1998).

252. Feinberg, A.P. & Tycko, B. The history of cancer epigenetics. Nat Rev Cancer 4, 143–53 (2004).

253. Pappalardo, X.G. & Barra, V. Losing DNA methylation at repetitive elements and breaking bad. Epigenetics Chromatin 14, 25 (2021).

254. Eymery, A. et al. A transcriptomic analysis of human centromeric and pericentric sequences in normal and tumor cells. Nucleic Acids Res 37, 6340–54 (2009).

255. Solovyov, A. et al. Global Cancer Transcriptome Quantifies Repeat Element Polarization between Immunotherapy Responsive and T Cell Suppressive Classes. Cell Rep 23, 512–521 (2018).

256. Perelli, L. et al. Evolutionary fingerprints of epithelial-to-mesenchymal transition. Nature 640, 1083–1092 (2025).

257. Guo, H. et al. DNA hypomethylation silences anti-tumor immune genes in early prostate cancer and CTCs. Cell 186, 2765–2782 e28 (2023).

258. Lakatos, E. et al. Epigenetically driven and early immune evasion in colorectal cancer evolution. Nat Genet 57, 3039–3049 (2025).

259. Hanahan, D. & Weinberg, R.A. Hallmarks of cancer: the next generation. Cell 144, 646–74 (2011).

260. Gaynor, L. et al. Crypt density and recruited enhancers underlie intestinal tumour initiation. Nature 640, 231–239 (2025).

261. Haga, Y. et al. Whole-genome sequencing reveals the molecular implications of the stepwise progression of lung adenocarcinoma. Nat Commun 14, 8375 (2023).

262. Berns, A. Transforming lung cancer types. Science 383, 590–591 (2024).

263. Azad, G.K. et al. PARP1-dependent eviction of the linker histone H1 mediates immediate early gene expression during neuronal activation. J Cell Biol 217, 473–481 (2018).

264. Willcockson, M.A. et al. H1 histones control the epigenetic landscape by local chromatin compaction. Nature 589, 293–298 (2021).

265. Matthews, R.E. et al. CRAMP1 drives linker histone expression to enable Polycomb repression. Mol Cell (2025).

266. Parmentier, R. et al. Global genome decompaction leads to stochastic activation of gene expression as a first step toward fate commitment in human hematopoietic cells. PLoS Biol 20, e3001849 (2022).

267. Gao, N.P., Gandrillon, O., Paldi, A., Herbach, U. & Gunawan, R. Single-cell transcriptional uncertainty landscape of cell differentiation. F1000Res 12, 426 (2023).

268. Beato, M., Wright, R.H.G. & Dily, F.L. 90 YEARS OF PROGESTERONE: Molecular mechanisms of progesterone receptor action on the breast cancer genome. J Mol Endocrinol 65, T65–T79 (2020).

269. Weinberg, D.N. et al. Two competing mechanisms of DNMT3A recruitment regulate the dynamics of de novo DNA methylation at PRC1-targeted CpG islands. Nat Genet 53, 794–800 (2021).

270. Padilla, R. et al. H3K36 Methylation as a Guardian of Epigenome Integrity. Nat Commun (2025).

271. Yuan, W. et al. H3K36 methylation antagonizes PRC2-mediated H3K27 methylation. J Biol Chem 286, 7983–7989 (2011).

272. Spracklin, G. et al. Diverse silent chromatin states modulate genome compartmentalization and loop extrusion barriers. Nat Struct Mol Biol 30, 38–51 (2023).

273. Keniry, A. et al. Setdb1-mediated H3K9 methylation is enriched on the inactive X and plays a role in its epigenetic silencing. Epigenetics Chromatin 9, 16 (2016).

274. Horard, B. et al. Global analysis of DNA methylation and transcription of human repetitive sequences. Epigenetics 4, 339–50 (2009).

275. Du, Q. et al. DNA methylation is required to maintain both DNA replication timing precision and 3D genome organization integrity. Cell Rep 36, 109722 (2021).

276. Haggerty, C. et al. Dnmt1 has de novo activity targeted to transposable elements. Nat Struct Mol Biol 28, 594–603 (2021).

277. Baubec, T. et al. Genomic profiling of DNA methyltransferases reveals a role for DNMT3B in genic methylation. Nature 520, 243–7 (2015).

278. Saksouk, N. et al. Redundant mechanisms to form silent chromatin at pericentromeric regions rely on BEND3 and DNA methylation. Mol Cell 56, 580–94 (2014).

279. Fukuda, K. et al. Epigenetic plasticity safeguards heterochromatin configuration in mammals. Nucleic Acids Res 51, 6190–6207 (2023).

280. Szczurek, A.T., Dimitrova, E., Kelley, J.R. & Klose, R.J. Polycomb sustains promoters in a deep OFF-state by limiting PIC formation to counteract transcription. bioRxiv, 2023.06.13.544762 (2023).

281. Feinberg, A.P. & Levchenko, A. Epigenetics as a mediator of plasticity in cancer. Science 379, eaaw3835 (2023).

282. Richard, A. et al. Single-Cell-Based Analysis Highlights a Surge in Cell-to-Cell Molecular Variability Preceding Irreversible Commitment in a Differentiation Process. PLoS Biol 14, e1002585 (2016).

283. Niskanen, H. et al. Endothelial cell differentiation is encompassed by changes in long range interactions between inactive chromatin regions. Nucleic Acids Res 46, 1724–1740 (2018).

284. Nishibuchi, G. et al. Mitotic phosphorylation of HP1alpha regulates its cell cycle-dependent chromatin binding. J Biochem 165, 433–446 (2019).

285. Noh, K.M. et al. ATRX tolerates activity-dependent histone H3 methyl/phos switching to maintain repetitive element silencing in neurons. Proc Natl Acad Sci U S A 112, 6820–7 (2015).

286. Liu, S.Y. & Ikegami, K. Nuclear lamin phosphorylation: an emerging role in gene regulation and pathogenesis of laminopathies. Nucleus 11, 299–314 (2020).

287. Wesley, C.C., North, D.V. & Levy, D.L. Protein kinase C activity modulates nuclear Lamin A/C dynamics in HeLa cells. Sci Rep 14, 6388 (2024).

288. Nava, M.M. et al. Heterochromatin-Driven Nuclear Softening Protects the Genome against Mechanical Stress-Induced Damage. Cell 181, 800–817 e22 (2020).

289. Krigerts, J. et al. Differentiating cancer cells reveal early large-scale genome regulation by pericentric domains. Biophys J 120, 711–724 (2021).

290. Morales Torres, C. et al. Selective inhibition of cancer cell self-renewal through a Quisinostat-histone H1.0 axis. Nat Commun 11, 1792 (2020).

291. Bagert, J.D. et al. Oncohistone mutations enhance chromatin remodeling and alter cell fates. Nat Chem Biol 17, 403–411 (2021).

292. Dyer, M.A., Qadeer, Z.A., Valle-Garcia, D. & Bernstein, E. ATRX and DAXX: Mechanisms and Mutations. Cold Spring Harb Perspect Med 7(2017).

293. Carraro, M. et al. DAXX adds a de novo H3.3K9me3 deposition pathway to the histone chaperone network. Mol Cell 83, 1075–1092 e9 (2023).

294. Cammareri, P. et al. Loss of colonic fidelity enables multilineage plasticity and metastasis. Nature (2025).

295. Ni, K. et al. LSH mediates gene repression through macroH2A deposition. Nat Commun 11, 5647 (2020).

296. Lee, S.C. et al. Chromatin remodeling of histone H3 variants by DDM1 underlies epigenetic inheritance of DNA methylation. Cell 186, 4100–4116 e15 (2023).

297. Guckelberger, P., Haut, L., Tornisiello, R., Kretzmer, H. & Meissner, A. HELLS is required for maintaining proper DNA modification at human satellite repeats. Genome Biol 26, 211 (2025).

298. Salinas-Luypaert, C. et al. DNA methylation influences human centromere positioning and function. Nat Genet 57, 2509–2521 (2025).

299. Bintu, L. et al. Dynamics of epigenetic regulation at the single-cell level. Science 351, 720–4 (2016).

300. Househam, J. et al. Phenotypic plasticity and genetic control in colorectal cancer evolution. Nature 611, 744–753 (2022).

301. Mathur, R. et al. Glioblastoma evolution and heterogeneity from a 3D whole-tumor perspective. Cell 187, 446–463 e16 (2024).

302. Gorbunova, V. et al. The role of retrotransposable elements in ageing and age-associated diseases. Nature 596, 43–53 (2021).

303. Simon, M. et al. A rare human centenarian variant of SIRT6 enhances genome stability and interaction with Lamin A. EMBO J 42, e113326 (2023).

304. Morandini, F. et al. Transposable element 5mC methylation state of blood cells predicts age and disease. Nat Aging 5, 193–204 (2025).

305. Amaral, M.L. et al. Single-Cell Epigenomics Uncovers Heterochromatin Instability and Transcription Factor Dysfunction during Mouse Brain Aging. bioRxiv, 2025.04.21.649585 (2025).

306. Pastushenko, I. et al. Identification of the tumour transition states occurring during EMT. Nature 556, 463–468 (2018).

307. Parreno, V. et al. Transient loss of Polycomb components induces an epigenetic cancer fate. Nature 629, 688–696 (2024).

308. Bersani, F. et al. Pericentromeric satellite repeat expansions through RNA-derived DNA intermediates in cancer. Proc Natl Acad Sci U S A 112, 15148–53 (2015).

309. Pachva, M.C., Lai, H., Jia, A., Rouleau, M. & Sorensen, P.H. Extracellular Vesicles in Reprogramming of the Ewing Sarcoma Tumor Microenvironment. Front Cell Dev Biol 9, 726205 (2021).

310. Ariffen, N.A. et al. Amplification of different satellite-DNAs in prostate cancer. Pathol Res Pract 256, 155269 (2024).

311. Jassim, A., Rahrmann, E.P., Simons, B.D. & Gilbertson, R.J. Cancers make their own luck: theories of cancer origins. Nat Rev Cancer 23, 710–724 (2023).

312. Karr, J.P., Ferrie, J.J., Tjian, R. & Darzacq, X. The transcription factor activity gradient (TAG) model: contemplating a contact-independent mechanism for enhancer-promoter communication. Genes Dev 36, 7–16 (2022).

313. Cullen, K.E., Kladde, M.P. & Seyfred, M.A. Interaction between transcription regulatory regions of prolactin chromatin. Science 261, 203–6 (1993).

314. Zaugg, J.B. et al. Current challenges in understanding the role of enhancers in disease. Nat Struct Mol Biol (2022).

315. Fourel, G. et al. Evidence for long-range oncogene activation by hepadnavirus insertion. EMBO J 13, 2526–34 (1994).

316. Walters, M.C. et al. Transcriptional enhancers act in cis to suppress position-effect variegation. Genes Dev 10, 185–95 (1996).

317. Fourel, G. et al. An activation-independent role of transcription factors in insulator function. EMBO Rep 2, 124–32 (2001).

318. Miron, E. et al. Chromatin arranges in chains of mesoscale domains with nanoscale functional topography independent of cohesin. Sci Adv 6(2020).

319. Minami, K. et al. Replication-dependent histone labeling dissects the physical properties of euchromatin/heterochromatin in living human cells. Sci Adv 11, eadu8400 (2025).

320. Fritton, H.P., Sippel, A.E. & Igo-Kemenes, T. Nuclease-hypersensitive sites in the chromatin domain of the chicken lysozyme gene. Nucleic Acids Res 11, 3467–85 (1983).

321. Elgin, S.C. Chromatin structure and gene activity. Curr Opin Cell Biol 2, 437–45 (1990).

322. Girdhar, K. et al. Chromatin domain alterations linked to 3D genome organization in a large cohort of schizophrenia and bipolar disorder brains. Nat Neurosci 25, 474–483 (2022).

323. Factor, D.C. et al. Cell Type-Specific Intralocus Interactions Reveal Oligodendrocyte Mechanisms in MS. Cell 181, 382–395 e21 (2020).

324. Chatterjee, S. et al. Enhancer Variants Synergistically Drive Dysfunction of a Gene Regulatory Network In Hirschsprung Disease. Cell 167, 355–368 e10 (2016).

325. Zhou, S. et al. Noncoding mutations target cis-regulatory elements of the FOXA1 plexus in prostate cancer. Nat Commun 11, 441 (2020).

326. Morris, J.A. et al. Discovery of target genes and pathways at GWAS loci by pooled single-cell CRISPR screens. Science, eadh7699 (2023).

327. Connally, N.J. et al. The missing link between genetic association and regulatory function. Elife 11(2022).

328. Judd, J., Sanderson, H. & Feschotte, C. Evolution of mouse circadian enhancers from transposable elements. Genome Biol 22, 193 (2021).

329. Neumayr, C. et al. Differential cofactor dependencies define distinct types of human enhancers. Nature (2022).

330. Zeller, P. et al. Single-cell sortChIC identifies hierarchical chromatin dynamics during hematopoiesis. Nat Genet 55, 333–345 (2023).

331. Zuckerkandl, E. A possible role of “inert” heterochromatin in cell differentiation. Action of and competition for “locking” molecules. Biochimie 56, 937–54 (1974).

332. Zuckerkandl, E. & Cavalli, G. Combinatorial epigenetics, “junk DNA”, and the evolution of complex organisms. Gene 390, 232–42 (2007).

333. Zuckerkandl, E. Junk DNA and sectorial gene repression. Gene 205, 323–43 (1997).

334. Benabdallah, N.S. et al. Decreased Enhancer-Promoter Proximity Accompanying Enhancer Activation. Mol Cell 76, 473–484 e7 (2019).

335. Bulut-Karslioglu, A. et al. A transcription factor-based mechanism for mouse heterochromatin formation. Nat Struct Mol Biol 19, 1023–30 (2012).

336. Velazquez Camacho, O. et al. Major satellite repeat RNA stabilize heterochromatin retention of Suv39h enzymes by RNA-nucleosome association and RNA:DNA hybrid formation. Elife 6(2017).

337. Thakur, J., Packiaraj, J. & Henikoff, S. Sequence, Chromatin and Evolution of Satellite DNA. Int J Mol Sci 22(2021).

338. Feng, Y. et al. Simultaneous epigenetic perturbation and genome imaging reveal distinct roles of H3K9me3 in chromatin architecture and transcription. Genome Biol 21, 296 (2020).

339. Noma, K., Cam, H.P., Maraia, R.J. & Grewal, S.I. A role for TFIIIC transcription factor complex in genome organization. Cell 125, 859–72 (2006).

340. Hiraga, S., Botsios, S., Donze, D. & Donaldson, A.D. TFIIIC localizes budding yeast ETC sites to the nuclear periphery. Mol Biol Cell 23, 2741–54 (2012).

341. Yokoshi, M., Segawa, K. & Fukaya, T. Visualizing the Role of Boundary Elements in Enhancer-Promoter Communication. Mol Cell 78, 224–235 e5 (2020).

342. Stutzman, A.V., Liang, A.S., Beilinson, V. & Ikegami, K. Transcription-independent TFIIIC-bound sites cluster near heterochromatin boundaries within lamina-associated domains in C. elegans. Epigenetics Chromatin 13, 1 (2020).

343. Kundu, T.K., Wang, Z. & Roeder, R.G. Human TFIIIC relieves chromatin-mediated repression of RNA polymerase III transcription and contains an intrinsic histone acetyltransferase activity. Mol Cell Biol 19, 1605–15 (1999).

344. Kaaij, L.J.T., Mohn, F., van der Weide, R.H., de Wit, E. & Buhler, M. The ChAHP Complex Counteracts Chromatin Looping at CTCF Sites that Emerged from SINE Expansions in Mouse. Cell 178, 1437–1451 e14 (2019).

345. Ahel, J. et al. ChAHP2 and ChAHP control diverse retrotransposons by complementary activities. bioRxiv, 2024.02.05.578923 (2024).

346. Ahel, J. et al. ChAHP2 and ChAHP control diverse retrotransposons by complementary activities. Genes Dev 38, 554–568 (2024).

347. Siddaway, R. et al. The in vivo Interaction Landscape of Histones H3.1 and H3.3. Mol Cell Proteomics 21, 100411 (2022).

348. Oughtred, R. et al. The BioGRID database: A comprehensive biomedical resource of curated protein, genetic, and chemical interactions. Protein Sci 30, 187–200 (2021).

349. Olsen, J.V. et al. Global, in vivo, and site-specific phosphorylation dynamics in signaling networks. Cell 127, 635–48 (2006).

350. Reverendo, M. et al. Polymerase III transcription is necessary for T cell priming by dendritic cells. Proc Natl Acad Sci U S A 116, 22721–22729 (2019).

351. Linker, S.B. et al. Identification of bona fide B2 SINE retrotransposon transcription through single-nucleus RNA-seq of the mouse hippocampus. Genome Res 30, 1643–1654 (2020).

352. Oki, M., Valenzuela, L., Chiba, T., Ito, T. & Kamakaka, R.T. Barrier proteins remodel and modify chromatin to restrict silenced domains. Mol Cell Biol 24, 1956–67 (2004).

353. Cotterman, R. et al. N-Myc regulates a widespread euchromatic program in the human genome partially independent of its role as a classical transcription factor. Cancer Res 68, 9654–62 (2008).

354. Lu, J.Y. et al. Genomic Repeats Categorize Genes with Distinct Functions for Orchestrated Regulation. Cell Rep 30, 3296–3311 e5 (2020).

355. Travers, A.A., Vaillant, C., Arneodo, A. & Muskhelishvili, G. DNA structure, nucleosome placement and chromatin remodelling: a perspective. Biochem Soc Trans 40, 335–40 (2012).

356. Abdulhay, N.J. et al. Massively multiplex single-molecule oligonucleosome footprinting. Elife 9(2020).

357. Kilic, S. et al. Single-molecule FRET reveals multiscale chromatin dynamics modulated by HP1alpha. Nat Commun 9, 235 (2018).

358. Scacchetti, A. et al. CHRAC/ACF contribute to the repressive ground state of chromatin. Life Sci Alliance 1, e201800024 (2018).

359. Takahata, S. et al. Two secured FACT recruitment mechanisms are essential for heterochromatin maintenance. Cell Rep 36, 109540 (2021).

360. Murawska, M. et al. The histone chaperone FACT facilitates heterochromatin spreading by regulating histone turnover and H3K9 methylation states. Cell Rep 37, 109944 (2021).

361. Xue, Y. et al. Mot1, Ino80C, and NC2 Function Coordinately to Regulate Pervasive Transcription in Yeast and Mammals. Mol Cell 67, 594–607 e4 (2017).

362. Ranjan, A. et al. Live-cell single particle imaging reveals the role of RNA polymerase II in histone H2A.Z eviction. Elife 9(2020).

363. Zhang, C. et al. H3K56 deacetylation and H2A.Z deposition are required for aberrant heterochromatin spreading. Nucleic Acids Res 50, 3852–3866 (2022).

364. Imre, L. et al. Fundamental Role Of The H2A.Z C-Terminal Tail In The Formation Of Constitutive Heterochromatin. bioRxiv, 2021.02.22.432230 (2021).

365. Tartour, K. et al. Mammalian PERIOD2 regulates H2A.Z incorporation in chromatin to orchestrate circadian negative feedback. Nat Struct Mol Biol 29, 549–562 (2022).

366. Hardy, S. & Robert, F. Random deposition of histone variants: A cellular mistake or a novel regulatory mechanism? Epigenetics 5, 368–72 (2010).

367. Chen, Z. et al. High-resolution and high-accuracy topographic and transcriptional maps of the nucleosome barrier. Elife 8(2019).

368. Collings, C.K. & Anderson, J.N. Links between DNA methylation and nucleosome occupancy in the human genome. Epigenetics Chromatin 10, 18 (2017).

369. Fourel, G., Miyake, T., Defossez, P.A., Li, R. & Gilson, E. General regulatory factors (GRFs) as genome partitioners. J Biol Chem 277, 41736–43 (2002).

370. Dion, M.F. et al. Dynamics of replication-independent histone turnover in budding yeast. Science 315, 1405–8 (2007).

371. Dodd, I.B. & Sneppen, K. Barriers and silencers: a theoretical toolkit for control and containment of nucleosome-based epigenetic states. J Mol Biol 414, 624–37 (2011).

372. McClintock, B. The significance of responses of the genome to challenge. Science 226, 792–801 (1984).

373. Kaplow, I.M. et al. Relating enhancer genetic variation across mammals to complex phenotypes using machine learning. Science 380, eabm7993 (2023).

374. Griffin, G.K. et al. Epigenetic silencing by SETDB1 suppresses tumour intrinsic immunogenicity. Nature 595, 309–314 (2021).

375. Soares, M.L. et al. Targeted deletion of a 170-kb cluster of LINE-1 repeats and implications for regional control. Genome Res 28, 345–56 (2018).

376. Kvikstad, E.M. & Makova, K.D. The (r)evolution of SINE versus LINE distributions in primate genomes: sex chromosomes are important. Genome Res 20, 600–13 (2010).

377. Fedoroff, N.V. Presidential address. Transposable elements, epigenetics, and genome evolution. Science 338, 758–67 (2012).

378. Zhu, Y., van Essen, D. & Saccani, S. Cell-type-specific control of enhancer activity by H3K9 trimethylation. Mol Cell 46, 408–23 (2012).

379. Yelagandula, R. et al. ZFP462 safeguards neural lineage specification by targeting G9A/GLP-mediated heterochromatin to silence enhancers. Nat Cell Biol (2023).

380. Tomkova, M. et al. Human DNA polymerase epsilon is a source of C>T mutations at CpG dinucleotides. Nat Genet (2024).

381. Hoyt, S.J. et al. From telomere to telomere: The transcriptional and epigenetic state of human repeat elements. Science 376, eabk3112 (2022).

382. Lanciano, S. et al. Locus-level L1 DNA methylation profiling reveals the epigenetic and transcriptional interplay between L1s and their integration sites. Cell Genom, 100498 (2024).

383. Mehdipour, P. et al. Epigenetic therapy induces transcription of inverted SINEs and ADAR1 dependency. Nature 588, 169–173 (2020).

384. Deniz, O., Frost, J.M. & Branco, M.R. Regulation of transposable elements by DNA modifications. Nat Rev Genet 20, 417–431 (2019).

385. Nishibuchi, G. & Dejardin, J. The molecular basis of the organization of repetitive DNA-containing constitutive heterochromatin in mammals. Chromosome Res 25, 77–87 (2017).

386. Muller, I. & Helin, K. Keep quiet: the HUSH complex in transcriptional silencing and disease. Nat Struct Mol Biol 31, 11–22 (2024).

387. Wang, L. et al. Histone Modifications Regulate Chromatin Compartmentalization by Contributing to a Phase Separation Mechanism. Mol Cell 76, 646–659 e6 (2019).

388. Liu, N. et al. Selective silencing of euchromatic L1s revealed by genome-wide screens for L1 regulators. Nature 553, 228–232 (2018).

389. Tan, K. et al. The Rhox gene cluster suppresses germline LINE1 transposition. Proc Natl Acad Sci U S A 118(2021).

390. Shaban, H.A., Barth, R., Recoules, L. & Bystricky, K. Hi-D: nanoscale mapping of nuclear dynamics in single living cells. Genome Biol 21, 95 (2020).

391. Zhang, W. et al. METTL3-dependent m6A RNA methylation regulates transposable elements and represses human naive pluripotency through transposable element-derived enhancers. Nucleic Acids Res 53(2025).

392. Li, Y., Chen, X. & Lu, C. The interplay between DNA and histone methylation: molecular mechanisms and disease implications. EMBO Rep 22, e51803 (2021).

393. Buckberry, S. et al. Transient naive reprogramming corrects hiPS cells functionally and epigenetically. Nature (2023).

394. Clark, S.J. et al. Single-cell multi-omics profiling links dynamic DNA methylation to cell fate decisions during mouse early organogenesis. Genome Biol 23, 202 (2022).

395. Francastel, C. & Magdinier, F. DNA methylation in satellite repeats disorders. Essays Biochem 63, 757–771 (2019).

396. Guo, H. et al. The DNA methylation landscape of human early embryos. Nature 511, 606–10 (2014).

397. Argelaguet, R. et al. Multi-omics profiling of mouse gastrulation at single-cell resolution. Nature 576, 487–491 (2019).

398. Gopi, L.K. & Kidder, B.L. Integrative pan cancer analysis reveals epigenomic variation in cancer type and cell specific chromatin domains. Nat Commun 12, 1419 (2021).

399. Souaifan, H. et al. Targeting of the nuclear RNA exosome to chromatin by HP1 affects the transcriptional programs of liver cells. bioRxiv, 2025.02.14.638307 (2025).

400. Martins, F. et al. A Cluster of Evolutionarily Recent KRAB Zinc Finger Proteins Protects Cancer Cells from Replicative Stress-Induced Inflammation. Cancer Res 84, 808–826 (2024).

401. Cuellar, T.L. et al. Silencing of retrotransposons by SETDB1 inhibits the interferon response in acute myeloid leukemia. J Cell Biol 216, 3535–3549 (2017).

402. Guler, G.D. et al. Repression of Stress-Induced LINE-1 Expression Protects Cancer Cell Subpopulations from Lethal Drug Exposure. Cancer Cell 32, 221–237 e13 (2017).

403. Moore, P.C., Henderson, K.W. & Classon, M. The epigenome and the many facets of cancer drug tolerance. Adv Cancer Res 158, 1–39 (2023).

404. Baratchian, M. et al. H3K9 methylation drives resistance to androgen receptor-antagonist therapy in prostate cancer. Proc Natl Acad Sci U S A 119, e2114324119 (2022).

405. Canat, A. et al. DAXX safeguards heterochromatin formation in embryonic stem cells. bioRxiv, 2021.04.28.441827 (2023).

406. Lindholm, H.T., Chen, R. & De Carvalho, D.D. Endogenous retroelements as alarms for disruptions to cellular homeostasis. Trends Cancer (2022).

407. Fan, D.N. et al. Histone lysine methyltransferase, suppressor of variegation 3-9 homolog 1, promotes hepatocellular carcinoma progression and is negatively regulated by microRNA-125b. Hepatology 57, 637–47 (2013).

408. Gu, Z. et al. Silencing of LINE-1 retrotransposons is a selective dependency of myeloid leukemia. Nat Genet 53, 672–682 (2021).

409. Liu, Y. et al. DNA hypomethylation promotes UHRF1-and SUV39H1/H2-dependent crosstalk between H3K18ub and H3K9me3 to reinforce heterochromatin states. Mol Cell 85, 394–412 e12 (2025).

410. Deblois, G. et al. Epigenetic Switch-Induced Viral Mimicry Evasion in Chemotherapy-Resistant Breast Cancer. Cancer Discov (2020).

411. Liang, L. et al. Complementary Alu sequences mediate enhancer-promoter selectivity. Nature 619, 868–875 (2023).

412. Virstedt, J., Berge, T., Henderson, R.M., Waring, M.J. & Travers, A.A. The influence of DNA stiffness upon nucleosome formation. J Struct Biol 148, 66–85 (2004).

413. Vinayachandran, V., Shivaswamy, S., Shukla, A., Kendyala, N. & Bhargava, P. Terminator-dependent facilitated recycling of RNA polymerase III couples transcriptional activation and chromatin remodeling in vivo. bioRxiv, 2023.02.28.530554 (2023).

414. Tramantano, M. et al. Constitutive turnover of histone H2A.Z at yeast promoters requires the preinitiation complex. Elife 5(2016).

415. Drillon, G., Audit, B., Argoul, F. & Arneodo, A. Evidence of selection for an accessible nucleosomal array in human. BMC Genomics 17, 526 (2016).

416. Brunet, F.G. et al. Evidence for DNA Sequence Encoding of an Accessible Nucleosomal Array across Vertebrates. Biophys J 114, 2308–2316 (2018).

417. Collings, C.K., Waddell, P.J. & Anderson, J.N. Effects of DNA methylation on nucleosome stability. Nucleic Acids Res 41, 2918–31 (2013).

418. Roulois, D. et al. DNA-Demethylating Agents Target Colorectal Cancer Cells by Inducing Viral Mimicry by Endogenous Transcripts. Cell 162, 961–73 (2015).

419. Zapatka, M. et al. The landscape of viral associations in human cancers. Nat Genet 52, 320–330 (2020).

420. Colombo, A.R., Triche, T., Jr. & Ramsingh, G. Transposable Element Expression in Acute Myeloid Leukemia Transcriptome and Prognosis. Sci Rep 8, 16449 (2018).

421. Porter, R.L. et al. Satellite repeat RNA expression in epithelial ovarian cancer associates with a tumor-immunosuppressive phenotype. J Clin Invest 132(2022).

422. Deniz, O. et al. Endogenous retroviruses are a source of enhancers with oncogenic potential in acute myeloid leukaemia. Nat Commun 11, 3506 (2020).

423. Larsen, F. et al. Viral mimicry acts as a tumor suppressor in colitis. Nat Commun 17, 1313 (2026).

424. Colombo, A.R. et al. Suppression of Transposable Elements in Leukemic Stem Cells. Sci Rep 7, 7029 (2017).

425. Torres-Padilla, M.E. On transposons and totipotency. Philos Trans R Soc Lond B Biol Sci 375, 20190339 (2020).

426. Lawson, K.A. et al. Functional genomic landscape of cancer-intrinsic evasion of killing by T cells. Nature 586, 120–126 (2020).

427. Hosseini, A. et al. Retroelement decay by the exonuclease XRN1 is a viral mimicry dependency in cancer. Cell Rep 43, 113684 (2024).

428. Zhao, Y. et al. Transposon-triggered innate immune response confers cancer resistance to the blind mole rat. Nat Immunol 22, 1219–1230 (2021).

429. Lamers, M.M., van den Hoogen, B.G. & Haagmans, B.L. ADAR1: “Editor-in-Chief” of Cytoplasmic Innate Immunity. Front Immunol 10, 1763 (2019).

430. Guo, Y.L. The underdeveloped innate immunity in embryonic stem cells: The molecular basis and biological perspectives from early embryogenesis. Am J Reprod Immunol 81, e13089 (2019).

431. Poirier, E.Z. et al. An isoform of Dicer protects mammalian stem cells against multiple RNA viruses. Science 373, 231–236 (2021).

432. Chung, H. et al. Human ADAR1 Prevents Endogenous RNA from Triggering Translational Shutdown. Cell 172, 811–824 e14 (2018).

433. Bonnin, M. et al. Toll-like receptor 3 downregulation is an escape mechanism from apoptosis during hepatocarcinogenesis. J Hepatol 71, 763–772 (2019).

434. Wu, M.J. et al. Mutant IDH1 inhibition induces dsDNA sensing to activate tumor immunity. Science 385, eadl6173 (2024).

435. Chen, R., Ishak, C.A. & De Carvalho, D.D. Endogenous Retroelements and the Viral Mimicry Response in Cancer Therapy and Cellular Homeostasis. Cancer Discov 11, 2707–2725 (2021).

436. Loo Yau, H., Ettayebi, I. & De Carvalho, D.D. The Cancer Epigenome: Exploiting Its Vulnerabilities for Immunotherapy. Trends Cell Biol 29, 31–43 (2019).

437. Sheng, W. et al. LSD1 Ablation Stimulates Anti-tumor Immunity and Enables Checkpoint Blockade. Cell 174, 549–563 e19 (2018).

438. Jang, H.J. et al. Epigenetic therapy potentiates transposable element transcription to create tumor-enriched antigens in glioblastoma cells. Nat Genet 56, 1903–1913 (2024).

439. Huang, D., Petrykowska, H.M., Miller, B.F., Elnitski, L. & Ovcharenko, I. Identification of human silencers by correlating cross-tissue epigenetic profiles and gene expression. Genome Res 29, 657–667 (2019).

440. Boyle, E.I. et al. GO::TermFinder--open source software for accessing Gene Ontology information and finding significantly enriched Gene Ontology terms associated with a list of genes. Bioinformatics 20, 3710–5 (2004).

441. Cheetham, S.W., Kindlova, M. & Ewing, A.D. Methylartist: tools for visualizing modified bases from nanopore sequence data. Bioinformatics 38, 3109–3112 (2022).

442. Lee, B.T. et al. The UCSC Genome Browser database: 2022 update. Nucleic Acids Res 50, D1115–D1122 (2022).

443. Castro-Diaz, N. et al. Evolutionally dynamic L1 regulation in embryonic stem cells. Genes Dev 28, 1397–409 (2014).

444. Bulyk, M.L., Drouin, J., Harrison, M.M., Taipale, J. & Zaret, K.S. Pioneer factors - key regulators of chromatin and gene expression. Nat Rev Genet 24, 809–815 (2023).

445. Kubo, N. et al. H3K4me1 facilitates promoter-enhancer interactions and gene activation during embryonic stem cell differentiation. Mol Cell 84, 1742–1752 e5 (2024).

446. Bozek, M. & Gompel, N. Developmental Transcriptional Enhancers: A Subtle Interplay between Accessibility and Activity: Considering Quantitative Accessibility Changes between Different Regulatory States of an Enhancer Deconvolutes the Complex Relationship between Accessibility and Activity. Bioessays 42, e1900188 (2020).

447. Loyfer, N. et al. A DNA methylation atlas of normal human cell types. Nature 613, 355–364 (2023).

