## Supplementary.Text, Figures_Graphical.Abstract for "ProA and ProB repeat sequences shape 3D genome organization in eukaryotes"

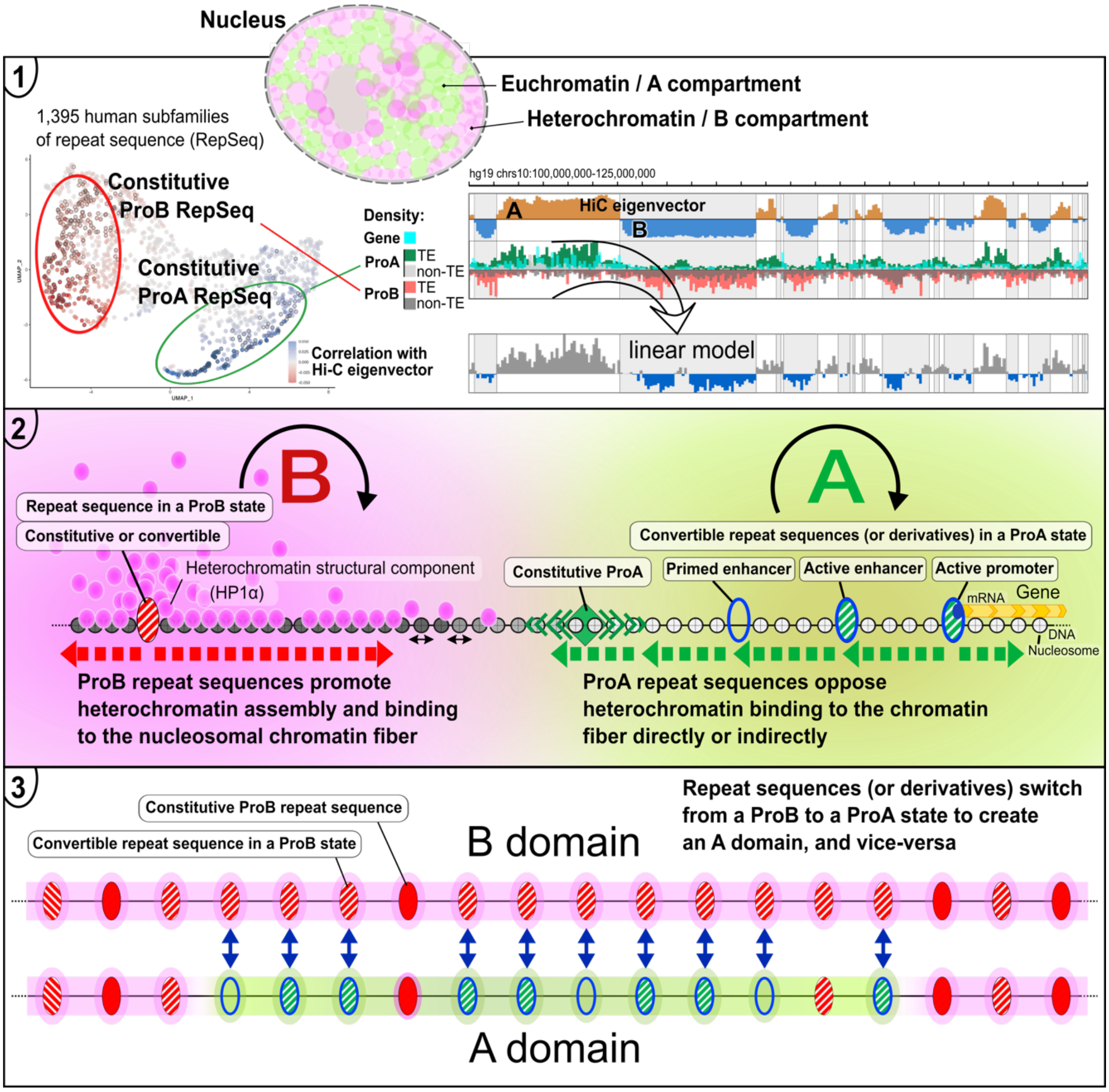

**Repeat sequences (RepSeqs) shape genome organization by driving A/B compartmentalization.**

(1) We identify sets of RepSeqs (TE and non-TE) whose genomic distribution is sufficient to predict A/B compartment profiles genome-wide (as captured by the Hi-C eigenvector).

(2) These RepSeqs are functionally antagonistic: ProB elements promote the assembly and spreading of heterochromatin, whereas ProA elements oppose heterochromatin spreading. Self-reinforcing interaction networks stabilize A and B chromatin states (circular arrows), forming a molecular toggle switch.

(3) While some RepSeqs are constitutively ProA or ProB, others ("convertible" RepSeqs, best known as cis-regulatory elements for gene transcription, or enhancers) can switch states and cooperatively fold or unfold chromatin domains, thereby modulating gene transcription at the level of genome organization.

SUPPLEMENTARY TEXT

**Comments on Extended Data Fig. 2:**
***GO analysis***
In order to determine whether individual chromatin subclasses display functional specializations, a GO analysis was performed for gene-containing subclasses 1-16 as defined in our study (using data obtained with HUVEC cells). GO terms referred to herein as "enriched GO terms" (or eGO) are those for which enrichment reaches significance ($P < 0.05$) at the level of the subclass considered, amounting to 1938 GO terms in total for the 16 subclasses (Extended Data Fig. 1C). To address whether genes embedded in individual bin subclasses display any specific patterns with regard to functional specialization, we created a list of 34 terms that are more cross-sectional than GO terms, dubbed "GF-GO.slim", such that it is possible to assign a unique GF-GO.slim to each GO term by following a procedure described in the Resources and Methods section. The GF-GO.slim are divided into the following 4 clades: I (GF-GO.slim terms 01-04): Crosscutting principles; II (05-16): Basic cell functions (housekeeping), such as metabolism and intracellular transport (also includes cell cycle, apoptosis, autophagy); III (17-21): Basic cell function associated with a particular cellular substructure, possibly associated with a specialization; IV (22-34): Cell communication with its environment (23,24), chromatin and transcription (25), development and differentiation, and differentiated cell functions, in particular related to Immunity (34). Clade IV thus encompasses what may be called "evolutionarily specialized function", as opposed to housekeeping functions, which are represented, most prominently, in clade II, and to some extent also in clade III.

Our GO analysis shows that GF-GO.slim belonging to clade II, and to some extent clades I and III, which broadly speaking correspond to housekeeping functions or processes at work in most tissues, clearly appear overrepresented among genes in AlwaysA bin subclasses (Extended Data Fig. 2B, dark green bars). In contrast, clade IV GF-GO.slim, which are associated with evolutionarily evolved functions and in particular tissue-specific functions, appear enriched in more diverse genomic contexts, and more specifically in genes present in AorB regions (Extended Data Fig. 2B, light green and light pink bars). This differential distribution of housekeeping and context-specific genes coincides with a differential composition and *modus operandi* of associated gene promoters. Thus, housekeeping genes have promoters containing CpG islands (CGI) and are regulated by Polycomb/H3K27me3-based heterochromatin, whereas AorB regions are enriched in genes with CGI-minus promoters, for which regulation involves HP1/H3K9me3-based heterochromatin and CpG methylation[1,2]. In addition, transcription cofactors are distinct for the two types of promoters [1,2].

***Gene complexes***
Gene-containing bin subclasses falling in AlwaysB (subclasses 7-8 and 15-16), which represent 7% of the total number of genes, as well as AorB:B high gene density bin subclasses (subclasses 5-6), display high g.eGO/g.GO ratios, suggestive of strong specialization. These bins overlap with "gene complexes", which are loci that emerged through the successive duplication of an ancestral segment containing one or more genes [3]. This may account for both the high gene density and the specialization observed in these regions. This is the case for the loci encoding KZFPs, found on different chromosomes but enriched on chromosome 19. KZFPs are TFs chiefly involved in TE repression, hence the unique GF-GO.slim emerging for bin subclass 7: "25-chromatin, transcription, chromosome". This is also the case for various loci involved in immunity ("34-Immunity"). Gene expression within these loci is strongly regulated, involving cooperation between the two main types of heterochromatin, based on HP1/H3K9me3 and Polycomb/H3K27me3, respectively, and an abundance of both ProA and ProB RepSeq (see Extended Data Figs. 2-3). Within gene complexes, isolated active genes may be observed within an otherwise inactive and strongly heterochromatin-marked locus, consistent with the seemingly paradoxical “AlwaysB, Active” character of bin subclass 7. Some gene complexes display regular gene densities, such as the olfactory receptor gene complexes, and a number of complexes encoding other cell membrane components (e.g. G-protein coupled receptors other than olfactory receptors (GPCRs)). Some of these gene-complex regions score as B in our analysis because HUVEC are endothelial cells, whereas olfactory receptor genes, for instance, are essentially only expressed in the olfactory mucosa.

**Comments on Extended Data Fig. 3 and Table S2: Correlation with Hi-C EV for RepSeq subfamilies, calculated in seven other cell lines in addition to HUVEC, shows that values are generally similar in different cell lines**
In a Euclidean plane with HUVEC values on the x axis, the points are distributed around a regression line close to the diagonal. For the majority of points, deviation from the diagonal is within a 0.015-unit-wide stripe, i.e., the deviations from the values obtained in HUVEC are below 0.007. Therefore, the subfamily composition of the CorrA and CorrB sets would not have been very different had they been generated using data from a line other than HUVEC.

Variability is seemingly higher in the range (-0.01/+0.01), which is the range we excluded when defining the corrA and corrB sets. However, this is partly an optical effect due to the fact that many RepSeq subfamilies are concentrated in this range (approximately 50% for HUVEC), in particular subfamilies with low copy numbers and therefore imprecise Corr.Hi-C EV calculations. On the other hand, it is interesting to note that even in this range where the corr.Hi-C EV for most RepSeq subfamilies displays P-values above 0.05 , the distribution of a majority of points remains within the 0.015 unit-wide stripe, such that these RepSeq families having no clear ProA or ProB character still can be said to display a "reproducibly undefined" character. Of note, for almost every cell line there are a few points across the whole distribution that clearly deviate from the regression line, although it is never a marked shift, suggesting that the corresponding RepSeq subfamilies have distinctive functional characteristics depending on the cell context, displaying either a more ProA or a more ProB character in the corresponding cell line compared to HUVEC.

The same conclusion can be reached for these data by using the original table (Table S2) with an appropriate color code. Thus, corrB RepSeq as calculated in HUVEC ("sharp" colors, red for corr.Hi-C EV below -0.05; yellow for corr.Hi-C EV between -0.05 and -0.01) do not appear on side A (corr.Hi-C EV > 0 in the column in question, colors tending towards green), with very rare exceptions but which often correspond to cases not reaching the significance threshold of P <0.05. Similarly, corrA RepSeq ("sharp" colors, ultramarine green for corr.Hi-C EV above 0.05; almond green for corr.Hi-C.EV between 0.01 and 0.05) do not appear on side B (corr.Hi-C EV below 0 in the column in question). On the contrary, in the intermediate zone (corr.Hi-CEV range -0.01/0.01, "grayed" greenish and reddish colors), colors are much more mixed. This can also be clearly seen if the analysis is limited to ERVs, which make up a large proportion of subfamilies in the intermediate range. Many of these subfamilies have counts below 1000, but there are also subfamilies with counts above 1000 and possibly displaying "disparate" correlations. For instance:

- MER65A (line 406), 1,413 copies/hg19, scores on A side (P<0.01) in GM12878, and on B side (P<0.05) in IMR90, and nothing clear otherwise in the other lines.
- LTR5 Hs (line 458), 645 copies hg19, scores on A side in GM12878, KBM7 and NHEK (P<0.05), and nothing clear otherwise in the other lines.

Here it should be remembered that most corr.Hi-C EV points in the range -0.01/0.01 display P>0.05.

It is notable that regression lines in Extended Data Fig. 3 do not have the exact same slope across all cell lines (upper left panels). This is likely due at least in part to the fact that the degree of separation between A and B compartments ("compartmentalization") can be quite different between cell lines, which impacts the Hi-C EV values and therefore the value of the calculated correlation. In order to be able to use this combinatorial representation while taking this point into account, we rotated panels around the origin (0;0) so as to superimpose the 7 regression lines (right panel). As can be seen, almost all the points fall within a band width of approx. 0.015 units around the obtained diagonal.

**Comments on Extended Data Fig. 6: Evidence for a specific functional role of CpG methylation at Alu and SVA TEs**

Regression lines indicate that there is a general trend for individual TE subfamilies by which the higher the GC content, the higher the CpG content. LINEs and DNA TEs are essentially AT-rich, whereas ERVs, and in particular the LTRs of young ERVs, are more GC- and CpG-rich. Alu TEs and even more so SVA are extremely GC-rich, with both Alu and SVA subfamilies each displaying a peculiar regime for the %GC to %CpG ratio, by comparison with other TE subclasses, suggestive of a specific role of CpG dinucleotides. There is also a trend by which the higher the CpG content, the higher the DNA methylation. Again, Alu and SVA subclasses are outliers, displaying extremely high methylation levels, suggesting that the specific role of CpG at Alu and SVA may depend on CpG methylation. Young ERVs tend to harbor a high CpG content while displaying remarkably variable CpG methylation levels. Young ERVs with an exceptionally high CpG content, close to the CpG content of CpG islands, such as LTR12C and HERH-int, tend to display variable CpG methylation between individuals, which may be explained by variation in KZFP activity opposing CpG island-type activator mechanisms that maintain a demethylated status by recruiting TET enzymes [4].

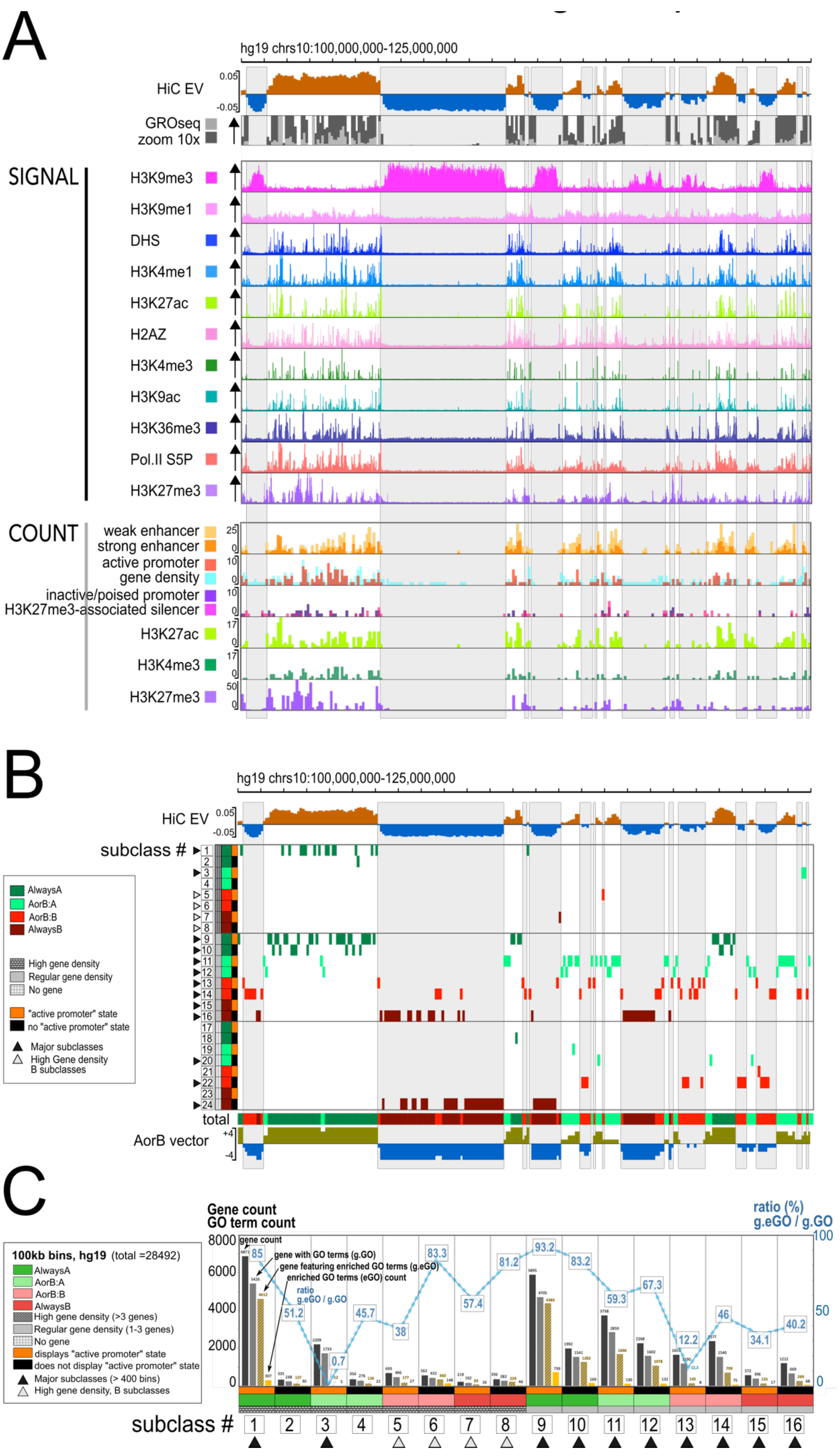
A
hg19 chrs10:100,000,000-125,000,000
HiC EV
0.05
-0.05
GROseq
zoom 10x
SIGNAL
H3K9me3
H3K9me1
DHS
H3K4me1
H3K27ac
H2AZ
H3K4me3
H3K9ac
H3K36me3
Pol.II S5P
H3K27me3
COUNT
weak enhancer
strong enhancer
active promoter
gene density
inactive/poised promoter
H3K27me3-associated silencer
H3K27ac
H3K4me3
H3K27me3
B
hg19 chrs10:100,000,000-125,000,000
HiC EV
subclass #
AlwaysA
AorB:A
AorB:B
AlwaysB
High gene density
Regular gene density
No gene
"active promoter" state
no "active promoter" state
Major subclasses
High Gene density B subclasses
total
AorB vector
C
Gene count
GO term count
ratio (%) g.eGO / g.GO
gene count
gene with GO terms (g.GO)
gene featuring enriched GO terms (g.eGO)
enriched GO terms (eGO) count
ratio g.eGO / g.GO
100kb bins, hg19 (total =28492)
AlwaysA
AorB:A
AorB:B
AlwaysB
High gene density (>3 genes)
Regular gene density (1-3 genes)
No gene
displays "active promoter" state
does not display "active promoter" state
Major subclasses (> 400 bins)
High gene density, B subclasses
subclass #

**Extended Data Fig. 1: Partitioning the human genome into chromatin subclasses.**

(A) Upper panel, GRO-seq and ChIP–seq signal for the histone marks indicated on the left across the same 25-Mb region of human chromosome 10 as in Fig. 1, in HUVEC cells (ENCODE tracks, visualized in IGV). Lower panel, ChromHMM segment counts or peak counts in 100-kb bins for the features indicated on the left.

(B) Chromatin subclasses assigned to each individual 100-kb bin across the same 25-Mb chromosomal region as in panel A. For additional details, see the legend to Fig. 1.

(C) GO analysis of bin subclasses. Histogram representation showing, for each subclass: total gene count (dark), genes associated with GO terms (g.GO, light gray), genes associated with at least one GO term enriched in the subclass (g.eGO, stippled yellow), and the count of GO terms enriched in the subclass (eGO, yellow; $P<0.05$). Left y-axis, counts (values indicated above bars). Blue curve, g.eGO/g.GO ratio (%) shown on the right y-axis, indicating the fraction of genes within a subclass that contribute to GO-term enrichment. A dotted blue line connects successive subclasses to emphasize trends. Low g.eGO/g.GO ratios in gene-rich subclasses indicate broad functional diversity (e.g. subclass 3, high gene density, AorB:A, active, containing over 2000 genes), whereas smaller subclasses with high g.eGO/g.GO ratios indicate strong functional specialization (e.g. subclasses 7 and 8, gene complexes displaying high gene density and embedded in AlwaysB regions). In these subclasses, a high fraction of genes is often associated with individual enriched GO terms (>10%, not shown).

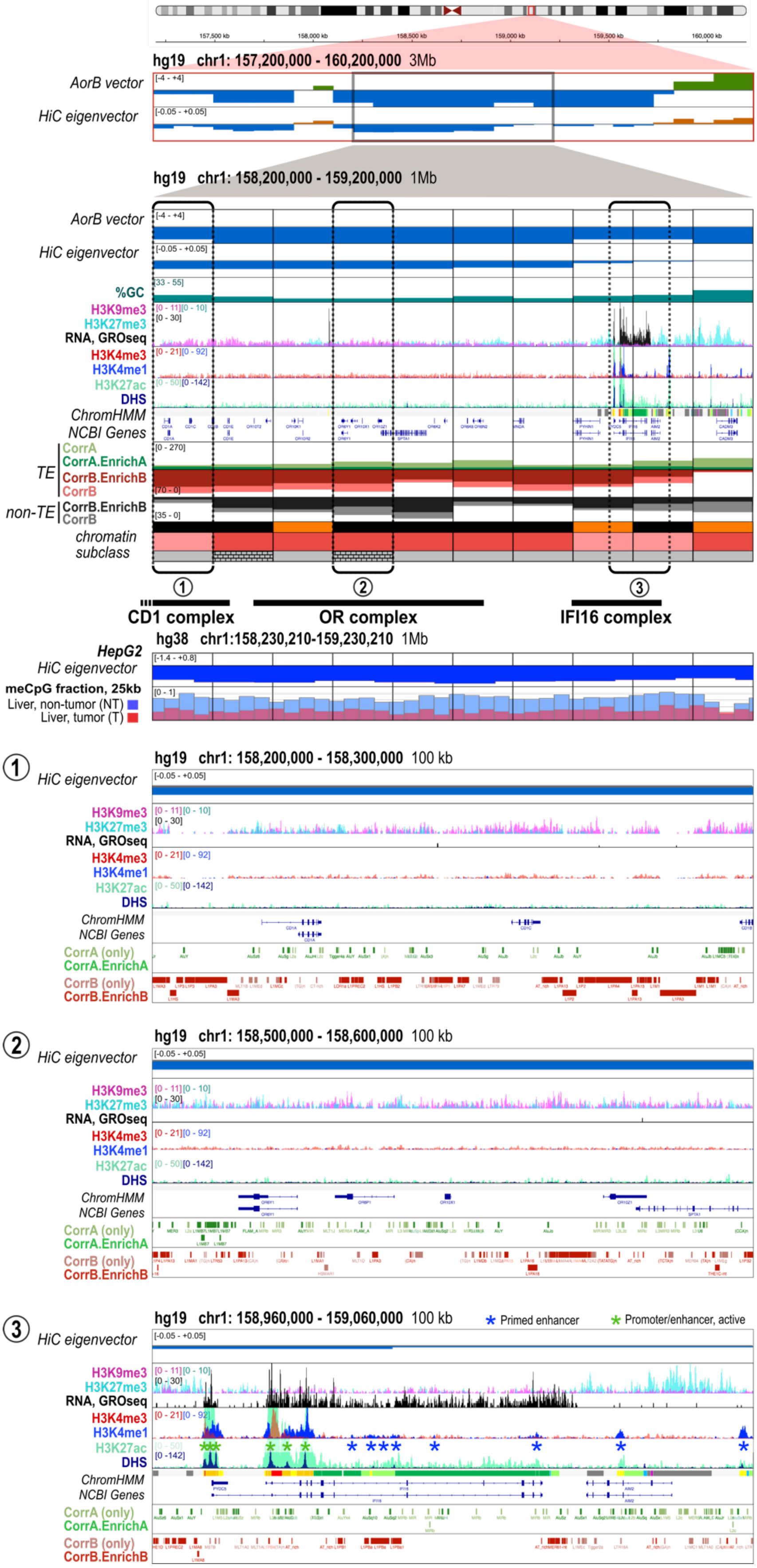
hg19 chr1: 157,200,000 - 160,200,000 3Mb
AorB vector
HiC eigenvector
hg19 chr1: 158,200,000 - 159,200,000 1Mb
AorB vector
HiC eigenvector
%GC
H3K9me3
H3K27me3
RNA, GROseq
H3K4me3
H3K4me1
H3K27ac
DHS
ChromHMM
NCBI Genes
TE
CorrA
CorrA.EnrichA
CorrB.EnrichB
CorrB
non-TE
CorrB.EnrichB
CorrB
chromatin subclass
CD1 complex
OR complex
IFI16 complex
hg38 chr1:158,230,210-159,230,210 1Mb
HepG2
HiC eigenvector
meCpG fraction, 25kb
Liver, non-tumor (NT)
Liver, tumor (T)
hg19 chr1: 158,200,000 - 158,300,000 100 kb
CorrA (only)
CorrA.EnrichA
CorrB (only)
CorrB.EnrichB
hg19 chr1: 158,500,000 - 158,600,000 100 kb
hg19 chr1: 158,960,000 - 159,060,000 100 kb
Primed enhancer
Promoter/enhancer, active

**Extended Data Fig. 2: chr1:158,200,000-159,200,000 (hg19), DNA and chromatin composition.**

The 1 Mb region shown is entirely part of the B compartment in HUVEC cells and exhibits a marked B trend, mostly scoring as AlwaysB (as indicated by strongly negative values of the AorB vector), which is also true for the surrounding regions (3 Mb panel at the top of the figure). However, the absolute values of the Hi-C EV are relatively low, and the last three bins on the right, which overlap with enlargement (3), exhibit Hi-C EV values close to zero. This is also the case for the five bins further to the right (3 Mb panel), collectively constituting an 800 kb region that toggles between A and B compartments.

The region has several notable features:
- It contains multiple gene complexes, corresponding to gene families that emerged through segmental duplication, forming a single locus. Gene complexes frequently score in the B compartment, but constitute a distinct type of B region, characterized by a relatively high gene density, whereas gene density is generally low or null in B regions.
- It includes two immune-related genes, CD1A and IFI16, whose repression during carcinogenesis is considered a widespread, early, and pivotal event [5].

The 1 Mb region shows a central AlwaysB domain, which corresponds to an Olfactory Receptor (OR) complex. This complex seemingly hosts two clusters of non-OR genes that themselves arose as a result of segmental duplication. From left to right, the region contains: five CD1 genes (CD1 molecules present lipid antigens to Natural Killer T cells) (see enlargement (1)), a central cluster of 13 olfactory receptor genes (see enlargement (2)), and the "IFI16 complex", consisting of four interferon-inducible genes encoding AIM2-like receptors, involved in innate immunity, among which: the AIM2 gene itself; the genes IFI16 and PYDC5, which are the only ones expressed in the 1 Mb region in HUVEC. They are massively transcribed, which is remarkable in a B context (see enlargement (3)). The IFI16 complex overlaps with the region displaying a very low value of the Hi-C EV. The IFI16 protein is ubiquitously expressed and plays a crucial role in immune surveillance of cancer emergence by binding both non-self and TE dsDNA, in particular L1 retrotransposition intermediates, and subsequently activating interferon signaling [6]. Note that the enlargements only contain a subset of the genes of the afore-mentioned gene families.
Binding of Polycomb-type heterochromatin across a chromosomal region is revealed by the H3K27me3 mark. The three “low Hi-C EV, B” bins on the right display a strong H3K27me3 signal outside of the two expressed genes. Such a pattern is commonly observed in regions that toggle between A and B compartments, particularly at boundaries between A and B domains, owing to the cooperation between H3K9me3/HP1α-based - and H3K27me3/Polycomb-based types of heterochromatin (see Extended Data Fig. 5). Notably, in the specific case of this locus, which is organized around a central OR complex, both H3K9me3 and H3K27me3 marks are pervasive across the entire region (see below).

Examining the DNA composition across the 1 Mb region reveals:
- A relatively low GC content (below 40%, except for the rightmost bin), which is commonly observed in AlwaysB regions, with an inverse bell shape such that GC content is lowest within the central OR complex and reaches 38-40% in the CD1 and IFI16 complexes.
- A RepSeq composition markedly unbalanced in favor of ProB RepSeq elements (except for the rightmost bin). ProB TEs are predominantly L1 elements, with densities among the highest in the human genome (enlargements (1-3)). ProB non-TE densities are also very high and largely correlate with GC content. There are a few ProA RepSeq elements, which are relatively well distributed along the locus.
Thus, both GC content and RepSeq composition are consistent with the fact that this region is embedded in a larger domain with a marked B-trend (see 3 Mb panel, upper part).

Both H3K9me3 and H3K27me3 heterochromatin marks are pervasive across the locus, though at modest levels, suggesting global repressive control through cooperation between Polycomb-based and HP1α-based types of heterochromatin, with ProB sequences playing a pivotal role, as independently supported [7]. The principles for cooperation between ProB elements over large chromosomal regions were established in [8] and are further illustrated in Fig. 8. There are 874 OR genes in the human genome (including approximately 50% pseudogenes), all embedded within similar L1-rich gene complexes in which every OR gene is repressed in all cell types except for a single allele in each olfactory sensory neuron of the olfactory mucosa [9]. OR complexes, together with the ProB sequences that enforce their repression, can therefore be viewed as providing a generally repressive chromatin environment that also supports the repressive control of non-OR embedded genes.
A well-known example is the β-globin locus, which resides within an OR complex on chromosome 11. Intriguingly, the α-globin locus lies in the subtelomeric region of chromosome 16p, which is likewise believed to confer a repressive chromatin environment [10,11]. Taken together, these observations suggest that the ProB power associated with OR complexes contributes not only to OR gene repression itself but also to higher-order genome organization.

The enlarged region (3), which contains the two active genes IFI16 and PYDC5, exhibits a remarkably dense array of open enhancers in primed or active states, distributed broadly across the locus (asterisks). DHS signals are faint or barely detectable at many of the primed enhancers (H3K4me1 peaks), possibly suggesting their transient existence. In contrast, the DHS, H3K27ac, and H3K4me3 signals are exceptionally strong at the promoter regions of both genes, collectively forming a ~20-kb super-enhancer.
Strikingly, the H3K27ac signal extends continuously across the entire IFI16 gene body, spanning more than 50 kb. These observations support a model in which enhancers and their associated TFs/cofactors counteract the surrounding B-compartment influence. In this locus, the unusually high density and activity of enhancers appear necessary to allow an A-type chromatin state to emerge despite the strong B-trend of the region. In addition, the high transcriptional output of IFI16 and PYDC5 provides a substantial ProA contribution, sustaining a feed-forward loop that maintains the region open in an otherwise unfavorable genomic environment.

The 1Mb locus appears to markedly lose CpG methylation in liver cancer, as expected for a locus with a marked B character (middle panel). Early and profound DNA hypomethylation of the whole locus is actually a consistent feature across multiple cancers [5]. Strikingly, such loss of CpG methylation is closely associated with repression of the CD1 and IFI16-related genes shown in enlargements (1) and (3), which was shown to be a key event of tumorigenesis for a variety of cancer types [5]. By contrast, OR genes are frequently found extraneously expressed in cancer [12].

These observations are consistent with a model in which, during early tumorigenesis, a subset of ProB RepSeqs - such as the abundant young L1 elements present at this locus - partially lose CpG methylation as they shift toward a more ProA-like configuration, with a twofold consequence: repression of OR genes is weakened, while ProB RepSeqs simultaneously gain the ability to recruit Polycomb heterochromatin, whose encroachment requires GC-rich, non-methylated DNA [13]. This recruitment remains highly dynamic and generally does not manifest as a detectable H3K27me3 ChIP peak (not shown). This process is further favored by the active delocalization of Polycomb heterochromatin away from the A compartment, which instead undergoes a chronic gain in CpG methylation during early tumorigenesis. Notably, even modest increases in CpG methylation at prominent targets such as the HOX loci (Extended Data Fig. 14) are predicted to trigger rapid disaggregation of Polycomb heterochromatin assemblies. The Polycomb components thus released readily accumulate at loci already targeted by Polycomb, in a “connecting-vessel” effect, as experimentally demonstrated [14].
Consistent with this framework, Polycomb invasion into B domains in cancer preferentially occurs at A/B compartment transition regions, which generally exhibit intermediate GC content, as exemplified by the IFI16 complex. The resulting increase in Polycomb heterochromatin occupancy appears sufficient to override the presence of multiple active enhancers, leading to IFI16 repression, as commonly observed across cancer types [5] and in HepG2 cells (not shown). A similar scenario is likely to apply to the CD1 locus in the relevant cell types [5]. By contrast, the OR locus largely resists Polycomb invasion, as observed in HepG2 cells (not shown), a property that may be explained by its low GC content.

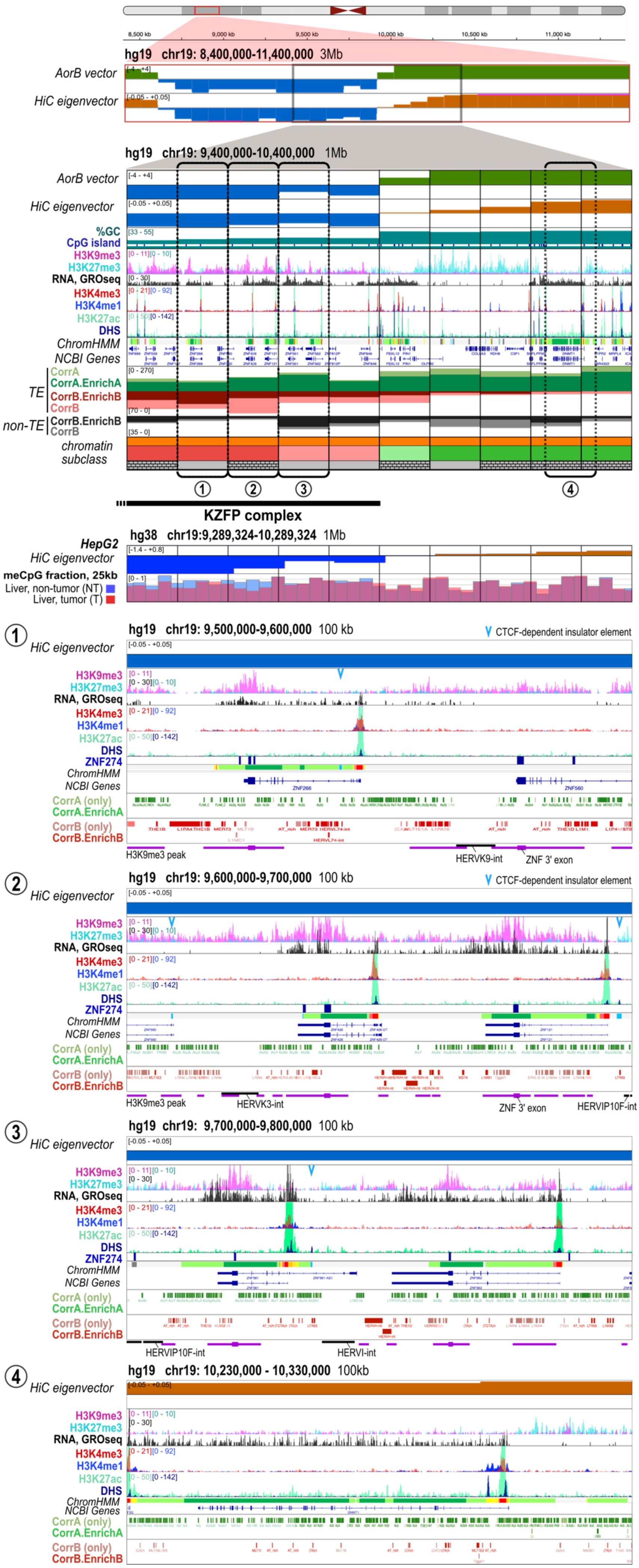

hg19 chr19: 8,400,000-11,400,000 3Mb
AorB vector
HiC eigenvector
hg19 chr19: 9,400,000-10,400,000 1Mb
AorB vector
HiC eigenvector
%GC
CpG island
H3K9me3
H3K27me3
RNA, GROseq
H3K4me3
H3K4me1
H3K27ac
DHS
ChromHMM
NCBI Genes
TE
CorrA
CorrA.EnrichA
CorrB.EnrichB
CorrB
non-TE
CorrB.EnrichB
CorrB
chromatin subclass
KZFP complex
HepG2
HiC eigenvector
meCpG fraction, 25kb
Liver, non-tumor (NT)
Liver, tumor (T)
hg38 chr19:9,289,324-10,289,324 1Mb
hg19 chr19: 9,500,000-9,600,000 100 kb
CTCF-dependent insulator element
ZNF274
CorrA (only)
CorrB (only)
H3K9me3 peak
HERVK9-int
ZNF 3' exon
hg19 chr19: 9,600,000-9,700,000 100 kb
HERVK3-int
HERVIP10F-int
hg19 chr19: 9,700,000-9,800,000 100 kb
HERVI-int
hg19 chr19: 10,230,000 - 10,330,000 100kb

**Extended Data Fig. 3: chr19:9,400,000-10,400,000 (hg19), DNA and chromatin composition.**

The left portion of the 1 Mb region shown overlaps with a KZFP complex (1 Mb panel). KZFPs constitute a superfamily of nearly 400 TFs in human and represents, for the most part, pivotal transcription-factor effectors of HP1α heterochromatin-mediated repression of RepSeqs [15]. The complex shown is relatively conserved over evolution, although it has been lost in the mouse lineage. While scoring as AlwaysB, with a pronounced B-character (indicated by the strongly negative values of the Hi-C EV), the 1 Mb region shown shares many characteristics with AlwaysA regions, similar to the one located on the right half of the 1 Mb region, namely: high to very high gene density, high transcriptional activity, gene promoters coinciding with CpG islands, near absence of active enhancers, and very high ProA RepSeq density. However, strikingly, these characteristics are associated with a significant presence of HP1α-based heterochromatin, as revealed by the prominent ChIP signal for H3K9me3, manifesting as discrete peaks, but also as a pervasive signal spreading across the domain. In contrast, H3K27me3, indicative of Polycomb-type heterochromatin, is nearly absent, appearing only as small, sparse patches that predominantly coincide with young ERV insertions (human endogenous retroviruses, HERVs; indicated by black lines in enlargements 1–3; see below). This portion of the KZFP locus shown is representative of the entire complex, which extends further to the left for a total of 1.2 Mb (3 Mb panel). In this context, a key function of heterochromatin is to prevent recombination between genes with highly similar sequences. Consistent with this function, the KZFP complex also contains the MUC16 gene, which contains internal repeats (not shown), as well as several olfactory-receptor genes, for which suppression of illegitimate recombination is likewise essential. ZNF genes are notably upregulated upon loss of the H3K9 methyltransferase SETDB1, demonstrating that heterochromatin also acts to restrain their transcription [16].
This region is also an outlier with respect to CpG-methylation loss in cancer: it shows only limited demethylation, in sharp contrast to standard AlwaysB regions. Nevertheless, a mild loss is detectable, especially when compared to the adjacent AlwaysA region on the right, which tends instead to display a slight gain in methylation. These observations suggest that, among the various A-like features of the KZFP complex, one or more confer an enhanced ability to resist meCpG erosion in cancer relative to canonical AlwaysB regions. A related conclusion is that prominent HP1α-heterochromatin occupancy is not, in itself, systematically associated with strong meCpG loss in cancerous B-compartment regions, an exception that again may reflect such compensatory effects.

Examining the RepSeq composition across the KZFP complex reveals the following:
- The RepSeq density in this region is exceptionally high, as nearly all sequences outside of coding or gene promoter regions correspond to RepSeq elements as identified with the RepBase library (not shown). The density of ProA elements is remarkably high, consisting of CorrA.enrichA elements rather than “CorrA only,” unlike the AlwaysA region further to the right or most regions genome-wide, which typically display both types. This may reflect the rapid evolution of KZFP complexes, which are associated with more recent insertions.
- The elevated presence of HP1α-based heterochromatin in the KZFP complex, can be explained by an overall high density of elements capable of nucleating heterochromatin (i.e. ProB elements), coinciding with H3K9me3 peaks (purple line beneath the 100-kb panels) (enlargements (1-3)). These include ProB TEs as well as an element located in the 3' exon of each ZNF gene. This exon encodes the zinc finger containing-DNA binding domain and shows the highest DNA sequence similarity between ZNF genes. Notably, the spreading of HP1α/H3K9me3 along the locus does not appear to be hindered by the high ProA density, suggesting that a balanced ratio of ProA and ProB permits a form of spreading over an extended region, according to principles described in [8] and illustrated in Fig. 8.
- Regarding the element located in the 3' exon of ZNF genes, the H3K9me3 peak at this element is present regardless of whether the gene is transcribed, but is more pronounced upon transcription (see enlargements (1-3)), with ZNF560 being the only untranscribed gene in this region. This aligns with the hypothesis that this well-described element, which depends on the ATRX remodeler, functions to prevent recombination between adjacent ZNF genes. The risk of recombination is higher when transcription occurs, requiring enhanced heterochromatin assembly [17-19]. Moreover, this element is bound by ZNF274 (see ZNF274 ChIP-seq peak track, enlargements (1-3)), which dynamically anchors the locus to the nucleolar surface, thereby facilitating the formation of a distinct heterochromatin-associated compartment that brings together multiple KZFP complexes as well as other gene complexes [17,19].
- In contrast to other gene complexes like OR loci, where the majority of ProB TEs consist of L1 elements, human KZFP complexes harbor both L1 and HERVs, particularly primate-specific HERVs, consistent with the fact that these are rapidly evolving loci [20]. A number of these are "HERV-int" inserts, where the LTRs have not recombined and thus contain remnants of internal coding sequences. Notably, some of these HERV-int inserts clearly play a ProB role within the KZFP complex, as they are decorated with H3K9me3 peaks (e.g., HERVK9-int, HERVK3-int, position indicated with a black line beneath enlargements 1 and 2), while these RepSeq subfamilies do not score as ProB because they do not meet the CorrB $< -0.01$ threshold

criteria. Others act as Polycomb-associated silencers, as indicated by a H3K27me3 signal (e.g., HERVIP10F-int, HERVI-int, enlargements (2) and (3)).
- During development and in cancer, many of these TEs, whether in ProB or H3K27me3-associated silencer configuration in HUVEC, can switch to a ProA/enhancer configuration due to the action of specific TFs, thereby activating nearby KZFP genes [21]. These TEs also form the basis for self-regulation loops of ZNF genes, and young TEs are often transcribed themselves in these contexts [20]. As a result, in cancer, and particularly in liver cancer, upregulation of KZFP genes is commonly observed (although this is not the case in HepG2 cells for the KZFP complex shown, nor for the ZNF genes or TEs; not shown) [21].
- Notably, although the overall %GC of the KZFP complex shown appears intermediate to low, it is in fact highly heterogeneous, with HERV inserts constituting the only sequences larger than 0.5 kb that display uniformly high GC content.

The distinctive genetic and epigenomic characteristics identified here for the KZFP locus are strikingly similar to those described for a 1.3 Mb region on chromosome 4 of Drosophila, with the entire chromosome 4 being well-known for its absence of recombination. Interestingly, within this 1.3 Mb region of the Drosophila genome, HP1a promotes high-level transcription of genes, associated with the combinatorial signature H3K9me3/H3K36me3, and the genes similarly display markedly open promoters. This region further contains interspersed TEs, which are classically repressed by HP1a-based heterochromatin [22,23]. It is therefore tempting to consider that, at the level of KZFP complexes, HP1α may have a similar dual role, acting differentially on both interspersed TEs and transcription units.

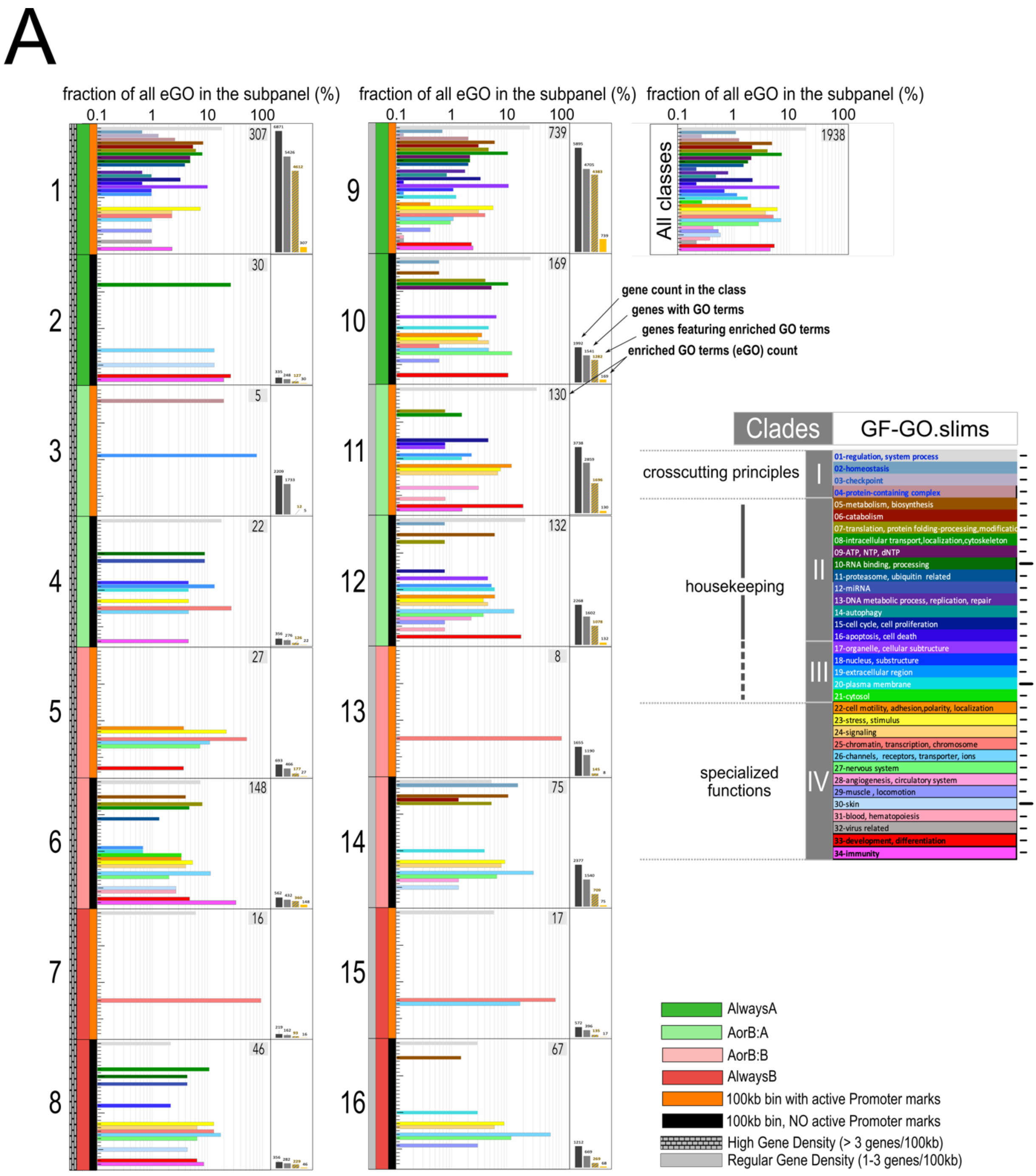
A
fraction of all eGO in the subpanel (%)
0.1
1
10
100
307
30
5
22
27
148
16
46
739
169
130
132
8
75
17
67
All classes
1938
gene count in the class
genes with GO terms
genes featuring enriched GO terms
enriched GO terms (eGO) count
Clades
GF-GO.slims
crosscutting principles
housekeeping
specialized functions
I
II
III
IV
01-regulation, system process
02-homeostasis
03-checkpoint
04-protein-containing complex
05-metabolism, biosynthesis
06-catabolism
07-translation, protein folding-processing,modificatio
08-intracellular transport,localization,cytoskeleton
09-ATP, NTP, dNTP
10-RNA binding, processing
11-proteasome, ubiquitin related
12-miRNA
13-DNA metabolic process, replication, repair
14-autophagy
15-cell cycle, cell proliferation
16-apoptosis, cell death
17-organelle, cellular subtructure
18-nucleus, substructure
19-extracellular region
20-plasma membrane
21-cytosol
22-cell motility, adhesion,polarity, localization
23-stress, stimulus
24-signaling
25-chromatin, transcription, chromosome
26-channels, receptors, transporter, ions
27-nervous system
28-angiogenesis, circulatory system
29-muscle , locomotion
30-skin
31-blood, hematopoiesis
32-virus related
33-development, differentiation
34-immunity
AlwaysA
AorB:A
AorB:B
AlwaysB
100kb bin with active Promoter marks
100kb bin, NO active Promoter marks
High Gene Density (> 3 genes/100kb)
Regular Gene Density (1-3 genes/100kb)

B

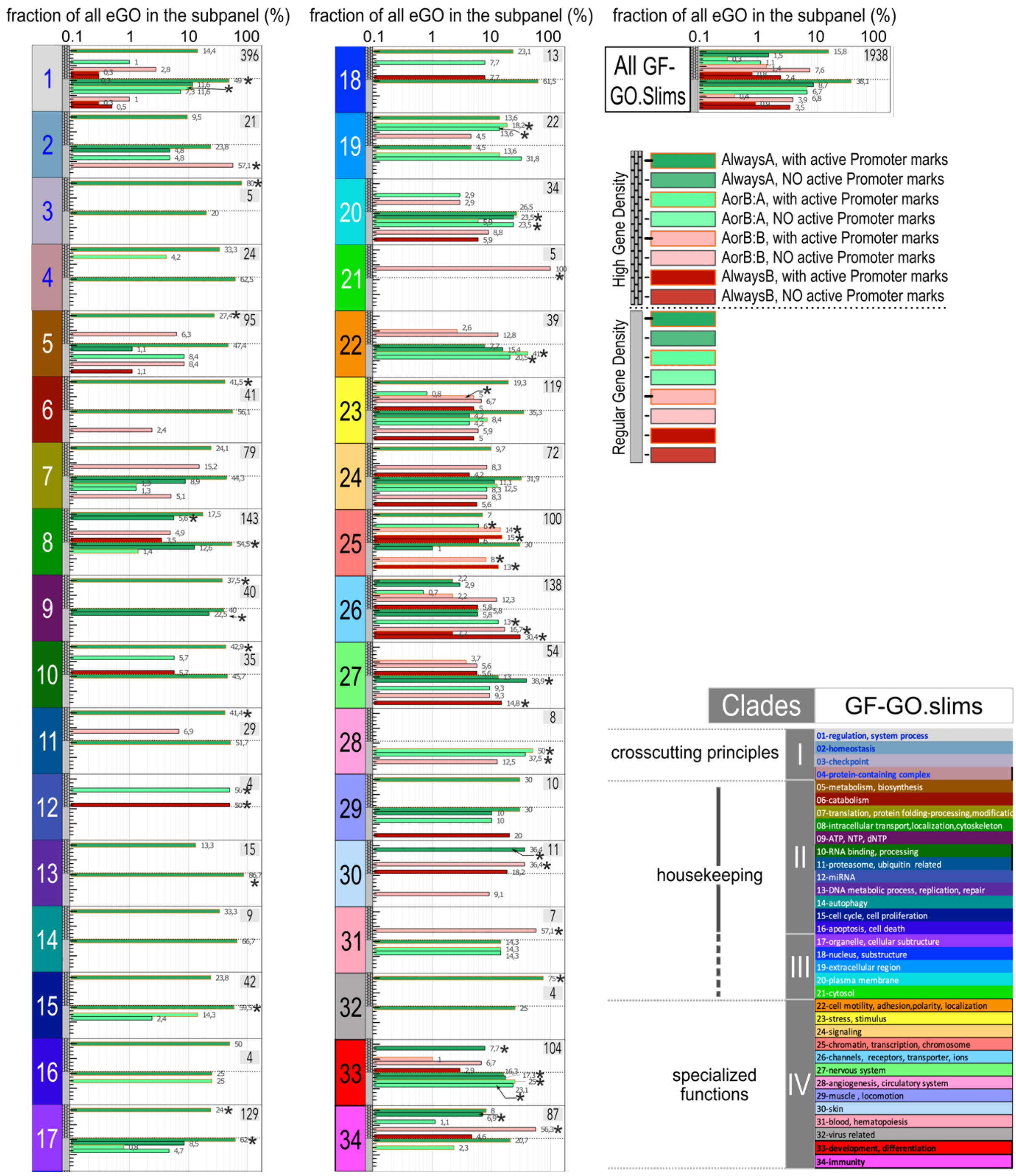
fraction of all eGO in the subpanel (%)
0.1
1
10
100
All GF-GO.Slims
1938
High Gene Density
Regular Gene Density
AlwaysA, with active Promoter marks
AlwaysA, NO active Promoter marks
AorB:A, with active Promoter marks
AorB:A, NO active Promoter marks
AorB:B, with active Promoter marks
AorB:B, NO active Promoter marks
AlwaysB, with active Promoter marks
AlwaysB, NO active Promoter marks
Clades
GF-GO.slims
crosscutting principles
housekeeping
specialized functions
I
II
III
IV
01-regulation, system process
02-homeostasis
03-checkpoint
04-protein-containing complex
05-metabolism, biosynthesis
06-catabolism
07-translation, protein folding-processing,modificatio
08-intracellular transport,localization,cytoskeleton
09-ATP, NTP, dNTP
10-RNA binding, processing
11-proteasome, ubiquitin related
12-miRNA
13-DNA metabolic process, replication, repair
14-autophagy
15-cell cycle, cell proliferation
16-apoptosis, cell death
17-organelle, cellular subtructure
18-nucleus, substructure
19-extracellular region
20-plasma membrane
21-cytosol
22-cell motility, adhesion,polarity, localization
23-stress, stimulus
24-signaling
25-chromatin, transcription, chromosome
26-channels, receptors, transporter, ions
27-nervous system
28-angiogenesis, circulatory system
29-muscle , locomotion
30-skin
31-blood, hematopoiesis
32-virus related
33-development, differentiation
34-immunity

**Extended Data Fig. 4: GO analysis reveals enrichment of AlwaysA regions in housekeeping genes, enrichment of AorB regions in genes with evolutionarily specialized function, and the presence of gene complexes in AlwaysB environments.**

(A) Subclass-centered GO analysis, shown in 16 subpanels. For each bin subclass, information found in panel C is indicated again on the right, with the total number of eGO terms (found significantly enriched in that class) indicated in the upper right corner. The distribution of these GO terms among the 34 GF-GO.slim terms is displayed as a percentage, with the color of the histogram bar indicating the GF-GO.slim term according to the legend. The upper right subpanel labelled "All subclasses" displays the compiled distribution of all eGOs (GO terms found enriched in this analysis in any one of the bin subclasses). Notably, more than 50% of the population of eGOs are contributed by the AlwaysA, active bin subclasses 1 and 9. Note the logarithmic scale. An example of how to read this panel is as follows: Bin subclass 3 (high gene density, AorB:A, active) features only 5 GO terms enriched, one of which (20%) is associated with GF-GO.slim 04 (protein-containing complex), and 4 (80%) are associated with GF-GO.slim 19 (extracellular region).

(B) Whether and which epigenomic landscape correlates with a particular type of function may be better appreciated by displaying the same information as shown in panel A in a GF-GO.slim-centered manner. In other words, for each GF-GO.slim, the total number of GO terms recovered as enriched in our GO analysis and falling in this GF-GO.slim is indicated in the upper right corner, and histogram bars represent the contribution in eGOs for each individual bin subclass, expressed as a percentage. The upper right subpanel labelled "All GF-GO.slim" displays the compiled distribution of all eGOs as a function of the bin subclasses where they are found enriched. A star (FDR < 0.05) indicates that enrichment of a GF-GO.slim in a particular bin subclass is significantly stronger than the value observed for the compilation of all eGOs. Let's consider for instance GF-GO.slim 22 ("cell motility, adhesion, polarity, localization"). This GF-GO.slim is associated with GO terms found enriched in the bin subclasses 5 and 6 (high gene density, AorB:B) and 9-12 (regular gene density, AlwaysA and AorB:A). However, only for bin subclasses 11 and 12 (regular gene density, AorB:A) is the enrichment remarkable, meaning that in the other four bin subclasses this GF-GO.slim is only weakly enriched relative to other GF-GO.slims.

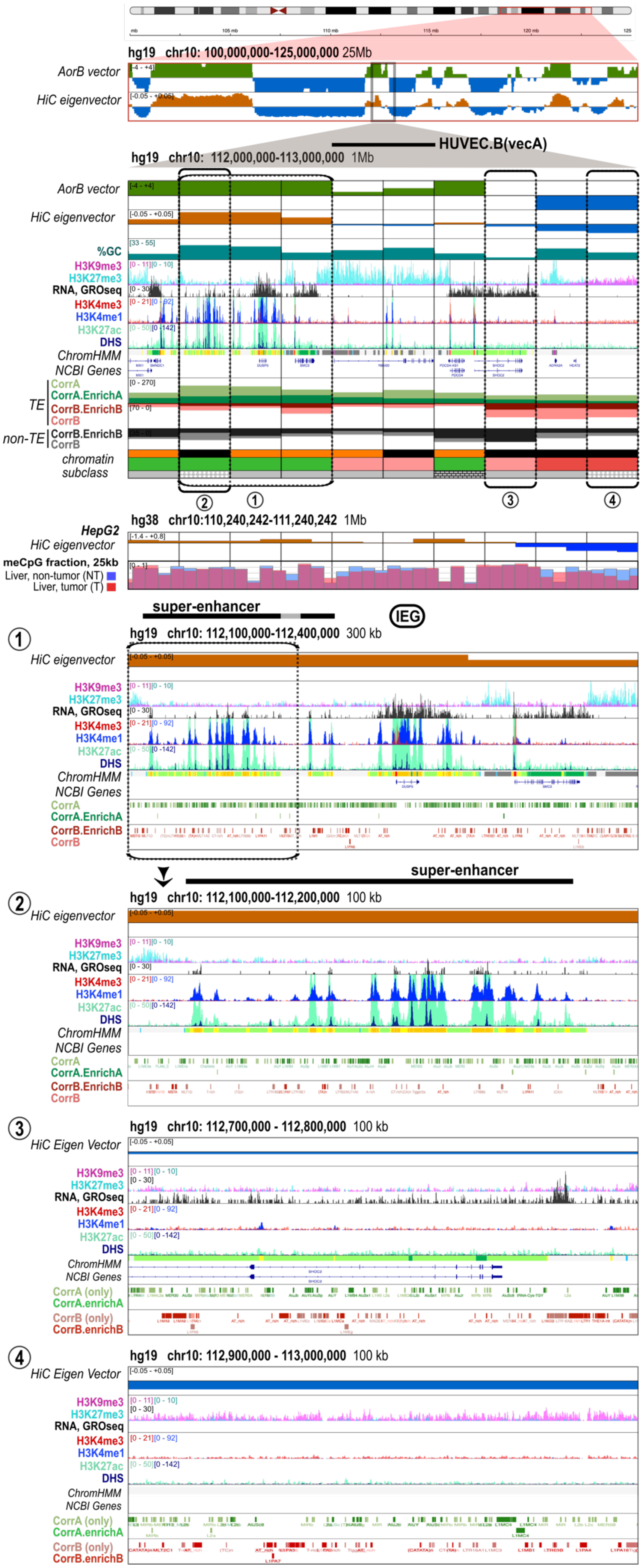
hg19 chr10: 100,000,000-125,000,000 25Mb
AorB vector
HiC eigenvector
HUVEC.B(vecA)
hg19 chr10: 112,000,000-113,000,000 1Mb
%GC
H3K9me3
H3K27me3
RNA, GROseq
H3K4me3
H3K4me1
H3K27ac
DHS
ChromHMM
NCBI Genes
TE
CorrA
CorrA.EnrichA
CorrB.EnrichB
CorrB
non-TE
chromatin subclass
hg38 chr10:110,240,242-111,240,242 1Mb
HepG2
meCpG fraction, 25kb
Liver, non-tumor (NT)
Liver, tumor (T)
super-enhancer
IEG
hg19 chr10: 112,100,000-112,400,000 300 kb
hg19 chr10: 112,100,000-112,200,000 100 kb
hg19 chr10: 112,700,000 - 112,800,000 100 kb
HiC Eigen Vector
CorrA (only)
CorrA.enrichA
CorrB (only)
CorrB.enrichB
hg19 chr10: 112,900,000 - 113,000,000 100 kb

**Extended Data Fig. 5: chr10:112,000,000-113,000,000 (hg19), DNA and chromatin composition.**

The 1 Mb region shown is located within the representative 25 Mb region on chromosome 10 also shown in Figures 1-7 and repeated here at the top. This 1 Mb region represents a transition zone between A and B compartments, transitioning from the AlwaysA class on the left to the AlwaysB class on the right. Central bins display Hi-C EV absolute values close to zero and belong to the AorB class.

The region exhibits a relatively standard and homogeneous gene density, except for the bin shown in enlargement (2) (see below), which displays paradoxical characteristics at first glance, as it scores in AlwaysA but contains no genes, belonging to chromatin subclass 18. This bin harbors a cluster of enhancers and in fact represents a typical super-enhancer driving expression of the neighboring gene DUSP5, which is an immediate early gene (IEG) [24] (see below).

The RepSeq composition is remarkably progressive and parallels the Hi-C EV profile, transitioning from high-density ProA/low-density ProB on the left to low-density ProA/high-density ProB on the right. This trend holds true with the exception of the bin shown in enlargement (3) (see below). Additionally, the super-enhancer shown in enlargement (2) exhibits a relatively high density of ProA elements. However, these are predominantly TEs from evolutionarily ancient subfamilies that are not significantly enriched in the A compartment (CorrA, pale green), in contrast to other super-enhancers (e.g., SOX2, HOXA, Extended Data Figs. 7 and 14), which typically exhibit a phenomenal density of CorrA.enrichA elements, specifically Alu sequences.

CpG methylation is relatively homogeneous across the region, with a few "valleys" displaying low meCpG levels, corresponding to CpG islands (not shown). CpG methylation levels remain remarkably unchanged in liver cancer compared to normal liver, consistent with the intermediate position of this region between sequences at the core of an AlwaysA domain on the left (which gain some methylation) and AlwaysB sequences on the right (which lose methylation more clearly), as well as with genome-wide averages shown in Fig. 7E.

The H3K27me3 mark is spread across most of the region, sparing only the transcribed areas, which aligns with the well-established observation that Polycomb heterochromatin typically installs by default in non-transcribed regions of the A compartment. However, the H3K27me3 signal is particularly strong in the central portion (112.35-112.65), coating the RBM20 gene which is not expressed in HUVEC. RBM20 overlaps with a HUVEC.B(vecA) segment, being part of the B compartment in HUVEC and part of the A compartment in the majority of the seven other cell lines of our reference panel. Such a strong H3K27me3 signal is typical of A-B transition regions, resulting in characteristic shoulder-like signals flanking each A domain in the H3K27me3 ChIP data (see Extended Data Fig. 1). It is also commonly observed over HUVEC.B(vecA) segments (Fig. 5C), which are frequently found in these transition regions. Considering processes in 3D within the nucleus of a living cell, the omnipresence of the H3K27me3 mark at A-B transition regions suggests that the entire region resides within a nucleoplasmic environment where the binding of structural components of Polycomb heterochromatin to the nucleosomal fiber is favored, owing to the activity of partner enzymes but also enzymes from the neighboring HP1α-based heterochromatin, particularly HDACs (see Extended Data Fig. 18).

The regulation of DUSP5 by a super-enhancer is characteristic not only of IEGs but also of genes crucial for cell identity, particularly those encoding pioneer TFs. The TFs bound to the super-enhancer and recruiting coactivators act both to directly activate DUSP5 transcription and to oppose the establishment of Polycomb heterochromatin, through the formation of phases and HAT activity that counteracts the effects of HDACs (see Extended Data Fig. 18) [25,26]. This principle of anti-repression enables fine-tuning of transcription, even for genes transcribed at high levels, and is predicted to be particularly efficient for DUSP5 due to its location in an A-B transition region.

The bin shown in enlargement (3) exhibits a relatively high density of ProB elements, both TE and non-TE, coinciding with a low GC content (panel 1Mb). This bin is an outlier in that it scores as AorB, yet its composition more closely resembles that of a bin located at the core of an AlwaysB domain (see enlargement (4) for instance). However, this bin, along with the adjacent bin to the left, contains three ubiquitously and strongly expressed genes: SHOC2, PDCD4, and PDCD4-AS1. These can be considered strong ProA elements, as the transcription process itself is associated with ProA forces [27].
Interpreting these data in light of the molecular mechanisms by which ProA and ProB elements act (Extended Data Fig. 18) and earlier work [8] suggests that the bin shown in enlargement (3) acts as a relay, extending the "ProB influence" of the adjacent AlwaysB domain on the right into the A domain on the left through its ProB elements. It constitutes a boundary in the sense that it is the place where the two chromatin logics, A and B, confront each other. Notably, the absence of the H3K9me3 mark over this bin (compare with

the bin shown in enlargement (4)) can be attributed to the destabilization of HP1α-based heterochromatin installation, due to the strong ProA effect associated with SHOC2 gene transcription. The bins shown in enlargements (3) and (4) also illustrate how two bins with similar DNA compositions, which are located close to each other along the genome, can exhibit very different epigenomic characteristics because one lies at the boundary of a B domain, while the other is at the core of the domain.

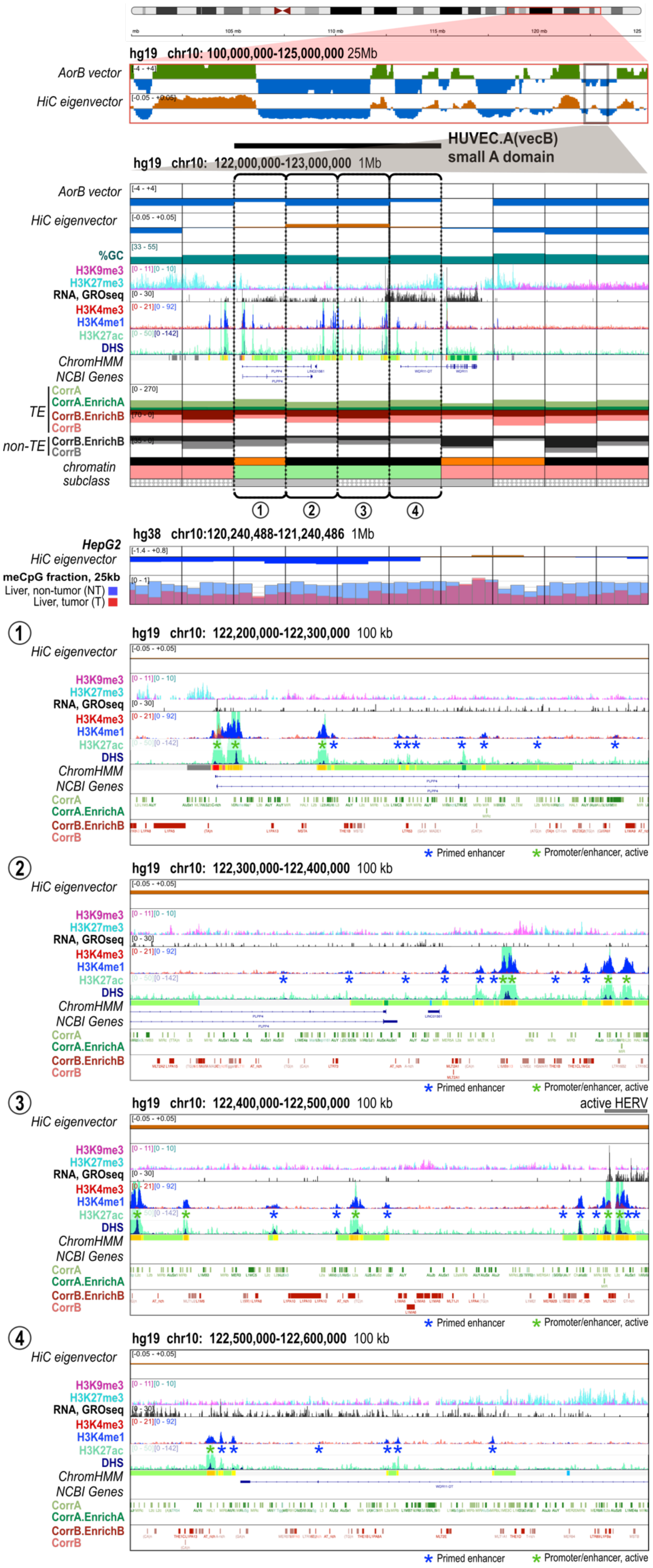

hg19 chr10: 100,000,000-125,000,000 25Mb
AorB vector
HiC eigenvector
HUVEC.A(vecB)
small A domain
hg19 chr10: 122,000,000-123,000,000 1Mb
%GC
H3K9me3
H3K27me3
RNA, GROseq
H3K4me3
H3K4me1
H3K27ac
DHS
ChromHMM
NCBI Genes
TE
CorrA
CorrA.EnrichA
CorrB.EnrichB
CorrB
non-TE
chromatin subclass
① ② ③ ④
hg38 chr10:120,240,488-121,240,486 1Mb
HepG2
meCpG fraction, 25kb
Liver, non-tumor (NT)
Liver, tumor (T)
hg19 chr10: 122,200,000-122,300,000 100 kb
hg19 chr10: 122,300,000-122,400,000 100 kb
hg19 chr10: 122,400,000-122,500,000 100 kb
active HERV
hg19 chr10: 122,500,000-122,600,000 100 kb
* Primed enhancer
* Promoter/enhancer, active

**Extended Data Fig. 6: chr10:122,000,000-123,000,000 (hg19), DNA and chromatin composition.**

The 1 Mb region shown is located within the representative 25 Mb region on chromosome 10, also depicted in Figures 1-7 and repeated here at the top. The central 400 kb portion (shown in enlargements (1-4)) is remarkable in that it constitutes a small A domain in HUVEC cells, whereas it is part of the B compartment in the majority of the other cell lines in our panel (HUVEC.A(vecB) domain). The 1 Mb region contains only three genes (PLPP4, WDR11, and WDR11-OT), clustered within or in the immediate vicinity of the HUVEC.A(vecB) domain, all of which are transcribed in HUVEC cells.
The entire 1 Mb region exhibits a chimeric DNA sequence composition, combining A-type GC content with a ProA/ProB RepSeq ratio that favors ProB. This is consistent with the observed B-trend within this extended AorB region, as indicated by negative values of the AorB vector. The density of ProB non-TE elements clearly decreases over the HUVEC.A(vecB) domain, reaching a level more commonly found in the A compartment. Notably, a majority of HUVEC.A(vecB) domains instead display an A-type ProB TE density and a GC content that corresponds to a B-type level (Fig. 5C).

Both CpG methylation levels in normal liver and CpG methylation loss in liver cancer are relatively homogeneous across the entire 1 Mb region (Hi-C EV in HepG2 cells), consistent with the relatively uniform underlying DNA composition (see above). meCpG loss in liver cancer is pronounced, as expected for an AorB region with a B trend (see Fig. 7). This pattern holds across the region except around the WDR11 gene, which scores as A and is actively transcribed in HepG2 cells, whereas PLPP4 is not (not shown). This subregion maintains a high CpG methylation level and remains transcriptionally active within a larger domain that undergoes marked methylation loss and becomes embedded in Polycomb heterochromatin (in HepG2 cells, not shown). This organization is generally consistent with the notion that, in cancer, Polycomb heterochromatin is redistributed away from the A compartment toward AorB regions with typically intermediate DNA composition and that lose CpG methylation (see Extended Data Fig. 2). The unusually low GC content across most of the WDR11 gene (not shown) likely contributes to its resistance to Polycomb heterochromatin invasion. This type of pattern is referred to as a Preserved Methylation Island and is frequently observed for cancer-associated genes [5]. WDR11 is indeed a polymorphic factor that participates in the Hedgehog signaling pathway, which is commonly implicated in liver tumors.

The HUVEC.A(vecB) domain exhibits a remarkably high density of DHS, some appearing as primed enhancers and others as active enhancers or promoters (indicated by asterisks in enlargements (1-4)), with an average of 12 such enhancers per 100 kb for the bins shown in enlargements (1-3), corresponding to approximately one every 8 kb. Such a high density of active DHS/enhancers is a characteristic feature of HUVEC.A(vecB) domains (see Fig. 5C) as well as small A domains (see Fig. 6B). This can generally be interpreted as the requirement for strong ProA activity to counteract the B-influence in the region and thereby open an A domain. The relatively even distribution of DHS along the HUVEC.A(vecB) domain further supports this view. Altogether, the low absolute Hi-C EV value of the HUVEC.A(vecB) domain suggests that the domain toggles between A and B compartments and may be part of the A compartment when enough of the multiple DHS/enhancers cooperatively repel the B-influence arising from both around and within - first, by acting as primed enhancers, and second, possibly by initiating the assembly of a coactivator phase as active enhancers.
Finally, this pattern of numerous enhancers distributed across the region is strikingly reminiscent of super-enhancers (see Extended Data Fig. 2), suggesting that a similar logic of anti-repression applies in both cases. The difference in the intensity of ChIP signals for active chromatin marks very likely reflects only differences in the local chromatin environment.

In conclusion, the entire 1 Mb region shown, which exhibits low absolute Hi-C EV values and thus toggles between A and B compartments over an extended length, constitutes a distinctive chromatin habitat with an intermediate character. This coincides with a mixed DNA composition, with an intermediate RepSeq combination displaying a B-trend and GC content showing an A-trend. This composition is relatively neutral, but the fact that the larger environment is generally B-trend suggests that it is a configuration conducive to dynamic regulation via anti-repression, where domain opening requires strong ProA activity, corresponding to the DHS/enhancers found distributed along the central small A domain. This configuration is quite similar to that observed at A-B junctions (see Extended Data Figs. 2 and 5).

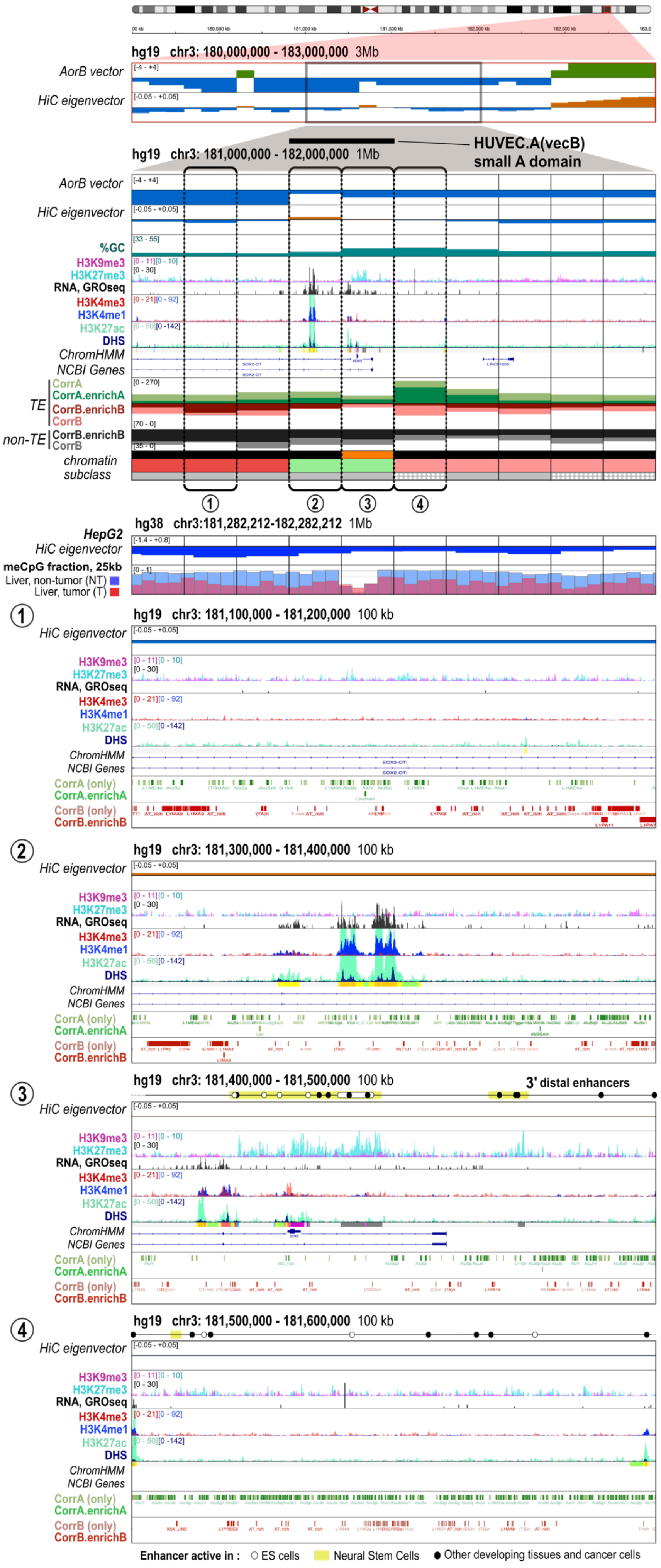
hg19 chr3: 180,000,000 - 183,000,000 3Mb
AorB vector
HiC eigenvector
hg19 chr3: 181,000,000 - 182,000,000 1Mb
HUVEC.A(vecB) small A domain
%GC
H3K9me3
H3K27me3
RNA, GROseq
H3K4me3
H3K4me1
H3K27ac
DHS
ChromHMM
NCBI Genes
TE
CorrA
CorrA.enrichA
CorrB.enrichB
CorrB
non-TE
chromatin subclass
hg38 chr3:181,282,212-182,282,212 1Mb
HepG2
meCpG fraction, 25kb
Liver, non-tumor (NT)
Liver, tumor (T)
hg19 chr3: 181,100,000 - 181,200,000 100 kb
CorrA (only)
CorrB (only)
hg19 chr3: 181,300,000 - 181,400,000 100 kb
hg19 chr3: 181,400,000 - 181,500,000 100 kb
3' distal enhancers
hg19 chr3: 181,500,000 - 181,600,000 100 kb
Enhancer active in : ES cells Neural Stem Cells Other developing tissues and cancer cells

**Extended Data Fig. 7: chr3:181,000,000-182,000,000 (hg19), DNA and chromatin composition.**

The SOX2 gene is located in the middle of a 2 Mb region, which is part of the B compartment in the majority of the eight cell lines in our reference panel. As visible on the 3 Mb panel, the SOX2 gene features an AlwaysB block on the left (5' of the gene) and an AorB:B block on the right (3' of the gene) clearly representing a B-trend environment. However, the entire 1 Mb region shown displays Hi-C EV values close to 0 in HUVECs, indicating that this extended region actually toggles between A and B compartments.

We selected four 100 kb bins to display in enlargements. Bin (3) contains the SOX2 gene. Bins (2) and (3) show positive values of the Hi-C EV in HUVEC, but negative values of the AorB vector, thus constituting a small HUVEC.A(vecB) region. Bins (3) and (4) overlap with the major regulatory region of SOX2, spanning approximately 200 kb and containing multiple enhancers responsible for gene expression in various cell types, as indicated, although none of these enhancers is active in HUVEC cells.

The SOX2 gene is very small (4.5 kb), a size comparable to that of IEGs. It is silent in HUVEC and harbors a poised promoter, with a combination of H3K4me3/H3K27me3 marks. In addition, there are two long non-coding RNA (lncRNA) genes in the 1 Mb region. SOX2- OT is a very long non-coding "overlapping transcript" for the SOX2 gene, transcribed in the same direction, visible on the left half of the 1 Mb panel and overlapping with bins (1-3). It may be involved in assisting the induction of SOX2 [28]. The other lncRNA, LINC01206, overlapping with bin (4), is expressed only in the testis and its function is unknown.

Examining the DNA composition across the 1 Mb region reveals:
- A remarkably low GC content over the left and right regions, around 36-38%, which is associated with an enrichment in AT-rich ProB non-TE elements. These are the only DNA features that align with B-type characteristics. Indeed, both ProA and ProB TEs densities are relatively low. Notably, the few ProB TE elements found in this 1 Mb region are all L1 elements, which are highly AT-rich across most of their length (enlargements (1) and (2); not shown).
- A significant rise in ProA density over the central region (chr3:181.3-181.7 Mb; with the exception of Bin (3)), primarily composed of Alu sequences, peaking at a very high level in bin (4). The general ProA profile in this portion aligns with the GC profile, reaching 41-42%, a typical level for the A compartment. Bin (4) ranks among the human genome regions richest in Alu sequences.
- A near absence of both ProA and ProB TEs in bin (3), similar to what can be observed at the HOXA locus, another locus known to be strongly regulated by Polycomb heterochromatin (see Extended Data Fig. 14). Interestingly, the region located 3' of Sox2 in mice harbors a super-enhancer in mouse embryonic stem (mES) cells and similarly exhibits a very high density of SINE B1 and B2 elements [29,30]. SINE B1 and B2 are the mouse equivalents of Alu, but they evolved independently. Thus, the high density of SINEs at both the mouse and human Sox2 loci represents a case of convergent evolution [31] providing strong support for the notion of a SINE-specific mechanism performing ProA function (see below).

Only bin (2) has a clearly positive Hi-C EV value in HUVECs, and it appears to draw the neighboring bins into the A compartment. This bin contains a distinct super-enhancer approximately 30 kb in length, that is massively transcribed in HUVECs.

Bin (3) exhibits notably low levels of CpG methylation and appears as a DNA methylation valley (DMV) in normal liver. This aligns with the strong and extended H3K27me3 ChIP signal over the SOX2 gene. Indeed, Polycomb heterochromatin preferentially associates with regions of high GC content and unmethylated CpG dinucleotides, accounting for the mutual exclusion between meCpG and H3K27me3 signals as observed [32]. Across the 1 Mb region, meCpG levels drop markedly in cancer, as expected for a B-trend region (see Fig. 7). However, in Bin (3), which contains the SOX2 gene, CpG methylation remains unchanged, or even slightly increases, in cancer. As a result, the meCpG level in Bin (3) becomes nearly indistinguishable from that of the surrounding 1 Mb region.
This pattern is consistent with partial delocalization of Polycomb heterochromatin from SOX2 during tumorigenesis (see Extended Data Fig.2 and main text). As a result, heterochromatin-mediated repression of SOX2 is predicted to weaken in liver cancer. Nonetheless, SOX2 is not expressed in HepG2 cells (not shown).

Overall, the DNA composition of the locus suggests the following regulatory model for the SOX2 gene.

The SOX2 gene is subject to strong global repressive control due to its location in a chromosomal environment with marked B-character, including a large 600-kb AlwaysB region on its 5′ side, and an overall high AT content, which favors the establishment of H3K9me3/HP1α heterochromatin. In cell types in which the locus is open, such as HUVECs, SOX2 remains under strong repressive control via Polycomb

heterochromatin, with its promoter maintained in a poised state, awaiting an instructive trigger. This ensures extremely low background expression, which is essential given that SOX2 is an oncogene [33].
Notably, SOX2 environment is characterized by the absence of a strongly defined A- or B-compartment identity. This supports efficient Polycomb heterochromatin function, and its installation is further facilitated by the relatively high GC content in the immediate SOX2 neighborhood (adjacent bins), in contrast to the rest of the region.

Although no H3K27me3 ChIP signal is detected over the 3′ region of SOX2, particularly in Bin (4), our preferred model is that the entire locus resides within a nuclear subvolume enriched in Polycomb heterochromatin components. Polycomb installation is facilitated in the immediate vicinity of SOX2 due to the absence of TEs, allowing effective spreading and stabilizing the chromatin state through self-reinforcement loops. In contrast, in regions with very high Alu density, especially Bin (4), Polycomb components are envisioned to bind in a highly dynamic manner, as spreading is inhibited by Alu repeats. This labile repression permits the rapid emergence of active enhancers upon a trigger and associated TF induction, facilitating the swift activation of SOX2 transcription, as observed in diverse regenerative contexts [33].

In carcinogenesis, a local increase in CpG methylation across the SOX2 gene gives the impression of partial methylation homogenization within a region that otherwise exhibits a global decrease consistent with its B trend. This creates a sensitized situation in which, counterintuitively, an increase in CpG methylation results in the SOX2 gene itself being less efficiently repressed, due to inhibition of Polycomb heterochromatin encroachment (see Extended Data Fig. 2 and main text). As a consequence, SOX2 may be induced by a minimal amount of inducer signal and associated TFs,, particularly Sox2 protein itself [33], generating a feed-forward loop that can lead to high SOX2 expression, as observed in some cancers.
Our explanations further clarify why strong long-range interactions between the SOX2 gene and its enhancers can readily be detected, and why contacts between Sox2 and the super-enhancer in murine ES cells are visible even by microscopy [29,30,34]. Indeed, the surrounding B-trend regions anchor nuclear subvolumes enriched in HP1α-heterochromatin components, which create a form of spatial exclusion that allows an active chromatin phase to form more easily and to coalesce more visibly.

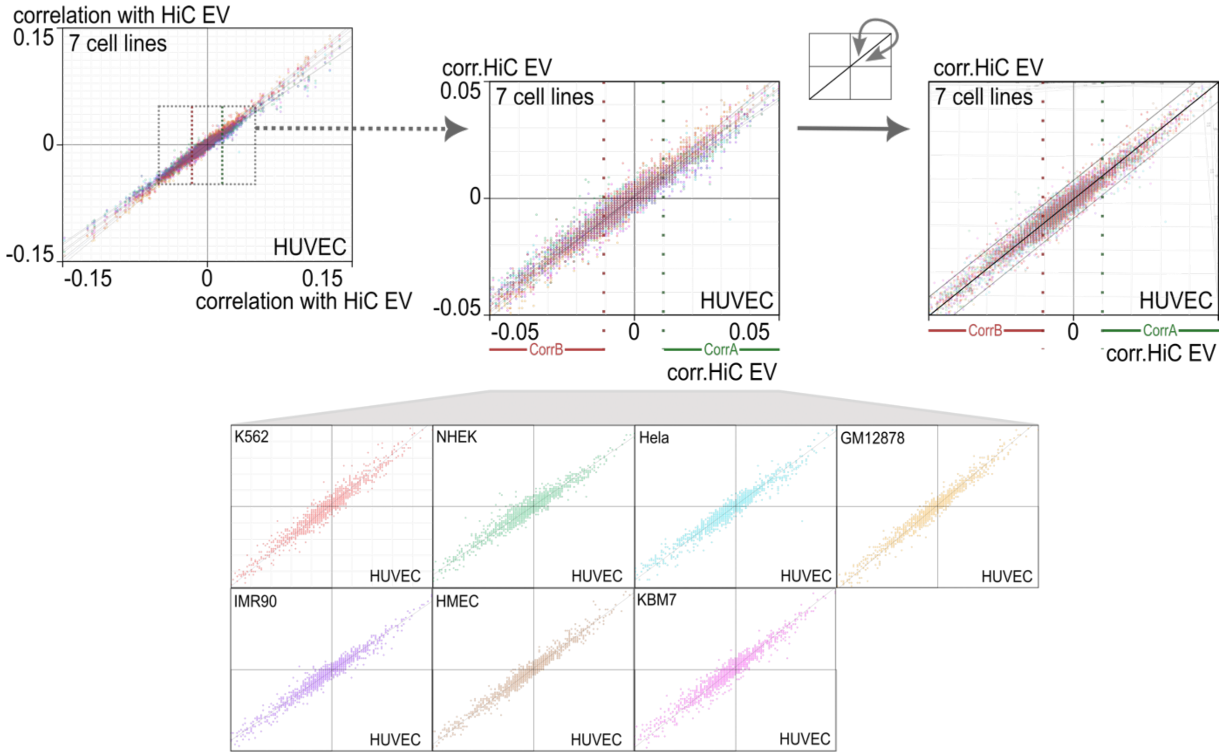

**Extended Data Fig. 8: Spearman correlation of RepSeq subfamilies with Hi-C EV is generally similar across cell lines.**

Each RepSeq subfamily is represented as a point in Euclidean space according to its Spearman correlation with the Hi-C EV in HUVEC cells and in the seven indicated cell lines, each shown in a different color. Center panel: magnified view of the –0.05 to +0.05 correlation range. Right panel: values obtained in the seven cell lines are slightly rotated relative to one another to render regression lines superimposable.

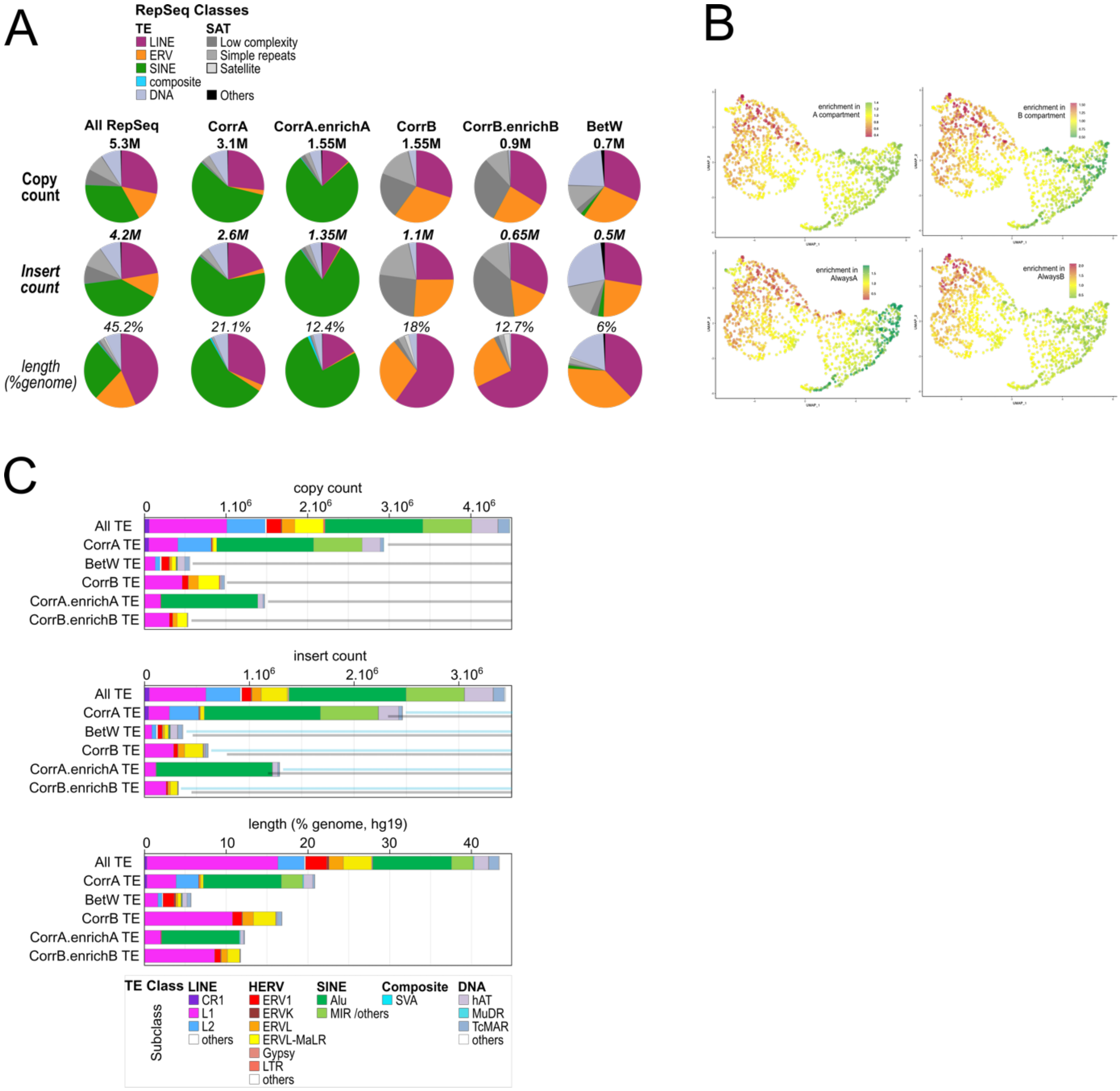

**Extended Data Fig. 9: Identification of ProA and ProB RepSeqs in the human genome: composition of RepSeq sets, UMAP analysis.**

(A) Pie charts showing composition of the full complement of RepSeq elements in the human genome (hg19), and of the RepSeq sets defined in this study, according to RepSeq class. Copy count corresponds to the total number of segments of continuous homology with a subfamily consensus sequence, as defined in the reference NCBI RepBase reference file. TE inserts frequently consist of multiple such segments, and are quantified as insert count (see Materials and Methods). For additional details, see legend to Fig. 3A.

(B) UMAP analysis of all RepSeq subfamilies from RepBase, based on the following parameters: enrichment (in A, in B, in AlwaysA, in AlwaysB); correlation (with Hi-C EV; with H3K9me3 ChIP-seq signal ; with H3K27me3 ChIP-seq signal); 100-kb binning; HUVEC data; age and copy-count categories. Coloring indicates enrichment. The panel "enrichment in B compartment" is also shown in Fig.3B and is reproduced here for reference.

(C) TE abundance in RepSeq sets, displayed by subfamily, shown as absolute counts (copy count or insert count, as defined in panel A) and as genomic coverage (% of total genome). Blue and black semi-transparent lines are included to facilitate visual comparison between copy count and insert count panels. For additional details, see panel A.

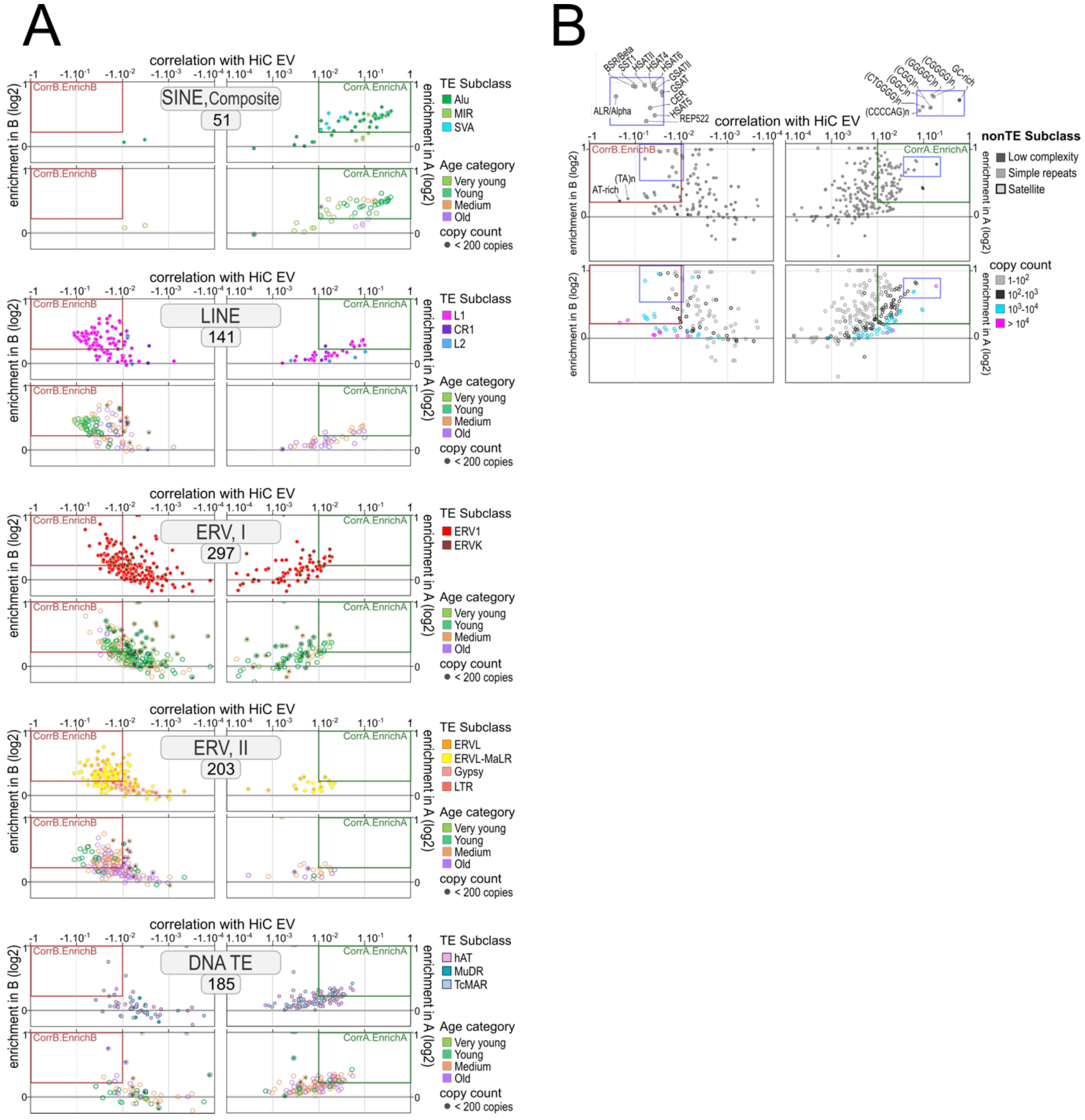

**Extended Data Fig. 10: Identification of ProA and ProB RepSeqs in the human genome: representation according to families.**

Euclidean representation of RepSeq subfamilies according to their correlation with Hi-C EV and enrichment in A (right) or B (left) compartments (as in Fig.3D, except that TE (A) and non-TE (B) subclasses are displayed in separate panels).

(A) Boxed number, total count of subfamilies represented in the panel. Lower panels, the colors indicate the median age category of each subfamily. Central black fill, subfamily copy count below 200. Note that each ERV subfamily is represented by two points, corresponding to LTR and internal sequences, respectively. For clarity, 91 TE subfamilies scoring predominantly in the Medium and Old age categories are omitted.

(B) Lower panels, the colors indicate the genomic copy count of each subfamily. Blue boxes, magnified above the Euclidean plane, highlight non-TE elements most strongly enriched in the B compartment (centromeric and pericentromeric satellites, and the telomere-associated composite element REP522) and in the A compartment (GC-rich low-complexity and simple-repeat satellites). Additional details are provided in the legend to Fig. 3D.

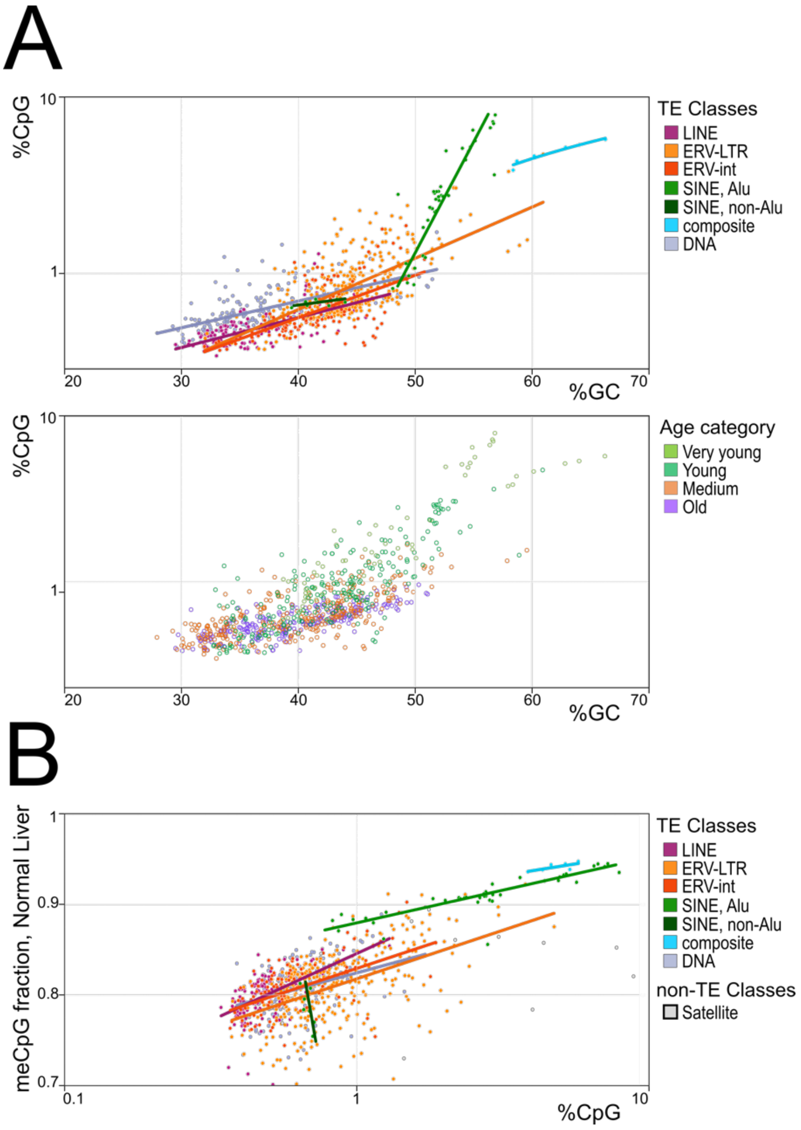

**Extended Data Fig. 11: GC content and DNA methylation of human RepSeqs.**

(A) Euclidean representation of TE subfamilies based on mean GC content (% nucleotides, linear scale) and CpG content (% dinucleotides, log scale). Lower panel, colors indicate the median age category of each subfamily. Each ERV subfamily is represented by two points corresponding to LTR (ERV-LTR) and internal (ERV-int) sequences. Regression lines are shown for individual TE classes.

(B) Euclidean representation of RepSeq subfamilies based on mean CpG content and meCpG fraction in normal liver. See Fig. 3D legend for additional details.

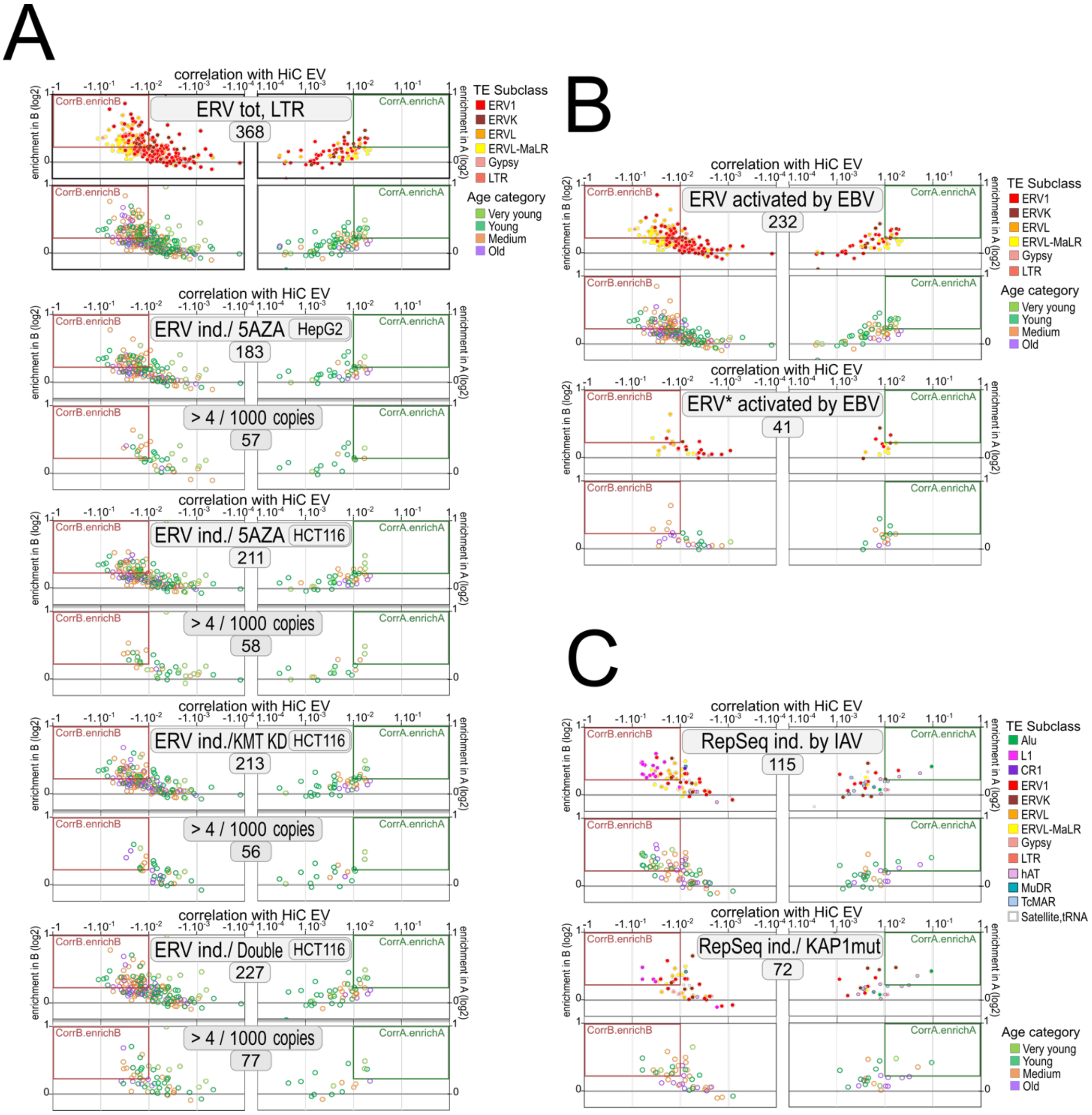

**Extended Data Fig. 12: Subsets of RepSeqs switch from a ProB to a ProA state upon chromatin perturbation and viral infection.**

(A-C) As in Extended Data Fig. 3A, except that only panels colored by subfamily age are shown in selected cases, for RepSeq sets functionally defined as follows.

(A) ERV subfamilies transcriptionally induced (>2-fold, RNA-seq) upon chromatin perturbation as described in [35]: inhibition of DNA methyltransferases with 5-azacytidine (5AZA), knockdown of major H3K9 methyltransferases (KMT KD; SUV39H1, SETDB1, G9a), or combined treatment (5AZA + KMT KD, "double"). Only RNA-seq reads mapping to LTRs of intergenic ERV inserts were considered to exclude intronic TE signal. For the combined treatment, only ERVs induced >2-fold relative to the single 5AZA treatment are shown. Data are shown for HepG2 (5AZA) and HCT116 (all treatments), as indicated. For each condition, upper panels show all subfamilies with at least one induced insert, whereas lower panels show subfamilies with >4 induced inserts per 1,000, indicative of subfamily-level induction.

(B) ERVs activated by EBV. ERV subfamilies with at least one pseudo-unique insert showing a marked gain of H3K4me3 ChIP-seq signal over the LTR in EBV-immortalized human B cells (LCLs, lymphoblastoid cell lines), compared with primary B cells from the same donor [36] (1,134 ERV inserts total). These LTRs generally show reduced CpG methylation, and 31% overlap a CAGE signal in the ENCODE LCL reference cell line GM12878, consistent with bona fide promoter activity. ERV* activated by EBV, same analysis restricted to ERV subfamilies reaching significance at the subfamily level ($P<0.05$).

(C) RepSeq subfamilies transcriptionally induced in the cancer-derived A549 cell line upon influenza A virus (IAV) infection (upper panels) or expression of a non-SUMOylatable KAP1 mutant as the sole KAP1 species (KAP1 mut, lower panels; RNA-seq fold induction >1.17; P<0.05 at the subfamily level) [37]. IAV infection is associated with loss of KAP1 sumoylation as part of the cellular response to infection, and the strong overlap between IAV- and KAP1 mut-induced RepSeq induction patterns supports this mechanism. Median age corresponds to the median age of RepSeq subfamilies, not of the individual induced copies.

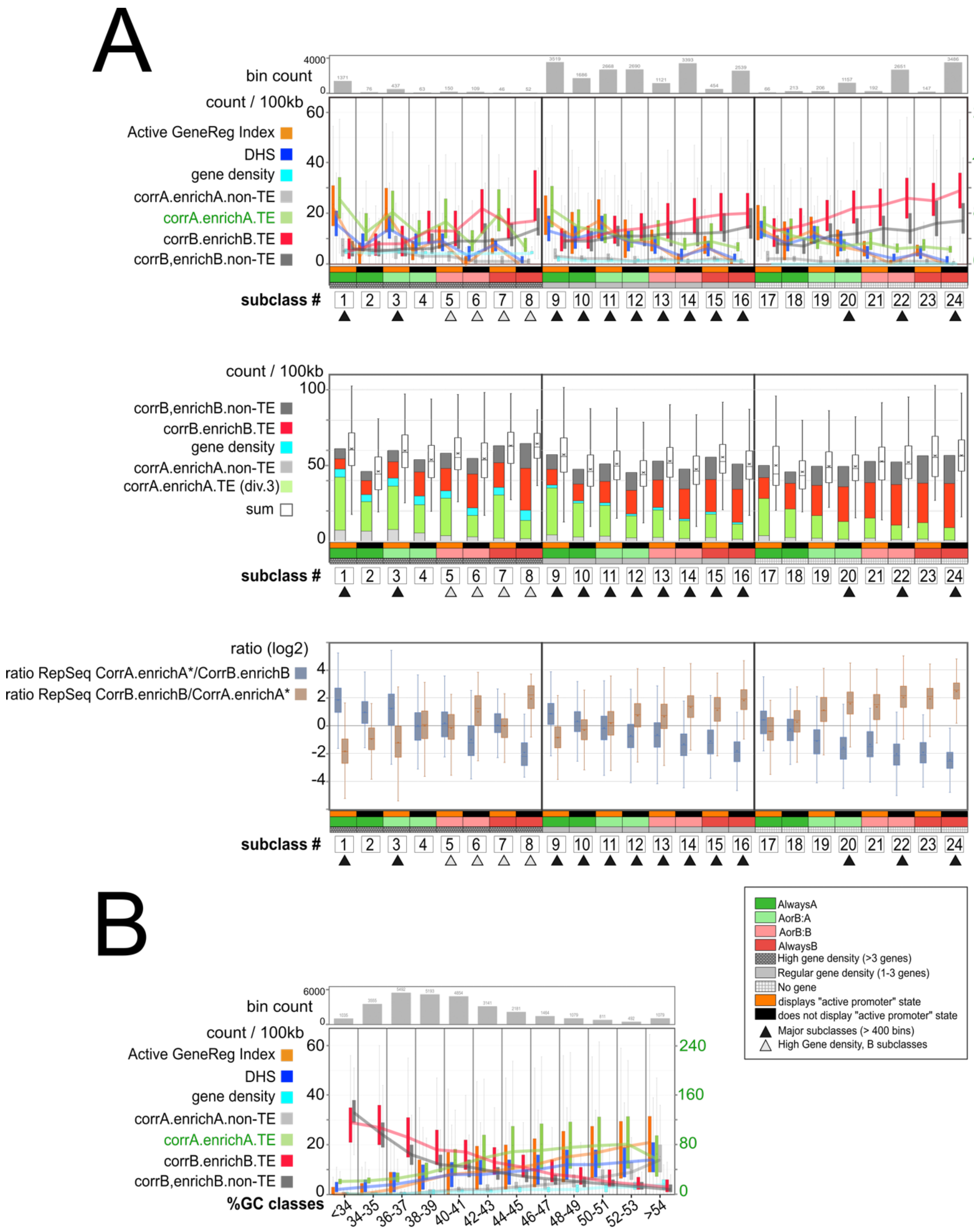

C

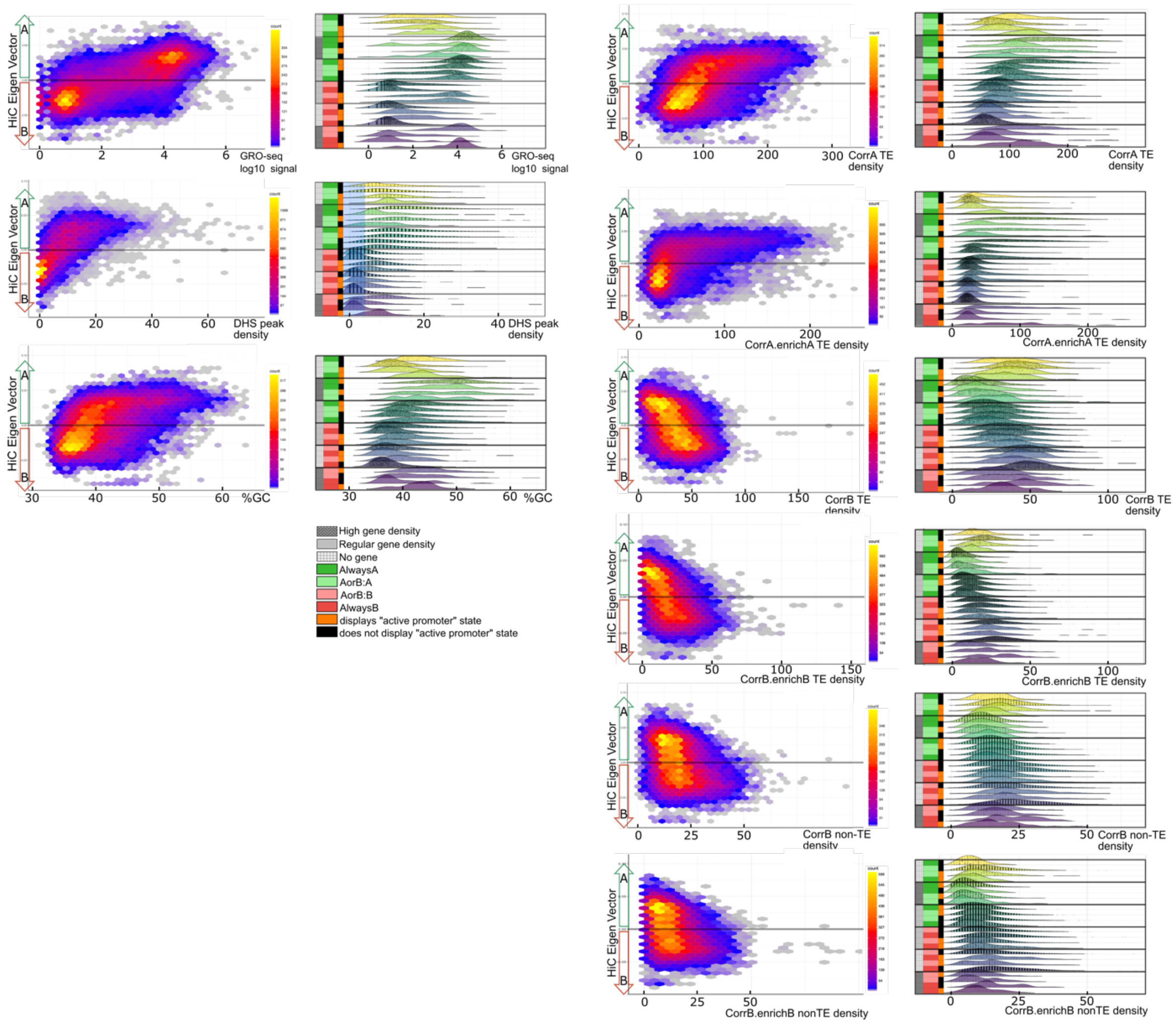

**Extended Data Fig. 13: Compositional genomics reveals general trends for the local densities of ProA and ProB elements.**

(A) Top: Boxplot representation of the density of DNA and chromatin features in all 24 bin subclasses. The Active GeneReg Index is a sum calculated from ChromHMM segment counts as follows: 0.5*WeakPromoter + 1.5*StrongPromoter + 0.5*WeakEnhancer + 1.5*StrongEnhancer. Lines connect successive median values. The bin count in each subclass is indicated. Note that the scale is different for CorrA.enrichA TE density, as indicated in green on the right of the graph. For other details, see also the legend to Fig. 1 and Fig.4.
Middle: The colored stacked bar representation shows the mean value for the features indicated, in each bin subclass. Corr.enrichA TE* denotes that the Corr.enrichA TE density value is divided by 3. Boxplot representation (white) of the sum of the features indicated on the left of the graph, for each individual bin in each bin subclass.
Bottom: Boxplot representation of the ratio between CorrA.enrichA* and CorrB.enrichB RepSeq density, as well as the inverse ratio, for each individual bin in each bin subclass. CorrA.enrichA* is the sum of CorrA.enrichA non-TE counts, and CorrA.enrichA TE counts divided by 3. CorrB.enrichB is the sum of the counts of CorrB.enrichB TE and non-TE.

(B) Same as panel A except that the genome is partitioned into 12 bin subclasses according to %GC.

(C) Scatter plot (left) and ridge plot representation (right) for DNA and chromatin features as indicated, in individual bins.
Scatter plots: Each bin is shown as a point. The y-axis indicates the Hi-C EV value in a bin, and the x-axis indicates the value (GRO-seq summed signal, %GC) or density (others) of the feature in the same bin.
Ridge plot representation for the 24 bin subclasses, with points corresponding to individual bins shown in black. For other details, see also the legend to Fig. 4.

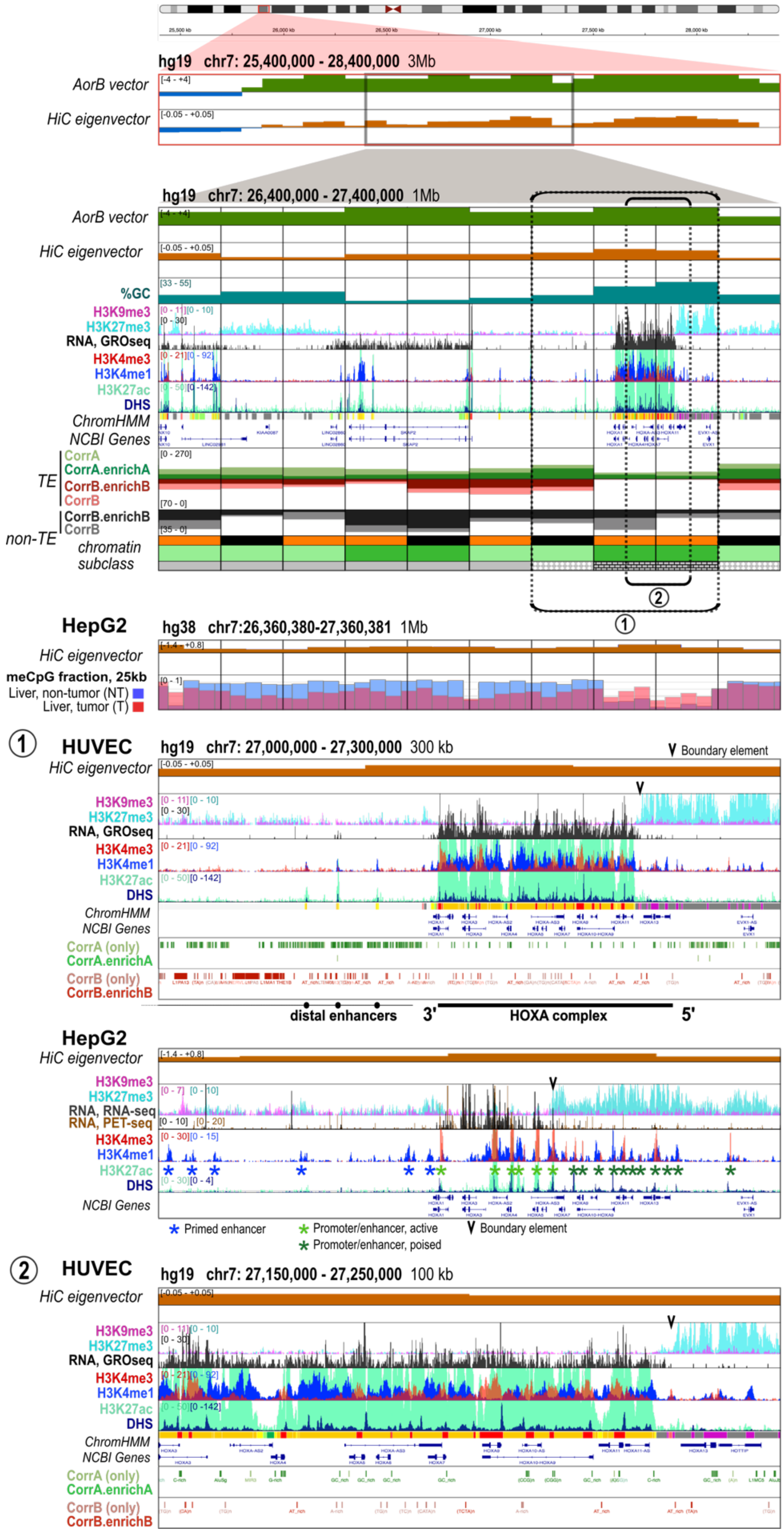
hg19 chr7: 25,400,000 - 28,400,000 3Mb
AorB vector
HiC eigenvector
hg19 chr7: 26,400,000 - 27,400,000 1Mb
AorB vector
HiC eigenvector
%GC
H3K9me3
H3K27me3
RNA, GROseq
H3K4me3
H3K4me1
H3K27ac
DHS
ChromHMM
NCBI Genes
TE
CorrA
CorrA.enrichA
CorrB.enrichB
CorrB
non-TE
CorrB.enrichB
CorrB
chromatin subclass
HepG2
hg38 chr7:26,360,380-27,360,381 1Mb
HiC eigenvector
meCpG fraction, 25kb
Liver, non-tumor (NT)
Liver, tumor (T)
HUVEC
hg19 chr7: 27,000,000 - 27,300,000 300 kb
Boundary element
HiC eigenvector
CorrA (only)
CorrA.enrichA
CorrB (only)
CorrB.enrichB
distal enhancers
3'
HOXA complex
5'
HepG2
HiC eigenvector
RNA, RNA-seq
RNA, PET-seq
Primed enhancer
Promoter/enhancer, active
Promoter/enhancer, poised
Boundary element
HUVEC
hg19 chr7: 27,150,000 - 27,250,000 100 kb
HiC eigenvector

**Extended Data Fig. 14: chr7:26,400,000-27,400,000 (hg19), DNA and chromatin composition.**

HOXA is one of the four HOX gene complexes in humans. All HOX genes are transcribed in the same orientation within a locus (hence the terminology "5'-to-3' " for their genomic order), but there are also both sense and antisense long non-coding RNAs located between or overlapping with these genes, which play regulatory roles.

The 1 Mb region containing the HOXA locus is entirely part of the A compartment in most of the eight ENCODE reference cell lines in our panel (and thus essentially scores as AlwaysA). It is located within an extended region that shows a clear A-trend (see the 3Mb enlargement). However, it shares many characteristics with AorB regions, notably a relatively low absolute Hi-C EV value in both HUVEC and HepG2 cells (see 1Mb enlargements, HUVEC and HepG2), as well as a balanced composition of ProA and ProB RepSeqs (further details below). The GC content is highly heterogeneous, peaking in the 200kb region encompassing the HOXA locus, which also contains a large number of CpG islands, both inside and outside promoters (not shown), and displays very low CpG methylation in normal liver, appearing as a large DMV (HepG2/Liver 1Mb panel). This is consistent with the fact that HOX loci are prominently regulated by Polycomb heterochromatin, whose binding to the chromatin fiber requires high GC content and the absence of CpG methylation.

In HepG2 cells, an H3K27me3 signal is observed across a portion of the HOXA complex that is not transcribed, on the 5′ side of HOXA6. In HUVEC cells, the H3K27me3 signal is only observed at HOXA13, at the very 5′ end of the locus (see enlargement (1)), which is the only inactive HOXA gene in these cells and encodes a repressor for the other HOXA genes. The H3K27me3 signal further extends to the neighboring 5′ gene EVX1, where it is particularly strong, paralleling the fact that this ~70 kb HOXA13–EVX1 region exhibits very high GC content and is virtually unmethylated at CpG sites in normal liver tissue. Individual HOX genes are believed to recruit and promote Polycomb encroachment at the nucleosomal fiber both autonomously and cooperatively, owing to their high GC content and the presence of multiple CpG islands. However, it is possible that the HOXA13–EVX1 region exhibits a greater degree of autonomy and capacity in this regard, functioning as a master silencer that imposes a dominant, Polycomb-associated repressive control over the entire locus. This mechanism could help explain, at least in part, why the locus tends to open in continuous portions starting from the 3′ end, with the 5′ portion being more resistant to opening, both during development and in differentiated cells [38], as previously proposed [8].

HOX gene complexes are highly compact, and, unsurprisingly, a conserved feature throughout evolution is that each transcriptional unit, including a HOX gene and its nearby enhancers, is flanked by insulator elements. These insulators provide a degree of autonomy in the regulation of the gene by acting as dynamic boundaries both for the action of enhancers within the domain and for the spreading of Polycomb heterochromatin from outside the domain. Insulator elements in HOX loci are largely dependent on the CTCF factor, both in vertebrates and invertebrates [39-41]. Two insulator elements are indicated with arrowheads in enlargements, serving as boundaries against the spreading of Polycomb heterochromatin from the EVX1/HOXA13 region in HUVEC and HepG2 cells. CTCF-dependent insulators are similarly found at other gene complexes, such as ZNF loci (see Extended Data Fig. 3).

***RepSeq composition and its impact on the epigenomics landscape:***

While HOX loci have been extensively used as models for gene regulation studies, they are actually significant outliers compared to the rest of the genome in many respects. These loci are characterized by a notably high gene density, and in this sense, are similar to other gene complexes. More intriguingly, however, they exhibit a near-complete absence of ProA and ProB TEs, as seen here for the HOXA locus (enlargements (1) and (2)). A low density of ProB non-TE elements is observed, however, along with ProA non-TE elements which correspond to GC-rich simple repeats (see Extended Data Fig. 10) within HOX gene promoters, which overlap with CpG islands (not shown). The absence of transposable elements within HOX loci suggests that evolution has actively selected against their presence, implying that any TE insertion would disrupt the precise regulatory mechanisms governing the spatiotemporal expression of genes within these loci, and very likely also their repression by Polycomb heterochromatin.

Interestingly, a substantial density of ProA and ProB RepSeqs is found outside the HOXA locus itself, specifically in the 3' and 5' distal regions (1Mb panel and enlargement (1)), similar to what is observed within the locus for other gene complexes. Thus, two blocks of ProA elements flank the HOXA locus, as also observed at the SOX2 locus (Extended Data Fig. 7). Additionally, a region of approximately 250 kb 3' of the HOXA locus, between the HOXA1 and SKAP2 genes and partially overlapping with SKAP2, exhibits a mixed RepSeq composition (enlargement (1)). The density of ProA and ProB is moderate to high, including both

ProB TE (L1s, as well as a few ERVs) and ProB non-TE elements. Notably, this is quite similar to the ProA and ProB RepSeq composition within the ZNF gene complex (Extended Data Fig. 3). Strikingly, the SKAP2 gene itself shows a very low GC content.

This composition and distribution of RepSeqs suggest the following functional scenario: The 3' and 5' distal regions of the locus are known to contain pivotal enhancers for the regulation of the HOXA locus. These regions are partially protected from Polycomb spreading by the presence of insulators within the HOXA locus. Furthermore, the high local density of Alu elements is consistent with the notion that these repeats prevent efficient Polycomb heterochromatin encroachment in these regions, and thereby subject them to a more dynamic mode of repression, as also proposed for the extended SOX2 super-enhancer (Extended Data Fig. 7). Moreover, the fact that the 3' region additionally contains a significant density of both ProB TE and non-TE elements suggests that it likely serves as a nucleation center for HP1α-type heterochromatin, operating as a relay for this type of heterochromatin along the chromosome arm [8], facilitating repression of the HOXA locus by Polycomb heterochromatin due to the cooperation between the two types of heterochromatin, as is also clearly visible at the interfaces between A and B compartments (Extended Data Fig. 5).

***Alteration of meCpG patterns in cancer:***

A notable loss of CpG methylation is observed in liver cancer across most of the 1 Mb region (middle panel), consistent with the AorB character of the region (see Extended Data Fig. 15A). Remarkably, however, meCpG levels are very low over the ~200 kb segment containing the HOXA locus in normal liver tissue but increase in cancer, reaching levels nearly equivalent to the rest of the 1 Mb region. Such an increase in CpG methylation at HOX loci is a general phenomenon observed in a variety of cancer types [42], as is also the case in other Polycomb-repressed regions of the genome (see SOX2 locus, Extended Data Fig. 7). This seemingly modest increase in meCpG density is predicted to heavily alter the nucleation of Polycomb heterochromatin, leading to a near-collapse of Polycomb assembly through a “house-of-cards” effect, thereby impairing the transcriptional repression of underlying sequences [13]. This observation aligns with the well-established finding that HOX genes are frequently derepressed and can act as key driver oncogenes in cancer, as observed for HOXA9 in AML and HOXB13 in prostate cancer [42].

Fine changes in the H3K27me3 pattern over the HOXA13–EVX1 region in HepG2 cells, as well as the presence of clear DHS and marked H3K4me3 peaks at poised promoters in the extended H3K27me3-marked HOXA7–HOXA13 region, further support the idea of progressive erosion of Polycomb-mediated repression at the HOXA locus in liver cancer (enlargement (1)). Conversely, a basal H3K27me3 signal is observed in the region located 3′ of the HOXA locus in HepG2 cells, suggesting that the slight decrease in CpG methylation in this region may, on the contrary, facilitate Polycomb heterochromatin binding, although other chromatin alterations could also contribute.

These observations align with the notion that, in cancer, Polycomb heterochromatin tends to bind to regions where it previously did not, driven in part by CpG hypomethylation (see Fig. 7) [32] (see also Extended Data Fig. 2). This pervasive binding reduces the pool of factors available for specific targeting, thereby globally weakening the Polycomb heterochromatin.

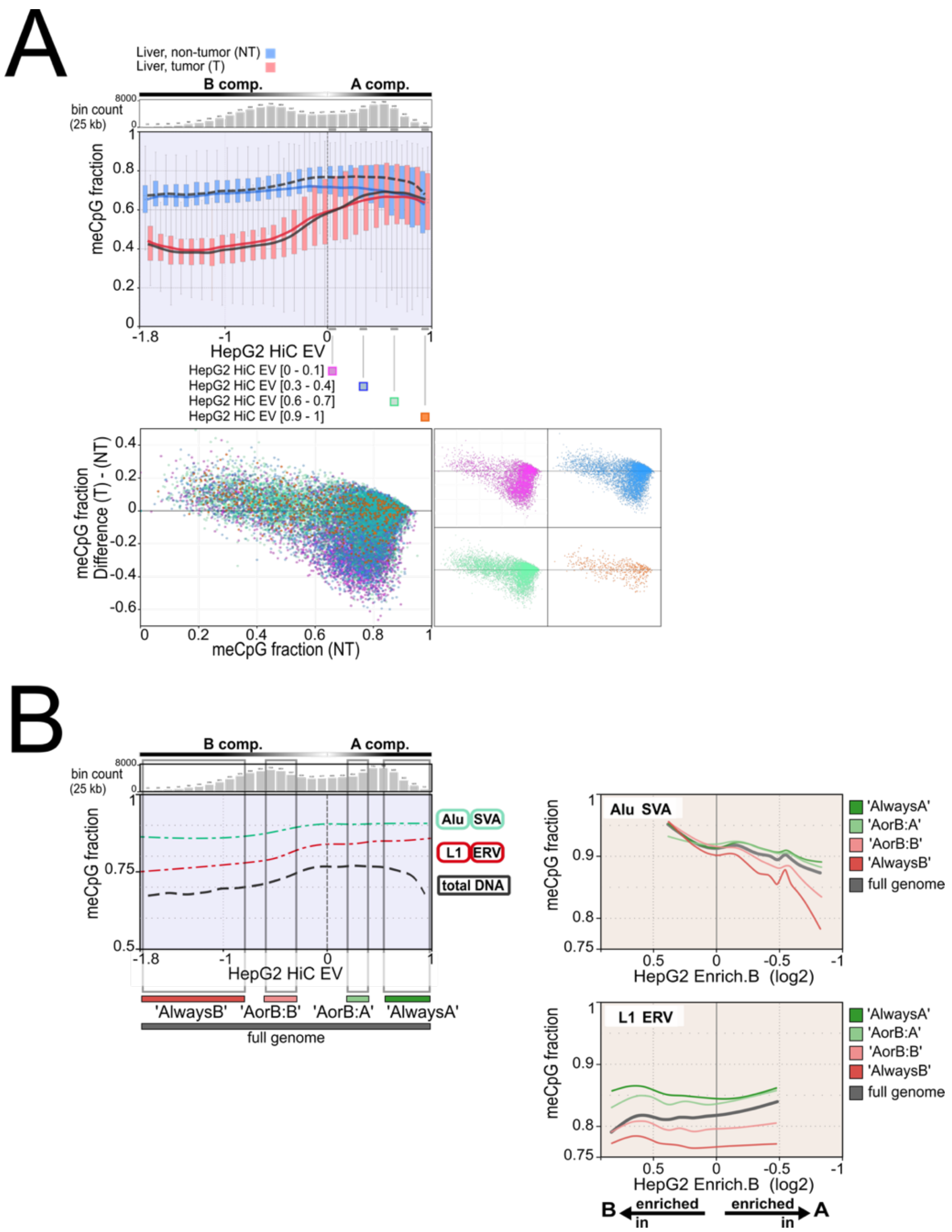

**Extended Data Fig. 15: meCpG fraction for total DNA, and for ProA and ProB RepSeqs.**

**(A) meCpG fraction in total DNA from tumor and non-tumor liver tissue.**
Upper panel: For the hg38 genome, 25 kb bins were grouped into intervals of width 0.1 based on their Hi-C EV values in HepG2 cells. For each interval, boxplots show the distribution of meCpG fractions in tumor (T) and non-tumor (NT) liver tissue. A curve connects the mean values (blue, NT; red, T) and the median values (grey dashed line, NT; grey solid line, T). The number of bins in each interval is shown as a bar plot.
Lower panel: Each bin (classified into four Hi-C EV intervals) is plotted according to its meCpG fraction in the NT sample (x-axis) and the NT-T difference (y-axis). Miniatures on the right display each interval separately.

**(B and C) meCpG fraction of ProA and ProB RepSeqs in normal liver, compared with total DNA.**
(B): Each TE copy in hg38 is plotted according to the Hi-C EV of its 25-kb bin (HepG2) and its meCpG fraction in normal liver. Only the regression curves are shown: Alu and SVA elements are grouped (green curve), and L1 and ERV elements are grouped (red curve). For reference, the dashed black line represents the meCpG fraction of total DNA in the NT sample (reproduced from panel A).
(C) Regression curves generated as in panel B, except that the x-axis now corresponds to the mean enrichment of the TE subfamily in the HepG2 B compartment. Curves are shown separately for four chromatin classes: AlwaysB (EV < -0.8), AorB:B (–0.6 < EV < -0.3), AorB:A (0.2 < EV < 0.4), and AlwaysA (EV > 0.55) (see Fig. 7E). Upper panel: Alu and SVA elements; lower panel: L1 and ERV elements. Curves shown here are identical to those in Fig. 7E (middle panels) but displayed in a different coordinate representation.

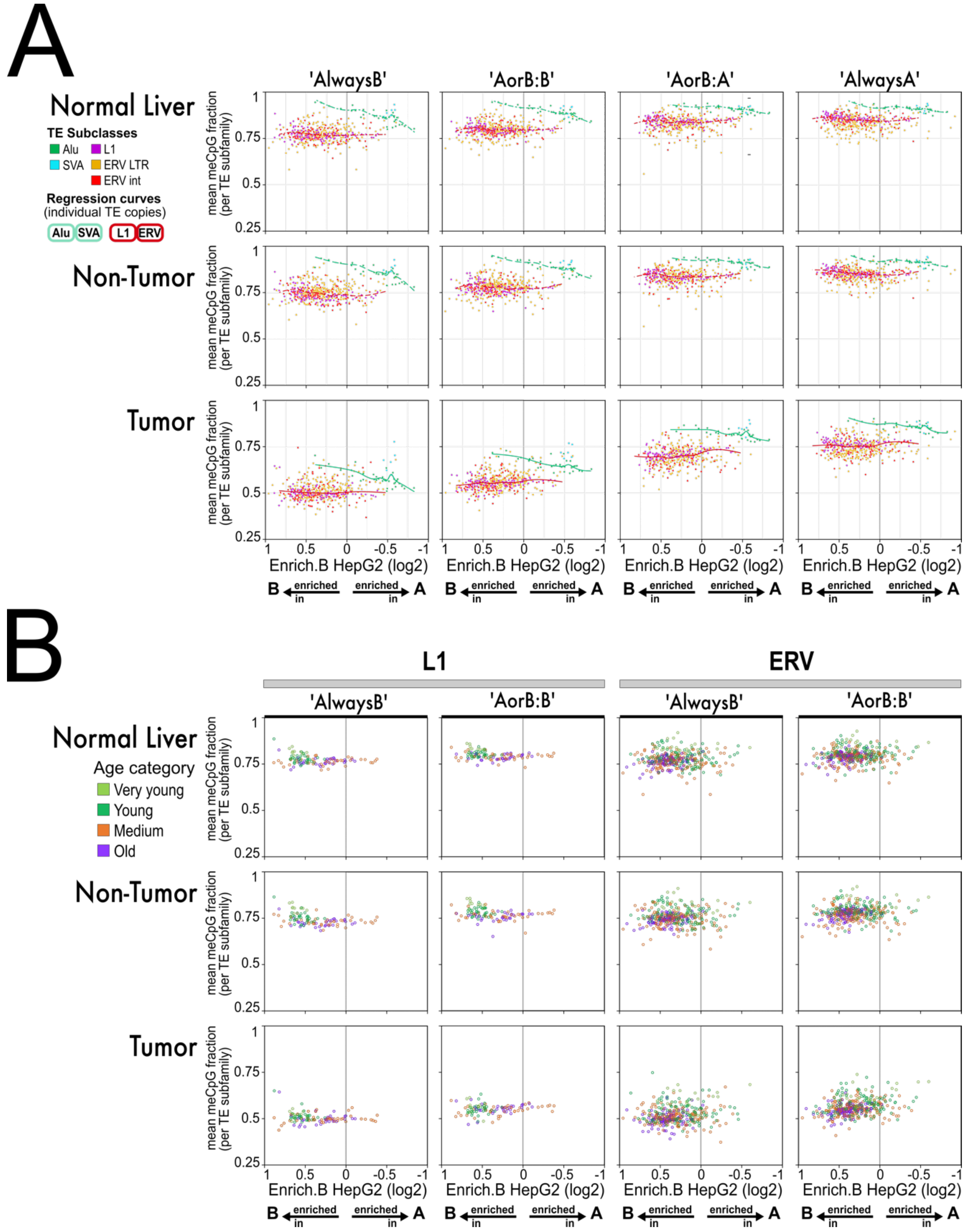
A
Normal Liver
TE Subclasses
Alu
L1
SVA
ERV LTR
ERV int
Regression curves
(individual TE copies)
Alu
SVA
L1
ERV
Non-Tumor
Tumor
'AlwaysB'
'AorB:B'
'AorB:A'
'AlwaysA'
mean meCpG fraction (per TE subfamily)
Enrich.B HepG2 (log2)
B enriched in
enriched in A
B
L1
ERV
Age category
Very young
Young
Medium
Old

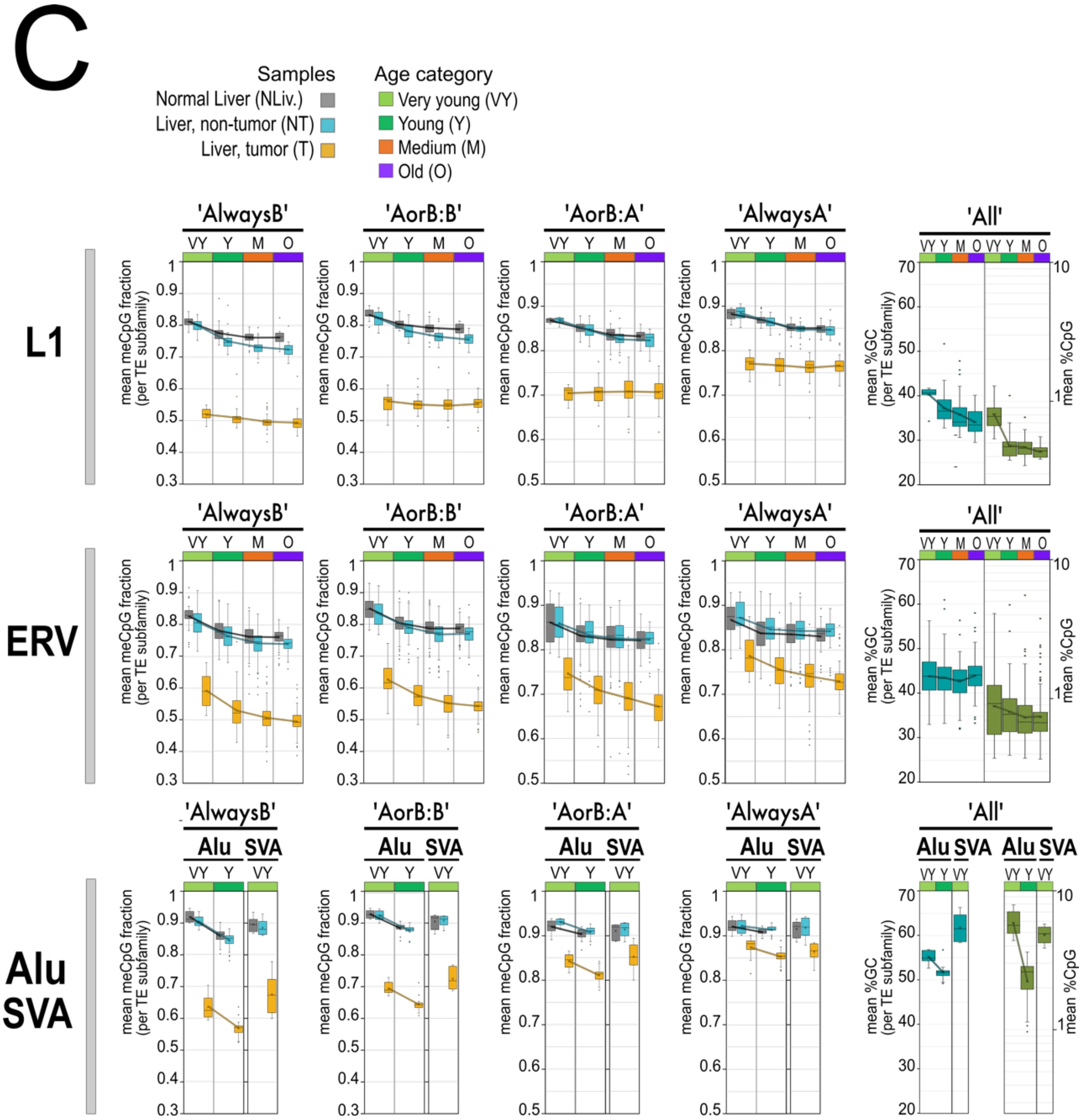

**Extended Data Fig. 16: Mean meCpG fraction per TE subfamily in normal and tumor tissue across chromatin classes.**

(A) Each TE subfamily from the Alu, SVA, L1, and ERV subclasses is represented as a point on a Euclidean plane according to its mean enrichment in the HepG2 B compartment (x-axis) and its mean meCpG fraction in tumor (T), non-tumor (NT), and normal liver (NLiv) tissue (y-axis), as indicated in the upper left. For ERVs, LTR sequences (“LTR”) and internal sequences (“int”) are shown separately. Note that enrichment in B is plotted in reverse order, such that subfamilies enriched in the A compartment appear to the right of the central 0 axis. Regression curves obtained from individual TE copies (Alu and SVA together; L1 and ERV together), as shown in Fig. 7E and Extended Data Fig. 15C, are overlaid for reference.
The analysis was performed separately for four chromatin classes defined by extreme or intermediate Hi-C EV values (see Fig. 7E; Extended Data Fig. 15): 'AlwaysB' (EV < -0.8), 'AorB:B' (-0.6 < EV < -0.3), 'AorB:A' (0.2 < EV < 0.4), and 'AlwaysA' (EV > 0.55). Only subfamilies with more than 30 genomic copies in the region considered were included.

(B) Each L1 and ERV subfamily is plotted according to its mean enrichment in the B compartment (x-axis) and its mean meCpG fraction in T, NT, and NLiv tissue (y-axis), as indicated on the left. Point colors reflect the median age of each subfamily (as in Fig. 3). Shown only for the 'AlwaysB' and 'AorB:B' genome portions. Other details follow those described for panel A.

(C) Boxplots showing, for each TE subfamily (L1, ERV, Alu, SVA), the mean meCpG fraction in T, NT, and NLiv tissue, together with mean %GC and mean %CpG, across age categories, for the same four chromatin classes as in panel A. ERV subfamilies (“LTR” and “int”) were combined. Only subfamilies with at least 30 copies per genome portion were included. %CpG, log10 scale.

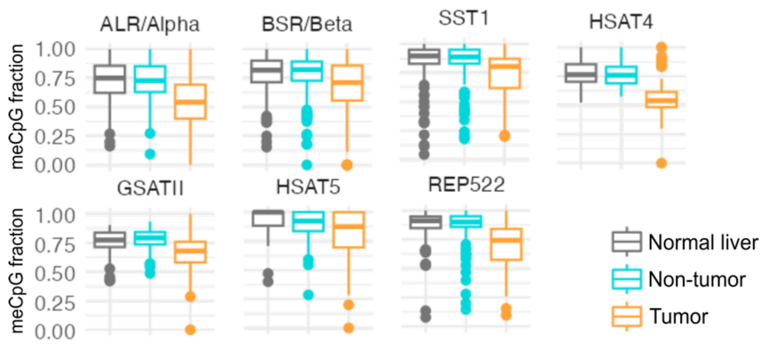

**Extended Data Fig. 17: (Peri)centromeric and (sub)telomeric satellites lose CpG methylation in cancer.**

Boxplot representation of the meCpG fraction for individual copies of the indicated satellite families, in tumor, non-tumor, and normal liver tissue, as in Fig. 7. REP522 is a subtelomeric satellite; all others correspond to centromeric or pericentromeric satellites. Only CorrB.enrichB satellite subfamilies with more than 50 copies in hg38 are shown.

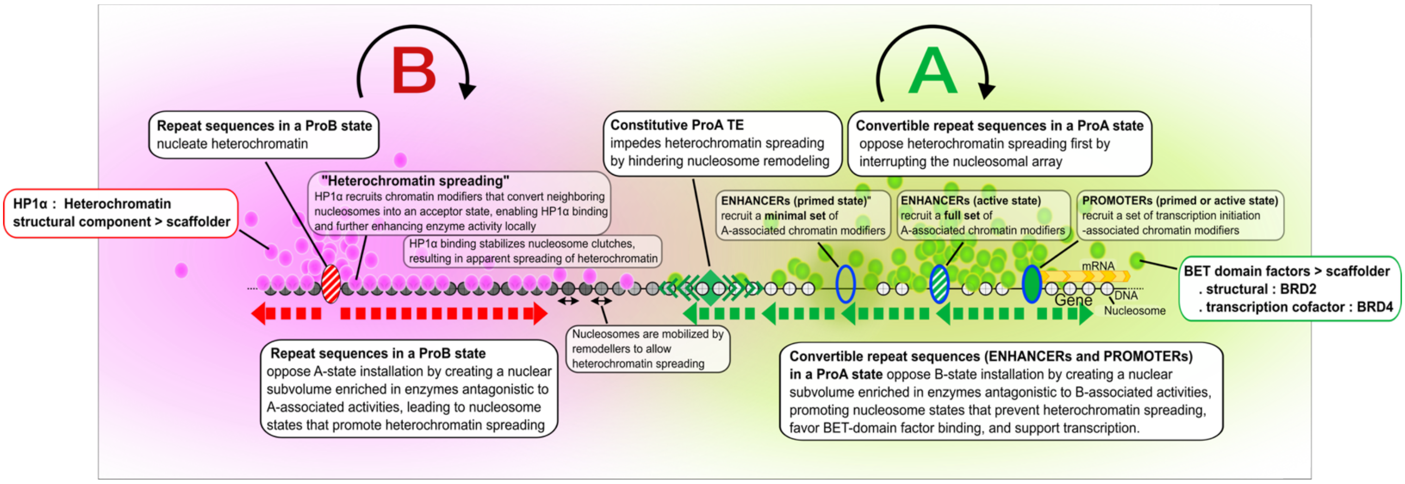

**Extended data Fig. 18: Molecular mechanisms by which ProA and ProB RepSeqs promote A-type and B-type chromatin: a molecular toggle switch.**

Schematic summary of the molecular mechanisms underlying ProA and ProB functions, presented as a complement to Fig. 8. Both A- and B-chromatin states are governed by similar general rules involving self-reinforcing interaction networks orchestrated by ProA and ProB elements, respectively, and actively oppose conversion to the alternate state, thereby forming a molecular toggle switch, as previously proposed and recapitulated in vitro [43-45]. The scheme is drawn as a one-dimensional representation of the chromatin fiber; the corresponding three-dimensional interpretation is explicitly described in the text below.

**Left: ProB-driven B-chromatin system**

ProB elements act as nucleation points for the assembly of dynamic heterochromatin condensates, from which heterochromatin appears to "spread" along the chromatin fiber.

When examined in 3D, this apparent heterochromatin "spreading" does not correspond to a single linear process, as schematically depicted, but instead reflects the combined effects of: (i) diffusion of heterochromatin factors away from the ProB element, together with stochastic contacts between the ProB element and distal loci along the same chromatin fiber - both processes being more efficient in the immediate vicinity of ProB elements [46,47]; and (ii) compaction and reduced dynamics of the chromatin fiber, arising from the stabilization of nucleosome clutches upon binding of heterochromatin structural factors, essentially HP1α and H1.0 [48].

Heterochromatin spreading requires active nucleosome mobilization by dedicated ATP-dependent remodeling complexes and a nucleosomal state (grey disks) permissive for both HP1α binding and nucleosome clutch formation. Such states are typically associated with low histone acetylation levels and the presence of the linker histone H1.0, and more generally depend on the activity of heterochromatin-enriched chromatin modifiers that deposit histone marks stabilizing HP1α binding - most notably H3K9me3 [49]. Conversely, acetylated nucleosomes (white disks; increasing whiteness indicates higher acetylation levels) impair clutch formation [44] and oppose heterochromatin spreading.

The coalescence model shown in Fig. 8 provides a conceptual representation of cooperativity among ProB elements within a chromatin domain: increasing ProB density results in more effective ProB function and more robust spreading, owing to functional interactions occurring in 3D. In practice, coalescence is thought to occur primarily between nucleosome clutches stabilized by HP1α, rather than between individual ProB elements. Because nucleosome clutches display a fractal-like organization involving nucleosome associations across multiple length scales [50-52], any mechanism that stabilizes such associations beyond a critical threshold is expected to enhance chromatin cohesion in a non-linear manner. Consistently, nucleosome clutches within heterochromatin-dominated chromatin nanodomains appear markedly over-stabilized by heterochromatin structural factors [48], providing a physical basis for synergistic cooperativity among ProB elements.
ProB elements distributed across a domain behave collectively as a higher-order ProB structure, establishing a potent feed-forward loop: they increase the local probability of presence of heterochromatin factors within a nuclear subvolume ("sphere of influence", pink shading), thereby further reinforcing ProB activity. HP1α, in turn, anchors this sphere of influence to the chromatin fiber.

Altogether, the B-chromatin system promotes the binding of heterochromatin components to the chromatin fiber and nucleosome-clutch stabilization, through interwoven positive feed-forward loops (circular arrow), thereby installing a chromatin environment intrinsically unfavorable to gene expression.

**Right: ProA-driven A-chromatin system**
ProA elements antagonize heterochromatin spreading by altering the composition of the chromatin fiber and its local molecular environment. Alu repeats and DHSs impede nucleosome translational sliding, albeit through distinct mechanisms. DHSs introduce a discontinuity in the nucleosomal array via nucleosome eviction or sliding by chromatin remodelers, as recruited by TFs (blue oval), whereas Alu elements (green diamond) locally constrain nucleosome mobility through the presence of one or two strongly positioned nucleosomes.

Gene regulatory elements (enhancers and promoters), which are largely derived from RepSeqs, are DHSs at which TFs recruit transcription-associated cofactors (green shading). The A-chromatin state associated with these ProA elements is stabilized by multiple interwoven feed-forward loops (circular arrow) [43,53,54]. Many transcription cofactors further counteract core B-system mechanisms. In the active enhancer state, as compared with the primed state, local concentrations of these factors are higher, thereby further destabilizing chromatin-fiber configurations compatible with heterochromatin assembly.
In particular, acetylation, phosphorylation, and PARP1-dependent PARylation of both core and linker histones (white disks) impair nucleosome-clutch formation, and disrupt HP1α binding and chromatin fiber bridging [44,54-58]. HP1α may itself be acetylated by enhancer-associated histone acetyltransferases (HATs), curtailing its ability to bind to chromatin in an A environment [59,60].
In parallel, BET-domain proteins (e.g. BRD2, BRD4) (green spheres), which bind acetylated histones, play a functional role that is in several respects a mirror image of HP1α, echoing the early-described functional opposition between yeast Bdf1 and Sir3 [61]. Notably, BRD2 binding requires H4K16 acetylation, a conserved hallmark of the A compartment [62,63]. Through its condensate-forming properties, BRD2 can provide local cohesion within the A compartment [64], and further recruit chromatin remodelers that also recognize acetylated histones, stabilizing the A state [65]. Importantly, however, BET-domain factors, and more generally A-associated chromatin factors, exhibit limited spreading along chromatin away from active promoters and enhancers [66], although some, such as BRD4, can recruit HATs, providing a potential read-write mechanism [67]. This likely reflects the fact that the bulk of HAT activity is tightly confined to active enhancers and promoters, where HAT function requires allosteric activation through interactions with multiple TFs [66,68,69]. Another plausible explanation is that in differentiated cells, A-associated chromatin factors are far less abundant than HP1α -heterochromatin associated components [70]. Accordingly, the A-chromatin system can be viewed as a regulatory layer superimposed on the B-chromatin system, which is otherwise established by default.

Altogether, the A-chromatin state is stabilized by feed-forward loops (circular arrow), and A domains emerge by preventing the physical conditions required for heterochromatin spreading, thereby establishing a chromatin environment favorable to gene expression.

### REFERENCES

1. Haberle, V. *et al.* Transcriptional cofactors display specificity for distinct types of core promoters. *Nature* **570**, 122-126 (2019).
2. Lee, J.Y. *et al.* Conserved dual-mode gene regulation programs in higher eukaryotes. *Nucleic Acids Res* **49**, 2583-2597 (2021).
3. Dennis, M.Y. & Eichler, E.E. Human adaptation and evolution by segmental duplication. *Curr Opin Genet Dev* **41**, 44-52 (2016).
4. Costello, K.R. *et al.* Sequence features of retrotransposons allow for epigenetic variability. *Elife* **10**(2021).
5. Guo, H. *et al.* DNA hypomethylation silences anti-tumor immune genes in early prostate cancer and CTCs. *Cell* **186**, 2765-2782 e28 (2023).
6. Luqman-Fatah, A. *et al.* The interferon stimulated gene-encoded protein HELZ2 inhibits human LINE-1 retrotransposition and LINE-1 RNA-mediated type I interferon induction. *Nat Commun* **14**, 203 (2023).
7. Ormundo, L.F., Machado, C.F., Sakamoto, E.D., Simoes, V. & Armelin-Correa, L. LINE-1 specific nuclear organization in mice olfactory sensory neurons. *Mol Cell Neurosci* **105**, 103494 (2020).
8. Fourel, G., Lebrun, E. & Gilson, E. Protosilencers as building blocks for heterochromatin. *Bioessays* **24**, 828-35 (2002).
9. Pourmorady, A. & Lomvardas, S. Olfactory receptor choice: a case study for gene regulation in a multi-enhancer system. *Curr Opin Genet Dev* **72**, 101-109 (2022).
10. Rey-Millet, M. *et al.* Senescence-associated transcriptional derepression in subtelomeres is determined in a chromosome-end-specific manner. *Aging Cell* **22**, e13804 (2023).
11. Elgin, S.C. & Reuter, G. Position-effect variegation, heterochromatin formation, and gene silencing in Drosophila. *Cold Spring Harb Perspect Biol* **5**, a017780 (2013).
12. Meir, Z., Mukamel, Z., Chomsky, E., Lifshitz, A. & Tanay, A. Single-cell analysis of clonal maintenance of transcriptional and epigenetic states in cancer cells. *Nat Genet* **52**, 709-718 (2020).
13. Owen, B.M. & Davidovich, C. DNA binding by polycomb-group proteins: searching for the link to CpG islands. *Nucleic Acids Res* **50**, 4813-4839 (2022).
14. Weber, C.M. *et al.* mSWI/SNF promotes Polycomb repression both directly and through genome-wide redistribution. *Nat Struct Mol Biol* **28**, 501-511 (2021).
15. de Tribolet-Hardy, J. *et al.* Genetic features and genomic targets of human KRAB-zinc finger proteins. *Genome Res* (2023).
16. Griffin, G.K. *et al.* Epigenetic silencing by SETDB1 suppresses tumour intrinsic immunogenicity. *Nature* **595**, 309-314 (2021).
17. Valle-Garcia, D. *et al.* ATRX binds to atypical chromatin domains at the 3' exons of zinc finger genes to preserve H3K9me3 enrichment. *Epigenetics* **11**, 398-414 (2016).
18. Carraro, M. *et al.* DAXX adds a de novo H3.3K9me3 deposition pathway to the histone chaperone network. *Mol Cell* **83**, 1075-1092 e9 (2023).
19. Begnis, M. *et al.* Clusters of lineage-specific genes are anchored by ZNF274 in repressive perinucleolar compartments. *bioRxiv*, 2024.01.04.574183 (2024).
20. Pontis, J. *et al.* Primate-specific transposable elements shape transcriptional networks during human development. *Nat Commun* **13**, 7178 (2022).
21. Ito, J. *et al.* Endogenous retroviruses drive KRAB zinc-finger protein family expression for tumor suppression. *Sci Adv* **6**(2020).
22. Riddle, N.C. *et al.* Enrichment of HP1a on Drosophila chromosome 4 genes creates an alternate chromatin structure critical for regulation in this heterochromatic domain. *PLoS Genet* **8**, e1002954 (2012).
23. Riddle, N.C. & Elgin, S.C.R. The Drosophila Dot Chromosome: Where Genes Flourish Amidst Repeats. *Genetics* **210**, 757-772 (2018).
24. Tyssowski, K.M. *et al.* Different Neuronal Activity Patterns Induce Different Gene Expression Programs. *Neuron* **98**, 530-546 e11 (2018).
25. Sabari, B.R. *et al.* Coactivator condensation at super-enhancers links phase separation and gene control. *Science* **361**(2018).
26. Atsumi, Y. *et al.* Repetitive CREB-DNA interactions at gene loci predetermined by CBP induce activity-dependent gene expression in human cortical neurons. *Cell Rep* **43**, 113576 (2024).
27. Fourel, G., Miyake, T., Defossez, P.A., Li, R. & Gilson, E. General regulatory factors (GRFs) as genome partitioners. *J Biol Chem* **277**, 41736-43 (2002).
28. Ferrer, J. & Dimitrova, N. Transcription regulation by long non-coding RNAs: mechanisms and disease relevance. *Nat Rev Mol Cell Biol* **25**, 396-415 (2024).

29. Chakraborty, S. *et al.* Enhancer-promoter interactions can bypass CTCF-mediated boundaries and contribute to phenotypic robustness. *Nat Genet* **55**, 280-290 (2023).
30. Du, M. *et al.* Direct observation of a condensate effect on super-enhancer controlled gene bursting. *Cell* **187**, 2595-2598 (2024).
31. Vassetzky, N.S., Ten, O.A. & Kramerov, D.A. B1 and related SINEs in mammalian genomes. *Gene* **319**, 149-60 (2003).
32. Richard Albert, J. *et al.* DNA methylation shapes the Polycomb landscape during the exit from naive pluripotency. *Nat Struct Mol Biol* (2024).
33. Abatti, L.E. *et al.* Epigenetic reprogramming of a distal developmental enhancer cluster drives SOX2 overexpression in breast and lung adenocarcinoma. *Nucleic Acids Res* **51**, 10109-10131 (2023).
34. Ordonez, R. *et al.* Genomic context sensitizes regulatory elements to genetic disruption. *Mol Cell* **84**, 1842-1854 e7 (2024).
35. Ohtani, H., Liu, M., Zhou, W., Liang, G. & Jones, P.A. Switching roles for DNA and histone methylation depend on evolutionary ages of human endogenous retroviruses. *Genome Res* **28**, 1147-1157 (2018).
36. Leung, A. *et al.* LTRs activated by Epstein-Barr virus-induced transformation of B cells alter the transcriptome. *Genome Res* **28**, 1791-1798 (2018).
37. Schmidt, N. *et al.* An influenza virus-triggered SUMO switch orchestrates co-opted endogenous retroviruses to stimulate host antiviral immunity. *Proc Natl Acad Sci U S A* **116**, 17399-17408 (2019).
38. Zeller, P. *et al.* Single-cell sortChIC identifies hierarchical chromatin dynamics during hematopoiesis. *Nat Genet* **55**, 333-345 (2023).
39. Maeda, R.K. & Karch, F. The open for business model of the bithorax complex in Drosophila. *Chromosoma* **124**, 293-307 (2015).
40. Deschamps, J. & Duboule, D. Embryonic timing, axial stem cells, chromatin dynamics, and the Hox clock. *Genes Dev* **31**, 1406-1416 (2017).
41. Dekker, J. & Mirny, L.A. The chromosome folding problem and how cells solve it. *Cell* **187**, 6424-6450 (2024).
42. Terekhanova, N.V. *et al.* Epigenetic regulation during cancer transitions across 11 tumour types. *Nature* **623**, 432-441 (2023).
43. Fourel, G., Magdinier, F. & Gilson, E. Insulator dynamics and the setting of chromatin domains. *Bioessays* **26**, 523-32 (2004).
44. Gibson, B.A. *et al.* Organization of Chromatin by Intrinsic and Regulated Phase Separation. *Cell* **179**, 470-484 e21 (2019).
45. Hsieh, L.J., Lou, T., Gourdet, M.A., Wong, E. & Narlikar, G.J. A Biochemical Screening Platform to Target Chromatin States Using Condensates as a Tool. *SLAS Discov*, 100236 (2025).
46. Lebrun, E., Fourel, G., Defossez, P.A. & Gilson, E. A methyltransferase targeting assay reveals silencer-telomere interactions in budding yeast. *Mol Cell Biol* **23**, 1498-508 (2003).
47. Lee, Y.C.G. *et al.* Pericentromeric heterochromatin is hierarchically organized and spatially contacts H3K9me2 islands in euchromatin. *PLoS Genet* **16**, e1008673 (2020).
48. Minami, K. *et al.* Replication-dependent histone labeling dissects the physical properties of euchromatin/heterochromatin in living human cells. *Sci Adv* **11**, eadu8400 (2025).
49. Allshire, R.C. & Madhani, H.D. Ten principles of heterochromatin formation and function. *Nat Rev Mol Cell Biol* **19**, 229-244 (2018).
50. Socol, M. *et al.* Rouse model with transient intramolecular contacts on a timescale of seconds recapitulates folding and fluctuation of yeast chromosomes. *Nucleic Acids Res* **47**, 6195-6207 (2019).
51. Barth, R., Bystricky, K. & Shaban, H.A. Coupling chromatin structure and dynamics by live super-resolution imaging. *Sci Adv* **6**(2020).
52. Almassalha, L.M. *et al.* Chromatin conformation, gene transcription, and nucleosome remodeling as an emergent system. *Sci Adv* **11**, eadq6652 (2025).
53. Rada-Iglesias, A., Grosveld, F.G. & Papantonis, A. Forces driving the three-dimensional folding of eukaryotic genomes. *Mol Syst Biol* **14**, e8214 (2018).
54. Karr, J.P., Ferrie, J.J., Tjian, R. & Darzacq, X. The transcription factor activity gradient (TAG) model: contemplating a contact-independent mechanism for enhancer-promoter communication. *Genes Dev* **36**, 7-16 (2022).
55. Azad, G.K. *et al.* PARP1-dependent eviction of the linker histone H1 mediates immediate early gene expression during neuronal activation. *J Cell Biol* **217**, 473-481 (2018).
56. Nosella, M.L. *et al.* Poly(ADP-ribosyl)ation enhances nucleosome dynamics and organizes DNA damage repair components within biomolecular condensates. *Mol Cell* **84**, 429-446 e17 (2024).
57. Benabdallah, N.S. *et al.* Decreased Enhancer-Promoter Proximity Accompanying Enhancer Activation. *Mol Cell* **76**, 473-484 e7 (2019).

58. Kunowska, N., Rotival, M., Yu, L., Choudhary, J. & Dillon, N. Identification of protein complexes that bind to histone H3 combinatorial modifications using super-SILAC and weighted correlation network analysis. *Nucleic Acids Res* **43**, 1418-32 (2015).
59. LeRoy, G. *et al.* Heterochromatin protein 1 is extensively decorated with histone code-like post-translational modifications. *Mol Cell Proteomics* **8**, 2432-42 (2009).
60. Ukmar-Godec, T. *et al.* Multimodal interactions drive chromatin phase separation and compaction. *Proc Natl Acad Sci U S A* **120**, e2308858120 (2023).
61. Ladurner, A.G., Inouye, C., Jain, R. & Tjian, R. Bromodomains mediate an acetyl-histone encoded antisilencing function at heterochromatin boundaries. *Mol Cell* **11**, 365-76 (2003).
62. Zheng, B. *et al.* BRD2 bridges TFIID and histone acetylation to promote transcriptional initiation. *bioRxiv*, 2025.09.20.676678 (2025).
63. Radzisheuskaya, A. *et al.* Complex-dependent histone acetyltransferase activity of KAT8 determines its role in transcription and cellular homeostasis. *Mol Cell* **81**, 1749-1765 e8 (2021).
64. Xie, L. *et al.* BRD2 compartmentalizes the accessible genome. *Nat Genet* **54**, 481-491 (2022).
65. Zhou, M.M. & Cole, P.A. Targeting lysine acetylation readers and writers. *Nat Rev Drug Discov* **24**, 112-133 (2025).
66. Crump, N.T. *et al.* BET inhibition disrupts transcription but retains enhancer-promoter contact. *Nat Commun* **12**, 223 (2021).
67. Wu, T., Kamikawa, Y.F. & Donohoe, M.E. Brd4's Bromodomains Mediate Histone H3 Acetylation and Chromatin Remodeling in Pluripotent Cells through P300 and Brg1. *Cell Rep* **25**, 1756-1771 (2018).
68. Ortega, E. *et al.* Transcription factor dimerization activates the p300 acetyltransferase. *Nature* **562**, 538-544 (2018).
69. Ferrie, J.J. *et al.* p300 is an obligate integrator of combinatorial transcription factor inputs. *Mol Cell* (2023).
70. Gillespie, M.A. *et al.* Absolute Quantification of Transcription Factors Reveals Principles of Gene Regulation in Erythropoiesis. *Mol Cell* **78**, 960-974 e11 (2020).
